# ClusTRace, a bioinformatic pipeline for analyzing clusters in virus phylogenies

**DOI:** 10.1101/2021.12.09.471941

**Authors:** Ilya Plyusnin, Phuoc Thien Truong Nguyen, Tarja Sironen, Olli Vapalahti, Teemu Smura, Ravi Kant

**Affiliations:** Department of Veterinary Bioscience, University of Helsinki, 00014 Helsinki, Finland; Department of Virology, University of Helsinki, 00014 Helsinki, Finland; Department of Virology and Immunology, University of Helsinki and Helsinki University Hospital, 00014 Helsinki, Finland

## Abstract

**Summary:** SARS-CoV-2 is the highly transmissible etiologic agent of coronavirus disease 2019 (COVID-19) and has become a global scientific and public health challenge since December 2019. Several new variants of SARS-CoV-2 have emerged globally raising concern about prevention and treatment of COVID-19. Early detection and in depth analysis of the emerging variants allowing pre-emptive alert and mitigation efforts are thus of paramount importance.

Here we present ClusTRace, a novel bioinformatic pipeline for a fast and scalable analysis of sequence clusters or clades in large viral phylogenies. ClusTRace offers several high level functionalities including outlier filtering, aligning, phylogenetic tree reconstruction, cluster or clade extraction, variant calling, visualization and reporting. ClusTRace was developed as an aid for COVID-19 transmission chain tracing in Finland and the main emphasis has been on fast and unsupervised screening of phylogenies for markers of super-spreading events and other features of concern, such as high rates of cluster growth and/or accumulation of novel mutations.

**Availability:** All code is freely available from https://bitbucket.org/plyusnin/clustrace/

## 1 INTRODUCTION

Emerging pathogens are a constant threat to mankind, as shown by the Ebola (Dixon *et al*., 2014) and Zika (Kindhauser *et al*., 2016) virus outbreaks in 2014 and 2015 respectively, and the ongoing severe acute respiratory syndrome coronavirus 2 (SARS-CoV-2) pandemic. These viruses are of zoonotic origin, like the majority of emerging pathogens (Woolhouse and Gowtage-Sequeria). Environmental changes, new forms of land use, increasing human and production animal densities, trans-boundary movements, travel and globalization have dramatically enhanced the rate of zoonotic disease emergence (Morens and Fauci, 2020). All viruses, including SARS-CoV-2, change over time, however most changes have negligible effect on the virus’s phenotype. However, some mutations may affect key pathogenic properties such as transmissibility and disease severity, or the performance of vaccines, therapeutic agents or diagnostic tools. SARS-CoV-2 is the causative agent of coronavirus disease 2019 (COVID-19) (Wu et al., 2020). The SARS-CoV-2 pandemic has already infected more than 262 million people in 224 countries, causing over 5 million deaths globally as of 1st of December 2021 (https://www.worldometers.info/coronavirus/).

SARS-CoV-2 is a global challenge, which is further complicated by the continuous emergence of new Variants of Concern (VOCs) or Variants of Interest (VOI). Variants that have carried VOC status include Alpha (B.1.1.7) (Wise, 2020), Beta (B1.351) (Tegally *et al*., 2020), Gamma (P.1), Delta (B.1.617.2) (Kirola, 2021) and, as of writing this, we are experiencing early spread of Omicron (Latif *et al*., 2021b). These VOCs pose an increased public health risk due to having one or more of the following characteristics: higher transmissibility (Campbell *et al*., 2021), immune escape properties for antibodies from previous infection (Virtanen *et al*., 2021), lower response towards current vaccines compared to the original wild type strains these vaccines were based on (Jalkanen *et al*., 2021). Detecting and monitoring these novel variants is essential in SARS-CoV-2 surveillance.

COVID-19 is the first pandemic with the pathogen being under surveillance using full genome sequencing on a global scale and over an extensive time period. Surveillance of the pandemic creates demand for fast and scalable sequencing, genome assembling, viral strain assignment, phylogenetic analysis, variant calling and molecular epidemiology to inform contact tracing and non-pharmaceutical interventions. Although bioinformatics offers an abundance of methods and tools for sequence analysis, their employment in virology and epidemiology can be hindered by the developer-user gap between bioinformatics and other fields (Mangul *et al*., 2019). This gap can be bridged by pipelines tailored specifically for the analysis of viral sequences and equipped with intuitive interface and output reporting. A number of bioinformatic software packages are already available to help with detection, tracking and tracing of VOCs e.g. Pangolin (O’Toole *et al*., 2021), Nextstrain (Hadfield *et al*., 2018), Nextclade (Aksamentov *et al*., 2021), Jovian (Zwagemaker *et al*., 2021) and HaVoC (Nguyen *et al*., 2021). At the same time co-infections can be tracked with meta-genomic approaches like Lazypipe (Plyusnin *et al*., 2020). Such tools are certainly helping the global effort for COVID-19 surveillance but they are not void of limitations. Tools like Pangolin and Nextclade are primarily designed for tracking large accumulations of mutation events that are rare and may be preceded by the less visible sub-lineage genetic changes. Nextstrain offers a comprehensive analysis, but is heavily dependent on sequence metadata, dataset pre-filtering and time-consuming manual inspection of the produced interactive reports. Here we introduce ClusTRace (https://www2.helsinki.fi/en/projects/clustrace), a novel bioinformatic pipeline for Unix/Linux environments that complements the existing toolkits with unsupervised clade or cluster analysis, intuitive visualizations and reporting. ClusTRace can help with surveillance of the current ongoing COVID-19 pandemic and for any upcoming future epidemic or pandemic.

## 2 MATERIAL AND METHODS

ClusTRace is a bioinformatic software package implemented primarily in perl. ClusTRace supports several tasks that can be executed one by one or combined into pipelines (Figure 1).

**Figure 1.**
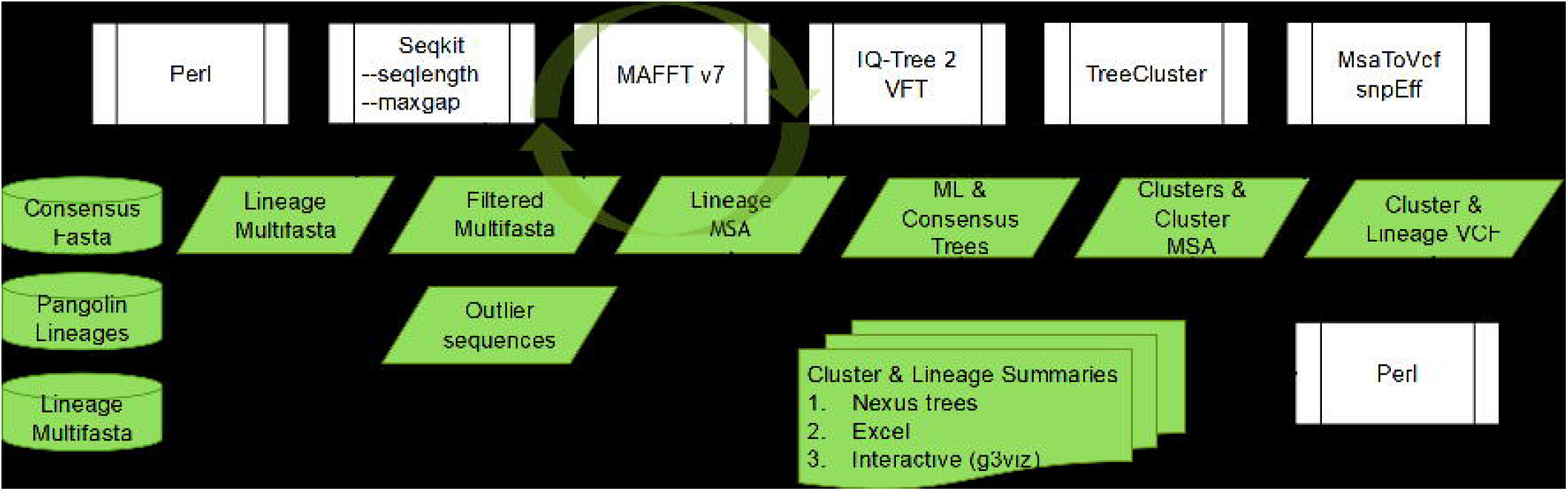
ClusTRace flowchart

### 2.1 Basic workflow

The analysis starts with consensus SARS-CoV-2 genomic sequences output by a given sequencing platform (e.g., Illumina). In the first step, ClusTRace assigns genomic sequences to a dynamic Pango lineage classification with Pangolin (O’Toole *et al*., 2021). Then, ClusTRace sorts sequences into multi-fasta files according to the assigned lineage file. Although we use Pangolin as the default lineage assigner, classification file can be produced with any method preferred by the user (the pipeline will accept any csv-file that includes a named column labelled as “lineage”).

Multi-fasta files are then pruned from outliers with the help of SeqKit (Shen *et al*., 2016). By default, we remove all sequences that deviate more than 10% from the median length of the sequence set or that have more than 10% gaps (these parameters can be modified on the command line with *--minlen, -- maxlen* and *--maxgap*).

In the next step, filtered sequence sets for each lineage are aligned with MAFFT v7 (Katoh and Standley, 2013). Multiple sequence alignments (MSAs) are then used to construct phylogenetic trees with IQ-TREE 2 (Minh *et al*., 2020). IQ-TREE 2 supports a wide range of substitution models and will, by default, use ModelFinder to determine the best-fit model (Minh *et al*., 2020). The user can choose to create bootstrapped consensus trees with IQ-TREE 2 Ultra-Fast Bootstrapping (ClusTRace *--ufboot* option). For very large sequence sets, the user can choose to run VeryFastTree (Piñeiro *et al*., 2020) with GTR model (ClusTRace --*tree vftree* option).

In the next step, sequence clusters are extracted with TreeCluster (Balaban *et al*., 2019). Clusters are extracted with MaxClade-method at several pairwise distance cut-offs. We use two cut-off thresholds that are scaled to the size of SARS-CoV-19 genome (∼30,000 nt) and roughly correspond to twenty and thirty mutations between pairs of sequences. Next, ClusTRace creates custom nexus trees in which sequences are assigned labels and colours according to the assigned cluster.

ClusTRace can read date annotations from sequence ids and will accept common date formats (e.g. “|YYYY-MM-DD|”). For date annotated sequences ClusTRace will trace the speed of growth for the extracted clusters. This is done by assigning sequences to time periods (calendar months or weeks) and by tracing the number of sequences that are assigned to each cluster and that are dated up to the given time period. For each lineage ClusTRace will print a separate cluster summary file with detailed information on the extracted clusters. These excel summaries include *clustSeqN, clustSeqId* and *clustGR* data sheets. The first and second data sheets report the number and ids of sequences in each cluster for each time period, while the third reports cluster size, median and maximal growth rates, and support value for the corresponding sub-phylogeny for each cluster. Separate *clustGR* data sheets are printed for each cluster cut-off threshold (by default twenty and thirty).

In the last step, ClusTRace extracts MSA(s) and runs variant calling for the extracted clusters. Nucleotide mutations are called from these against a reference genome with MsaToVcf (Page *et al*., 2016). Nucleotide variants are filtered to exclude 100 nt from the start and the end of the genome, as well as any regions that have over 30nt continuous stretches of below 75% coverage (these are assumed to represent sequencing errors). We also exclude variants with support below 50%. These filtering options are specified in the pipeline options and can be modified. Amino acid (aa) variants are called with snpEff (Cingolani *et al*., 2012). Finally, aa variants in all clusters are parsed and added to the cluster Excel summaries as *clustMutations* and *clustMutationTable* data sheets. The *clustMutations* sheet reports nt and aa mutations for each cluster, reference aa mutations and non-reference aa mutations. Reporting reference and non-reference mutations requires supplying reference mutations in a separate file. For genes of interest non-reference mutations can be reported separately (current version reports mutations for the S-gene). The *clustMutationTable* sheet reports aa mutations for the fastest growing clusters in a binary matrix. The top row lists aa mutations in genomic order with non-reference mutations highlighted in bold.

ClusTRace also supports extracting nt and aa, reference aa and non-reference aa mutations for lineage MSA(s) or for any other collection of MSA(s). Lineage mutations are reported with excel summary tables similar to the cluster mutation summaries.

ClusTRace also offers an interface to g3viz R library (Guo *et al*., 2020). Using this interface, the user can generate in R environment interactive mutation plots for both cluster and lineage vcf-files. These interactive plots can be saved in the form of simple html files to complement excel reports.

### 2.2 Adding new sequences to your analysis

ClusTRace supports updating your analysis with new sequence batches. Using this option, the user can update MSA(s) and tree(s) with novel data without repeating the whole analysis and by keeping exactly the same clusters for the updated dataset. This reduces subsequent run times and allows for more rapid analysis with up-to-date data. In this mode sequences are added to reference MSA(s) and new trees are restricted with reference topologies. The same set of clusters are identified in the updated trees by locating the last common ancestor (LCA) for old sequences and by assigning new sequences to the LCA with the shortest path in the tree. Due to topology changes, some clusters may shift to include other clusters; these are marked in the summary tables to avoid overestimation of the growth rates.

### 2.3 Selecting outgroup sequences

ClusTRace uses outgroup sequences to reroot phylogenetic trees. Several options are offered for the user. The default option is to use COVID-19 reference genome (NC_045512.2), however the user can also specify a given genomic sequence or a collection of sequences with options --*outgroup file* and -- *outgroup directory*. If *--outgroup* is a directory, the pipeline will look for an outgroup fasta file for each lineage. This can be an exact match (e.g. B.1.1.7.fasta) or a less specific match (e.g. B.1.fasta for B.1.1.7 lineage). Outgroup sequence will be added to all resulting MSA(s) and all phylogenetic trees will be rerooted using the outgroup.

## 3 RESULTS

We used ClusTRace to analyze a dataset of COVID-19 full genome sequences from patient samples collected in Finland from January to the end of May 2021. We started by running ClusTRace Pangolin mapping, multi-fasta construction and outlier filtering. In the downstream analysis, we focused on the dominant variants of concern, which for this dataset were the Alpha VOC with 5,379 sequences and Beta VOC with 1,051 sequences (GISAID accessions are listed in STable 1). We then run ClusTRace alignment, phylogeny with ultrafast bootstrapping (*--ufboot* option), default clustering and variant calling for these two lineages. As our outgroup sequences we used *EPI_ISL_601443* for Alpha and *EPI_ISL_660190* for Beta. All files output by ClusTRace for this analysis are available in SFile-1.

**Table 1.** Growth rates for ten Alpha clusters with maximal monthly growth rates.

| Cluster | Time Period |  |  |  |  |  | medGR | maxGR | maxGR % |
| --- | --- | --- | --- | --- | --- | --- | --- | --- | --- |
|  | 2020-12 | 2021-01 | 2021-02 | 2021-03 | 2021-04 | 2021-05 |  |  |  |
| Clust 68 | 0 | 69 | **379** | 462 | 479 | 479 | 69 | 310 | 64.7% |
| Clust 81 | 0 | 19 | **219** | 389 | 440 | 449 | 51 | 200 | 44.5% |
| Clust 70 | 0 | 9 | **188** | 278 | 315 | 324 | 37 | 179 | 55.2% |
| Clust 80 | 0 | 50 | 158 | **337** | 385 | 388 | 50 | 179 | 46.1% |
| Clust 56 | 0 | 0 | 47 | **224** | 302 | 314 | 47 | 177 | 56.4% |
| Clust 24 | 0 | 9 | 67 | **238** | 392 | 402 | 58 | 171 | 42.5% |
| Clust 22 | 0 | 9 | 104 | **246** | 277 | 283 | 31 | 142 | 50.2% |
| Clust 38 | 1 | 26 | **139** | 223 | 228 | 230 | 25 | 113 | 49.1% |
| Clust 74 | 2 | 39 | **125** | 169 | 177 | 177 | 37 | 86 | 48.6% |
| Clust 15 | 0 | 1 | 3 | 22 | **96** | 100 | 4 | 74 | 74.0% |
| Singletons | 2 | 8 | 21 | 37 | 42 | 43 |  |  |  |
| Total | 39 | 480 | 2316 | **4320** | 5241 | 5380 | 921 | 2004 | 37.2% |

Cluster, cluster id, Time Period, time period by month, medGR, median absolute growth rate per month, maxGR, maximal absolute growth rate per month, maxGR%, maximal relative growth rate per month. Values in bold indicate the month of the maximal absolute growth rate.

To get a quick summary on the lineage mutations, we start with g3viz visualisation (Figure 2). For Alpha we see that most high frequency aa mutations follow the GISAID reference (Figure 2.A). These include T1001I, A1708D, I2230T, 3675_3677del and P4715L in *ORF1ab*, 69_70del, N501Y, A570D, D614G, P681H, T716I, S982A and D1118H in *S*, D3L and S235F in *N*. For Alpha, there are just five aa variants specific for Finnish data with frequency 10% or higher: K5784R and E6272G in *ORF1ab*, N119H in *ORF3a* and G96S and RG203KP in *N*.

**Figure 2.**
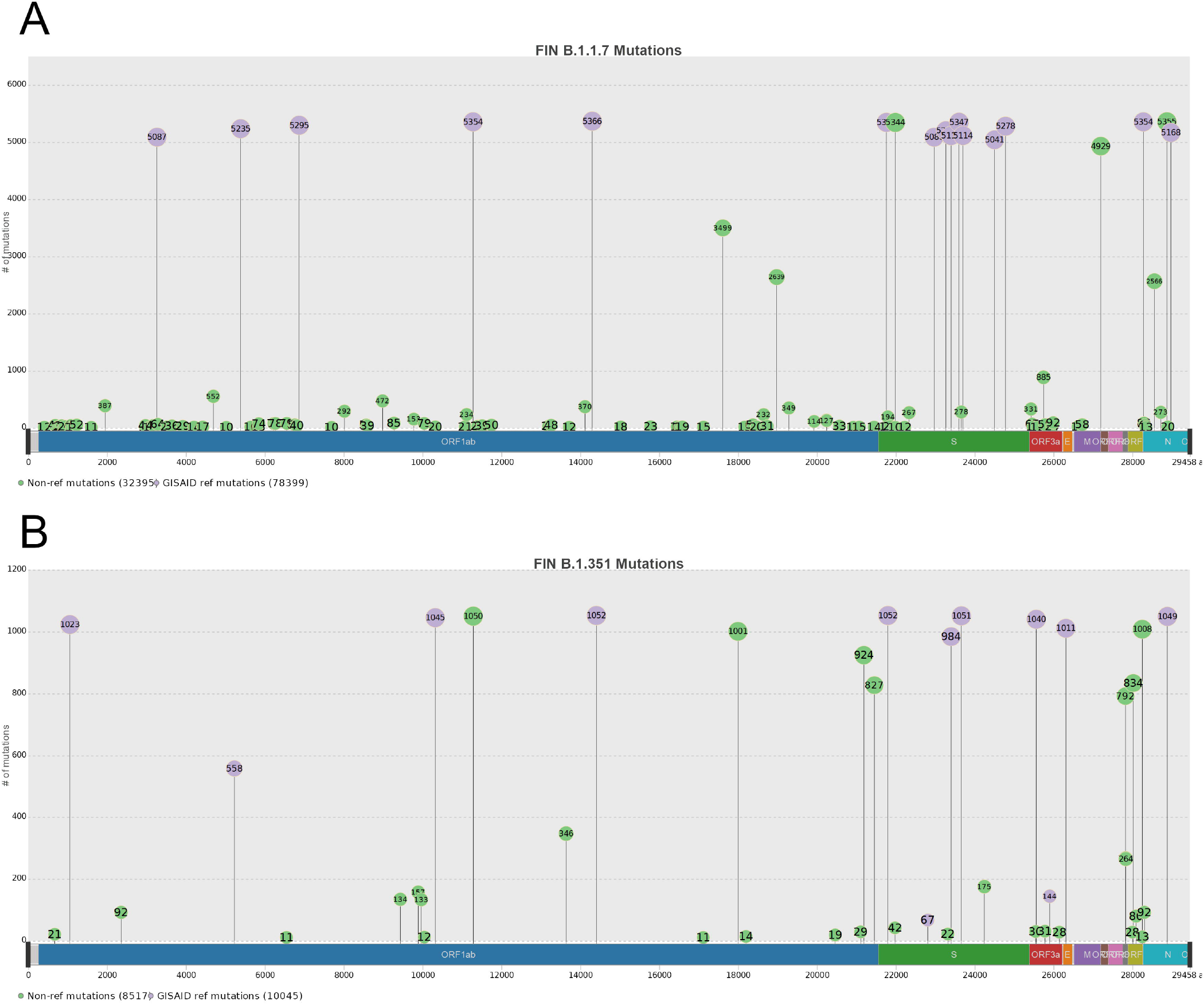
Amino acid mutations for Finnish Alpha (A) and Beta (B) datasets All mutations found in at least ten sequences are plotted. GISAID characteristic mutations (Latif *et al*., 2021a, 1, 2021c) are plotted in purple, mutations not found in the GISAID reference are plotted in green. Graphics were created with the ClusTRace interface to g3viz (Guo *et al*., 2020).

For Beta, approximately half of mutations with frequency 10% or higher were not covered by the GISAID reference (Figure 2.B). The GISAID reference aa mutations for Beta were: T265I, K1655N, K3353R and P4715L in *ORF1ab*, D80A, D614G and A701V in *S*, Q57H and S171L in *ORF3a*, P71L in *E*, T205I in *N*, while the non-reference aa mutations were: T3058I, A3209V, A3235S, D4459A, T5912I and A6976V in *ORF1ab*, T19I and I896L in *S*, M24V, I26V and I27V in *ORF7b*, K44R and I121L in *ORF8*. Note that Beta has non-GISAID mutations in Spike protein, which may potentially affect their receptor binding: T19I in 789 (75%) and I896L in 175 (16.7%) sequences.

Cluster analysis with TreeCluster (Balaban *et al*., 2019) max-clade method yielded 110 clusters for Alpha and nineteen clusters for Beta (Figures 3 and 4). Here we take a closer look at the ten clusters for Alpha and Beta that had the highest per month growth rate peaks over the analysed time period.

**Figure 3.**
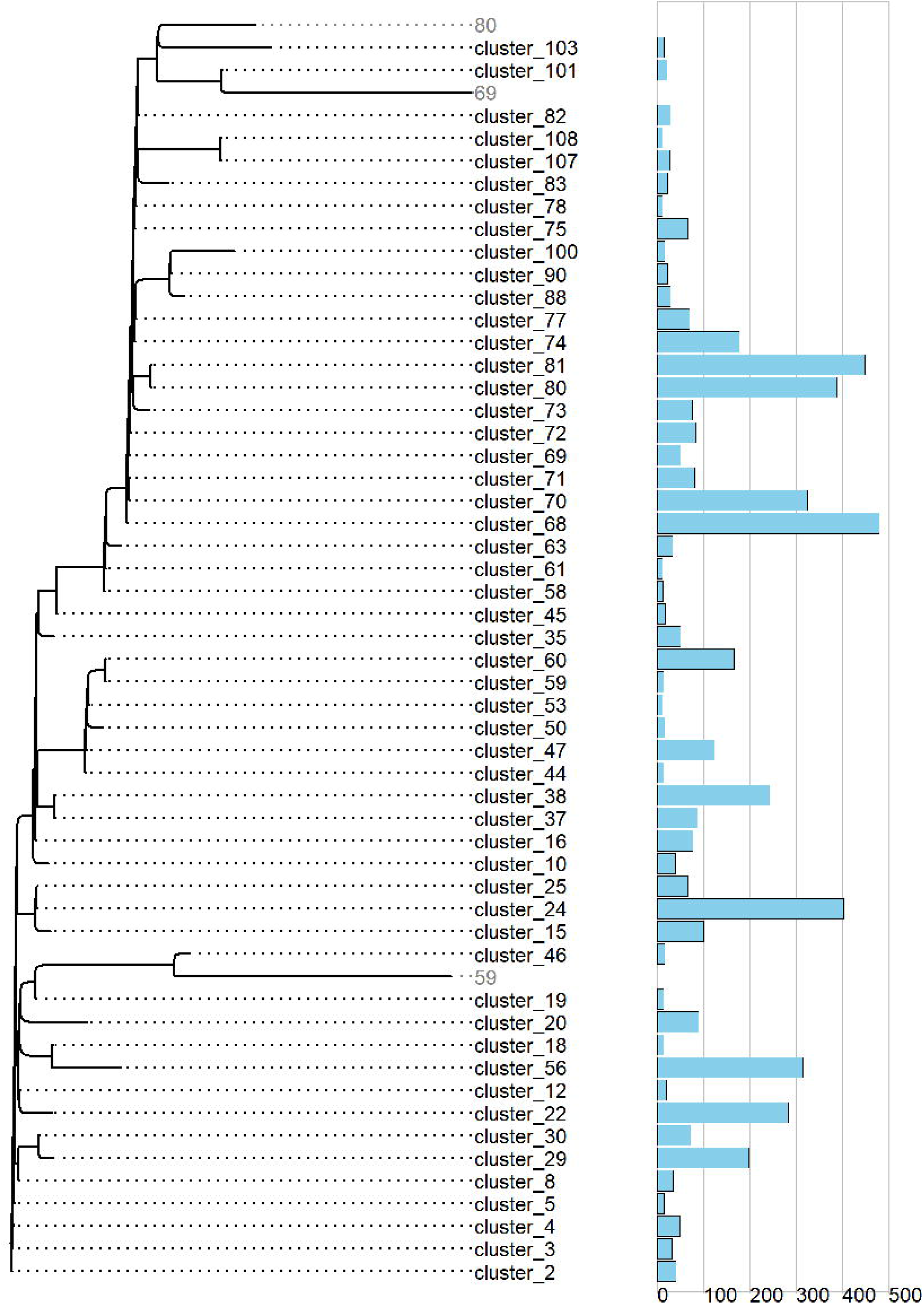
Consensus tree for Finnish Alpha dataset with clusters collapsed Bar plots on the right indicate the number of sequences in each cluster. For clarity, clusters with less than ten sequences and singletons were removed. Inner nodes with no large cluster descendants are plotted in grey.

**Figure 4.**
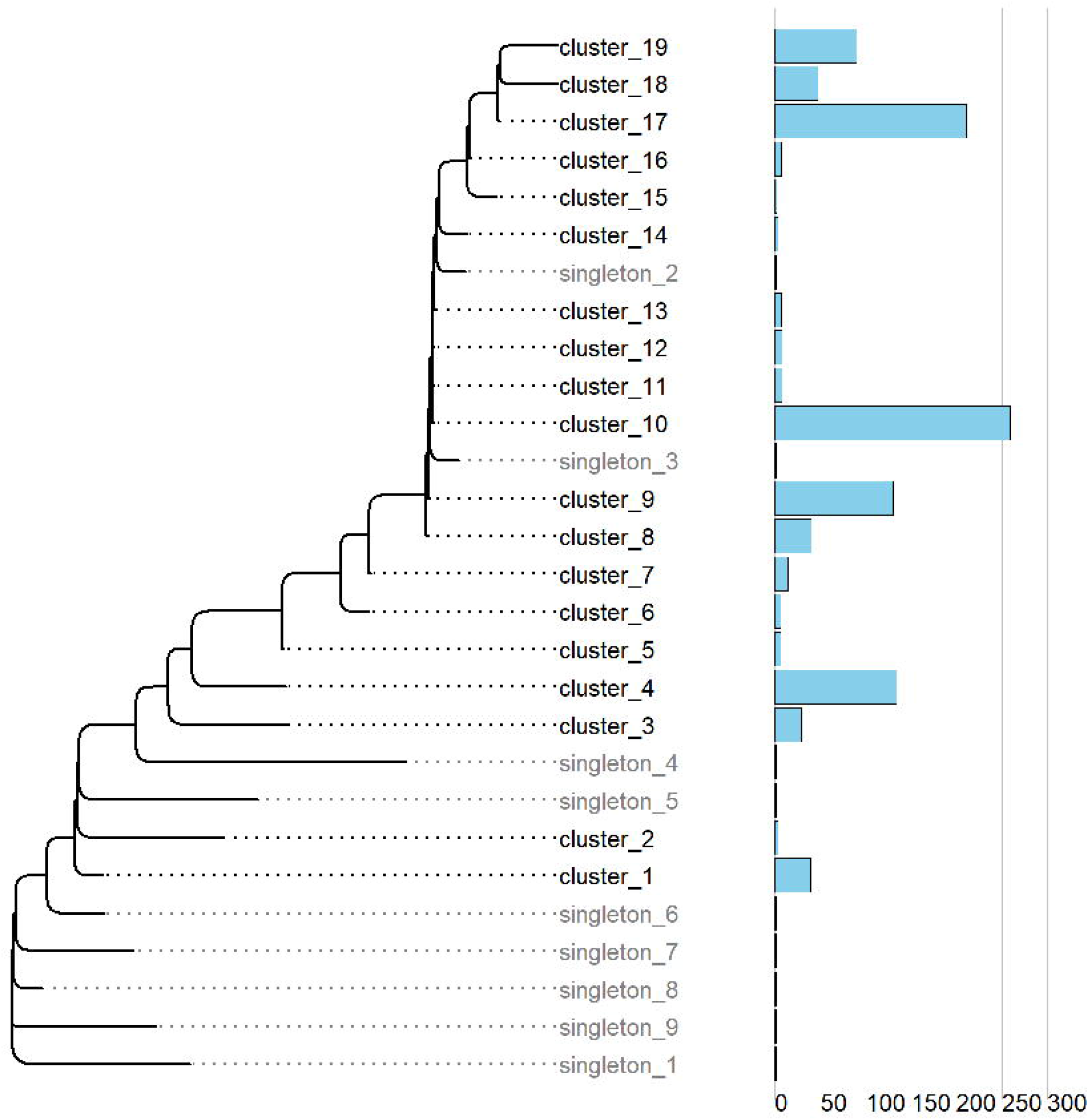
Consensus tree for Finnish Beta dataset with clusters collapsed Bar plots on the right indicate the number of sequences in each cluster. For clarity, clusters with less than ten sequences were removed.

We start by discussing Alpha clusters. Clusters with the largest per month growth peaks covered 3,243 (58.5%) of all Alpha sequences. Cluster size was between 100 (1.9%) and 479 (8.9%) sequences (Table 1). Maximal absolute growth rates (i.e. the number of sequences added to the cluster) ranged between 74 and 310 sequences per month and peak growth was during February and March. Number of non-GISAID aa mutations introduced in these clusters ranged from one to six. Solitary non-GISAID mutations in S-gene were found in clusters 56 (D80Y), 38 (D287G) and 22 (A701V) (Table 2).

**Table 2.**
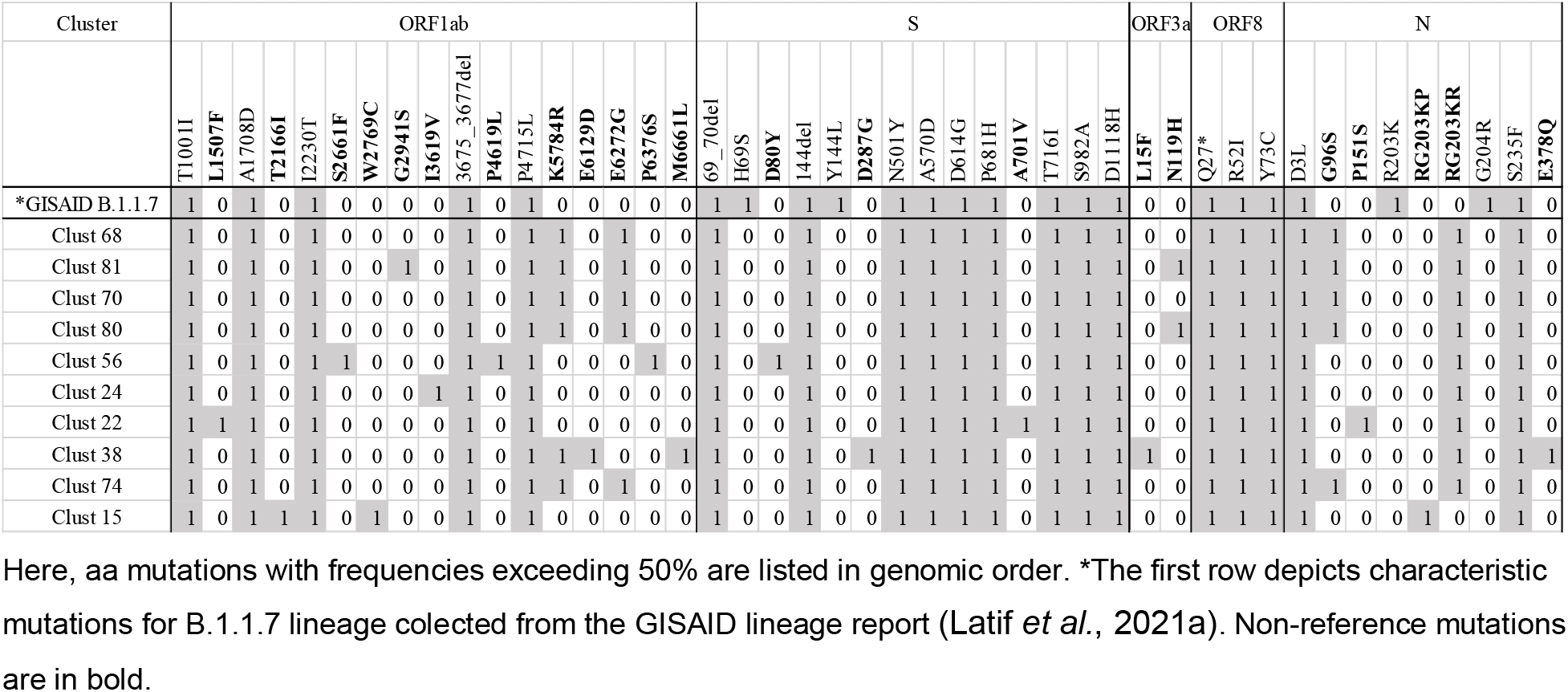
Mutations in Alpha clusters

The ten fastest growing clusters covered 979 (94.5%) of Beta sequences. Cluster size was between fourteen (1.3%) and 259 (24.6%) sequences (Table 3). Maximal growth rates ranged between 11 and 148 sequences per month and maximal growth was during February (clusters 3 and 8), March (clusters 1,4,7,10,17,18 and 19) and April (cluster 9). Number of non-GISAID aa mutations introduced in these clusters ranged from three to eight. Several clusters had non-GISAID mutations in S-gene: L18F (cluster 1), T19I (clusters 8-10, 17 and 19) and I896L (cluster 9) (Table 4).

**Table 3.** Growth rates for ten Beta clusters with maximal monthly growth rates.

| Cluster | Time Period |  |  |  |  |  | medGR | maxGR | maxGR% |
| --- | --- | --- | --- | --- | --- | --- | --- | --- | --- |
|  | 2020-12 | 2021-01 | 2021-02 | 2021-03 | 2021-04 | 2021-05 |  |  |  |
| Clust 10 | 0 | 2 | 79 | **227** | 255 | 259 | 28 | 148 | 57.1% |
| Clust 17 | 0 | 0 | 4 | **134** | 207 | 211 | 4 | 130 | 61.6% |
| Clust 19 | 0 | 0 | 0 | **83** | 90 | 90 | 0 | 83 | 92.2% |
| Clust 9 | 0 | 0 | 3 | 55 | **129** | 130 | 3 | 74 | 56.9% |
| Clust 4 | 0 | 0 | 20 | **81** | 132 | 134 | 20 | 61 | 45.5% |
| Clust 1 | 0 | 4 | 5 | **31** | 39 | 39 | 4 | 26 | 66.7% |
| Clust 8 | 0 | 0 | **21** | 34 | 39 | 40 | 5 | 21 | 52.5% |
| Clust 18 | 0 | 0 | 2 | **22** | 42 | 47 | 5 | 20 | 42.6% |
| Clust 3 | 0 | 2 | **19** | 29 | 29 | 29 | 2 | 17 | 58.6% |
| Clust 7 | 0 | 0 | 2 | **13** | 14 | 14 | 1 | 11 | 78.6% |
| Singletons | 3 | 5 | 5 | 9 | 9 | 9 |  |  |  |
| Total | 3 | 19 | 189 | **762** | 1034 | 1052 | 170 | 573 | 54.5% |

Annotation as in Table 1.

**Table 4.**
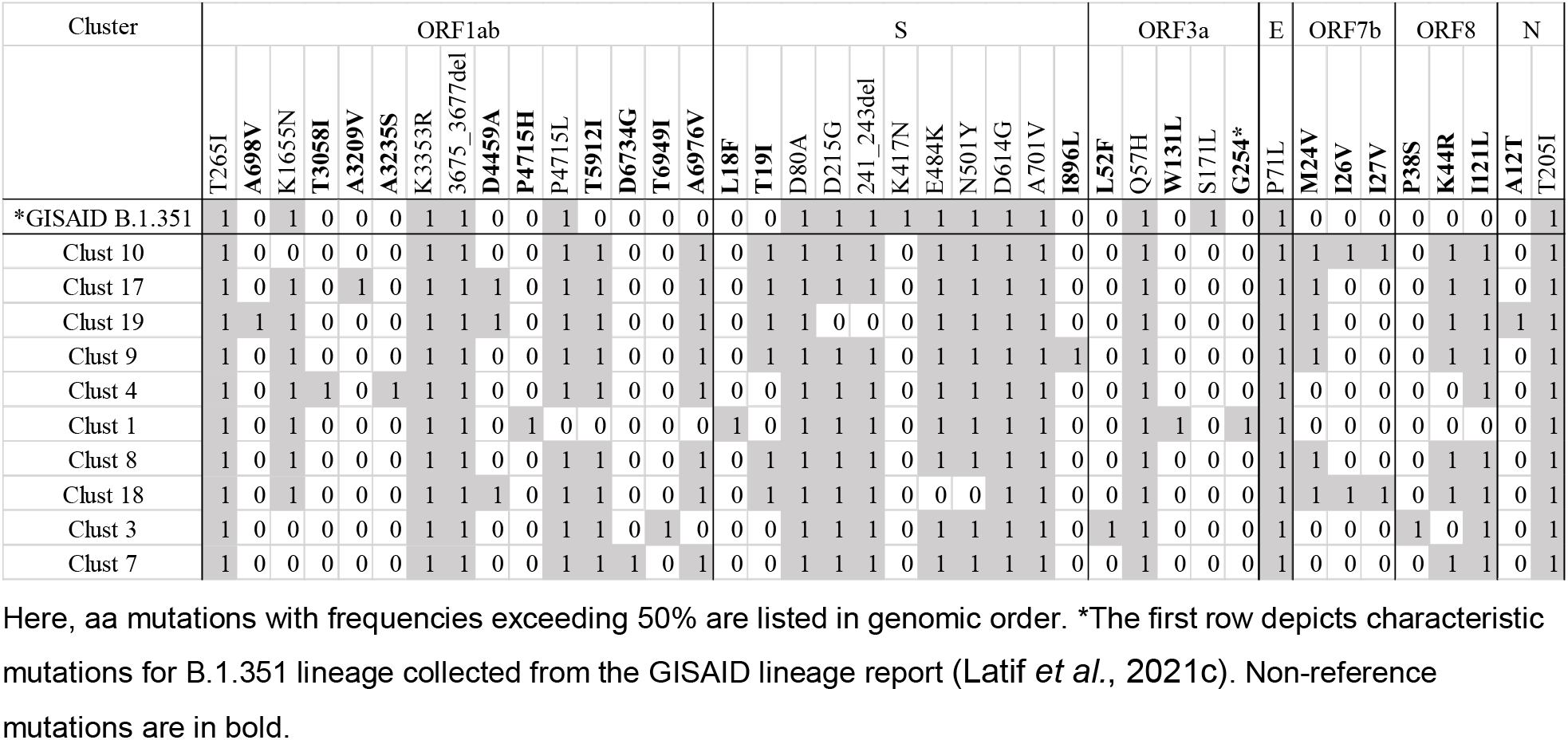
Mutations in Beta clusters

## 4 DISCUSSION

The years 2020 and 2021 could arguably be referred to as a turning point in the history of global health. The COVID-19 pandemic has demonstrated that emerging pathogens can cause havoc in our globalised world. On the other hand, the pandemic has accelerated development in many fields including genomic sequencing, bioinformatics, diagnostic testing and vaccins. The ongoing pandemic has emphasised the need for fast, scalable and, ideally pipelined, analysis of viral genomic sequences. For health authorities, it is important to be able to streamline processing large amounts of genomic sequence data into various summaries and reports that can help to make rational decisions concerning e.g. restrictions, non-pharmaceutical interventions and border control measures to minimize further spread of SARS-CoV-2. Researchers also struggle with the continuous inflow of SARS-CoV-2 sequences that need to be organized into lineages, alignments and phylogenetic trees in order to make sense of the constantly evolving pandemic.

Here, we have presented ClusTRace, a novel bioinformatic pipeline for fast and scalable analysis of large collections of SARS-CoV-2 sequences. ClusTRace supports many types of relevant analyses. These include lineage assignment, generation of multi-fasta collections, outlier filtering, generation of multiple sequence alignments and phylogenetic trees, extraction and matching of monophyletic sequence clusters, estimation of cluster growth rates and support values, nt and aa variant calling for lineages and clusters, as well as cluster and mutation reporting and visualizations. Although most of these steps can be performed separately with designated bioinformatic tools, pipelining with a high-level interface helps to cut down on the learning and operating costs of complex bioinformatic analysis. Several authors have commented on the developer-user gap between bioinformatics and other fields in biology and biomedical research (Mangul *et al*., 2019). Pipelines that are tailored to the needs of virus research are an important way to bridge this gap.

Notable pipelines for tracking viral outbreak phylodynamics include Augur, Auspice, Nextstrain, Nextclade and Pangolin (Huddleston *et al*., 2021; Aksamentov *et al*., 2021; Hadfield *et al*., 2018; O’Toole *et al*., 2021). Here, we reflect on key similarities and differences of ClusTRace to these toolkits. Pangolin and Nextclade are primarily concerned with classifying viral genomes into major lineages, while ClusTRace is designed to track more in-depth changes within lineages. Nextclade also offers mutation calling for large clades, which is similar to ClusTRace mutation calling for lineages. Nextstrain is an integration of several toolkits, including Augur for analysing sequence and phylogeographical data, and Auspice for visualising results. Like ClusTRace, Augur offers functionalities for filtering, aligning, phylogenetic reconstruction, rerooting and refinement of the obtained phylogenies, and also offers functionalities to estimate mutation frequencies. However, unlike ClusTRace, Augur also infers sequences and ancestral traits for the ancestral tree nodes. Auspice is designed to visualise phylogenetic and phylogeographic data output by Augur in an interactive webpage format. In ClusTRace, we provide different visualizations, namely Excel summaries and interactive g3viz plots for high growth-rate and/or mutation-rate clades. Unlike Nextstrain/Auspice visualizations, ClusTRace focuses directly on parts of the phylogeny that are picked out by the unsupervised clustering and analysis procedure, although this does not provide any details on the likely origin of the mutations in the tree. This approach has its advantages, such as simplicity, speed and scalability; unlike Nextstrain/Augur, ClusTRace has no need for downsampling the sequence sets. ClusTRace analysis is also largely unsupervised, i.e. clades are selected and examined for mutations and growth-peaks automatically, in effect filtering clades with alarming features that can then be checked manually more in detail.

In this work, we illustrated the intended scenario for ClusTRace usage on Finnish Alpha and Beta variants of concern. We run ClusTRace lineage assignment, generation of multi-fasta collections, outlier filtering, alignment, phylogenetic tree building, cluster extraction and variant calling as described in the results section. Overall, this approach can be described as an unsupervised phylogeny-based cluster analysis. ClusTRace uses automated clustering coupled with cluster growth rate analysis and variant calling to scan through the phylogeny. Clusters that display elevated growth rates, elevated non-reference mutation content or mutations in genomic regions that are of accentuated concern, such as the S-gene, can then be easily flagged for downstream analysis.

The current SARS-CoV-2 pandemic might endure to the unforeseeable future, and new viral variants will likely continue to emerge. Therefore, the global response has to continue to adapt and improve to mitigate the costs of the pandemic. The progress made since the start of the pandemic in early 2020 with the global implementation of full genome sequencing can be consolidated by developing efficient and scalable bioinformatic tools that are specifically tailored for genomic surveillance of viral pathogens. These tools must deliver fast, scalable and, ideally, unsupervised analysis and reporting on the pandemic events of concern. Our pipeline, ClusTRace, adds to the available toolbox the option for fast, scalable and unsupervised screening and reporting of the within or local lineage events of concern, such as elevated growth and mutation rates. ClusTRace can also be adapted for the surveillance of viral pathogens other than the SARS-CoV-2, which may prove useful in future epidemic emergencies.

## Supporting information

SFile 1

STable 1

## DATA AVAILABILITY

ClusTRace user manual and other resources are hosted at the project’s website (https://www2.helsinki.fi/en/projects/clustrace). ClusTRace source code is available from https://bitbucket.org/plyusnin/clustrace/.

## REFERENCES

Aksamentov, I. et al. (2021) Nextclade: clade assignment, mutation calling and quality control for viral genomes.

Balaban, M. et al. (2019) TreeCluster: Clustering biological sequences using phylogenetic trees. PLOS ONE, 14, e0221068.

Campbell, F. et al. (2021) Increased transmissibility and global spread of SARS-CoV-2 variants of concern as at June 2021. Eurosurveillance, 26, 2100509.

Cingolani, P. et al. (2012) A program for annotating and predicting the effects of single nucleotide polymorphisms, SnpEff: SNPs in the genome of Drosophila melanogaster strain w1118; iso-2; iso-3. Fly (Austin), 6, 80–92.

Dixon, M.G. et al. (2014) Ebola viral disease outbreak--West Africa, 2014. MMWR Morb. Mortal. Wkly. Rep., 63, 548–551.

Guo, X. et al. (2020) G3viz: an R package to interactively visualize genetic mutation data using a lollipop-diagram. Bioinformatics, 36, 928–929.

Hadfield, J. et al. (2018) Nextstrain: real-time tracking of pathogen evolution. Bioinformatics, 34, 4121–4123.

Huddleston, J. et al. (2021) Augur: a bioinformatics toolkit for phylogenetic analyses of human pathogens. J. Open Source Softw., 6, 2906.

Jalkanen, P. et al. (2021) COVID-19 mRNA vaccine induced antibody responses against three SARS-CoV-2 variants. Nat. Commun., 12, 3991.

Katoh, K. and Standley, D.M. (2013) MAFFT Multiple Sequence Alignment Software Version 7: Improvements in Performance and Usability. Mol. Biol. Evol., 30, 772–780.

Kindhauser, M.K. et al. (2016) Zika: the origin and spread of a mosquito-borne virus. Bull. World Health Organ., 94, 675–686C.

Kirola, L. (2021) Genetic emergence of B.1.617.2 in COVID-19. New Microbes New Infect., 43, 100929.

Latif, A.A. et al. (2021a) B.1.1.7 Lineage Report. outbreak.info, (https://outbreak.info/situation-reports?pango=B.1.1.7). Accessed 28 September 2021.

Latif, A.A. et al. (2021b) B.1.1.529 Lineage Report (available at https://outbreak.info/situation-reports?pango=B.1.1.529). Accessed 30 November 2021.

Latif, A.A. et al. (2021c) B.1.351 Lineage Report. outbreak.info, (https://outbreak.info/situation-reports?pango=B.1.351). Accessed 28 September 2021.

Mangul, S. et al. (2019) Systematic benchmarking of omics computational tools. Nat. Commun., 10, 1–11.

Morens, D.M. and Fauci, A.S. (2020) Emerging pandemic diseases: how we got to COVID-19. Cell.

Minh, B.Q. et al. (2020) IQ-TREE 2: New Models and Efficient Methods for Phylogenetic Inference in the Genomic Era. Mol. Biol. Evol., 37, 1530–1534.

Nguyen, P.T. et al. (2021) HaVoC, a bioinformatic pipeline for reference-based consensus assembly and lineage assignment for SARS-CoV-2 sequence.

O’Toole, Á. et al. (2021) Assignment of epidemiological lineages in an emerging pandemic using the pangolin tool. Virus Evol.

Page, A.J. et al. (2016) SNP-sites: rapid efficient extraction of SNPs from multi-FASTA alignments. Microb. Genomics, 2, e000056.

Piñeiro, C. et al. (2020) Very Fast Tree: speeding up the estimation of phylogenies for large alignments through parallelization and vectorization strategies. Bioinformatics, 36, 4658–4659.

Plyusnin, I. et al. (2020) Novel NGS pipeline for virus discovery from a wide spectrum of hosts and sample types. Virus Evol., 6, veaa091.

Shen, W. et al. (2016) SeqKit: A Cross-Platform and Ultrafast Toolkit for FASTA/Q File Manipulation. PLOS ONE, 11, e0163962.

Tegally, H. et al. (2020) Emergence and rapid spread of a new severe acute respiratory syndrome-related coronavirus 2 (SARS-CoV-2) lineage with multiple spike mutations in South Africa.

Virtanen, J. et al. (2021) Kinetics of Neutralizing Antibodies of COVID-19 Patients Tested Using Clinical D614G, B.1.1.7, and B 1.351 Isolates in Microneutralization Assays. Viruses, 13, 996.

Wise, J. (2020) Covid-19: New coronavirus variant is identified in UK. BMJ, 371, m4857. Woolhouse, M.E.J. and Gowtage-Sequeria, S. Host Range and Emerging and Reemerging Pathogens - Volume 11, Number 12—December 2005 - Emerging Infectious Diseases journal - CDC.

Wu, F. et al. (2020) A new coronavirus associated with human respiratory disease in China. Nature, 579, 265–269.

Zwagemaker, F. et al. (2021) DennisSchmitz/Jovian: 1.2.07 Zenodo.

