## Supplementary material for "ClusTRace, a bioinformatic pipeline for analyzing clusters in virus phylogenies": SFile 1: d3.v4.min.js

```
// https://d3js.org Version 4.13.0. Copyright 2018 Mike Bostock.
(function(t,n){"object"==typeof exports&&"undefined"!=typeof module?n(exports):"function"==typeof define&&define.amd?define(["exports"],n):n(t.d3=t.d3||{})})(this,function(t){"use strict";function n(t,n){return t<n?-1:t>n?1:t>=n?0:NaN}function e(t){return 1===t.length&&(t=function(t){return function(e,r){return n(t(e),r)}}(t)),{left:function(n,e,r,i){for(null==r&&(r=0),null==i&&(i=n.length);r<i;){var o=r+i>>>1;t(n[o],e)<0?r=o+1:i=o}return r},right:function(n,e,r,i){for(null==r&&(r=0),null==i&&(i=n.length);r<i;){var o=r+i>>>1;t(n[o],e)>0?i=o:r=o+1}return r}}}function r(t,n){return[t,n]}function i(t){return null===t?NaN:+t}function o(t,n){var e,r,o=t.length,u=0,a=-1,c=0,s=0;if(null==n)for(;++a<o;)isNaN(e=i(t[a]))||(s+=(r=e-c)*(e-(c+=r/++u)));else for(;++a<o;)isNaN(e=i(n(t[a],a,t)))||(s+=(r=e-c)*(e-(c+=r/++u)));if(u>1)return s/(u-1)}function u(t,n){var e=o(t,n);return e?Math.sqrt(e):e}function a(t,n){var e,r,i,o=t.length,u=-1;if(null==n){for(;++u<o;)if(null!=(e=t[u])&&e>=e)for(r=i=e;++u<o;)null!=(e=t[u])&&(r>e&&(r=e),i<e&&(i=e))}else for(;++u<o;)if(null!=(e=n(t[u],u,t))&&e>=e)for(r=i=e;++u<o;)null!=(e=n(t[u],u,t))&&(r>e&&(r=e),i<e&&(i=e));return[r,i]}function c(t){return function(){return t}}function s(t){return t}function f(t,n,e){t=+t,n=+n,e=(i=arguments.length)<2?(n=t,t=0,1):i<3?1:+e;for(var r=-1,i=0|Math.max(0,Math.ceil((n-t)/e)),o=new Array(i);++r<i;)o[r]=t+r*e;return o}function l(t,n,e){var r,i,o,u,a=-1;if(n=+n,t=+t,e=+e,t===n&&e>0)return[t];if((r=n<t)&&(i=t,t=n,n=i),0===(u=h(t,n,e))||!isFinite(u))return[];if(u>0)for(t=Math.ceil(t/u),n=Math.floor(n/u),o=new Array(i=Math.ceil(n-t+1));++a<i;)o[a]=(t+a)*u;else for(t=Math.floor(t*u),n=Math.ceil(n*u),o=new Array(i=Math.ceil(t-n+1));++a<i;)o[a]=(t-a)/u;return r&&o.reverse(),o}function h(t,n,e){var r=(n-t)/Math.max(0,e),i=Math.floor(Math.log(r)/Math.LN10),o=r/Math.pow(10,i);return i>=0?(o>=Hs?10:o>=js?5:o>=Xs?2:1)*Math.pow(10,i):-Math.pow(10,-i)/(o>=Hs?10:o>=js?5:o>=Xs?2:1)}function p(t,n,e){var r=Math.abs(n-t)/Math.max(0,e),i=Math.pow(10,Math.floor(Math.log(r)/Math.LN10)),o=r/i;return o>=Hs?i*=10:o>=js?i*=5:o>=Xs&&(i*=2),n<t?-i:i}function d(t){return Math.ceil(Math.log(t.length)/Math.LN2)+1}function v(t,n,e){if(null==e&&(e=i),r=t.length){if((n=+n)<=0||r<2)return+e(t[0],0,t);if(n>=1)return+e(t[r-1],r-1,t);var r,o=(r-1)*n,u=Math.floor(o),a=+e(t[u],u,t);return a+(+e(t[u+1],u+1,t)-a)*(o-u)}}function g(t){for(var n,e,r,i=t.length,o=-1,u=0;++o<i;)u+=t[o].length;for(e=new Array(u);--i>=0;)for(n=(r=t[i]).length;--n>=0;)e[--u]=r[n];return e}function _(t,n){var e,r,i=t.length,o=-1;if(null==n){for(;++o<i;)if(null!=(e=t[o])&&e>=e)for(r=e;++o<i;)null!=(e=t[o])&&r>e&&(r=e)}else for(;++o<i;)if(null!=(e=n(t[o],o,t))&&e>=e)for(r=e;++o<i;)null!=(e=n(t[o],o,t))&&r>e&&(r=e);return r}function y(t){if(!(i=t.length))return[];for(var n=-1,e=_(t,m),r=new Array(e);++n<e;)for(var i,o=-1,u=r[n]=new Array(i);++o<i;)u[o]=t[o][n];return r}function m(t){return t.length}function x(t){return t}function b(t){return"translate("+(t+.5)+",0)"}function w(t){return"translate(0,"+(t+.5)+")"}function M(){return!this.__axis}function T(t,n){function e(e){var h=null==i?n.ticks?n.ticks.apply(n,r):n.domain():i,p=null==o?n.tickFormat?n.tickFormat.apply(n,r):x:o,d=Math.max(u,0)+c,v=n.range(),g=+v[0]+.5,_=+v[v.length-1]+.5,y=(n.bandwidth?function(t){var n=Math.max(0,t.bandwidth()-1)/2;return t.round()&&(n=Math.round(n)),function(e){return+t(e)+n}}:function(t){return function(n){return+t(n)}})(n.copy()),m=e.selection?e.selection():e,b=m.selectAll(".domain").data([null]),w=m.selectAll(".tick").data(h,n).order(),T=w.exit(),N=w.enter().append("g").attr("class","tick"),k=w.select("line"),S=w.select("text");b=b.merge(b.enter().insert("path",".tick").attr("class","domain").attr("stroke","#000")),w=w.merge(N),k=k.merge(N.append("line").attr("stroke","#000").attr(f+"2",s*u)),S=S.merge(N.append("text").attr("fill","#000").attr(f,s*d).attr("dy",t===$s?"0em":t===Zs?"0.71em":"0.32em")),e!==m&&(b=b.transition(e),w=w.transition(e),k=k.transition(e),S=S.transition(e),T=T.transition(e).attr("opacity",Qs).attr("transform",function(t){return isFinite(t=y(t))?l(t):this.getAttribute("transform")}),N.attr("opacity",Qs).attr("transform",function(t){var n=this.parentNode.__axis;return l(n&&isFinite(n=n(t))?n:y(t))})),T.remove(),b.attr("d",t===Gs||t==Ws?"M"+s*a+","+g+"H0.5V"+_+"H"+s*a:"M"+g+","+s*a+"V0.5H"+_+"V"+s*a),w.attr("opacity",1).attr("transform",function(t){return l(y(t))}),k.attr(f+"2",s*u),S.attr(f,s*d).text(p),m.filter(M).attr("fill","none").attr("font-size",10).attr("font-family","sans-serif").attr("text-anchor",t===Ws?"start":t===Gs?"end":"middle"),m.each(function(){this.__axis=y})}var r=[],i=null,o=null,u=6,a=6,c=3,s=t===$s||t===Gs?-1:1,f=t===Gs||t===Ws?"x":"y",l=t===$s||t===Zs?b:w;return e.scale=function(t){return arguments.length?(n=t,e):n},e.ticks=function(){return r=Vs.call(arguments),e},e.tickArguments=function(t){return arguments.length?(r=null==t?[]:Vs.call(t),e):r.slice()},e.tickValues=function(t){return arguments.length?(i=null==t?null:Vs.call(t),e):i&&i.slice()},e.tickFormat=function(t){return arguments.length?(o=t,e):o},e.tickSize=function(t){return arguments.length?(u=a=+t,e):u},e.tickSizeInner=function(t){return arguments.length?(u=+t,e):u},e.tickSizeOuter=function(t){return arguments.length?(a=+t,e):a},e.tickPadding=function(t){return arguments.length?(c=+t,e):c},e}function N(){for(var t,n=0,e=arguments.length,r={};n<e;++n){if(!(t=arguments[n]+"")||t in r)throw new Error("illegal type: "+t);r[t]=[]}return new k(r)}function k(t){this._=t}function S(t,n,e){for(var r=0,i=t.length;r<i;++r)if(t[r].name===n){t[r]=Js,t=t.slice(0,r).concat(t.slice(r+1));break}return null!=e&&t.push({name:n,value:e}),t}function E(t){var n=t+="",e=n.indexOf(":");return e>=0&&"xmlns"!==(n=t.slice(0,e))&&(t=t.slice(e+1)),tf.hasOwnProperty(n)?{space:tf[n],local:t}:t}function A(t){var n=E(t);return(n.local?function(t){return function(){return this.ownerDocument.createElementNS(t.space,t.local)}}:function(t){return function(){var n=this.ownerDocument,e=this.namespaceURI;return e===Ks&&n.documentElement.namespaceURI===Ks?n.createElement(t):n.createElementNS(e,t)}})(n)}function C(){}function z(t){return null==t?C:function(){return this.querySelector(t)}}function P(){return[]}function R(t){return null==t?P:function(){return this.querySelectorAll(t)}}function L(t){return new Array(t.length)}function q(t,n){this.ownerDocument=t.ownerDocument,this.namespaceURI=t.namespaceURI,this._next=null,this._parent=t,this.__data__=n}function D(t,n,e,r,i,o){for(var u,a=0,c=n.length,s=o.length;a<s;++a)(u=n[a])?(u.__data__=o[a],r[a]=u):e[a]=new q(t,o[a]);for(;a<c;++a)(u=n[a])&&(i[a]=u)}function U(t,n,e,r,i,o,u){var a,c,s,f={},l=n.length,h=o.length,p=new Array(l);for(a=0;a<l;++a)(c=n[a])&&(p[a]=s=uf+u.call(c,c.__data__,a,n),s in f?i[a]=c:f[s]=c);for(a=0;a<h;++a)(c=f[s=uf+u.call(t,o[a],a,o)])?(r[a]=c,c.__data__=o[a],f[s]=null):e[a]=new q(t,o[a]);for(a=0;a<l;++a)(c=n[a])&&f[p[a]]===c&&(i[a]=c)}function O(t,n){return t<n?-1:t>n?1:t>=n?0:NaN}function F(t){return t.ownerDocument&&t.ownerDocument.defaultView||t.document&&t||t.defaultView}function I(t,n){return t.style.getPropertyValue(n)||F(t).getComputedStyle(t,null).getPropertyValue(n)}function Y(t){return t.trim().split(/^|\s+/)}function B(t){return t.classList||new H(t)}function H(t){this._node=t,this._names=Y(t.getAttribute("class")||"")}function j(t,n){for(var e=B(t),r=-1,i=n.length;++r<i;)e.add(n[r])}function X(t,n){for(var e=B(t),r=-1,i=n.length;++r<i;)e.remove(n[r])}function V(){this.textContent=""}function $(){this.innerHTML=""}function W(){this.nextSibling&&this.parentNode.appendChild(this)}function Z(){this.previousSibling&&this.parentNode.insertBefore(this,this.parentNode.firstChild)}function G(){return null}function Q(){var t=this.parentNode;t&&t.removeChild(this)}function J(){return this.parentNode.insertBefore(this.cloneNode(!1),this.nextSibling)}function K(){return this.parentNode.insertBefore(this.cloneNode(!0),this.nextSibling)}function tt(t,n,e){return t=nt(t,n,e),function(n){var e=n.relatedTarget;e&&(e===this||8&e.compareDocumentPosition(this))||t.call(this,n)}}function nt(n,e,r){return function(i){var o=t.event;t.event=i;try{n.call(this,this.__data__,e,r)}finally{t.event=o}}}function et(t){return function(){var n=this.__on;if(n){for(var e,r=0,i=-1,o=n.length;r<o;++r)e=n[r],t.type&&e.type!==t.type||e.name!==t.name?n[++i]=e:this.removeEventListener(e.type,e.listener,e.capture);++i?n.length=i:delete this.__on}}}function rt(t,n,e){var r=af.hasOwnProperty(t.type)?tt:nt;return function(i,o,u){var a,c=this.__on,s=r(n,o,u);if(c)for(var f=0,l=c.length;f<l;++f)if((a=c[f]).type===t.type&&a.name===t.name)return this.removeEventListener(a.type,a.listener,a.capture),this.addEventListener(a.type,a.listener=s,a.capture=e),void(a.value=n);this.addEventListener(t.type,s,e),a={type:t.type,name:t.name,value:n,listener:s,capture:e},c?c.push(a):this.__on=[a]}}function it(n,e,r,i){var o=t.event;n.sourceEvent=t.event,t.event=n;try{return e.apply(r,i)}finally{t.event=o}}function ot(t,n,e){var r=F(t),i=r.CustomEvent;"function"==typeof i?i=new i(n,e):(i=r.document.createEvent("Event"),e?(i.initEvent(n,e.bubbles,e.cancelable),i.detail=e.detail):i.initEvent(n,!1,!1)),t.dispatchEvent(i)}function ut(t,n){this._groups=t,this._parents=n}function at(){return new ut([[document.documentElement]],cf)}function ct(t){return"string"==typeof t?new ut([[document.querySelector(t)]],[document.documentElement]):new ut([[t]],cf)}function st(){return new ft}function ft(){this._="@"+(++sf).toString(36)}function lt(){for(var n,e=t.event;n=e.sourceEvent;)e=n;return e}function ht(t,n){var e=t.ownerSVGElement||t;if(e.createSVGPoint){var r=e.createSVGPoint();return r.x=n.clientX,r.y=n.clientY,r=r.matrixTransform(t.getScreenCTM().inverse()),[r.x,r.y]}var i=t.getBoundingClientRect();return[n.clientX-i.left-t.clientLeft,n.clientY-i.top-t.clientTop]}function pt(t){var n=lt();return n.changedTouches&&(n=n.changedTouches[0]),ht(t,n)}function dt(t,n,e){arguments.length<3&&(e=n,n=lt().changedTouches);for(var r,i=0,o=n?n.length:0;i<o;++i)if((r=n[i]).identifier===e)return ht(t,r);return null}function vt(){t.event.stopImmediatePropagation()}function gt(){t.event.preventDefault(),t.event.stopImmediatePropagation()}function _t(t){var n=t.document.documentElement,e=ct(t).on("dragstart.drag",gt,!0);"onselectstart"in n?e.on("selectstart.drag",gt,!0):(n.__noselect=n.style.MozUserSelect,n.style.MozUserSelect="none")}function yt(t,n){var e=t.document.documentElement,r=ct(t).on("dragstart.drag",null);n&&(r.on("click.drag",gt,!0),setTimeout(function(){r.on("click.drag",null)},0)),"onselectstart"in e?r.on("selectstart.drag",null):(e.style.MozUserSelect=e.__noselect,delete e.__noselect)}function mt(t){return function(){return t}}function xt(t,n,e,r,i,o,u,a,c,s){this.target=t,this.type=n,this.subject=e,this.identifier=r,this.active=i,this.x=o,this.y=u,this.dx=a,this.dy=c,this._=s}function bt(){return!t.event.button}function wt(){return this.parentNode}function Mt(n){return null==n?{x:t.event.x,y:t.event.y}:n}function Tt(){return"ontouchstart"in this}function Nt(t,n,e){t.prototype=n.prototype=e,e.constructor=t}function kt(t,n){var e=Object.create(t.prototype);for(var r in n)e[r]=n[r];return e}function St(){}function Et(t){var n;return t=(t+"").trim().toLowerCase(),(n=pf.exec(t))?(n=parseInt(n[1],16),new Rt(n>>8&15|n>>4&240,n>>4&15|240&n,(15&n)<<4|15&n,1)):(n=df.exec(t))?At(parseInt(n[1],16)):(n=vf.exec(t))?new Rt(n[1],n[2],n[3],1):(n=gf.exec(t))?new Rt(255*n[1]/100,255*n[2]/100,255*n[3]/100,1):(n=_f.exec(t))?Ct(n[1],n[2],n[3],n[4]):(n=yf.exec(t))?Ct(255*n[1]/100,255*n[2]/100,255*n[3]/100,n[4]):(n=mf.exec(t))?Lt(n[1],n[2]/100,n[3]/100,1):(n=xf.exec(t))?Lt(n[1],n[2]/100,n[3]/100,n[4]):bf.hasOwnProperty(t)?At(bf[t]):"transparent"===t?new Rt(NaN,NaN,NaN,0):null}function At(t){return new Rt(t>>16&255,t>>8&255,255&t,1)}function Ct(t,n,e,r){return r<=0&&(t=n=e=NaN),new Rt(t,n,e,r)}function zt(t){return t instanceof St||(t=Et(t)),t?(t=t.rgb(),new Rt(t.r,t.g,t.b,t.opacity)):new Rt}function Pt(t,n,e,r){return 1===arguments.length?zt(t):new Rt(t,n,e,null==r?1:r)}function Rt(t,n,e,r){this.r=+t,this.g=+n,this.b=+e,this.opacity=+r}function Lt(t,n,e,r){return r<=0?t=n=e=NaN:e<=0||e>=1?t=n=NaN:n<=0&&(t=NaN),new Dt(t,n,e,r)}function qt(t,n,e,r){return 1===arguments.length?function(t){if(t instanceof Dt)return new Dt(t.h,t.s,t.l,t.opacity);if(t instanceof St||(t=Et(t)),!t)return new Dt;if(t instanceof Dt)return t;var n=(t=t.rgb()).r/255,e=t.g/255,r=t.b/255,i=Math.min(n,e,r),o=Math.max(n,e,r),u=NaN,a=o-i,c=(o+i)/2;return a?(u=n===o?(e-r)/a+6*(e<r):e===o?(r-n)/a+2:(n-e)/a+4,a/=c<.5?o+i:2-o-i,u*=60):a=c>0&&c<1?0:u,new Dt(u,a,c,t.opacity)}(t):new Dt(t,n,e,null==r?1:r)}function Dt(t,n,e,r){this.h=+t,this.s=+n,this.l=+e,this.opacity=+r}function Ut(t,n,e){return 255*(t<60?n+(e-n)*t/60:t<180?e:t<240?n+(e-n)*(240-t)/60:n)}function Ot(t){if(t instanceof It)return new It(t.l,t.a,t.b,t.opacity);if(t instanceof Vt){var n=t.h*wf;return new It(t.l,Math.cos(n)*t.c,Math.sin(n)*t.c,t.opacity)}t instanceof Rt||(t=zt(t));var e=jt(t.r),r=jt(t.g),i=jt(t.b),o=Yt((.4124564*e+.3575761*r+.1804375*i)/Tf),u=Yt((.2126729*e+.7151522*r+.072175*i)/Nf);return new It(116*u-16,500*(o-u),200*(u-Yt((.0193339*e+.119192*r+.9503041*i)/kf)),t.opacity)}function Ft(t,n,e,r){return 1===arguments.length?Ot(t):new It(t,n,e,null==r?1:r)}function It(t,n,e,r){this.l=+t,this.a=+n,this.b=+e,this.opacity=+r}function Yt(t){return t>Cf?Math.pow(t,1/3):t/Af+Sf}function Bt(t){return t>Ef?t*t*t:Af*(t-Sf)}function Ht(t){return 255*(t<=.0031308?12.92*t:1.055*Math.pow(t,1/2.4)-.055)}function jt(t){return(t/=255)<=.04045?t/12.92:Math.pow((t+.055)/1.055,2.4)}function Xt(t,n,e,r){return 1===arguments.length?function(t){if(t instanceof Vt)return new Vt(t.h,t.c,t.l,t.opacity);t instanceof It||(t=Ot(t));var n=Math.atan2(t.b,t.a)*Mf;return new Vt(n<0?n+360:n,Math.sqrt(t.a*t.a+t.b*t.b),t.l,t.opacity)}(t):new Vt(t,n,e,null==r?1:r)}function Vt(t,n,e,r){this.h=+t,this.c=+n,this.l=+e,this.opacity=+r}function $t(t,n,e,r){return 1===arguments.length?function(t){if(t instanceof Wt)return new Wt(t.h,t.s,t.l,t.opacity);t instanceof Rt||(t=zt(t));var n=t.r/255,e=t.g/255,r=t.b/255,i=(Df*r+Lf*n-qf*e)/(Df+Lf-qf),o=r-i,u=(Rf*(e-i)-zf*o)/Pf,a=Math.sqrt(u*u+o*o)/(Rf*i*(1-i)),c=a?Math.atan2(u,o)*Mf-120:NaN;return new Wt(c<0?c+360:c,a,i,t.opacity)}(t):new Wt(t,n,e,null==r?1:r)}function Wt(t,n,e,r){this.h=+t,this.s=+n,this.l=+e,this.opacity=+r}function Zt(t,n,e,r,i){var o=t*t,u=o*t;return((1-3*t+3*o-u)*n+(4-6*o+3*u)*e+(1+3*t+3*o-3*u)*r+u*i)/6}function Gt(t){var n=t.length-1;return function(e){var r=e<=0?e=0:e>=1?(e=1,n-1):Math.floor(e*n),i=t[r],o=t[r+1],u=r>0?t[r-1]:2*i-o,a=r<n-1?t[r+2]:2*o-i;return Zt((e-r/n)*n,u,i,o,a)}}function Qt(t){var n=t.length;return function(e){var r=Math.floor(((e%=1)<0?++e:e)*n),i=t[(r+n-1)%n],o=t[r%n],u=t[(r+1)%n],a=t[(r+2)%n];return Zt((e-r/n)*n,i,o,u,a)}}function Jt(t){return function(){return t}}function Kt(t,n){return function(e){return t+e*n}}function tn(t,n){var e=n-t;return e?Kt(t,e>180||e<-180?e-360*Math.round(e/360):e):Jt(isNaN(t)?n:t)}function nn(t){return 1==(t=+t)?en:function(n,e){return e-n?function(t,n,e){return t=Math.pow(t,e),n=Math.pow(n,e)-t,e=1/e,function(r){return Math.pow(t+r*n,e)}}(n,e,t):Jt(isNaN(n)?e:n)}}function en(t,n){var e=n-t;return e?Kt(t,e):Jt(isNaN(t)?n:t)}function rn(t){return function(n){var e,r,i=n.length,o=new Array(i),u=new Array(i),a=new Array(i);for(e=0;e<i;++e)r=Pt(n[e]),o[e]=r.r||0,u[e]=r.g||0,a[e]=r.b||0;return o=t(o),u=t(u),a=t(a),r.opacity=1,function(t){return r.r=o(t),r.g=u(t),r.b=a(t),r+""}}}function on(t,n){var e,r=n?n.length:0,i=t?Math.min(r,t.length):0,o=new Array(i),u=new Array(r);for(e=0;e<i;++e)o[e]=fn(t[e],n[e]);for(;e<r;++e)u[e]=n[e];return function(t){for(e=0;e<i;++e)u[e]=o[e](t);return u}}function un(t,n){var e=new Date;return t=+t,n-=t,function(r){return e.setTime(t+n*r),e}}function an(t,n){return t=+t,n-=t,function(e){return t+n*e}}function cn(t,n){var e,r={},i={};null!==t&&"object"==typeof t||(t={}),null!==n&&"object"==typeof n||(n={});for(e in n)e in t?r[e]=fn(t[e],n[e]):i[e]=n[e];return function(t){for(e in r)i[e]=r[e](t);return i}}function sn(t,n){var e,r,i,o=Vf.lastIndex=$f.lastIndex=0,u=-1,a=[],c=[];for(t+="",n+="";(e=Vf.exec(t))&&(r=$f.exec(n));)(i=r.index)>o&&(i=n.slice(o,i),a[u]?a[u]+=i:a[++u]=i),(e=e[0])===(r=r[0])?a[u]?a[u]+=r:a[++u]=r:(a[++u]=null,c.push({i:u,x:an(e,r)})),o=$f.lastIndex;return o<n.length&&(i=n.slice(o),a[u]?a[u]+=i:a[++u]=i),a.length<2?c[0]?function(t){return function(n){return t(n)+""}}(c[0].x):function(t){return function(){return t}}(n):(n=c.length,function(t){for(var e,r=0;r<n;++r)a[(e=c[r]).i]=e.x(t);return a.join("")})}function fn(t,n){var e,r=typeof n;return null==n||"boolean"===r?Jt(n):("number"===r?an:"string"===r?(e=Et(n))?(n=e,Hf):sn:n instanceof Et?Hf:n instanceof Date?un:Array.isArray(n)?on:"function"!=typeof n.valueOf&&"function"!=typeof n.toString||isNaN(n)?cn:an)(t,n)}function ln(t,n){return t=+t,n-=t,function(e){return Math.round(t+n*e)}}function hn(t,n,e,r,i,o){var u,a,c;return(u=Math.sqrt(t*t+n*n))&&(t/=u,n/=u),(c=t*e+n*r)&&(e-=t*c,r-=n*c),(a=Math.sqrt(e*e+r*r))&&(e/=a,r/=a,c/=a),t*r<n*e&&(t=-t,n=-n,c=-c,u=-u),{translateX:i,translateY:o,rotate:Math.atan2(n,t)*Wf,skewX:Math.atan(c)*Wf,scaleX:u,scaleY:a}}function pn(t,n,e,r){function i(t){return t.length?t.pop()+" ":""}return function(o,u){var a=[],c=[];return o=t(o),u=t(u),function(t,r,i,o,u,a){if(t!==i||r!==o){var c=u.push("translate(",null,n,null,e);a.push({i:c-4,x:an(t,i)},{i:c-2,x:an(r,o)})}else(i||o)&&u.push("translate("+i+n+o+e)}(o.translateX,o.translateY,u.translateX,u.translateY,a,c),function(t,n,e,o){t!==n?(t-n>180?n+=360:n-t>180&&(t+=360),o.push({i:e.push(i(e)+"rotate(",null,r)-2,x:an(t,n)})):n&&e.push(i(e)+"rotate("+n+r)}(o.rotate,u.rotate,a,c),function(t,n,e,o){t!==n?o.push({i:e.push(i(e)+"skewX(",null,r)-2,x:an(t,n)}):n&&e.push(i(e)+"skewX("+n+r)}(o.skewX,u.skewX,a,c),function(t,n,e,r,o,u){if(t!==e||n!==r){var a=o.push(i(o)+"scale(",null,",",null,")");u.push({i:a-4,x:an(t,e)},{i:a-2,x:an(n,r)})}else 1===e&&1===r||o.push(i(o)+"scale("+e+","+r+")")}(o.scaleX,o.scaleY,u.scaleX,u.scaleY,a,c),o=u=null,function(t){for(var n,e=-1,r=c.length;++e<r;)a[(n=c[e]).i]=n.x(t);return a.join("")}}}function dn(t){return((t=Math.exp(t))+1/t)/2}function vn(t,n){var e,r,i=t[0],o=t[1],u=t[2],a=n[0],c=n[1],s=n[2],f=a-i,l=c-o,h=f*f+l*l;if(h<nl)r=Math.log(s/u)/Jf,e=function(t){return[i+t*f,o+t*l,u*Math.exp(Jf*t*r)]};else{var p=Math.sqrt(h),d=(s*s-u*u+tl*h)/(2*u*Kf*p),v=(s*s-u*u-tl*h)/(2*s*Kf*p),g=Math.log(Math.sqrt(d*d+1)-d),_=Math.log(Math.sqrt(v*v+1)-v);r=(_-g)/Jf,e=function(t){var n=t*r,e=dn(g),a=u/(Kf*p)*(e*function(t){return((t=Math.exp(2*t))-1)/(t+1)}(Jf*n+g)-function(t){return((t=Math.exp(t))-1/t)/2}(g));return[i+a*f,o+a*l,u*e/dn(Jf*n+g)]}}return e.duration=1e3*r,e}function gn(t){return function(n,e){var r=t((n=qt(n)).h,(e=qt(e)).h),i=en(n.s,e.s),o=en(n.l,e.l),u=en(n.opacity,e.opacity);return function(t){return n.h=r(t),n.s=i(t),n.l=o(t),n.opacity=u(t),n+""}}}function _n(t){return function(n,e){var r=t((n=Xt(n)).h,(e=Xt(e)).h),i=en(n.c,e.c),o=en(n.l,e.l),u=en(n.opacity,e.opacity);return function(t){return n.h=r(t),n.c=i(t),n.l=o(t),n.opacity=u(t),n+""}}}function yn(t){return function n(e){function r(n,r){var i=t((n=$t(n)).h,(r=$t(r)).h),o=en(n.s,r.s),u=en(n.l,r.l),a=en(n.opacity,r.opacity);return function(t){return n.h=i(t),n.s=o(t),n.l=u(Math.pow(t,e)),n.opacity=a(t),n+""}}return e=+e,r.gamma=n,r}(1)}function mn(){return pl||(gl(xn),pl=vl.now()+dl)}function xn(){pl=0}function bn(){this._call=this._time=this._next=null}function wn(t,n,e){var r=new bn;return r.restart(t,n,e),r}function Mn(){mn(),++cl;for(var t,n=Yf;n;)(t=pl-n._time)>=0&&n._call.call(null,t),n=n._next;--cl}function Tn(){pl=(hl=vl.now())+dl,cl=sl=0;try{Mn()}finally{cl=0,function(){var t,n,e=Yf,r=1/0;for(;e;)e._call?(r>e._time&&(r=e._time),t=e,e=e._next):(n=e._next,e._next=null,e=t?t._next=n:Yf=n);Bf=t,kn(r)}(),pl=0}}function Nn(){var t=vl.now(),n=t-hl;n>ll&&(dl-=n,hl=t)}function kn(t){if(!cl){sl&&(sl=clearTimeout(sl));t-pl>24?(t<1/0&&(sl=setTimeout(Tn,t-vl.now()-dl)),fl&&(fl=clearInterval(fl))):(fl||(hl=vl.now(),fl=setInterval(Nn,ll)),cl=1,gl(Tn))}}function Sn(t,n,e){var r=new bn;return n=null==n?0:+n,r.restart(function(e){r.stop(),t(e+n)},n,e),r}function En(t,n,e,r,i,o){var u=t.__transition;if(u){if(e in u)return}else t.__transition={};(function(t,n,e){function r(c){var s,f,l,h;if(e.state!==xl)return o();for(s in a)if((h=a[s]).name===e.name){if(h.state===wl)return Sn(r);h.state===Ml?(h.state=Nl,h.timer.stop(),h.on.call("interrupt",t,t.__data__,h.index,h.group),delete a[s]):+s<n&&(h.state=Nl,h.timer.stop(),delete a[s])}if(Sn(function(){e.state===wl&&(e.state=Ml,e.timer.restart(i,e.delay,e.time),i(c))}),e.state=bl,e.on.call("start",t,t.__data__,e.index,e.group),e.state===bl){for(e.state=wl,u=new Array(l=e.tween.length),s=0,f=-1;s<l;++s)(h=e.tween[s].value.call(t,t.__data__,e.index,e.group))&&(u[++f]=h);u.length=f+1}}function i(n){for(var r=n<e.duration?e.ease.call(null,n/e.duration):(e.timer.restart(o),e.state=Tl,1),i=-1,a=u.length;++i<a;)u[i].call(null,r);e.state===Tl&&(e.on.call("end",t,t.__data__,e.index,e.group),o())}function o(){e.state=Nl,e.timer.stop(),delete a[n];for(var r in a)return;delete t.__transition}var u,a=t.__transition;a[n]=e,e.timer=wn(function(t){e.state=xl,e.timer.restart(r,e.delay,e.time),e.delay<=t&&r(t-e.delay)},0,e.time)})(t,e,{name:n,index:r,group:i,on:_l,tween:yl,time:o.time,delay:o.delay,duration:o.duration,ease:o.ease,timer:null,state:ml})}function An(t,n){var e=zn(t,n);if(e.state>ml)throw new Error("too late; already scheduled");return e}function Cn(t,n){var e=zn(t,n);if(e.state>bl)throw new Error("too late; already started");return e}function zn(t,n){var e=t.__transition;if(!e||!(e=e[n]))throw new Error("transition not found");return e}function Pn(t,n){var e,r,i,o=t.__transition,u=!0;if(o){n=null==n?null:n+"";for(i in o)(e=o[i]).name===n?(r=e.state>bl&&e.state<Tl,e.state=Nl,e.timer.stop(),r&&e.on.call("interrupt",t,t.__data__,e.index,e.group),delete o[i]):u=!1;u&&delete t.__transition}}function Rn(t,n,e){var r=t._id;return t.each(function(){var t=Cn(this,r);(t.value||(t.value={}))[n]=e.apply(this,arguments)}),function(t){return zn(t,r).value[n]}}function Ln(t,n){var e;return("number"==typeof n?an:n instanceof Et?Hf:(e=Et(n))?(n=e,Hf):sn)(t,n)}function qn(t,n,e,r){this._groups=t,this._parents=n,this._name=e,this._id=r}function Dn(t){return at().transition(t)}function Un(){return++Sl}function On(t){return((t*=2)<=1?t*t:--t*(2-t)+1)/2}function Fn(t){return((t*=2)<=1?t*t*t:(t-=2)*t*t+2)/2}function In(t){return(1-Math.cos(Pl*t))/2}function Yn(t){return((t*=2)<=1?Math.pow(2,10*t-10):2-Math.pow(2,10-10*t))/2}function Bn(t){return((t*=2)<=1?1-Math.sqrt(1-t*t):Math.sqrt(1-(t-=2)*t)+1)/2}function Hn(t){return(t=+t)<Ll?Hl*t*t:t<Dl?Hl*(t-=ql)*t+Ul:t<Fl?Hl*(t-=Ol)*t+Il:Hl*(t-=Yl)*t+Bl}function jn(t,n){for(var e;!(e=t.__transition)||!(e=e[n]);)if(!(t=t.parentNode))return Ql.time=mn(),Ql;return e}function Xn(t){return function(){return t}}function Vn(){t.event.stopImmediatePropagation()}function $n(){t.event.preventDefault(),t.event.stopImmediatePropagation()}function Wn(t){return{type:t}}function Zn(){return!t.event.button}function Gn(){var t=this.ownerSVGElement||this;return[[0,0],[t.width.baseVal.value,t.height.baseVal.value]]}function Qn(t){for(;!t.__brush;)if(!(t=t.parentNode))return;return t.__brush}function Jn(t){return t[0][0]===t[1][0]||t[0][1]===t[1][1]}function Kn(n){function e(t){var e=t.property("__brush",a).selectAll(".overlay").data([Wn("overlay")]);e.enter().append("rect").attr("class","overlay").attr("pointer-events","all").attr("cursor",uh.overlay).merge(e).each(function(){var t=Qn(this).extent;ct(this).attr("x",t[0][0]).attr("y",t[0][1]).attr("width",t[1][0]-t[0][0]).attr("height",t[1][1]-t[0][1])}),t.selectAll(".selection").data([Wn("selection")]).enter().append("rect").attr("class","selection").attr("cursor",uh.selection).attr("fill","#777").attr("fill-opacity",.3).attr("stroke","#fff").attr("shape-rendering","crispEdges");var i=t.selectAll(".handle").data(n.handles,function(t){return t.type});i.exit().remove(),i.enter().append("rect").attr("class",function(t){return"handle handle--"+t.type}).attr("cursor",function(t){return uh[t.type]}),t.each(r).attr("fill","none").attr("pointer-events","all").style("-webkit-tap-highlight-color","rgba(0,0,0,0)").on("mousedown.brush touchstart.brush",u)}function r(){var t=ct(this),n=Qn(this).selection;n?(t.selectAll(".selection").style("display",null).attr("x",n[0][0]).attr("y",n[0][1]).attr("width",n[1][0]-n[0][0]).attr("height",n[1][1]-n[0][1]),t.selectAll(".handle").style("display",null).attr("x",function(t){return"e"===t.type[t.type.length-1]?n[1][0]-h/2:n[0][0]-h/2}).attr("y",function(t){return"s"===t.type[0]?n[1][1]-h/2:n[0][1]-h/2}).attr("width",function(t){return"n"===t.type||"s"===t.type?n[1][0]-n[0][0]+h:h}).attr("height",function(t){return"e"===t.type||"w"===t.type?n[1][1]-n[0][1]+h:h})):t.selectAll(".selection,.handle").style("display","none").attr("x",null).attr("y",null).attr("width",null).attr("height",null)}function i(t,n){return t.__brush.emitter||new o(t,n)}function o(t,n){this.that=t,this.args=n,this.state=t.__brush,this.active=0}function u(){function e(){var t=pt(w);!L||x||b||(Math.abs(t[0]-D[0])>Math.abs(t[1]-D[1])?b=!0:x=!0),D=t,m=!0,$n(),o()}function o(){var t;switch(_=D[0]-q[0],y=D[1]-q[1],T){case th:case Kl:N&&(_=Math.max(C-a,Math.min(P-p,_)),s=a+_,d=p+_),k&&(y=Math.max(z-l,Math.min(R-v,y)),h=l+y,g=v+y);break;case nh:N<0?(_=Math.max(C-a,Math.min(P-a,_)),s=a+_,d=p):N>0&&(_=Math.max(C-p,Math.min(P-p,_)),s=a,d=p+_),k<0?(y=Math.max(z-l,Math.min(R-l,y)),h=l+y,g=v):k>0&&(y=Math.max(z-v,Math.min(R-v,y)),h=l,g=v+y);break;case eh:N&&(s=Math.max(C,Math.min(P,a-_*N)),d=Math.max(C,Math.min(P,p+_*N))),k&&(h=Math.max(z,Math.min(R,l-y*k)),g=Math.max(z,Math.min(R,v+y*k)))}d<s&&(N*=-1,t=a,a=p,p=t,t=s,s=d,d=t,M in ah&&F.attr("cursor",uh[M=ah[M]])),g<h&&(k*=-1,t=l,l=v,v=t,t=h,h=g,g=t,M in ch&&F.attr("cursor",uh[M=ch[M]])),S.selection&&(A=S.selection),x&&(s=A[0][0],d=A[1][0]),b&&(h=A[0][1],g=A[1][1]),A[0][0]===s&&A[0][1]===h&&A[1][0]===d&&A[1][1]===g||(S.selection=[[s,h],[d,g]],r.call(w),U.brush())}function u(){if(Vn(),t.event.touches){if(t.event.touches.length)return;c&&clearTimeout(c),c=setTimeout(function(){c=null},500),O.on("touchmove.brush touchend.brush touchcancel.brush",null)}else yt(t.event.view,m),I.on("keydown.brush keyup.brush mousemove.brush mouseup.brush",null);O.attr("pointer-events","all"),F.attr("cursor",uh.overlay),S.selection&&(A=S.selection),Jn(A)&&(S.selection=null,r.call(w)),U.end()}if(t.event.touches){if(t.event.changedTouches.length<t.event.touches.length)return $n()}else if(c)return;if(f.apply(this,arguments)){var a,s,l,h,p,d,v,g,_,y,m,x,b,w=this,M=t.event.target.__data__.type,T="selection"===(t.event.metaKey?M="overlay":M)?Kl:t.event.altKey?eh:nh,N=n===ih?null:sh[M],k=n===rh?null:fh[M],S=Qn(w),E=S.extent,A=S.selection,C=E[0][0],z=E[0][1],P=E[1][0],R=E[1][1],L=N&&k&&t.event.shiftKey,q=pt(w),D=q,U=i(w,arguments).beforestart();"overlay"===M?S.selection=A=[[a=n===ih?C:q[0],l=n===rh?z:q[1]],[p=n===ih?P:a,v=n===rh?R:l]]:(a=A[0][0],l=A[0][1],p=A[1][0],v=A[1][1]),s=a,h=l,d=p,g=v;var O=ct(w).attr("pointer-events","none"),F=O.selectAll(".overlay").attr("cursor",uh[M]);if(t.event.touches)O.on("touchmove.brush",e,!0).on("touchend.brush touchcancel.brush",u,!0);else{var I=ct(t.event.view).on("keydown.brush",function(){switch(t.event.keyCode){case 16:L=N&&k;break;case 18:T===nh&&(N&&(p=d-_*N,a=s+_*N),k&&(v=g-y*k,l=h+y*k),T=eh,o());break;case 32:T!==nh&&T!==eh||(N<0?p=d-_:N>0&&(a=s-_),k<0?v=g-y:k>0&&(l=h-y),T=th,F.attr("cursor",uh.selection),o());break;default:return}$n()},!0).on("keyup.brush",function(){switch(t.event.keyCode){case 16:L&&(x=b=L=!1,o());break;case 18:T===eh&&(N<0?p=d:N>0&&(a=s),k<0?v=g:k>0&&(l=h),T=nh,o());break;case 32:T===th&&(t.event.altKey?(N&&(p=d-_*N,a=s+_*N),k&&(v=g-y*k,l=h+y*k),T=eh):(N<0?p=d:N>0&&(a=s),k<0?v=g:k>0&&(l=h),T=nh),F.attr("cursor",uh[M]),o());break;default:return}$n()},!0).on("mousemove.brush",e,!0).on("mouseup.brush",u,!0);_t(t.event.view)}Vn(),Pn(w),r.call(w),U.start()}}function a(){var t=this.__brush||{selection:null};return t.extent=s.apply(this,arguments),t.dim=n,t}var c,s=Gn,f=Zn,l=N(e,"start","brush","end"),h=6;return e.move=function(t,e){t.selection?t.on("start.brush",function(){i(this,arguments).beforestart().start()}).on("interrupt.brush end.brush",function(){i(this,arguments).end()}).tween("brush",function(){function t(t){u.selection=1===t&&Jn(s)?null:f(t),r.call(o),a.brush()}var o=this,u=o.__brush,a=i(o,arguments),c=u.selection,s=n.input("function"==typeof e?e.apply(this,arguments):e,u.extent),f=fn(c,s);return c&&s?t:t(1)}):t.each(function(){var t=arguments,o=this.__brush,u=n.input("function"==typeof e?e.apply(this,t):e,o.extent),a=i(this,t).beforestart();Pn(this),o.selection=null==u||Jn(u)?null:u,r.call(this),a.start().brush().end()})},o.prototype={beforestart:function(){return 1==++this.active&&(this.state.emitter=this,this.starting=!0),this},start:function(){return this.starting&&(this.starting=!1,this.emit("start")),this},brush:function(){return this.emit("brush"),this},end:function(){return 0==--this.active&&(delete this.state.emitter,this.emit("end")),this},emit:function(t){it(new function(t,n,e){this.target=t,this.type=n,this.selection=e}(e,t,n.output(this.state.selection)),l.apply,l,[t,this.that,this.args])}},e.extent=function(t){return arguments.length?(s="function"==typeof t?t:Xn([[+t[0][0],+t[0][1]],[+t[1][0],+t[1][1]]]),e):s},e.filter=function(t){return arguments.length?(f="function"==typeof t?t:Xn(!!t),e):f},e.handleSize=function(t){return arguments.length?(h=+t,e):h},e.on=function(){var t=l.on.apply(l,arguments);return t===l?e:t},e}function te(t){return function(){return t}}function ne(){this._x0=this._y0=this._x1=this._y1=null,this._=""}function ee(){return new ne}function re(t){return t.source}function ie(t){return t.target}function oe(t){return t.radius}function ue(t){return t.startAngle}function ae(t){return t.endAngle}function ce(){}function se(t,n){var e=new ce;if(t instanceof ce)t.each(function(t,n){e.set(n,t)});else if(Array.isArray(t)){var r,i=-1,o=t.length;if(null==n)for(;++i<o;)e.set(i,t[i]);else for(;++i<o;)e.set(n(r=t[i],i,t),r)}else if(t)for(var u in t)e.set(u,t[u]);return e}function fe(){return{}}function le(t,n,e){t[n]=e}function he(){return se()}function pe(t,n,e){t.set(n,e)}function de(){}function ve(t,n){var e=new de;if(t instanceof de)t.each(function(t){e.add(t)});else if(t){var r=-1,i=t.length;if(null==n)for(;++r<i;)e.add(t[r]);else for(;++r<i;)e.add(n(t[r],r,t))}return e}function ge(t){return new Function("d","return {"+t.map(function(t,n){return JSON.stringify(t)+": d["+n+"]"}).join(",")+"}")}function _e(t){function n(t,n){function e(){if(s)return Mh;if(f)return f=!1,wh;var n,e,r=a;if(t.charCodeAt(r)===Th){for(;a++<u&&t.charCodeAt(a)!==Th||t.charCodeAt(++a)===Th;);return(n=a)>=u?s=!0:(e=t.charCodeAt(a++))===Nh?f=!0:e===kh&&(f=!0,t.charCodeAt(a)===Nh&&++a),t.slice(r+1,n-1).replace(/""/g,'"')}for(;a<u;){if((e=t.charCodeAt(n=a++))===Nh)f=!0;else if(e===kh)f=!0,t.charCodeAt(a)===Nh&&++a;else if(e!==o)continue;return t.slice(r,n)}return s=!0,t.slice(r,u)}var r,i=[],u=t.length,a=0,c=0,s=u<=0,f=!1;for(t.charCodeAt(u-1)===Nh&&--u,t.charCodeAt(u-1)===kh&&--u;(r=e())!==Mh;){for(var l=[];r!==wh&&r!==Mh;)l.push(r),r=e();n&&null==(l=n(l,c++))||i.push(l)}return i}function e(n){return n.map(r).join(t)}function r(t){return null==t?"":i.test(t+="")?'"'+t.replace(/"/g,'""')+'"':t}var i=new RegExp('["'+t+"\n\r]"),o=t.charCodeAt(0);return{parse:function(t,e){var r,i,o=n(t,function(t,n){if(r)return r(t,n-1);i=t,r=e?function(t,n){var e=ge(t);return function(r,i){return n(e(r),i,t)}}(t,e):ge(t)});return o.columns=i||[],o},parseRows:n,format:function(n,e){return null==e&&(e=function(t){var n=Object.create(null),e=[];return t.forEach(function(t){for(var r in t)r in n||e.push(n[r]=r)}),e}(n)),[e.map(r).join(t)].concat(n.map(function(n){return e.map(function(t){return r(n[t])}).join(t)})).join("\n")},formatRows:function(t){return t.map(e).join("\n")}}}function ye(t){return function(){return t}}function me(){return 1e-6*(Math.random()-.5)}function xe(t,n,e,r){if(isNaN(n)||isNaN(e))return t;var i,o,u,a,c,s,f,l,h,p=t._root,d={data:r},v=t._x0,g=t._y0,_=t._x1,y=t._y1;if(!p)return t._root=d,t;for(;p.length;)if((s=n>=(o=(v+_)/2))?v=o:_=o,(f=e>=(u=(g+y)/2))?g=u:y=u,i=p,!(p=p[l=f<<1|s]))return i[l]=d,t;if(a=+t._x.call(null,p.data),c=+t._y.call(null,p.data),n===a&&e===c)return d.next=p,i?i[l]=d:t._root=d,t;do{i=i?i[l]=new Array(4):t._root=new Array(4),(s=n>=(o=(v+_)/2))?v=o:_=o,(f=e>=(u=(g+y)/2))?g=u:y=u}while((l=f<<1|s)==(h=(c>=u)<<1|a>=o));return i[h]=p,i[l]=d,t}function be(t,n,e,r,i){this.node=t,this.x0=n,this.y0=e,this.x1=r,this.y1=i}function we(t){return t[0]}function Me(t){return t[1]}function Te(t,n,e){var r=new Ne(null==n?we:n,null==e?Me:e,NaN,NaN,NaN,NaN);return null==t?r:r.addAll(t)}function Ne(t,n,e,r,i,o){this._x=t,this._y=n,this._x0=e,this._y0=r,this._x1=i,this._y1=o,this._root=void 0}function ke(t){for(var n={data:t.data},e=n;t=t.next;)e=e.next={data:t.data};return n}function Se(t){return t.x+t.vx}function Ee(t){return t.y+t.vy}function Ae(t){return t.index}function Ce(t,n){var e=t.get(n);if(!e)throw new Error("missing: "+n);return e}function ze(t){return t.x}function Pe(t){return t.y}function Re(t,n){if((e=(t=n?t.toExponential(n-1):t.toExponential()).indexOf("e"))<0)return null;var e,r=t.slice(0,e);return[r.length>1?r[0]+r.slice(2):r,+t.slice(e+1)]}function Le(t){return(t=Re(Math.abs(t)))?t[1]:NaN}function qe(t,n){var e=Re(t,n);if(!e)return t+"";var r=e[0],i=e[1];return i<0?"0."+new Array(-i).join("0")+r:r.length>i+1?r.slice(0,i+1)+"."+r.slice(i+1):r+new Array(i-r.length+2).join("0")}function De(t){return new Ue(t)}function Ue(t){if(!(n=Bh.exec(t)))throw new Error("invalid format: "+t);var n,e=n[1]||" ",r=n[2]||">",i=n[3]||"-",o=n[4]||"",u=!!n[5],a=n[6]&&+n[6],c=!!n[7],s=n[8]&&+n[8].slice(1),f=n[9]||"";"n"===f?(c=!0,f="g"):Yh[f]||(f=""),(u||"0"===e&&"="===r)&&(u=!0,e="0",r="="),this.fill=e,this.align=r,this.sign=i,this.symbol=o,this.zero=u,this.width=a,this.comma=c,this.precision=s,this.type=f}function Oe(t){return t}function Fe(t){function n(t){function n(t){var n,r,u,f=g,x=_;if("c"===v)x=y(t)+x,t="";else{var b=(t=+t)<0;if(t=y(Math.abs(t),d),b&&0==+t&&(b=!1),f=(b?"("===s?s:"-":"-"===s||"("===s?"":s)+f,x=("s"===v?jh[8+Oh/3]:"")+x+(b&&"("===s?")":""),m)for(n=-1,r=t.length;++n<r;)if(48>(u=t.charCodeAt(n))||u>57){x=(46===u?i+t.slice(n+1):t.slice(n))+x,t=t.slice(0,n);break}}p&&!l&&(t=e(t,1/0));var w=f.length+t.length+x.length,M=w<h?new Array(h-w+1).join(a):"";switch(p&&l&&(t=e(M+t,M.length?h-x.length:1/0),M=""),c){case"<":t=f+t+x+M;break;case"=":t=f+M+t+x;break;case"^":t=M.slice(0,w=M.length>>1)+f+t+x+M.slice(w);break;default:t=M+f+t+x}return o(t)}var a=(t=De(t)).fill,c=t.align,s=t.sign,f=t.symbol,l=t.zero,h=t.width,p=t.comma,d=t.precision,v=t.type,g="$"===f?r[0]:"#"===f&&/[boxX]/.test(v)?"0"+v.toLowerCase():"",_="$"===f?r[1]:/[%p]/.test(v)?u:"",y=Yh[v],m=!v||/[defgprs%]/.test(v);return d=null==d?v?6:12:/[gprs]/.test(v)?Math.max(1,Math.min(21,d)):Math.max(0,Math.min(20,d)),n.toString=function(){return t+""},n}var e=t.grouping&&t.thousands?function(t,n){return function(e,r){for(var i=e.length,o=[],u=0,a=t[0],c=0;i>0&&a>0&&(c+a+1>r&&(a=Math.max(1,r-c)),o.push(e.substring(i-=a,i+a)),!((c+=a+1)>r));)a=t[u=(u+1)%t.length];return o.reverse().join(n)}}(t.grouping,t.thousands):Oe,r=t.currency,i=t.decimal,o=t.numerals?function(t){return function(n){return n.replace(/[0-9]/g,function(n){return t[+n]})}}(t.numerals):Oe,u=t.percent||"%";return{format:n,formatPrefix:function(t,e){var r=n((t=De(t),t.type="f",t)),i=3*Math.max(-8,Math.min(8,Math.floor(Le(e)/3))),o=Math.pow(10,-i),u=jh[8+i/3];return function(t){return r(o*t)+u}}}}function Ie(n){return Hh=Fe(n),t.format=Hh.format,t.formatPrefix=Hh.formatPrefix,Hh}function Ye(t){return Math.max(0,-Le(Math.abs(t)))}function Be(t,n){return Math.max(0,3*Math.max(-8,Math.min(8,Math.floor(Le(n)/3)))-Le(Math.abs(t)))}function He(t,n){return t=Math.abs(t),n=Math.abs(n)-t,Math.max(0,Le(n)-Le(t))+1}function je(){return new Xe}function Xe(){this.reset()}function Ve(t,n,e){var r=t.s=n+e,i=r-n,o=r-i;t.t=n-o+(e-i)}function $e(t){return t>1?0:t<-1?Np:Math.acos(t)}function We(t){return t>1?kp:t<-1?-kp:Math.asin(t)}function Ze(t){return(t=Fp(t/2))*t}function Ge(){}function Qe(t,n){t&&jp.hasOwnProperty(t.type)&&jp[t.type](t,n)}function Je(t,n,e){var r,i=-1,o=t.length-e;for(n.lineStart();++i<o;)r=t[i],n.point(r[0],r[1],r[2]);n.lineEnd()}function Ke(t,n){var e=-1,r=t.length;for(n.polygonStart();++e<r;)Je(t[e],n,1);n.polygonEnd()}function tr(t,n){t&&Hp.hasOwnProperty(t.type)?Hp[t.type](t,n):Qe(t,n)}function nr(){$p.point=rr}function er(){ir(Xh,Vh)}function rr(t,n){$p.point=ir,Xh=t,Vh=n,$h=t*=Cp,Wh=Lp(n=(n*=Cp)/2+Sp),Zh=Fp(n)}function ir(t,n){n=(n*=Cp)/2+Sp;var e=(t*=Cp)-$h,r=e>=0?1:-1,i=r*e,o=Lp(n),u=Fp(n),a=Zh*u,c=Wh*o+a*Lp(i),s=a*r*Fp(i);Xp.add(Rp(s,c)),$h=t,Wh=o,Zh=u}function or(t){return[Rp(t[1],t[0]),We(t[2])]}function ur(t){var n=t[0],e=t[1],r=Lp(e);return[r*Lp(n),r*Fp(n),Fp(e)]}function ar(t,n){return t[0]*n[0]+t[1]*n[1]+t[2]*n[2]}function cr(t,n){return[t[1]*n[2]-t[2]*n[1],t[2]*n[0]-t[0]*n[2],t[0]*n[1]-t[1]*n[0]]}function sr(t,n){t[0]+=n[0],t[1]+=n[1],t[2]+=n[2]}function fr(t,n){return[t[0]*n,t[1]*n,t[2]*n]}function lr(t){var n=Yp(t[0]*t[0]+t[1]*t[1]+t[2]*t[2]);t[0]/=n,t[1]/=n,t[2]/=n}function hr(t,n){ip.push(op=[Gh=t,Jh=t]),n<Qh&&(Qh=n),n>Kh&&(Kh=n)}function pr(t,n){var e=ur([t*Cp,n*Cp]);if(rp){var r=cr(rp,e),i=cr([r[1],-r[0],0],r);lr(i),i=or(i);var o,u=t-tp,a=u>0?1:-1,c=i[0]*Ap*a,s=zp(u)>180;s^(a*tp<c&&c<a*t)?(o=i[1]*Ap)>Kh&&(Kh=o):(c=(c+360)%360-180,s^(a*tp<c&&c<a*t)?(o=-i[1]*Ap)<Qh&&(Qh=o):(n<Qh&&(Qh=n),n>Kh&&(Kh=n))),s?t<tp?mr(Gh,t)>mr(Gh,Jh)&&(Jh=t):mr(t,Jh)>mr(Gh,Jh)&&(Gh=t):Jh>=Gh?(t<Gh&&(Gh=t),t>Jh&&(Jh=t)):t>tp?mr(Gh,t)>mr(Gh,Jh)&&(Jh=t):mr(t,Jh)>mr(Gh,Jh)&&(Gh=t)}else ip.push(op=[Gh=t,Jh=t]);n<Qh&&(Qh=n),n>Kh&&(Kh=n),rp=e,tp=t}function dr(){Zp.point=pr}function vr(){op[0]=Gh,op[1]=Jh,Zp.point=hr,rp=null}function gr(t,n){if(rp){var e=t-tp;Wp.add(zp(e)>180?e+(e>0?360:-360):e)}else np=t,ep=n;$p.point(t,n),pr(t,n)}function _r(){$p.lineStart()}function yr(){gr(np,ep),$p.lineEnd(),zp(Wp)>Mp&&(Gh=-(Jh=180)),op[0]=Gh,op[1]=Jh,rp=null}function mr(t,n){return(n-=t)<0?n+360:n}function xr(t,n){return t[0]-n[0]}function br(t,n){return t[0]<=t[1]?t[0]<=n&&n<=t[1]:n<t[0]||t[1]<n}function wr(t,n){t*=Cp;var e=Lp(n*=Cp);Mr(e*Lp(t),e*Fp(t),Fp(n))}function Mr(t,n,e){cp+=(t-cp)/++up,sp+=(n-sp)/up,fp+=(e-fp)/up}function Tr(){Gp.point=Nr}function Nr(t,n){t*=Cp;var e=Lp(n*=Cp);mp=e*Lp(t),xp=e*Fp(t),bp=Fp(n),Gp.point=kr,Mr(mp,xp,bp)}function kr(t,n){t*=Cp;var e=Lp(n*=Cp),r=e*Lp(t),i=e*Fp(t),o=Fp(n),u=Rp(Yp((u=xp*o-bp*i)*u+(u=bp*r-mp*o)*u+(u=mp*i-xp*r)*u),mp*r+xp*i+bp*o);ap+=u,lp+=u*(mp+(mp=r)),hp+=u*(xp+(xp=i)),pp+=u*(bp+(bp=o)),Mr(mp,xp,bp)}function Sr(){Gp.point=wr}function Er(){Gp.point=Cr}function Ar(){zr(_p,yp),Gp.point=wr}function Cr(t,n){_p=t,yp=n,t*=Cp,n*=Cp,Gp.point=zr;var e=Lp(n);mp=e*Lp(t),xp=e*Fp(t),bp=Fp(n),Mr(mp,xp,bp)}function zr(t,n){t*=Cp;var e=Lp(n*=Cp),r=e*Lp(t),i=e*Fp(t),o=Fp(n),u=xp*o-bp*i,a=bp*r-mp*o,c=mp*i-xp*r,s=Yp(u*u+a*a+c*c),f=We(s),l=s&&-f/s;dp+=l*u,vp+=l*a,gp+=l*c,ap+=f,lp+=f*(mp+(mp=r)),hp+=f*(xp+(xp=i)),pp+=f*(bp+(bp=o)),Mr(mp,xp,bp)}function Pr(t){return function(){return t}}function Rr(t,n){function e(e,r){return e=t(e,r),n(e[0],e[1])}return t.invert&&n.invert&&(e.invert=function(e,r){return(e=n.invert(e,r))&&t.invert(e[0],e[1])}),e}function Lr(t,n){return[t>Np?t-Ep:t<-Np?t+Ep:t,n]}function qr(t,n,e){return(t%=Ep)?n||e?Rr(Ur(t),Or(n,e)):Ur(t):n||e?Or(n,e):Lr}function Dr(t){return function(n,e){return n+=t,[n>Np?n-Ep:n<-Np?n+Ep:n,e]}}function Ur(t){var n=Dr(t);return n.invert=Dr(-t),n}function Or(t,n){function e(t,n){var e=Lp(n),a=Lp(t)*e,c=Fp(t)*e,s=Fp(n),f=s*r+a*i;return[Rp(c*o-f*u,a*r-s*i),We(f*o+c*u)]}var r=Lp(t),i=Fp(t),o=Lp(n),u=Fp(n);return e.invert=function(t,n){var e=Lp(n),a=Lp(t)*e,c=Fp(t)*e,s=Fp(n),f=s*o-c*u;return[Rp(c*o+s*u,a*r+f*i),We(f*r-a*i)]},e}function Fr(t){function n(n){return n=t(n[0]*Cp,n[1]*Cp),n[0]*=Ap,n[1]*=Ap,n}return t=qr(t[0]*Cp,t[1]*Cp,t.length>2?t[2]*Cp:0),n.invert=function(n){return n=t.invert(n[0]*Cp,n[1]*Cp),n[0]*=Ap,n[1]*=Ap,n},n}function Ir(t,n,e,r,i,o){if(e){var u=Lp(n),a=Fp(n),c=r*e;null==i?(i=n+r*Ep,o=n-c/2):(i=Yr(u,i),o=Yr(u,o),(r>0?i<o:i>o)&&(i+=r*Ep));for(var s,f=i;r>0?f>o:f<o;f-=c)s=or([u,-a*Lp(f),-a*Fp(f)]),t.point(s[0],s[1])}}function Yr(t,n){(n=ur(n))[0]-=t,lr(n);var e=$e(-n[1]);return((-n[2]<0?-e:e)+Ep-Mp)%Ep}function Br(){var t,n=[];return{point:function(n,e){t.push([n,e])},lineStart:function(){n.push(t=[])},lineEnd:Ge,rejoin:function(){n.length>1&&n.push(n.pop().concat(n.shift()))},result:function(){var e=n;return n=[],t=null,e}}}function Hr(t,n){return zp(t[0]-n[0])<Mp&&zp(t[1]-n[1])<Mp}function jr(t,n,e,r){this.x=t,this.z=n,this.o=e,this.e=r,this.v=!1,this.n=this.p=null}function Xr(t,n,e,r,i){var o,u,a=[],c=[];if(t.forEach(function(t){if(!((n=t.length-1)<=0)){var n,e,r=t[0],u=t[n];if(Hr(r,u)){for(i.lineStart(),o=0;o<n;++o)i.point((r=t[o])[0],r[1]);i.lineEnd()}else a.push(e=new jr(r,t,null,!0)),c.push(e.o=new jr(r,null,e,!1)),a.push(e=new jr(u,t,null,!1)),c.push(e.o=new jr(u,null,e,!0))}}),a.length){for(c.sort(n),Vr(a),Vr(c),o=0,u=c.length;o<u;++o)c[o].e=e=!e;for(var s,f,l=a[0];;){for(var h=l,p=!0;h.v;)if((h=h.n)===l)return;s=h.z,i.lineStart();do{if(h.v=h.o.v=!0,h.e){if(p)for(o=0,u=s.length;o<u;++o)i.point((f=s[o])[0],f[1]);else r(h.x,h.n.x,1,i);h=h.n}else{if(p)for(s=h.p.z,o=s.length-1;o>=0;--o)i.point((f=s[o])[0],f[1]);else r(h.x,h.p.x,-1,i);h=h.p}s=(h=h.o).z,p=!p}while(!h.v);i.lineEnd()}}}function Vr(t){if(n=t.length){for(var n,e,r=0,i=t[0];++r<n;)i.n=e=t[r],e.p=i,i=e;i.n=e=t[0],e.p=i}}function $r(t,n){var e=n[0],r=n[1],i=[Fp(e),-Lp(e),0],o=0,u=0;cd.reset();for(var a=0,c=t.length;a<c;++a)if(f=(s=t[a]).length)for(var s,f,l=s[f-1],h=l[0],p=l[1]/2+Sp,d=Fp(p),v=Lp(p),g=0;g<f;++g,h=y,d=x,v=b,l=_){var _=s[g],y=_[0],m=_[1]/2+Sp,x=Fp(m),b=Lp(m),w=y-h,M=w>=0?1:-1,T=M*w,N=T>Np,k=d*x;if(cd.add(Rp(k*M*Fp(T),v*b+k*Lp(T))),o+=N?w+M*Ep:w,N^h>=e^y>=e){var S=cr(ur(l),ur(_));lr(S);var E=cr(i,S);lr(E);var A=(N^w>=0?-1:1)*We(E[2]);(r>A||r===A&&(S[0]||S[1]))&&(u+=N^w>=0?1:-1)}}return(o<-Mp||o<Mp&&cd<-Mp)^1&u}function Wr(t,n,e,r){return function(i){function o(n,e){t(n,e)&&i.point(n,e)}function u(t,n){v.point(t,n)}function a(){x.point=u,v.lineStart()}function c(){x.point=o,v.lineEnd()}function s(t,n){d.push([t,n]),y.point(t,n)}function f(){y.lineStart(),d=[]}function l(){s(d[0][0],d[0][1]),y.lineEnd();var t,n,e,r,o=y.clean(),u=_.result(),a=u.length;if(d.pop(),h.push(d),d=null,a)if(1&o){if(e=u[0],(n=e.length-1)>0){for(m||(i.polygonStart(),m=!0),i.lineStart(),t=0;t<n;++t)i.point((r=e[t])[0],r[1]);i.lineEnd()}}else a>1&&2&o&&u.push(u.pop().concat(u.shift())),p.push(u.filter(Zr))}var h,p,d,v=n(i),_=Br(),y=n(_),m=!1,x={point:o,lineStart:a,lineEnd:c,polygonStart:function(){x.point=s,x.lineStart=f,x.lineEnd=l,p=[],h=[]},polygonEnd:function(){x.point=o,x.lineStart=a,x.lineEnd=c,p=g(p);var t=$r(h,r);p.length?(m||(i.polygonStart(),m=!0),Xr(p,Gr,t,e,i)):t&&(m||(i.polygonStart(),m=!0),i.lineStart(),e(null,null,1,i),i.lineEnd()),m&&(i.polygonEnd(),m=!1),p=h=null},sphere:function(){i.polygonStart(),i.lineStart(),e(null,null,1,i),i.lineEnd(),i.polygonEnd()}};return x}}function Zr(t){return t.length>1}function Gr(t,n){return((t=t.x)[0]<0?t[1]-kp-Mp:kp-t[1])-((n=n.x)[0]<0?n[1]-kp-Mp:kp-n[1])}function Qr(t){function n(t,n){return Lp(t)*Lp(n)>i}function e(t,n,e){var r=[1,0,0],o=cr(ur(t),ur(n)),u=ar(o,o),a=o[0],c=u-a*a;if(!c)return!e&&t;var s=i*u/c,f=-i*a/c,l=cr(r,o),h=fr(r,s);sr(h,fr(o,f));var p=l,d=ar(h,p),v=ar(p,p),g=d*d-v*(ar(h,h)-1);if(!(g<0)){var _=Yp(g),y=fr(p,(-d-_)/v);if(sr(y,h),y=or(y),!e)return y;var m,x=t[0],b=n[0],w=t[1],M=n[1];b<x&&(m=x,x=b,b=m);var T=b-x,N=zp(T-Np)<Mp;if(!N&&M<w&&(m=w,w=M,M=m),N||T<Mp?N?w+M>0^y[1]<(zp(y[0]-x)<Mp?w:M):w<=y[1]&&y[1]<=M:T>Np^(x<=y[0]&&y[0]<=b)){var k=fr(p,(-d+_)/v);return sr(k,h),[y,or(k)]}}}function r(n,e){var r=u?t:Np-t,i=0;return n<-r?i|=1:n>r&&(i|=2),e<-r?i|=4:e>r&&(i|=8),i}var i=Lp(t),o=6*Cp,u=i>0,a=zp(i)>Mp;return Wr(n,function(t){var i,o,c,s,f;return{lineStart:function(){s=c=!1,f=1},point:function(l,h){var p,d=[l,h],v=n(l,h),g=u?v?0:r(l,h):v?r(l+(l<0?Np:-Np),h):0;if(!i&&(s=c=v)&&t.lineStart(),v!==c&&(!(p=e(i,d))||Hr(i,p)||Hr(d,p))&&(d[0]+=Mp,d[1]+=Mp,v=n(d[0],d[1])),v!==c)f=0,v?(t.lineStart(),p=e(d,i),t.point(p[0],p[1])):(p=e(i,d),t.point(p[0],p[1]),t.lineEnd()),i=p;else if(a&&i&&u^v){var _;g&o||!(_=e(d,i,!0))||(f=0,u?(t.lineStart(),t.point(_[0][0],_[0][1]),t.point(_[1][0],_[1][1]),t.lineEnd()):(t.point(_[1][0],_[1][1]),t.lineEnd(),t.lineStart(),t.point(_[0][0],_[0][1])))}!v||i&&Hr(i,d)||t.point(d[0],d[1]),i=d,c=v,o=g},lineEnd:function(){c&&t.lineEnd(),i=null},clean:function(){return f|(s&&c)<<1}}},function(n,e,r,i){Ir(i,t,o,r,n,e)},u?[0,-t]:[-Np,t-Np])}function Jr(t,n,e,r){function i(i,o){return t<=i&&i<=e&&n<=o&&o<=r}function o(i,o,a,s){var f=0,l=0;if(null==i||(f=u(i,a))!==(l=u(o,a))||c(i,o)<0^a>0)do{s.point(0===f||3===f?t:e,f>1?r:n)}while((f=(f+a+4)%4)!==l);else s.point(o[0],o[1])}function u(r,i){return zp(r[0]-t)<Mp?i>0?0:3:zp(r[0]-e)<Mp?i>0?2:1:zp(r[1]-n)<Mp?i>0?1:0:i>0?3:2}function a(t,n){return c(t.x,n.x)}function c(t,n){var e=u(t,1),r=u(n,1);return e!==r?e-r:0===e?n[1]-t[1]:1===e?t[0]-n[0]:2===e?t[1]-n[1]:n[0]-t[0]}return function(u){function c(t,n){i(t,n)&&w.point(t,n)}function s(o,u){var a=i(o,u);if(l&&h.push([o,u]),x)p=o,d=u,v=a,x=!1,a&&(w.lineStart(),w.point(o,u));else if(a&&m)w.point(o,u);else{var c=[_=Math.max(ld,Math.min(fd,_)),y=Math.max(ld,Math.min(fd,y))],s=[o=Math.max(ld,Math.min(fd,o)),u=Math.max(ld,Math.min(fd,u))];!function(t,n,e,r,i,o){var u,a=t[0],c=t[1],s=0,f=1,l=n[0]-a,h=n[1]-c;if(u=e-a,l||!(u>0)){if(u/=l,l<0){if(u<s)return;u<f&&(f=u)}else if(l>0){if(u>f)return;u>s&&(s=u)}if(u=i-a,l||!(u<0)){if(u/=l,l<0){if(u>f)return;u>s&&(s=u)}else if(l>0){if(u<s)return;u<f&&(f=u)}if(u=r-c,h||!(u>0)){if(u/=h,h<0){if(u<s)return;u<f&&(f=u)}else if(h>0){if(u>f)return;u>s&&(s=u)}if(u=o-c,h||!(u<0)){if(u/=h,h<0){if(u>f)return;u>s&&(s=u)}else if(h>0){if(u<s)return;u<f&&(f=u)}return s>0&&(t[0]=a+s*l,t[1]=c+s*h),f<1&&(n[0]=a+f*l,n[1]=c+f*h),!0}}}}}(c,s,t,n,e,r)?a&&(w.lineStart(),w.point(o,u),b=!1):(m||(w.lineStart(),w.point(c[0],c[1])),w.point(s[0],s[1]),a||w.lineEnd(),b=!1)}_=o,y=u,m=a}var f,l,h,p,d,v,_,y,m,x,b,w=u,M=Br(),T={point:c,lineStart:function(){T.point=s,l&&l.push(h=[]),x=!0,m=!1,_=y=NaN},lineEnd:function(){f&&(s(p,d),v&&m&&M.rejoin(),f.push(M.result())),T.point=c,m&&w.lineEnd()},polygonStart:function(){w=M,f=[],l=[],b=!0},polygonEnd:function(){var n=function(){for(var n=0,e=0,i=l.length;e<i;++e)for(var o,u,a=l[e],c=1,s=a.length,f=a[0],h=f[0],p=f[1];c<s;++c)o=h,u=p,h=(f=a[c])[0],p=f[1],u<=r?p>r&&(h-o)*(r-u)>(p-u)*(t-o)&&++n:p<=r&&(h-o)*(r-u)<(p-u)*(t-o)&&--n;return n}(),e=b&&n,i=(f=g(f)).length;(e||i)&&(u.polygonStart(),e&&(u.lineStart(),o(null,null,1,u),u.lineEnd()),i&&Xr(f,a,n,o,u),u.polygonEnd()),w=u,f=l=h=null}};return T}}function Kr(){pd.point=pd.lineEnd=Ge}function ti(t,n){Qp=t*=Cp,Jp=Fp(n*=Cp),Kp=Lp(n),pd.point=ni}function ni(t,n){t*=Cp;var e=Fp(n*=Cp),r=Lp(n),i=zp(t-Qp),o=Lp(i),u=r*Fp(i),a=Kp*e-Jp*r*o,c=Jp*e+Kp*r*o;hd.add(Rp(Yp(u*u+a*a),c)),Qp=t,Jp=e,Kp=r}function ei(t){return hd.reset(),tr(t,pd),+hd}function ri(t,n){return dd[0]=t,dd[1]=n,ei(vd)}function ii(t,n){return!(!t||!_d.hasOwnProperty(t.type))&&_d[t.type](t,n)}function oi(t,n){return 0===ri(t,n)}function ui(t,n){var e=ri(t[0],t[1]);return ri(t[0],n)+ri(n,t[1])<=e+Mp}function ai(t,n){return!!$r(t.map(ci),si(n))}function ci(t){return(t=t.map(si)).pop(),t}function si(t){return[t[0]*Cp,t[1]*Cp]}function fi(t,n,e){var r=f(t,n-Mp,e).concat(n);return function(t){return r.map(function(n){return[t,n]})}}function li(t,n,e){var r=f(t,n-Mp,e).concat(n);return function(t){return r.map(function(n){return[n,t]})}}function hi(){function t(){return{type:"MultiLineString",coordinates:n()}}function n(){return f(qp(o/_)*_,i,_).map(p).concat(f(qp(s/y)*y,c,y).map(d)).concat(f(qp(r/v)*v,e,v).filter(function(t){return zp(t%_)>Mp}).map(l)).concat(f(qp(a/g)*g,u,g).filter(function(t){return zp(t%y)>Mp}).map(h))}var e,r,i,o,u,a,c,s,l,h,p,d,v=10,g=v,_=90,y=360,m=2.5;return t.lines=function(){return n().map(function(t){return{type:"LineString",coordinates:t}})},t.outline=function(){return{type:"Polygon",coordinates:[p(o).concat(d(c).slice(1),p(i).reverse().slice(1),d(s).reverse().slice(1))]}},t.extent=function(n){return arguments.length?t.extentMajor(n).extentMinor(n):t.extentMinor()},t.extentMajor=function(n){return arguments.length?(o=+n[0][0],i=+n[1][0],s=+n[0][1],c=+n[1][1],o>i&&(n=o,o=i,i=n),s>c&&(n=s,s=c,c=n),t.precision(m)):[[o,s],[i,c]]},t.extentMinor=function(n){return arguments.length?(r=+n[0][0],e=+n[1][0],a=+n[0][1],u=+n[1][1],r>e&&(n=r,r=e,e=n),a>u&&(n=a,a=u,u=n),t.precision(m)):[[r,a],[e,u]]},t.step=function(n){return arguments.length?t.stepMajor(n).stepMinor(n):t.stepMinor()},t.stepMajor=function(n){return arguments.length?(_=+n[0],y=+n[1],t):[_,y]},t.stepMinor=function(n){return arguments.length?(v=+n[0],g=+n[1],t):[v,g]},t.precision=function(n){return arguments.length?(m=+n,l=fi(a,u,90),h=li(r,e,m),p=fi(s,c,90),d=li(o,i,m),t):m},t.extentMajor([[-180,-90+Mp],[180,90-Mp]]).extentMinor([[-180,-80-Mp],[180,80+Mp]])}function pi(t){return t}function di(){xd.point=vi}function vi(t,n){xd.point=gi,td=ed=t,nd=rd=n}function gi(t,n){md.add(rd*t-ed*n),ed=t,rd=n}function _i(){gi(td,nd)}function yi(t,n){kd+=t,Sd+=n,++Ed}function mi(){qd.point=xi}function xi(t,n){qd.point=bi,yi(ud=t,ad=n)}function bi(t,n){var e=t-ud,r=n-ad,i=Yp(e*e+r*r);Ad+=i*(ud+t)/2,Cd+=i*(ad+n)/2,zd+=i,yi(ud=t,ad=n)}function wi(){qd.point=yi}function Mi(){qd.point=Ni}function Ti(){ki(id,od)}function Ni(t,n){qd.point=ki,yi(id=ud=t,od=ad=n)}function ki(t,n){var e=t-ud,r=n-ad,i=Yp(e*e+r*r);Ad+=i*(ud+t)/2,Cd+=i*(ad+n)/2,zd+=i,Pd+=(i=ad*t-ud*n)*(ud+t),Rd+=i*(ad+n),Ld+=3*i,yi(ud=t,ad=n)}function Si(t){this._context=t}function Ei(t,n){Bd.point=Ai,Ud=Fd=t,Od=Id=n}function Ai(t,n){Fd-=t,Id-=n,Yd.add(Yp(Fd*Fd+Id*Id)),Fd=t,Id=n}function Ci(){this._string=[]}function zi(t){return"m0,"+t+"a"+t+","+t+" 0 1,1 0,"+-2*t+"a"+t+","+t+" 0 1,1 0,"+2*t+"z"}function Pi(t){return function(n){var e=new Ri;for(var r in t)e[r]=t[r];return e.stream=n,e}}function Ri(){}function Li(t,n,e){var r=t.clipExtent&&t.clipExtent();return t.scale(150).translate([0,0]),null!=r&&t.clipExtent(null),tr(e,t.stream(Nd)),n(Nd.result()),null!=r&&t.clipExtent(r),t}function qi(t,n,e){return Li(t,function(e){var r=n[1][0]-n[0][0],i=n[1][1]-n[0][1],o=Math.min(r/(e[1][0]-e[0][0]),i/(e[1][1]-e[0][1])),u=+n[0][0]+(r-o*(e[1][0]+e[0][0]))/2,a=+n[0][1]+(i-o*(e[1][1]+e[0][1]))/2;t.scale(150*o).translate([u,a])},e)}function Di(t,n,e){return qi(t,[[0,0],n],e)}function Ui(t,n,e){return Li(t,function(e){var r=+n,i=r/(e[1][0]-e[0][0]),o=(r-i*(e[1][0]+e[0][0]))/2,u=-i*e[0][1];t.scale(150*i).translate([o,u])},e)}function Oi(t,n,e){return Li(t,function(e){var r=+n,i=r/(e[1][1]-e[0][1]),o=-i*e[0][0],u=(r-i*(e[1][1]+e[0][1]))/2;t.scale(150*i).translate([o,u])},e)}function Fi(t,n){return+n?function(t,n){function e(r,i,o,u,a,c,s,f,l,h,p,d,v,g){var _=s-r,y=f-i,m=_*_+y*y;if(m>4*n&&v--){var x=u+h,b=a+p,w=c+d,M=Yp(x*x+b*b+w*w),T=We(w/=M),N=zp(zp(w)-1)<Mp||zp(o-l)<Mp?(o+l)/2:Rp(b,x),k=t(N,T),S=k[0],E=k[1],A=S-r,C=E-i,z=y*A-_*C;(z*z/m>n||zp((_*A+y*C)/m-.5)>.3||u*h+a*p+c*d<jd)&&(e(r,i,o,u,a,c,S,E,N,x/=M,b/=M,w,v,g),g.point(S,E),e(S,E,N,x,b,w,s,f,l,h,p,d,v,g))}}return function(n){function r(e,r){e=t(e,r),n.point(e[0],e[1])}function i(){_=NaN,w.point=o,n.lineStart()}function o(r,i){var o=ur([r,i]),u=t(r,i);e(_,y,g,m,x,b,_=u[0],y=u[1],g=r,m=o[0],x=o[1],b=o[2],Hd,n),n.point(_,y)}function u(){w.point=r,n.lineEnd()}function a(){i(),w.point=c,w.lineEnd=s}function c(t,n){o(f=t,n),l=_,h=y,p=m,d=x,v=b,w.point=o}function s(){e(_,y,g,m,x,b,l,h,f,p,d,v,Hd,n),w.lineEnd=u,u()}var f,l,h,p,d,v,g,_,y,m,x,b,w={point:r,lineStart:i,lineEnd:u,polygonStart:function(){n.polygonStart(),w.lineStart=a},polygonEnd:function(){n.polygonEnd(),w.lineStart=i}};return w}}(t,n):function(t){return Pi({point:function(n,e){n=t(n,e),this.stream.point(n[0],n[1])}})}(t)}function Ii(t){return Yi(function(){return t})()}function Yi(t){function n(t){return t=s(t[0]*Cp,t[1]*Cp),[t[0]*v+u,a-t[1]*v]}function e(t,n){return t=o(t,n),[t[0]*v+u,a-t[1]*v]}function r(){s=Rr(c=qr(x,b,w),o);var t=o(y,m);return u=g-t[0]*v,a=_+t[1]*v,i()}function i(){return p=d=null,n}var o,u,a,c,s,f,l,h,p,d,v=150,g=480,_=250,y=0,m=0,x=0,b=0,w=0,M=null,T=sd,N=null,k=pi,S=.5,E=Fi(e,S);return n.stream=function(t){return p&&d===t?p:p=Xd(function(t){return Pi({point:function(n,e){var r=t(n,e);return this.stream.point(r[0],r[1])}})}(c)(T(E(k(d=t)))))},n.preclip=function(t){return arguments.length?(T=t,M=void 0,i()):T},n.postclip=function(t){return arguments.length?(k=t,N=f=l=h=null,i()):k},n.clipAngle=function(t){return arguments.length?(T=+t?Qr(M=t*Cp):(M=null,sd),i()):M*Ap},n.clipExtent=function(t){return arguments.length?(k=null==t?(N=f=l=h=null,pi):Jr(N=+t[0][0],f=+t[0][1],l=+t[1][0],h=+t[1][1]),i()):null==N?null:[[N,f],[l,h]]},n.scale=function(t){return arguments.length?(v=+t,r()):v},n.translate=function(t){return arguments.length?(g=+t[0],_=+t[1],r()):[g,_]},n.center=function(t){return arguments.length?(y=t[0]%360*Cp,m=t[1]%360*Cp,r()):[y*Ap,m*Ap]},n.rotate=function(t){return arguments.length?(x=t[0]%360*Cp,b=t[1]%360*Cp,w=t.length>2?t[2]%360*Cp:0,r()):[x*Ap,b*Ap,w*Ap]},n.precision=function(t){return arguments.length?(E=Fi(e,S=t*t),i()):Yp(S)},n.fitExtent=function(t,e){return qi(n,t,e)},n.fitSize=function(t,e){return Di(n,t,e)},n.fitWidth=function(t,e){return Ui(n,t,e)},n.fitHeight=function(t,e){return Oi(n,t,e)},function(){return o=t.apply(this,arguments),n.invert=o.invert&&function(t){return(t=s.invert((t[0]-u)/v,(a-t[1])/v))&&[t[0]*Ap,t[1]*Ap]},r()}}function Bi(t){var n=0,e=Np/3,r=Yi(t),i=r(n,e);return i.parallels=function(t){return arguments.length?r(n=t[0]*Cp,e=t[1]*Cp):[n*Ap,e*Ap]},i}function Hi(t,n){function e(t,n){var e=Yp(o-2*i*Fp(n))/i;return[e*Fp(t*=i),u-e*Lp(t)]}var r=Fp(t),i=(r+Fp(n))/2;if(zp(i)<Mp)return function(t){function n(t,n){return[t*e,Fp(n)/e]}var e=Lp(t);return n.invert=function(t,n){return[t/e,We(n*e)]},n}(t);var o=1+r*(2*i-r),u=Yp(o)/i;return e.invert=function(t,n){var e=u-n;return[Rp(t,zp(e))/i*Ip(e),We((o-(t*t+e*e)*i*i)/(2*i))]},e}function ji(){return Bi(Hi).scale(155.424).center([0,33.6442])}function Xi(){return ji().parallels([29.5,45.5]).scale(1070).translate([480,250]).rotate([96,0]).center([-.6,38.7])}function Vi(t){return function(n,e){var r=Lp(n),i=Lp(e),o=t(r*i);return[o*i*Fp(n),o*Fp(e)]}}function $i(t){return function(n,e){var r=Yp(n*n+e*e),i=t(r),o=Fp(i),u=Lp(i);return[Rp(n*o,r*u),We(r&&e*o/r)]}}function Wi(t,n){return[t,Up(Bp((kp+n)/2))]}function Zi(t){function n(){var n=Np*a(),u=o(Fr(o.rotate()).invert([0,0]));return s(null==f?[[u[0]-n,u[1]-n],[u[0]+n,u[1]+n]]:t===Wi?[[Math.max(u[0]-n,f),e],[Math.min(u[0]+n,r),i]]:[[f,Math.max(u[1]-n,e)],[r,Math.min(u[1]+n,i)]])}var e,r,i,o=Ii(t),u=o.center,a=o.scale,c=o.translate,s=o.clipExtent,f=null;return o.scale=function(t){return arguments.length?(a(t),n()):a()},o.translate=function(t){return arguments.length?(c(t),n()):c()},o.center=function(t){return arguments.length?(u(t),n()):u()},o.clipExtent=function(t){return arguments.length?(null==t?f=e=r=i=null:(f=+t[0][0],e=+t[0][1],r=+t[1][0],i=+t[1][1]),n()):null==f?null:[[f,e],[r,i]]},n()}function Gi(t){return Bp((kp+t)/2)}function Qi(t,n){function e(t,n){o>0?n<-kp+Mp&&(n=-kp+Mp):n>kp-Mp&&(n=kp-Mp);var e=o/Op(Gi(n),i);return[e*Fp(i*t),o-e*Lp(i*t)]}var r=Lp(t),i=t===n?Fp(t):Up(r/Lp(n))/Up(Gi(n)/Gi(t)),o=r*Op(Gi(t),i)/i;return i?(e.invert=function(t,n){var e=o-n,r=Ip(i)*Yp(t*t+e*e);return[Rp(t,zp(e))/i*Ip(e),2*Pp(Op(o/r,1/i))-kp]},e):Wi}function Ji(t,n){return[t,n]}function Ki(t,n){function e(t,n){var e=o-n,r=i*t;return[e*Fp(r),o-e*Lp(r)]}var r=Lp(t),i=t===n?Fp(t):(r-Lp(n))/(n-t),o=r/i+t;return zp(i)<Mp?Ji:(e.invert=function(t,n){var e=o-n;return[Rp(t,zp(e))/i*Ip(e),o-Ip(i)*Yp(t*t+e*e)]},e)}function to(t,n){var e=Lp(n),r=Lp(t)*e;return[e*Fp(t)/r,Fp(n)/r]}function no(t,n,e,r){return 1===t&&1===n&&0===e&&0===r?pi:Pi({point:function(i,o){this.stream.point(i*t+e,o*n+r)}})}function eo(t,n){var e=n*n,r=e*e;return[t*(.8707-.131979*e+r*(r*(.003971*e-.001529*r)-.013791)),n*(1.007226+e*(.015085+r*(.028874*e-.044475-.005916*r)))]}function ro(t,n){return[Lp(n)*Fp(t),Fp(n)]}function io(t,n){var e=Lp(n),r=1+Lp(t)*e;return[e*Fp(t)/r,Fp(n)/r]}function oo(t,n){return[Up(Bp((kp+n)/2)),-t]}function uo(t,n){return t.parent===n.parent?1:2}function ao(t,n){return t+n.x}function co(t,n){return Math.max(t,n.y)}function so(t){var n=0,e=t.children,r=e&&e.length;if(r)for(;--r>=0;)n+=e[r].value;else n=1;t.value=n}function fo(t,n){var e,r,i,o,u,a=new vo(t),c=+t.value&&(a.value=t.value),s=[a];for(null==n&&(n=lo);e=s.pop();)if(c&&(e.value=+e.data.value),(i=n(e.data))&&(u=i.length))for(e.children=new Array(u),o=u-1;o>=0;--o)s.push(r=e.children[o]=new vo(i[o])),r.parent=e,r.depth=e.depth+1;return a.eachBefore(po)}function lo(t){return t.children}function ho(t){t.data=t.data.data}function po(t){var n=0;do{t.height=n}while((t=t.parent)&&t.height<++n)}function vo(t){this.data=t,this.depth=this.height=0,this.parent=null}function go(t){for(var n,e,r=0,i=(t=function(t){for(var n,e,r=t.length;r;)e=Math.random()*r--|0,n=t[r],t[r]=t[e],t[e]=n;return t}(Wd.call(t))).length,o=[];r<i;)n=t[r],e&&yo(e,n)?++r:(e=function(t){switch(t.length){case 1:return function(t){return{x:t.x,y:t.y,r:t.r}}(t[0]);case 2:return xo(t[0],t[1]);case 3:return bo(t[0],t[1],t[2])}}(o=function(t,n){var e,r;if(mo(n,t))return[n];for(e=0;e<t.length;++e)if(_o(n,t[e])&&mo(xo(t[e],n),t))return[t[e],n];for(e=0;e<t.length-1;++e)for(r=e+1;r<t.length;++r)if(_o(xo(t[e],t[r]),n)&&_o(xo(t[e],n),t[r])&&_o(xo(t[r],n),t[e])&&mo(bo(t[e],t[r],n),t))return[t[e],t[r],n];throw new Error}(o,n)),r=0);return e}function _o(t,n){var e=t.r-n.r,r=n.x-t.x,i=n.y-t.y;return e<0||e*e<r*r+i*i}function yo(t,n){var e=t.r-n.r+1e-6,r=n.x-t.x,i=n.y-t.y;return e>0&&e*e>r*r+i*i}function mo(t,n){for(var e=0;e<n.length;++e)if(!yo(t,n[e]))return!1;return!0}function xo(t,n){var e=t.x,r=t.y,i=t.r,o=n.x,u=n.y,a=n.r,c=o-e,s=u-r,f=a-i,l=Math.sqrt(c*c+s*s);return{x:(e+o+c/l*f)/2,y:(r+u+s/l*f)/2,r:(l+i+a)/2}}function bo(t,n,e){var r=t.x,i=t.y,o=t.r,u=n.x,a=n.y,c=n.r,s=e.x,f=e.y,l=e.r,h=r-u,p=r-s,d=i-a,v=i-f,g=c-o,_=l-o,y=r*r+i*i-o*o,m=y-u*u-a*a+c*c,x=y-s*s-f*f+l*l,b=p*d-h*v,w=(d*x-v*m)/(2*b)-r,M=(v*g-d*_)/b,T=(p*m-h*x)/(2*b)-i,N=(h*_-p*g)/b,k=M*M+N*N-1,S=2*(o+w*M+T*N),E=w*w+T*T-o*o,A=-(k?(S+Math.sqrt(S*S-4*k*E))/(2*k):E/S);return{x:r+w+M*A,y:i+T+N*A,r:A}}function wo(t,n,e){var r=t.x,i=t.y,o=n.r+e.r,u=t.r+e.r,a=n.x-r,c=n.y-i,s=a*a+c*c;if(s){var f=.5+((u*=u)-(o*=o))/(2*s),l=Math.sqrt(Math.max(0,2*o*(u+s)-(u-=s)*u-o*o))/(2*s);e.x=r+f*a+l*c,e.y=i+f*c-l*a}else e.x=r+u,e.y=i}function Mo(t,n){var e=n.x-t.x,r=n.y-t.y,i=t.r+n.r;return i*i-1e-6>e*e+r*r}function To(t){var n=t._,e=t.next._,r=n.r+e.r,i=(n.x*e.r+e.x*n.r)/r,o=(n.y*e.r+e.y*n.r)/r;return i*i+o*o}function No(t){this._=t,this.next=null,this.previous=null}function ko(t){if(!(i=t.length))return 0;var n,e,r,i,o,u,a,c,s,f,l;if(n=t[0],n.x=0,n.y=0,!(i>1))return n.r;if(e=t[1],n.x=-e.r,e.x=n.r,e.y=0,!(i>2))return n.r+e.r;wo(e,n,r=t[2]),n=new No(n),e=new No(e),r=new No(r),n.next=r.previous=e,e.next=n.previous=r,r.next=e.previous=n;t:for(a=3;a<i;++a){wo(n._,e._,r=t[a]),r=new No(r),c=e.next,s=n.previous,f=e._.r,l=n._.r;do{if(f<=l){if(Mo(c._,r._)){e=c,n.next=e,e.previous=n,--a;continue t}f+=c._.r,c=c.next}else{if(Mo(s._,r._)){(n=s).next=e,e.previous=n,--a;continue t}l+=s._.r,s=s.previous}}while(c!==s.next);for(r.previous=n,r.next=e,n.next=e.previous=e=r,o=To(n);(r=r.next)!==e;)(u=To(r))<o&&(n=r,o=u);e=n.next}for(n=[e._],r=e;(r=r.next)!==e;)n.push(r._);for(r=go(n),a=0;a<i;++a)n=t[a],n.x-=r.x,n.y-=r.y;return r.r}function So(t){if("function"!=typeof t)throw new Error;return t}function Eo(){return 0}function Ao(t){return function(){return t}}function Co(t){return Math.sqrt(t.value)}function zo(t){return function(n){n.children||(n.r=Math.max(0,+t(n)||0))}}function Po(t,n){return function(e){if(r=e.children){var r,i,o,u=r.length,a=t(e)*n||0;if(a)for(i=0;i<u;++i)r[i].r+=a;if(o=ko(r),a)for(i=0;i<u;++i)r[i].r-=a;e.r=o+a}}}function Ro(t){return function(n){var e=n.parent;n.r*=t,e&&(n.x=e.x+t*n.x,n.y=e.y+t*n.y)}}function Lo(t){t.x0=Math.round(t.x0),t.y0=Math.round(t.y0),t.x1=Math.round(t.x1),t.y1=Math.round(t.y1)}function qo(t,n,e,r,i){for(var o,u=t.children,a=-1,c=u.length,s=t.value&&(r-n)/t.value;++a<c;)(o=u[a]).y0=e,o.y1=i,o.x0=n,o.x1=n+=o.value*s}function Do(t){return t.id}function Uo(t){return t.parentId}function Oo(t,n){return t.parent===n.parent?1:2}function Fo(t){var n=t.children;return n?n[0]:t.t}function Io(t){var n=t.children;return n?n[n.length-1]:t.t}function Yo(t,n,e){var r=e/(n.i-t.i);n.c-=r,n.s+=e,t.c+=r,n.z+=e,n.m+=e}function Bo(t,n,e){return t.a.parent===n.parent?t.a:e}function Ho(t,n){this._=t,this.parent=null,this.children=null,this.A=null,this.a=this,this.z=0,this.m=0,this.c=0,this.s=0,this.t=null,this.i=n}function jo(t,n,e,r,i){for(var o,u=t.children,a=-1,c=u.length,s=t.value&&(i-e)/t.value;++a<c;)(o=u[a]).x0=n,o.x1=r,o.y0=e,o.y1=e+=o.value*s}function Xo(t,n,e,r,i,o){for(var u,a,c,s,f,l,h,p,d,v,g,_=[],y=n.children,m=0,x=0,b=y.length,w=n.value;m<b;){c=i-e,s=o-r;do{f=y[x++].value}while(!f&&x<b);for(l=h=f,g=f*f*(v=Math.max(s/c,c/s)/(w*t)),d=Math.max(h/g,g/l);x<b;++x){if(f+=a=y[x].value,a<l&&(l=a),a>h&&(h=a),g=f*f*v,(p=Math.max(h/g,g/l))>d){f-=a;break}d=p}_.push(u={value:f,dice:c<s,children:y.slice(m,x)}),u.dice?qo(u,e,r,i,w?r+=s*f/w:o):jo(u,e,r,w?e+=c*f/w:i,o),w-=f,m=x}return _}function Vo(t,n,e){return(n[0]-t[0])*(e[1]-t[1])-(n[1]-t[1])*(e[0]-t[0])}function $o(t,n){return t[0]-n[0]||t[1]-n[1]}function Wo(t){for(var n=t.length,e=[0,1],r=2,i=2;i<n;++i){for(;r>1&&Vo(t[e[r-2]],t[e[r-1]],t[i])<=0;)--r;e[r++]=i}return e.slice(0,r)}function Zo(t){this._size=t,this._call=this._error=null,this._tasks=[],this._data=[],this._waiting=this._active=this._ended=this._start=0}function Go(t){if(!t._start)try{(function(t){for(;t._start=t._waiting&&t._active<t._size;){var n=t._ended+t._active,e=t._tasks[n],r=e.length-1,i=e[r];e[r]=function(t,n){return function(e,r){t._tasks[n]&&(--t._active,++t._ended,t._tasks[n]=null,null==t._error&&(null!=e?Qo(t,e):(t._data[n]=r,t._waiting?Go(t):Jo(t))))}}(t,n),--t._waiting,++t._active,e=i.apply(null,e),t._tasks[n]&&(t._tasks[n]=e||ev)}})(t)}catch(n){if(t._tasks[t._ended+t._active-1])Qo(t,n);else if(!t._data)throw n}}function Qo(t,n){var e,r=t._tasks.length;for(t._error=n,t._data=void 0,t._waiting=NaN;--r>=0;)if((e=t._tasks[r])&&(t._tasks[r]=null,e.abort))try{e.abort()}catch(n){}t._active=NaN,Jo(t)}function Jo(t){if(!t._active&&t._call){var n=t._data;t._data=void 0,t._call(t._error,n)}}function Ko(t){if(null==t)t=1/0;else if(!((t=+t)>=1))throw new Error("invalid concurrency");return new Zo(t)}function tu(){return Math.random()}function nu(t,n){function e(t){var n,e=s.status;if(!e&&function(t){var n=t.responseType;return n&&"text"!==n?t.response:t.responseText}(s)||e>=200&&e<300||304===e){if(o)try{n=o.call(r,s)}catch(t){return void a.call("error",r,t)}else n=s;a.call("load",r,n)}else a.call("error",r,t)}var r,i,o,u,a=N("beforesend","progress","load","error"),c=se(),s=new XMLHttpRequest,f=null,l=null,h=0;if("undefined"==typeof XDomainRequest||"withCredentials"in s||!/^(http(s)?:)?\/\//.test(t)||(s=new XDomainRequest),"onload"in s?s.onload=s.onerror=s.ontimeout=e:s.onreadystatechange=function(t){s.readyState>3&&e(t)},s.onprogress=function(t){a.call("progress",r,t)},r={header:function(t,n){return t=(t+"").toLowerCase(),arguments.length<2?c.get(t):(null==n?c.remove(t):c.set(t,n+""),r)},mimeType:function(t){return arguments.length?(i=null==t?null:t+"",r):i},responseType:function(t){return arguments.length?(u=t,r):u},timeout:function(t){return arguments.length?(h=+t,r):h},user:function(t){return arguments.length<1?f:(f=null==t?null:t+"",r)},password:function(t){return arguments.length<1?l:(l=null==t?null:t+"",r)},response:function(t){return o=t,r},get:function(t,n){return r.send("GET",t,n)},post:function(t,n){return r.send("POST",t,n)},send:function(n,e,o){return s.open(n,t,!0,f,l),null==i||c.has("accept")||c.set("accept",i+",*/*"),s.setRequestHeader&&c.each(function(t,n){s.setRequestHeader(n,t)}),null!=i&&s.overrideMimeType&&s.overrideMimeType(i),null!=u&&(s.responseType=u),h>0&&(s.timeout=h),null==o&&"function"==typeof e&&(o=e,e=null),null!=o&&1===o.length&&(o=function(t){return function(n,e){t(null==n?e:null)}}(o)),null!=o&&r.on("error",o).on("load",function(t){o(null,t)}),a.call("beforesend",r,s),s.send(null==e?null:e),r},abort:function(){return s.abort(),r},on:function(){var t=a.on.apply(a,arguments);return t===a?r:t}},null!=n){if("function"!=typeof n)throw new Error("invalid callback: "+n);return r.get(n)}return r}function eu(t,n){return function(e,r){var i=nu(e).mimeType(t).response(n);if(null!=r){if("function"!=typeof r)throw new Error("invalid callback: "+r);return i.get(r)}return i}}function ru(t,n){return function(e,r,i){arguments.length<3&&(i=r,r=null);var o=nu(e).mimeType(t);return o.row=function(t){return arguments.length?o.response(function(t,n){return function(e){return t(e.responseText,n)}}(n,r=t)):r},o.row(r),i?o.get(i):o}}function iu(t){function n(n){var o=n+"",u=e.get(o);if(!u){if(i!==yv)return i;e.set(o,u=r.push(n))}return t[(u-1)%t.length]}var e=se(),r=[],i=yv;return t=null==t?[]:_v.call(t),n.domain=function(t){if(!arguments.length)return r.slice();r=[],e=se();for(var i,o,u=-1,a=t.length;++u<a;)e.has(o=(i=t[u])+"")||e.set(o,r.push(i));return n},n.range=function(e){return arguments.length?(t=_v.call(e),n):t.slice()},n.unknown=function(t){return arguments.length?(i=t,n):i},n.copy=function(){return iu().domain(r).range(t).unknown(i)},n}function ou(){function t(){var t=i().length,r=u[1]<u[0],h=u[r-0],p=u[1-r];n=(p-h)/Math.max(1,t-c+2*s),a&&(n=Math.floor(n)),h+=(p-h-n*(t-c))*l,e=n*(1-c),a&&(h=Math.round(h),e=Math.round(e));var d=f(t).map(function(t){return h+n*t});return o(r?d.reverse():d)}var n,e,r=iu().unknown(void 0),i=r.domain,o=r.range,u=[0,1],a=!1,c=0,s=0,l=.5;return delete r.unknown,r.domain=function(n){return arguments.length?(i(n),t()):i()},r.range=function(n){return arguments.length?(u=[+n[0],+n[1]],t()):u.slice()},r.rangeRound=function(n){return u=[+n[0],+n[1]],a=!0,t()},r.bandwidth=function(){return e},r.step=function(){return n},r.round=function(n){return arguments.length?(a=!!n,t()):a},r.padding=function(n){return arguments.length?(c=s=Math.max(0,Math.min(1,n)),t()):c},r.paddingInner=function(n){return arguments.length?(c=Math.max(0,Math.min(1,n)),t()):c},r.paddingOuter=function(n){return arguments.length?(s=Math.max(0,Math.min(1,n)),t()):s},r.align=function(n){return arguments.length?(l=Math.max(0,Math.min(1,n)),t()):l},r.copy=function(){return ou().domain(i()).range(u).round(a).paddingInner(c).paddingOuter(s).align(l)},t()}function uu(t){var n=t.copy;return t.padding=t.paddingOuter,delete t.paddingInner,delete t.paddingOuter,t.copy=function(){return uu(n())},t}function au(t){return function(){return t}}function cu(t){return+t}function su(t,n){return(n-=t=+t)?function(e){return(e-t)/n}:au(n)}function fu(t,n,e,r){var i=t[0],o=t[1],u=n[0],a=n[1];return o<i?(i=e(o,i),u=r(a,u)):(i=e(i,o),u=r(u,a)),function(t){return u(i(t))}}function lu(t,n,e,r){var i=Math.min(t.length,n.length)-1,o=new Array(i),u=new Array(i),a=-1;for(t[i]<t[0]&&(t=t.slice().reverse(),n=n.slice().reverse());++a<i;)o[a]=e(t[a],t[a+1]),u[a]=r(n[a],n[a+1]);return function(n){var e=Os(t,n,1,i)-1;return u[e](o[e](n))}}function hu(t,n){return n.domain(t.domain()).range(t.range()).interpolate(t.interpolate()).clamp(t.clamp())}function pu(t,n){function e(){return i=Math.min(a.length,c.length)>2?lu:fu,o=u=null,r}function r(n){return(o||(o=i(a,c,f?function(t){return function(n,e){var r=t(n=+n,e=+e);return function(t){return t<=n?0:t>=e?1:r(t)}}}(t):t,s)))(+n)}var i,o,u,a=mv,c=mv,s=fn,f=!1;return r.invert=function(t){return(u||(u=i(c,a,su,f?function(t){return function(n,e){var r=t(n=+n,e=+e);return function(t){return t<=0?n:t>=1?e:r(t)}}}(n):n)))(+t)},r.domain=function(t){return arguments.length?(a=gv.call(t,cu),e()):a.slice()},r.range=function(t){return arguments.length?(c=_v.call(t),e()):c.slice()},r.rangeRound=function(t){return c=_v.call(t),s=ln,e()},r.clamp=function(t){return arguments.length?(f=!!t,e()):f},r.interpolate=function(t){return arguments.length?(s=t,e()):s},e()}function du(n){var e=n.domain;return n.ticks=function(t){var n=e();return l(n[0],n[n.length-1],null==t?10:t)},n.tickFormat=function(n,r){return function(n,e,r){var i,o=n[0],u=n[n.length-1],a=p(o,u,null==e?10:e);switch((r=De(null==r?",f":r)).type){case"s":var c=Math.max(Math.abs(o),Math.abs(u));return null!=r.precision||isNaN(i=Be(a,c))||(r.precision=i),t.formatPrefix(r,c);case"":case"e":case"g":case"p":case"r":null!=r.precision||isNaN(i=He(a,Math.max(Math.abs(o),Math.abs(u))))||(r.precision=i-("e"===r.type));break;case"f":case"%":null!=r.precision||isNaN(i=Ye(a))||(r.precision=i-2*("%"===r.type))}return t.format(r)}(e(),n,r)},n.nice=function(t){null==t&&(t=10);var r,i=e(),o=0,u=i.length-1,a=i[o],c=i[u];return c<a&&(r=a,a=c,c=r,r=o,o=u,u=r),(r=h(a,c,t))>0?r=h(a=Math.floor(a/r)*r,c=Math.ceil(c/r)*r,t):r<0&&(r=h(a=Math.ceil(a*r)/r,c=Math.floor(c*r)/r,t)),r>0?(i[o]=Math.floor(a/r)*r,i[u]=Math.ceil(c/r)*r,e(i)):r<0&&(i[o]=Math.ceil(a*r)/r,i[u]=Math.floor(c*r)/r,e(i)),n},n}function vu(){var t=pu(su,an);return t.copy=function(){return hu(t,vu())},du(t)}function gu(){function t(t){return+t}var n=[0,1];return t.invert=t,t.domain=t.range=function(e){return arguments.length?(n=gv.call(e,cu),t):n.slice()},t.copy=function(){return gu().domain(n)},du(t)}function _u(t,n){var e,r=0,i=(t=t.slice()).length-1,o=t[r],u=t[i];return u<o&&(e=r,r=i,i=e,e=o,o=u,u=e),t[r]=n.floor(o),t[i]=n.ceil(u),t}function yu(t,n){return(n=Math.log(n/t))?function(e){return Math.log(e/t)/n}:au(n)}function mu(t,n){return t<0?function(e){return-Math.pow(-n,e)*Math.pow(-t,1-e)}:function(e){return Math.pow(n,e)*Math.pow(t,1-e)}}function xu(t){return isFinite(t)?+("1e"+t):t<0?0:t}function bu(t){return 10===t?xu:t===Math.E?Math.exp:function(n){return Math.pow(t,n)}}function wu(t){return t===Math.E?Math.log:10===t&&Math.log10||2===t&&Math.log2||(t=Math.log(t),function(n){return Math.log(n)/t})}function Mu(t){return function(n){return-t(-n)}}function Tu(){function n(){return o=wu(i),u=bu(i),r()[0]<0&&(o=Mu(o),u=Mu(u)),e}var e=pu(yu,mu).domain([1,10]),r=e.domain,i=10,o=wu(10),u=bu(10);return e.base=function(t){return arguments.length?(i=+t,n()):i},e.domain=function(t){return arguments.length?(r(t),n()):r()},e.ticks=function(t){var n,e=r(),a=e[0],c=e[e.length-1];(n=c<a)&&(p=a,a=c,c=p);var s,f,h,p=o(a),d=o(c),v=null==t?10:+t,g=[];if(!(i%1)&&d-p<v){if(p=Math.round(p)-1,d=Math.round(d)+1,a>0){for(;p<d;++p)for(f=1,s=u(p);f<i;++f)if(!((h=s*f)<a)){if(h>c)break;g.push(h)}}else for(;p<d;++p)for(f=i-1,s=u(p);f>=1;--f)if(!((h=s*f)<a)){if(h>c)break;g.push(h)}}else g=l(p,d,Math.min(d-p,v)).map(u);return n?g.reverse():g},e.tickFormat=function(n,r){if(null==r&&(r=10===i?".0e":","),"function"!=typeof r&&(r=t.format(r)),n===1/0)return r;null==n&&(n=10);var a=Math.max(1,i*n/e.ticks().length);return function(t){var n=t/u(Math.round(o(t)));return n*i<i-.5&&(n*=i),n<=a?r(t):""}},e.nice=function(){return r(_u(r(),{floor:function(t){return u(Math.floor(o(t)))},ceil:function(t){return u(Math.ceil(o(t)))}}))},e.copy=function(){return hu(e,Tu().base(i))},e}function Nu(t,n){return t<0?-Math.pow(-t,n):Math.pow(t,n)}function ku(){var t=1,n=pu(function(n,e){return(e=Nu(e,t)-(n=Nu(n,t)))?function(r){return(Nu(r,t)-n)/e}:au(e)},function(n,e){return e=Nu(e,t)-(n=Nu(n,t)),function(r){return Nu(n+e*r,1/t)}}),e=n.domain;return n.exponent=function(n){return arguments.length?(t=+n,e(e())):t},n.copy=function(){return hu(n,ku().exponent(t))},du(n)}function Su(){function t(){var t=0,n=Math.max(1,i.length);for(o=new Array(n-1);++t<n;)o[t-1]=v(r,t/n);return e}function e(t){if(!isNaN(t=+t))return i[Os(o,t)]}var r=[],i=[],o=[];return e.invertExtent=function(t){var n=i.indexOf(t);return n<0?[NaN,NaN]:[n>0?o[n-1]:r[0],n<o.length?o[n]:r[r.length-1]]},e.domain=function(e){if(!arguments.length)return r.slice();r=[];for(var i,o=0,u=e.length;o<u;++o)null==(i=e[o])||isNaN(i=+i)||r.push(i);return r.sort(n),t()},e.range=function(n){return arguments.length?(i=_v.call(n),t()):i.slice()},e.quantiles=function(){return o.slice()},e.copy=function(){return Su().domain(r).range(i)},e}function Eu(){function t(t){if(t<=t)return u[Os(o,t,0,i)]}function n(){var n=-1;for(o=new Array(i);++n<i;)o[n]=((n+1)*r-(n-i)*e)/(i+1);return t}var e=0,r=1,i=1,o=[.5],u=[0,1];return t.domain=function(t){return arguments.length?(e=+t[0],r=+t[1],n()):[e,r]},t.range=function(t){return arguments.length?(i=(u=_v.call(t)).length-1,n()):u.slice()},t.invertExtent=function(t){var n=u.indexOf(t);return n<0?[NaN,NaN]:n<1?[e,o[0]]:n>=i?[o[i-1],r]:[o[n-1],o[n]]},t.copy=function(){return Eu().domain([e,r]).range(u)},du(t)}function Au(){function t(t){if(t<=t)return e[Os(n,t,0,r)]}var n=[.5],e=[0,1],r=1;return t.domain=function(i){return arguments.length?(n=_v.call(i),r=Math.min(n.length,e.length-1),t):n.slice()},t.range=function(i){return arguments.length?(e=_v.call(i),r=Math.min(n.length,e.length-1),t):e.slice()},t.invertExtent=function(t){var r=e.indexOf(t);return[n[r-1],n[r]]},t.copy=function(){return Au().domain(n).range(e)},t}function Cu(t,n,e,r){function i(n){return t(n=new Date(+n)),n}return i.floor=i,i.ceil=function(e){return t(e=new Date(e-1)),n(e,1),t(e),e},i.round=function(t){var n=i(t),e=i.ceil(t);return t-n<e-t?n:e},i.offset=function(t,e){return n(t=new Date(+t),null==e?1:Math.floor(e)),t},i.range=function(e,r,o){var u,a=[];if(e=i.ceil(e),o=null==o?1:Math.floor(o),!(e<r&&o>0))return a;do{a.push(u=new Date(+e)),n(e,o),t(e)}while(u<e&&e<r);return a},i.filter=function(e){return Cu(function(n){if(n>=n)for(;t(n),!e(n);)n.setTime(n-1)},function(t,r){if(t>=t)if(r<0)for(;++r<=0;)for(;n(t,-1),!e(t););else for(;--r>=0;)for(;n(t,1),!e(t););})},e&&(i.count=function(n,r){return xv.setTime(+n),bv.setTime(+r),t(xv),t(bv),Math.floor(e(xv,bv))},i.every=function(t){return t=Math.floor(t),isFinite(t)&&t>0?t>1?i.filter(r?function(n){return r(n)%t==0}:function(n){return i.count(0,n)%t==0}):i:null}),i}function zu(t){return Cu(function(n){n.setDate(n.getDate()-(n.getDay()+7-t)%7),n.setHours(0,0,0,0)},function(t,n){t.setDate(t.getDate()+7*n)},function(t,n){return(n-t-(n.getTimezoneOffset()-t.getTimezoneOffset())*Tv)/Nv})}function Pu(t){return Cu(function(n){n.setUTCDate(n.getUTCDate()-(n.getUTCDay()+7-t)%7),n.setUTCHours(0,0,0,0)},function(t,n){t.setUTCDate(t.getUTCDate()+7*n)},function(t,n){return(n-t)/Nv})}function Ru(t){if(0<=t.y&&t.y<100){var n=new Date(-1,t.m,t.d,t.H,t.M,t.S,t.L);return n.setFullYear(t.y),n}return new Date(t.y,t.m,t.d,t.H,t.M,t.S,t.L)}function Lu(t){if(0<=t.y&&t.y<100){var n=new Date(Date.UTC(-1,t.m,t.d,t.H,t.M,t.S,t.L));return n.setUTCFullYear(t.y),n}return new Date(Date.UTC(t.y,t.m,t.d,t.H,t.M,t.S,t.L))}function qu(t){return{y:t,m:0,d:1,H:0,M:0,S:0,L:0}}function Du(t){function n(t,n){return function(e){var r,i,o,u=[],a=-1,c=0,s=t.length;for(e instanceof Date||(e=new Date(+e));++a<s;)37===t.charCodeAt(a)&&(u.push(t.slice(c,a)),null!=(i=Mg[r=t.charAt(++a)])?r=t.charAt(++a):i="e"===r?" ":"0",(o=n[r])&&(r=o(e,i)),u.push(r),c=a+1);return u.push(t.slice(c,a)),u.join("")}}function e(t,n){return function(e){var i,o,u=qu(1900);if(r(u,t,e+="",0)!=e.length)return null;if("Q"in u)return new Date(u.Q);if("p"in u&&(u.H=u.H%12+12*u.p),"V"in u){if(u.V<1||u.V>53)return null;"w"in u||(u.w=1),"Z"in u?(i=(o=(i=Lu(qu(u.y))).getUTCDay())>4||0===o?og.ceil(i):og(i),i=eg.offset(i,7*(u.V-1)),u.y=i.getUTCFullYear(),u.m=i.getUTCMonth(),u.d=i.getUTCDate()+(u.w+6)%7):(i=(o=(i=n(qu(u.y))).getDay())>4||0===o?qv.ceil(i):qv(i),i=Pv.offset(i,7*(u.V-1)),u.y=i.getFullYear(),u.m=i.getMonth(),u.d=i.getDate()+(u.w+6)%7)}else("W"in u||"U"in u)&&("w"in u||(u.w="u"in u?u.u%7:"W"in u?1:0),o="Z"in u?Lu(qu(u.y)).getUTCDay():n(qu(u.y)).getDay(),u.m=0,u.d="W"in u?(u.w+6)%7+7*u.W-(o+5)%7:u.w+7*u.U-(o+6)%7);return"Z"in u?(u.H+=u.Z/100|0,u.M+=u.Z%100,Lu(u)):n(u)}}function r(t,n,e,r){for(var i,o,u=0,a=n.length,c=e.length;u<a;){if(r>=c)return-1;if(37===(i=n.charCodeAt(u++))){if(i=n.charAt(u++),!(o=T[i in Mg?n.charAt(u++):i])||(r=o(t,e,r))<0)return-1}else if(i!=e.charCodeAt(r++))return-1}return r}var i=t.dateTime,o=t.date,u=t.time,a=t.periods,c=t.days,s=t.shortDays,f=t.months,l=t.shortMonths,h=Fu(a),p=Iu(a),d=Fu(c),v=Iu(c),g=Fu(s),_=Iu(s),y=Fu(f),m=Iu(f),x=Fu(l),b=Iu(l),w={a:function(t){return s[t.getDay()]},A:function(t){return c[t.getDay()]},b:function(t){return l[t.getMonth()]},B:function(t){return f[t.getMonth()]},c:null,d:ua,e:ua,f:la,H:aa,I:ca,j:sa,L:fa,m:ha,M:pa,p:function(t){return a[+(t.getHours()>=12)]},Q:Ya,s:Ba,S:da,u:va,U:ga,V:_a,w:ya,W:ma,x:null,X:null,y:xa,Y:ba,Z:wa,"%":Ia},M={a:function(t){return s[t.getUTCDay()]},A:function(t){return c[t.getUTCDay()]},b:function(t){return l[t.getUTCMonth()]},B:function(t){return f[t.getUTCMonth()]},c:null,d:Ma,e:Ma,f:Ea,H:Ta,I:Na,j:ka,L:Sa,m:Aa,M:Ca,p:function(t){return a[+(t.getUTCHours()>=12)]},Q:Ya,s:Ba,S:za,u:Pa,U:Ra,V:La,w:qa,W:Da,x:null,X:null,y:Ua,Y:Oa,Z:Fa,"%":Ia},T={a:function(t,n,e){var r=g.exec(n.slice(e));return r?(t.w=_[r[0].toLowerCase()],e+r[0].length):-1},A:function(t,n,e){var r=d.exec(n.slice(e));return r?(t.w=v[r[0].toLowerCase()],e+r[0].length):-1},b:function(t,n,e){var r=x.exec(n.slice(e));return r?(t.m=b[r[0].toLowerCase()],e+r[0].length):-1},B:function(t,n,e){var r=y.exec(n.slice(e));return r?(t.m=m[r[0].toLowerCase()],e+r[0].length):-1},c:function(t,n,e){return r(t,i,n,e)},d:Gu,e:Gu,f:ea,H:Ju,I:Ju,j:Qu,L:na,m:Zu,M:Ku,p:function(t,n,e){var r=h.exec(n.slice(e));return r?(t.p=p[r[0].toLowerCase()],e+r[0].length):-1},Q:ia,s:oa,S:ta,u:Bu,U:Hu,V:ju,w:Yu,W:Xu,x:function(t,n,e){return r(t,o,n,e)},X:function(t,n,e){return r(t,u,n,e)},y:$u,Y:Vu,Z:Wu,"%":ra};return w.x=n(o,w),w.X=n(u,w),w.c=n(i,w),M.x=n(o,M),M.X=n(u,M),M.c=n(i,M),{format:function(t){var e=n(t+="",w);return e.toString=function(){return t},e},parse:function(t){var n=e(t+="",Ru);return n.toString=function(){return t},n},utcFormat:function(t){var e=n(t+="",M);return e.toString=function(){return t},e},utcParse:function(t){var n=e(t,Lu);return n.toString=function(){return t},n}}}function Uu(t,n,e){var r=t<0?"-":"",i=(r?-t:t)+"",o=i.length;return r+(o<e?new Array(e-o+1).join(n)+i:i)}function Ou(t){return t.replace(kg,"\\$&")}function Fu(t){return new RegExp("^(?:"+t.map(Ou).join("|")+")","i")}function Iu(t){for(var n={},e=-1,r=t.length;++e<r;)n[t[e].toLowerCase()]=e;return n}function Yu(t,n,e){var r=Tg.exec(n.slice(e,e+1));return r?(t.w=+r[0],e+r[0].length):-1}function Bu(t,n,e){var r=Tg.exec(n.slice(e,e+1));return r?(t.u=+r[0],e+r[0].length):-1}function Hu(t,n,e){var r=Tg.exec(n.slice(e,e+2));return r?(t.U=+r[0],e+r[0].length):-1}function ju(t,n,e){var r=Tg.exec(n.slice(e,e+2));return r?(t.V=+r[0],e+r[0].length):-1}function Xu(t,n,e){var r=Tg.exec(n.slice(e,e+2));return r?(t.W=+r[0],e+r[0].length):-1}function Vu(t,n,e){var r=Tg.exec(n.slice(e,e+4));return r?(t.y=+r[0],e+r[0].length):-1}function $u(t,n,e){var r=Tg.exec(n.slice(e,e+2));return r?(t.y=+r[0]+(+r[0]>68?1900:2e3),e+r[0].length):-1}function Wu(t,n,e){var r=/^(Z)|([+-]\d\d)(?::?(\d\d))?/.exec(n.slice(e,e+6));return r?(t.Z=r[1]?0:-(r[2]+(r[3]||"00")),e+r[0].length):-1}function Zu(t,n,e){var r=Tg.exec(n.slice(e,e+2));return r?(t.m=r[0]-1,e+r[0].length):-1}function Gu(t,n,e){var r=Tg.exec(n.slice(e,e+2));return r?(t.d=+r[0],e+r[0].length):-1}function Qu(t,n,e){var r=Tg.exec(n.slice(e,e+3));return r?(t.m=0,t.d=+r[0],e+r[0].length):-1}function Ju(t,n,e){var r=Tg.exec(n.slice(e,e+2));return r?(t.H=+r[0],e+r[0].length):-1}function Ku(t,n,e){var r=Tg.exec(n.slice(e,e+2));return r?(t.M=+r[0],e+r[0].length):-1}function ta(t,n,e){var r=Tg.exec(n.slice(e,e+2));return r?(t.S=+r[0],e+r[0].length):-1}function na(t,n,e){var r=Tg.exec(n.slice(e,e+3));return r?(t.L=+r[0],e+r[0].length):-1}function ea(t,n,e){var r=Tg.exec(n.slice(e,e+6));return r?(t.L=Math.floor(r[0]/1e3),e+r[0].length):-1}function ra(t,n,e){var r=Ng.exec(n.slice(e,e+1));return r?e+r[0].length:-1}function ia(t,n,e){var r=Tg.exec(n.slice(e));return r?(t.Q=+r[0],e+r[0].length):-1}function oa(t,n,e){var r=Tg.exec(n.slice(e));return r?(t.Q=1e3*+r[0],e+r[0].length):-1}function ua(t,n){return Uu(t.getDate(),n,2)}function aa(t,n){return Uu(t.getHours(),n,2)}function ca(t,n){return Uu(t.getHours()%12||12,n,2)}function sa(t,n){return Uu(1+Pv.count(Gv(t),t),n,3)}function fa(t,n){return Uu(t.getMilliseconds(),n,3)}function la(t,n){return fa(t,n)+"000"}function ha(t,n){return Uu(t.getMonth()+1,n,2)}function pa(t,n){return Uu(t.getMinutes(),n,2)}function da(t,n){return Uu(t.getSeconds(),n,2)}function va(t){var n=t.getDay();return 0===n?7:n}function ga(t,n){return Uu(Lv.count(Gv(t),t),n,2)}function _a(t,n){var e=t.getDay();return t=e>=4||0===e?Ov(t):Ov.ceil(t),Uu(Ov.count(Gv(t),t)+(4===Gv(t).getDay()),n,2)}function ya(t){return t.getDay()}function ma(t,n){return Uu(qv.count(Gv(t),t),n,2)}function xa(t,n){return Uu(t.getFullYear()%100,n,2)}function ba(t,n){return Uu(t.getFullYear()%1e4,n,4)}function wa(t){var n=t.getTimezoneOffset();return(n>0?"-":(n*=-1,"+"))+Uu(n/60|0,"0",2)+Uu(n%60,"0",2)}function Ma(t,n){return Uu(t.getUTCDate(),n,2)}function Ta(t,n){return Uu(t.getUTCHours(),n,2)}function Na(t,n){return Uu(t.getUTCHours()%12||12,n,2)}function ka(t,n){return Uu(1+eg.count(xg(t),t),n,3)}function Sa(t,n){return Uu(t.getUTCMilliseconds(),n,3)}function Ea(t,n){return Sa(t,n)+"000"}function Aa(t,n){return Uu(t.getUTCMonth()+1,n,2)}function Ca(t,n){return Uu(t.getUTCMinutes(),n,2)}function za(t,n){return Uu(t.getUTCSeconds(),n,2)}function Pa(t){var n=t.getUTCDay();return 0===n?7:n}function Ra(t,n){return Uu(ig.count(xg(t),t),n,2)}function La(t,n){var e=t.getUTCDay();return t=e>=4||0===e?cg(t):cg.ceil(t),Uu(cg.count(xg(t),t)+(4===xg(t).getUTCDay()),n,2)}function qa(t){return t.getUTCDay()}function Da(t,n){return Uu(og.count(xg(t),t),n,2)}function Ua(t,n){return Uu(t.getUTCFullYear()%100,n,2)}function Oa(t,n){return Uu(t.getUTCFullYear()%1e4,n,4)}function Fa(){return"+0000"}function Ia(){return"%"}function Ya(t){return+t}function Ba(t){return Math.floor(+t/1e3)}function Ha(n){return bg=Du(n),t.timeFormat=bg.format,t.timeParse=bg.parse,t.utcFormat=bg.utcFormat,t.utcParse=bg.utcParse,bg}function ja(t){return new Date(t)}function Xa(t){return t instanceof Date?+t:+new Date(+t)}function Va(t,n,r,i,o,u,a,c,s){function f(e){return(a(e)<e?g:u(e)<e?_:o(e)<e?y:i(e)<e?m:n(e)<e?r(e)<e?x:b:t(e)<e?w:M)(e)}function l(n,r,i,o){if(null==n&&(n=10),"number"==typeof n){var u=Math.abs(i-r)/n,a=e(function(t){return t[2]}).right(T,u);a===T.length?(o=p(r/Dg,i/Dg,n),n=t):a?(o=(a=T[u/T[a-1][2]<T[a][2]/u?a-1:a])[1],n=a[0]):(o=Math.max(p(r,i,n),1),n=c)}return null==o?n:n.every(o)}var h=pu(su,an),d=h.invert,v=h.domain,g=s(".%L"),_=s(":%S"),y=s("%I:%M"),m=s("%I %p"),x=s("%a %d"),b=s("%b %d"),w=s("%B"),M=s("%Y"),T=[[a,1,Cg],[a,5,5*Cg],[a,15,15*Cg],[a,30,30*Cg],[u,1,zg],[u,5,5*zg],[u,15,15*zg],[u,30,30*zg],[o,1,Pg],[o,3,3*Pg],[o,6,6*Pg],[o,12,12*Pg],[i,1,Rg],[i,2,2*Rg],[r,1,Lg],[n,1,qg],[n,3,3*qg],[t,1,Dg]];return h.invert=function(t){return new Date(d(t))},h.domain=function(t){return arguments.length?v(gv.call(t,Xa)):v().map(ja)},h.ticks=function(t,n){var e,r=v(),i=r[0],o=r[r.length-1],u=o<i;return u&&(e=i,i=o,o=e),e=l(t,i,o,n),e=e?e.range(i,o+1):[],u?e.reverse():e},h.tickFormat=function(t,n){return null==n?f:s(n)},h.nice=function(t,n){var e=v();return(t=l(t,e[0],e[e.length-1],n))?v(_u(e,t)):h},h.copy=function(){return hu(h,Va(t,n,r,i,o,u,a,c,s))},h}function $a(t){return t.match(/.{6}/g).map(function(t){return"#"+t})}function Wa(t){var n=t.length;return function(e){return t[Math.max(0,Math.min(n-1,Math.floor(e*n)))]}}function Za(t){function n(n){var o=(n-e)/(r-e);return t(i?Math.max(0,Math.min(1,o)):o)}var e=0,r=1,i=!1;return n.domain=function(t){return arguments.length?(e=+t[0],r=+t[1],n):[e,r]},n.clamp=function(t){return arguments.length?(i=!!t,n):i},n.interpolator=function(e){return arguments.length?(t=e,n):t},n.copy=function(){return Za(t).domain([e,r]).clamp(i)},du(n)}function Ga(t){return function(){return t}}function Qa(t){return t>=1?i_:t<=-1?-i_:Math.asin(t)}function Ja(t){return t.innerRadius}function Ka(t){return t.outerRadius}function tc(t){return t.startAngle}function nc(t){return t.endAngle}function ec(t){return t&&t.padAngle}function rc(t,n,e,r,i,o,u){var a=t-e,c=n-r,s=(u?o:-o)/n_(a*a+c*c),f=s*c,l=-s*a,h=t+f,p=n+l,d=e+f,v=r+l,g=(h+d)/2,_=(p+v)/2,y=d-h,m=v-p,x=y*y+m*m,b=i-o,w=h*v-d*p,M=(m<0?-1:1)*n_(Jg(0,b*b*x-w*w)),T=(w*m-y*M)/x,N=(-w*y-m*M)/x,k=(w*m+y*M)/x,S=(-w*y+m*M)/x,E=T-g,A=N-_,C=k-g,z=S-_;return E*E+A*A>C*C+z*z&&(T=k,N=S),{cx:T,cy:N,x01:-f,y01:-l,x11:T*(i/b-1),y11:N*(i/b-1)}}function ic(t){this._context=t}function oc(t){return new ic(t)}function uc(t){return t[0]}function ac(t){return t[1]}function cc(){function t(t){var a,c,s,f=t.length,l=!1;for(null==i&&(u=o(s=ee())),a=0;a<=f;++a)!(a<f&&r(c=t[a],a,t))===l&&((l=!l)?u.lineStart():u.lineEnd()),l&&u.point(+n(c,a,t),+e(c,a,t));if(s)return u=null,s+""||null}var n=uc,e=ac,r=Ga(!0),i=null,o=oc,u=null;return t.x=function(e){return arguments.length?(n="function"==typeof e?e:Ga(+e),t):n},t.y=function(n){return arguments.length?(e="function"==typeof n?n:Ga(+n),t):e},t.defined=function(n){return arguments.length?(r="function"==typeof n?n:Ga(!!n),t):r},t.curve=function(n){return arguments.length?(o=n,null!=i&&(u=o(i)),t):o},t.context=function(n){return arguments.length?(null==n?i=u=null:u=o(i=n),t):i},t}function sc(){function t(t){var n,f,l,h,p,d=t.length,v=!1,g=new Array(d),_=new Array(d);for(null==a&&(s=c(p=ee())),n=0;n<=d;++n){if(!(n<d&&u(h=t[n],n,t))===v)if(v=!v)f=n,s.areaStart(),s.lineStart();else{for(s.lineEnd(),s.lineStart(),l=n-1;l>=f;--l)s.point(g[l],_[l]);s.lineEnd(),s.areaEnd()}v&&(g[n]=+e(h,n,t),_[n]=+i(h,n,t),s.point(r?+r(h,n,t):g[n],o?+o(h,n,t):_[n]))}if(p)return s=null,p+""||null}function n(){return cc().defined(u).curve(c).context(a)}var e=uc,r=null,i=Ga(0),o=ac,u=Ga(!0),a=null,c=oc,s=null;return t.x=function(n){return arguments.length?(e="function"==typeof n?n:Ga(+n),r=null,t):e},t.x0=function(n){return arguments.length?(e="function"==typeof n?n:Ga(+n),t):e},t.x1=function(n){return arguments.length?(r=null==n?null:"function"==typeof n?n:Ga(+n),t):r},t.y=function(n){return arguments.length?(i="function"==typeof n?n:Ga(+n),o=null,t):i},t.y0=function(n){return arguments.length?(i="function"==typeof n?n:Ga(+n),t):i},t.y1=function(n){return arguments.length?(o=null==n?null:"function"==typeof n?n:Ga(+n),t):o},t.lineX0=t.lineY0=function(){return n().x(e).y(i)},t.lineY1=function(){return n().x(e).y(o)},t.lineX1=function(){return n().x(r).y(i)},t.defined=function(n){return arguments.length?(u="function"==typeof n?n:Ga(!!n),t):u},t.curve=function(n){return arguments.length?(c=n,null!=a&&(s=c(a)),t):c},t.context=function(n){return arguments.length?(null==n?a=s=null:s=c(a=n),t):a},t}function fc(t,n){return n<t?-1:n>t?1:n>=t?0:NaN}function lc(t){return t}function hc(t){this._curve=t}function pc(t){function n(n){return new hc(t(n))}return n._curve=t,n}function dc(t){var n=t.curve;return t.angle=t.x,delete t.x,t.radius=t.y,delete t.y,t.curve=function(t){return arguments.length?n(pc(t)):n()._curve},t}function vc(){return dc(cc().curve(u_))}function gc(){var t=sc().curve(u_),n=t.curve,e=t.lineX0,r=t.lineX1,i=t.lineY0,o=t.lineY1;return t.angle=t.x,delete t.x,t.startAngle=t.x0,delete t.x0,t.endAngle=t.x1,delete t.x1,t.radius=t.y,delete t.y,t.innerRadius=t.y0,delete t.y0,t.outerRadius=t.y1,delete t.y1,t.lineStartAngle=function(){return dc(e())},delete t.lineX0,t.lineEndAngle=function(){return dc(r())},delete t.lineX1,t.lineInnerRadius=function(){return dc(i())},delete t.lineY0,t.lineOuterRadius=function(){return dc(o())},delete t.lineY1,t.curve=function(t){return arguments.length?n(pc(t)):n()._curve},t}function _c(t,n){return[(n=+n)*Math.cos(t-=Math.PI/2),n*Math.sin(t)]}function yc(t){return t.source}function mc(t){return t.target}function xc(t){function n(){var n,a=a_.call(arguments),c=e.apply(this,a),s=r.apply(this,a);if(u||(u=n=ee()),t(u,+i.apply(this,(a[0]=c,a)),+o.apply(this,a),+i.apply(this,(a[0]=s,a)),+o.apply(this,a)),n)return u=null,n+""||null}var e=yc,r=mc,i=uc,o=ac,u=null;return n.source=function(t){return arguments.length?(e=t,n):e},n.target=function(t){return arguments.length?(r=t,n):r},n.x=function(t){return arguments.length?(i="function"==typeof t?t:Ga(+t),n):i},n.y=function(t){return arguments.length?(o="function"==typeof t?t:Ga(+t),n):o},n.context=function(t){return arguments.length?(u=null==t?null:t,n):u},n}function bc(t,n,e,r,i){t.moveTo(n,e),t.bezierCurveTo(n=(n+r)/2,e,n,i,r,i)}function wc(t,n,e,r,i){t.moveTo(n,e),t.bezierCurveTo(n,e=(e+i)/2,r,e,r,i)}function Mc(t,n,e,r,i){var o=_c(n,e),u=_c(n,e=(e+i)/2),a=_c(r,e),c=_c(r,i);t.moveTo(o[0],o[1]),t.bezierCurveTo(u[0],u[1],a[0],a[1],c[0],c[1])}function Tc(){}function Nc(t,n,e){t._context.bezierCurveTo((2*t._x0+t._x1)/3,(2*t._y0+t._y1)/3,(t._x0+2*t._x1)/3,(t._y0+2*t._y1)/3,(t._x0+4*t._x1+n)/6,(t._y0+4*t._y1+e)/6)}function kc(t){this._context=t}function Sc(t){this._context=t}function Ec(t){this._context=t}function Ac(t,n){this._basis=new kc(t),this._beta=n}function Cc(t,n,e){t._context.bezierCurveTo(t._x1+t._k*(t._x2-t._x0),t._y1+t._k*(t._y2-t._y0),t._x2+t._k*(t._x1-n),t._y2+t._k*(t._y1-e),t._x2,t._y2)}function zc(t,n){this._context=t,this._k=(1-n)/6}function Pc(t,n){this._context=t,this._k=(1-n)/6}function Rc(t,n){this._context=t,this._k=(1-n)/6}function Lc(t,n,e){var r=t._x1,i=t._y1,o=t._x2,u=t._y2;if(t._l01_a>e_){var a=2*t._l01_2a+3*t._l01_a*t._l12_a+t._l12_2a,c=3*t._l01_a*(t._l01_a+t._l12_a);r=(r*a-t._x0*t._l12_2a+t._x2*t._l01_2a)/c,i=(i*a-t._y0*t._l12_2a+t._y2*t._l01_2a)/c}if(t._l23_a>e_){var s=2*t._l23_2a+3*t._l23_a*t._l12_a+t._l12_2a,f=3*t._l23_a*(t._l23_a+t._l12_a);o=(o*s+t._x1*t._l23_2a-n*t._l12_2a)/f,u=(u*s+t._y1*t._l23_2a-e*t._l12_2a)/f}t._context.bezierCurveTo(r,i,o,u,t._x2,t._y2)}function qc(t,n){this._context=t,this._alpha=n}function Dc(t,n){this._context=t,this._alpha=n}function Uc(t,n){this._context=t,this._alpha=n}function Oc(t){this._context=t}function Fc(t){return t<0?-1:1}function Ic(t,n,e){var r=t._x1-t._x0,i=n-t._x1,o=(t._y1-t._y0)/(r||i<0&&-0),u=(e-t._y1)/(i||r<0&&-0),a=(o*i+u*r)/(r+i);return(Fc(o)+Fc(u))*Math.min(Math.abs(o),Math.abs(u),.5*Math.abs(a))||0}function Yc(t,n){var e=t._x1-t._x0;return e?(3*(t._y1-t._y0)/e-n)/2:n}function Bc(t,n,e){var r=t._x0,i=t._y0,o=t._x1,u=t._y1,a=(o-r)/3;t._context.bezierCurveTo(r+a,i+a*n,o-a,u-a*e,o,u)}function Hc(t){this._context=t}function jc(t){this._context=new Xc(t)}function Xc(t){this._context=t}function Vc(t){this._context=t}function $c(t){var n,e,r=t.length-1,i=new Array(r),o=new Array(r),u=new Array(r);for(i[0]=0,o[0]=2,u[0]=t[0]+2*t[1],n=1;n<r-1;++n)i[n]=1,o[n]=4,u[n]=4*t[n]+2*t[n+1];for(i[r-1]=2,o[r-1]=7,u[r-1]=8*t[r-1]+t[r],n=1;n<r;++n)e=i[n]/o[n-1],o[n]-=e,u[n]-=e*u[n-1];for(i[r-1]=u[r-1]/o[r-1],n=r-2;n>=0;--n)i[n]=(u[n]-i[n+1])/o[n];for(o[r-1]=(t[r]+i[r-1])/2,n=0;n<r-1;++n)o[n]=2*t[n+1]-i[n+1];return[i,o]}function Wc(t,n){this._context=t,this._t=n}function Zc(t,n){if((i=t.length)>1)for(var e,r,i,o=1,u=t[n[0]],a=u.length;o<i;++o)for(r=u,u=t[n[o]],e=0;e<a;++e)u[e][1]+=u[e][0]=isNaN(r[e][1])?r[e][0]:r[e][1]}function Gc(t){for(var n=t.length,e=new Array(n);--n>=0;)e[n]=n;return e}function Qc(t,n){return t[n]}function Jc(t){var n=t.map(Kc);return Gc(t).sort(function(t,e){return n[t]-n[e]})}function Kc(t){for(var n,e=0,r=-1,i=t.length;++r<i;)(n=+t[r][1])&&(e+=n);return e}function ts(t){return function(){return t}}function ns(t){return t[0]}function es(t){return t[1]}function rs(){this._=null}function is(t){t.U=t.C=t.L=t.R=t.P=t.N=null}function os(t,n){var e=n,r=n.R,i=e.U;i?i.L===e?i.L=r:i.R=r:t._=r,r.U=i,e.U=r,e.R=r.L,e.R&&(e.R.U=e),r.L=e}function us(t,n){var e=n,r=n.L,i=e.U;i?i.L===e?i.L=r:i.R=r:t._=r,r.U=i,e.U=r,e.L=r.R,e.L&&(e.L.U=e),r.R=e}function as(t){for(;t.L;)t=t.L;return t}function cs(t,n,e,r){var i=[null,null],o=D_.push(i)-1;return i.left=t,i.right=n,e&&fs(i,t,n,e),r&&fs(i,n,t,r),L_[t.index].halfedges.push(o),L_[n.index].halfedges.push(o),i}function ss(t,n,e){var r=[n,e];return r.left=t,r}function fs(t,n,e,r){t[0]||t[1]?t.left===e?t[1]=r:t[0]=r:(t[0]=r,t.left=n,t.right=e)}function ls(t,n,e,r,i){var o,u=t[0],a=t[1],c=u[0],s=u[1],f=0,l=1,h=a[0]-c,p=a[1]-s;if(o=n-c,h||!(o>0)){if(o/=h,h<0){if(o<f)return;o<l&&(l=o)}else if(h>0){if(o>l)return;o>f&&(f=o)}if(o=r-c,h||!(o<0)){if(o/=h,h<0){if(o>l)return;o>f&&(f=o)}else if(h>0){if(o<f)return;o<l&&(l=o)}if(o=e-s,p||!(o>0)){if(o/=p,p<0){if(o<f)return;o<l&&(l=o)}else if(p>0){if(o>l)return;o>f&&(f=o)}if(o=i-s,p||!(o<0)){if(o/=p,p<0){if(o>l)return;o>f&&(f=o)}else if(p>0){if(o<f)return;o<l&&(l=o)}return!(f>0||l<1)||(f>0&&(t[0]=[c+f*h,s+f*p]),l<1&&(t[1]=[c+l*h,s+l*p]),!0)}}}}}function hs(t,n,e,r,i){var o=t[1];if(o)return!0;var u,a,c=t[0],s=t.left,f=t.right,l=s[0],h=s[1],p=f[0],d=f[1],v=(l+p)/2,g=(h+d)/2;if(d===h){if(v<n||v>=r)return;if(l>p){if(c){if(c[1]>=i)return}else c=[v,e];o=[v,i]}else{if(c){if(c[1]<e)return}else c=[v,i];o=[v,e]}}else if(u=(l-p)/(d-h),a=g-u*v,u<-1||u>1)if(l>p){if(c){if(c[1]>=i)return}else c=[(e-a)/u,e];o=[(i-a)/u,i]}else{if(c){if(c[1]<e)return}else c=[(i-a)/u,i];o=[(e-a)/u,e]}else if(h<d){if(c){if(c[0]>=r)return}else c=[n,u*n+a];o=[r,u*r+a]}else{if(c){if(c[0]<n)return}else c=[r,u*r+a];o=[n,u*n+a]}return t[0]=c,t[1]=o,!0}function ps(t,n){var e=t.site,r=n.left,i=n.right;return e===i&&(i=r,r=e),i?Math.atan2(i[1]-r[1],i[0]-r[0]):(e===r?(r=n[1],i=n[0]):(r=n[0],i=n[1]),Math.atan2(r[0]-i[0],i[1]-r[1]))}function ds(t,n){return n[+(n.left!==t.site)]}function vs(t,n){return n[+(n.left===t.site)]}function gs(t){var n=t.P,e=t.N;if(n&&e){var r=n.site,i=t.site,o=e.site;if(r!==o){var u=i[0],a=i[1],c=r[0]-u,s=r[1]-a,f=o[0]-u,l=o[1]-a,h=2*(c*l-s*f);if(!(h>=-I_)){var p=c*c+s*s,d=f*f+l*l,v=(l*p-s*d)/h,g=(c*d-f*p)/h,_=U_.pop()||new function(){is(this),this.x=this.y=this.arc=this.site=this.cy=null};_.arc=t,_.site=i,_.x=v+u,_.y=(_.cy=g+a)+Math.sqrt(v*v+g*g),t.circle=_;for(var y=null,m=q_._;m;)if(_.y<m.y||_.y===m.y&&_.x<=m.x){if(!m.L){y=m.P;break}m=m.L}else{if(!m.R){y=m;break}m=m.R}q_.insert(y,_),y||(P_=_)}}}}function _s(t){var n=t.circle;n&&(n.P||(P_=n.N),q_.remove(n),U_.push(n),is(n),t.circle=null)}function ys(t){var n=O_.pop()||new function(){is(this),this.edge=this.site=this.circle=null};return n.site=t,n}function ms(t){_s(t),R_.remove(t),O_.push(t),is(t)}function xs(t){var n=t.circle,e=n.x,r=n.cy,i=[e,r],o=t.P,u=t.N,a=[t];ms(t);for(var c=o;c.circle&&Math.abs(e-c.circle.x)<F_&&Math.abs(r-c.circle.cy)<F_;)o=c.P,a.unshift(c),ms(c),c=o;a.unshift(c),_s(c);for(var s=u;s.circle&&Math.abs(e-s.circle.x)<F_&&Math.abs(r-s.circle.cy)<F_;)u=s.N,a.push(s),ms(s),s=u;a.push(s),_s(s);var f,l=a.length;for(f=1;f<l;++f)s=a[f],c=a[f-1],fs(s.edge,c.site,s.site,i);c=a[0],(s=a[l-1]).edge=cs(c.site,s.site,null,i),gs(c),gs(s)}function bs(t){for(var n,e,r,i,o=t[0],u=t[1],a=R_._;a;)if((r=ws(a,u)-o)>F_)a=a.L;else{if(!((i=o-function(t,n){var e=t.N;if(e)return ws(e,n);var r=t.site;return r[1]===n?r[0]:1/0}(a,u))>F_)){r>-F_?(n=a.P,e=a):i>-F_?(n=a,e=a.N):n=e=a;break}if(!a.R){n=a;break}a=a.R}(function(t){L_[t.index]={site:t,halfedges:[]}})(t);var c=ys(t);if(R_.insert(n,c),n||e){if(n===e)return _s(n),e=ys(n.site),R_.insert(c,e),c.edge=e.edge=cs(n.site,c.site),gs(n),void gs(e);if(e){_s(n),_s(e);var s=n.site,f=s[0],l=s[1],h=t[0]-f,p=t[1]-l,d=e.site,v=d[0]-f,g=d[1]-l,_=2*(h*g-p*v),y=h*h+p*p,m=v*v+g*g,x=[(g*y-p*m)/_+f,(h*m-v*y)/_+l];fs(e.edge,s,d,x),c.edge=cs(s,t,null,x),e.edge=cs(t,d,null,x),gs(n),gs(e)}else c.edge=cs(n.site,c.site)}}function ws(t,n){var e=t.site,r=e[0],i=e[1],o=i-n;if(!o)return r;var u=t.P;if(!u)return-1/0;var a=(e=u.site)[0],c=e[1],s=c-n;if(!s)return a;var f=a-r,l=1/o-1/s,h=f/s;return l?(-h+Math.sqrt(h*h-2*l*(f*f/(-2*s)-c+s/2+i-o/2)))/l+r:(r+a)/2}function Ms(t,n,e){return(t[0]-e[0])*(n[1]-t[1])-(t[0]-n[0])*(e[1]-t[1])}function Ts(t,n){return n[1]-t[1]||n[0]-t[0]}function Ns(t,n){var e,r,i,o=t.sort(Ts).pop();for(D_=[],L_=new Array(t.length),R_=new rs,q_=new rs;;)if(i=P_,o&&(!i||o[1]<i.y||o[1]===i.y&&o[0]<i.x))o[0]===e&&o[1]===r||(bs(o),e=o[0],r=o[1]),o=t.pop();else{if(!i)break;xs(i.arc)}if(function(){for(var t,n,e,r,i=0,o=L_.length;i<o;++i)if((t=L_[i])&&(r=(n=t.halfedges).length)){var u=new Array(r),a=new Array(r);for(e=0;e<r;++e)u[e]=e,a[e]=ps(t,D_[n[e]]);for(u.sort(function(t,n){return a[n]-a[t]}),e=0;e<r;++e)a[e]=n[u[e]];for(e=0;e<r;++e)n[e]=a[e]}}(),n){var u=+n[0][0],a=+n[0][1],c=+n[1][0],s=+n[1][1];(function(t,n,e,r){for(var i,o=D_.length;o--;)hs(i=D_[o],t,n,e,r)&&ls(i,t,n,e,r)&&(Math.abs(i[0][0]-i[1][0])>F_||Math.abs(i[0][1]-i[1][1])>F_)||delete D_[o]})(u,a,c,s),function(t,n,e,r){var i,o,u,a,c,s,f,l,h,p,d,v,g=L_.length,_=!0;for(i=0;i<g;++i)if(o=L_[i]){for(u=o.site,a=(c=o.halfedges).length;a--;)D_[c[a]]||c.splice(a,1);for(a=0,s=c.length;a<s;)d=(p=vs(o,D_[c[a]]))[0],v=p[1],l=(f=ds(o,D_[c[++a%s]]))[0],h=f[1],(Math.abs(d-l)>F_||Math.abs(v-h)>F_)&&(c.splice(a,0,D_.push(ss(u,p,Math.abs(d-t)<F_&&r-v>F_?[t,Math.abs(l-t)<F_?h:r]:Math.abs(v-r)<F_&&e-d>F_?[Math.abs(h-r)<F_?l:e,r]:Math.abs(d-e)<F_&&v-n>F_?[e,Math.abs(l-e)<F_?h:n]:Math.abs(v-n)<F_&&d-t>F_?[Math.abs(h-n)<F_?l:t,n]:null))-1),++s);s&&(_=!1)}if(_){var y,m,x,b=1/0;for(i=0,_=null;i<g;++i)(o=L_[i])&&(x=(y=(u=o.site)[0]-t)*y+(m=u[1]-n)*m)<b&&(b=x,_=o);if(_){var w=[t,n],M=[t,r],T=[e,r],N=[e,n];_.halfedges.push(D_.push(ss(u=_.site,w,M))-1,D_.push(ss(u,M,T))-1,D_.push(ss(u,T,N))-1,D_.push(ss(u,N,w))-1)}}for(i=0;i<g;++i)(o=L_[i])&&(o.halfedges.length||delete L_[i])}(u,a,c,s)}this.edges=D_,this.cells=L_,R_=q_=D_=L_=null}function ks(t){return function(){return t}}function Ss(t,n,e){this.k=t,this.x=n,this.y=e}function Es(t){return t.__zoom||Y_}function As(){t.event.stopImmediatePropagation()}function Cs(){t.event.preventDefault(),t.event.stopImmediatePropagation()}function zs(){return!t.event.button}function Ps(){var t,n,e=this;return e instanceof SVGElement?(t=(e=e.ownerSVGElement||e).width.baseVal.value,n=e.height.baseVal.value):(t=e.clientWidth,n=e.clientHeight),[[0,0],[t,n]]}function Rs(){return this.__zoom||Y_}function Ls(){return-t.event.deltaY*(t.event.deltaMode?120:1)/500}function qs(){return"ontouchstart"in this}function Ds(t,n,e){var r=t.invertX(n[0][0])-e[0][0],i=t.invertX(n[1][0])-e[1][0],o=t.invertY(n[0][1])-e[0][1],u=t.invertY(n[1][1])-e[1][1];return t.translate(i>r?(r+i)/2:Math.min(0,r)||Math.max(0,i),u>o?(o+u)/2:Math.min(0,o)||Math.max(0,u))}var Us=e(n),Os=Us.right,Fs=Us.left,Is=Array.prototype,Ys=Is.slice,Bs=Is.map,Hs=Math.sqrt(50),js=Math.sqrt(10),Xs=Math.sqrt(2),Vs=Array.prototype.slice,$s=1,Ws=2,Zs=3,Gs=4,Qs=1e-6,Js={value:function(){}};k.prototype=N.prototype={constructor:k,on:function(t,n){var e,r=this._,i=function(t,n){return t.trim().split(/^|\s+/).map(function(t){var e="",r=t.indexOf(".");if(r>=0&&(e=t.slice(r+1),t=t.slice(0,r)),t&&!n.hasOwnProperty(t))throw new Error("unknown type: "+t);return{type:t,name:e}})}(t+"",r),o=-1,u=i.length;{if(!(arguments.length<2)){if(null!=n&&"function"!=typeof n)throw new Error("invalid callback: "+n);for(;++o<u;)if(e=(t=i[o]).type)r[e]=S(r[e],t.name,n);else if(null==n)for(e in r)r[e]=S(r[e],t.name,null);return this}for(;++o<u;)if((e=(t=i[o]).type)&&(e=function(t,n){for(var e,r=0,i=t.length;r<i;++r)if((e=t[r]).name===n)return e.value}(r[e],t.name)))return e}},copy:function(){var t={},n=this._;for(var e in n)t[e]=n[e].slice();return new k(t)},call:function(t,n){if((e=arguments.length-2)>0)for(var e,r,i=new Array(e),o=0;o<e;++o)i[o]=arguments[o+2];if(!this._.hasOwnProperty(t))throw new Error("unknown type: "+t);for(o=0,e=(r=this._[t]).length;o<e;++o)r[o].value.apply(n,i)},apply:function(t,n,e){if(!this._.hasOwnProperty(t))throw new Error("unknown type: "+t);for(var r=this._[t],i=0,o=r.length;i<o;++i)r[i].value.apply(n,e)}};var Ks="http://www.w3.org/1999/xhtml",tf={svg:"http://www.w3.org/2000/svg",xhtml:Ks,xlink:"http://www.w3.org/1999/xlink",xml:"http://www.w3.org/XML/1998/namespace",xmlns:"http://www.w3.org/2000/xmlns/"},nf=function(t){return function(){return this.matches(t)}};if("undefined"!=typeof document){var ef=document.documentElement;if(!ef.matches){var rf=ef.webkitMatchesSelector||ef.msMatchesSelector||ef.mozMatchesSelector||ef.oMatchesSelector;nf=function(t){return function(){return rf.call(this,t)}}}}var of=nf;q.prototype={constructor:q,appendChild:function(t){return this._parent.insertBefore(t,this._next)},insertBefore:function(t,n){return this._parent.insertBefore(t,n)},querySelector:function(t){return this._parent.querySelector(t)},querySelectorAll:function(t){return this._parent.querySelectorAll(t)}};var uf="$";H.prototype={add:function(t){this._names.indexOf(t)<0&&(this._names.push(t),this._node.setAttribute("class",this._names.join(" ")))},remove:function(t){var n=this._names.indexOf(t);n>=0&&(this._names.splice(n,1),this._node.setAttribute("class",this._names.join(" ")))},contains:function(t){return this._names.indexOf(t)>=0}};var af={};if(t.event=null,"undefined"!=typeof document){"onmouseenter"in document.documentElement||(af={mouseenter:"mouseover",mouseleave:"mouseout"})}var cf=[null];ut.prototype=at.prototype={constructor:ut,select:function(t){"function"!=typeof t&&(t=z(t));for(var n=this._groups,e=n.length,r=new Array(e),i=0;i<e;++i)for(var o,u,a=n[i],c=a.length,s=r[i]=new Array(c),f=0;f<c;++f)(o=a[f])&&(u=t.call(o,o.__data__,f,a))&&("__data__"in o&&(u.__data__=o.__data__),s[f]=u);return new ut(r,this._parents)},selectAll:function(t){"function"!=typeof t&&(t=R(t));for(var n=this._groups,e=n.length,r=[],i=[],o=0;o<e;++o)for(var u,a=n[o],c=a.length,s=0;s<c;++s)(u=a[s])&&(r.push(t.call(u,u.__data__,s,a)),i.push(u));return new ut(r,i)},filter:function(t){"function"!=typeof t&&(t=of(t));for(var n=this._groups,e=n.length,r=new Array(e),i=0;i<e;++i)for(var o,u=n[i],a=u.length,c=r[i]=[],s=0;s<a;++s)(o=u[s])&&t.call(o,o.__data__,s,u)&&c.push(o);return new ut(r,this._parents)},data:function(t,n){if(!t)return p=new Array(this.size()),s=-1,this.each(function(t){p[++s]=t}),p;var e=n?U:D,r=this._parents,i=this._groups;"function"!=typeof t&&(t=function(t){return function(){return t}}(t));for(var o=i.length,u=new Array(o),a=new Array(o),c=new Array(o),s=0;s<o;++s){var f=r[s],l=i[s],h=l.length,p=t.call(f,f&&f.__data__,s,r),d=p.length,v=a[s]=new Array(d),g=u[s]=new Array(d);e(f,l,v,g,c[s]=new Array(h),p,n);for(var _,y,m=0,x=0;m<d;++m)if(_=v[m]){for(m>=x&&(x=m+1);!(y=g[x])&&++x<d;);_._next=y||null}}return u=new ut(u,r),u._enter=a,u._exit=c,u},enter:function(){return new ut(this._enter||this._groups.map(L),this._parents)},exit:function(){return new ut(this._exit||this._groups.map(L),this._parents)},merge:function(t){for(var n=this._groups,e=t._groups,r=n.length,i=e.length,o=Math.min(r,i),u=new Array(r),a=0;a<o;++a)for(var c,s=n[a],f=e[a],l=s.length,h=u[a]=new Array(l),p=0;p<l;++p)(c=s[p]||f[p])&&(h[p]=c);for(;a<r;++a)u[a]=n[a];return new ut(u,this._parents)},order:function(){for(var t=this._groups,n=-1,e=t.length;++n<e;)for(var r,i=t[n],o=i.length-1,u=i[o];--o>=0;)(r=i[o])&&(u&&u!==r.nextSibling&&u.parentNode.insertBefore(r,u),u=r);return this},sort:function(t){function n(n,e){return n&&e?t(n.__data__,e.__data__):!n-!e}t||(t=O);for(var e=this._groups,r=e.length,i=new Array(r),o=0;o<r;++o){for(var u,a=e[o],c=a.length,s=i[o]=new Array(c),f=0;f<c;++f)(u=a[f])&&(s[f]=u);s.sort(n)}return new ut(i,this._parents).order()},call:function(){var t=arguments[0];return arguments[0]=this,t.apply(null,arguments),this},nodes:function(){var t=new Array(this.size()),n=-1;return this.each(function(){t[++n]=this}),t},node:function(){for(var t=this._groups,n=0,e=t.length;n<e;++n)for(var r=t[n],i=0,o=r.length;i<o;++i){var u=r[i];if(u)return u}return null},size:function(){var t=0;return this.each(function(){++t}),t},empty:function(){return!this.node()},each:function(t){for(var n=this._groups,e=0,r=n.length;e<r;++e)for(var i,o=n[e],u=0,a=o.length;u<a;++u)(i=o[u])&&t.call(i,i.__data__,u,o);return this},attr:function(t,n){var e=E(t);if(arguments.length<2){var r=this.node();return e.local?r.getAttributeNS(e.space,e.local):r.getAttribute(e)}return this.each((null==n?e.local?function(t){return function(){this.removeAttributeNS(t.space,t.local)}}:function(t){return function(){this.removeAttribute(t)}}:"function"==typeof n?e.local?function(t,n){return function(){var e=n.apply(this,arguments);null==e?this.removeAttributeNS(t.space,t.local):this.setAttributeNS(t.space,t.local,e)}}:function(t,n){return function(){var e=n.apply(this,arguments);null==e?this.removeAttribute(t):this.setAttribute(t,e)}}:e.local?function(t,n){return function(){this.setAttributeNS(t.space,t.local,n)}}:function(t,n){return function(){this.setAttribute(t,n)}})(e,n))},style:function(t,n,e){return arguments.length>1?this.each((null==n?function(t){return function(){this.style.removeProperty(t)}}:"function"==typeof n?function(t,n,e){return function(){var r=n.apply(this,arguments);null==r?this.style.removeProperty(t):this.style.setProperty(t,r,e)}}:function(t,n,e){return function(){this.style.setProperty(t,n,e)}})(t,n,null==e?"":e)):I(this.node(),t)},property:function(t,n){return arguments.length>1?this.each((null==n?function(t){return function(){delete this[t]}}:"function"==typeof n?function(t,n){return function(){var e=n.apply(this,arguments);null==e?delete this[t]:this[t]=e}}:function(t,n){return function(){this[t]=n}})(t,n)):this.node()[t]},classed:function(t,n){var e=Y(t+"");if(arguments.length<2){for(var r=B(this.node()),i=-1,o=e.length;++i<o;)if(!r.contains(e[i]))return!1;return!0}return this.each(("function"==typeof n?function(t,n){return function(){(n.apply(this,arguments)?j:X)(this,t)}}:n?function(t){return function(){j(this,t)}}:function(t){return function(){X(this,t)}})(e,n))},text:function(t){return arguments.length?this.each(null==t?V:("function"==typeof t?function(t){return function(){var n=t.apply(this,arguments);this.textContent=null==n?"":n}}:function(t){return function(){this.textContent=t}})(t)):this.node().textContent},html:function(t){return arguments.length?this.each(null==t?$:("function"==typeof t?function(t){return function(){var n=t.apply(this,arguments);this.innerHTML=null==n?"":n}}:function(t){return function(){this.innerHTML=t}})(t)):this.node().innerHTML},raise:function(){return this.each(W)},lower:function(){return this.each(Z)},append:function(t){var n="function"==typeof t?t:A(t);return this.select(function(){return this.appendChild(n.apply(this,arguments))})},insert:function(t,n){var e="function"==typeof t?t:A(t),r=null==n?G:"function"==typeof n?n:z(n);return this.select(function(){return this.insertBefore(e.apply(this,arguments),r.apply(this,arguments)||null)})},remove:function(){return this.each(Q)},clone:function(t){return this.select(t?K:J)},datum:function(t){return arguments.length?this.property("__data__",t):this.node().__data__},on:function(t,n,e){var r,i,o=function(t){return t.trim().split(/^|\s+/).map(function(t){var n="",e=t.indexOf(".");return e>=0&&(n=t.slice(e+1),t=t.slice(0,e)),{type:t,name:n}})}(t+""),u=o.length;if(!(arguments.length<2)){for(a=n?rt:et,null==e&&(e=!1),r=0;r<u;++r)this.each(a(o[r],n,e));return this}var a=this.node().__on;if(a)for(var c,s=0,f=a.length;s<f;++s)for(r=0,c=a[s];r<u;++r)if((i=o[r]).type===c.type&&i.name===c.name)return c.value},dispatch:function(t,n){return this.each(("function"==typeof n?function(t,n){return function(){return ot(this,t,n.apply(this,arguments))}}:function(t,n){return function(){return ot(this,t,n)}})(t,n))}};var sf=0;ft.prototype=st.prototype={constructor:ft,get:function(t){for(var n=this._;!(n in t);)if(!(t=t.parentNode))return;return t[n]},set:function(t,n){return t[this._]=n},remove:function(t){return this._ in t&&delete t[this._]},toString:function(){return this._}},xt.prototype.on=function(){var t=this._.on.apply(this._,arguments);return t===this._?this:t};var ff="\\s*([+-]?\\d+)\\s*",lf="\\s*([+-]?\\d*\\.?\\d+(?:[eE][+-]?\\d+)?)\\s*",hf="\\s*([+-]?\\d*\\.?\\d+(?:[eE][+-]?\\d+)?)%\\s*",pf=/^#([0-9a-f]{3})$/,df=/^#([0-9a-f]{6})$/,vf=new RegExp("^rgb\\("+[ff,ff,ff]+"\\)$"),gf=new RegExp("^rgb\\("+[hf,hf,hf]+"\\)$"),_f=new RegExp("^rgba\\("+[ff,ff,ff,lf]+"\\)$"),yf=new RegExp("^rgba\\("+[hf,hf,hf,lf]+"\\)$"),mf=new RegExp("^hsl\\("+[lf,hf,hf]+"\\)$"),xf=new RegExp("^hsla\\("+[lf,hf,hf,lf]+"\\)$"),bf={aliceblue:15792383,antiquewhite:16444375,aqua:65535,aquamarine:8388564,azure:15794175,beige:16119260,bisque:16770244,black:0,blanchedalmond:16772045,blue:255,blueviolet:9055202,brown:10824234,burlywood:14596231,cadetblue:6266528,chartreuse:8388352,chocolate:13789470,coral:16744272,cornflowerblue:6591981,cornsilk:16775388,crimson:14423100,cyan:65535,darkblue:139,darkcyan:35723,darkgoldenrod:12092939,darkgray:11119017,darkgreen:25600,darkgrey:11119017,darkkhaki:12433259,darkmagenta:9109643,darkolivegreen:5597999,darkorange:16747520,darkorchid:10040012,darkred:9109504,darksalmon:15308410,darkseagreen:9419919,darkslateblue:4734347,darkslategray:3100495,darkslategrey:3100495,darkturquoise:52945,darkviolet:9699539,deeppink:16716947,deepskyblue:49151,dimgray:6908265,dimgrey:6908265,dodgerblue:2003199,firebrick:11674146,floralwhite:16775920,forestgreen:2263842,fuchsia:16711935,gainsboro:14474460,ghostwhite:16316671,gold:16766720,goldenrod:14329120,gray:8421504,green:32768,greenyellow:11403055,grey:8421504,honeydew:15794160,hotpink:16738740,indianred:13458524,indigo:4915330,ivory:16777200,khaki:15787660,lavender:15132410,lavenderblush:16773365,lawngreen:8190976,lemonchiffon:16775885,lightblue:11393254,lightcoral:15761536,lightcyan:14745599,lightgoldenrodyellow:16448210,lightgray:13882323,lightgreen:9498256,lightgrey:13882323,lightpink:16758465,lightsalmon:16752762,lightseagreen:2142890,lightskyblue:8900346,lightslategray:7833753,lightslategrey:7833753,lightsteelblue:11584734,lightyellow:16777184,lime:65280,limegreen:3329330,linen:16445670,magenta:16711935,maroon:8388608,mediumaquamarine:6737322,mediumblue:205,mediumorchid:12211667,mediumpurple:9662683,mediumseagreen:3978097,mediumslateblue:8087790,mediumspringgreen:64154,mediumturquoise:4772300,mediumvioletred:13047173,midnightblue:1644912,mintcream:16121850,mistyrose:16770273,moccasin:16770229,navajowhite:16768685,navy:128,oldlace:16643558,olive:8421376,olivedrab:7048739,orange:16753920,orangered:16729344,orchid:14315734,palegoldenrod:15657130,palegreen:10025880,paleturquoise:11529966,palevioletred:14381203,papayawhip:16773077,peachpuff:16767673,peru:13468991,pink:16761035,plum:14524637,powderblue:11591910,purple:8388736,rebeccapurple:6697881,red:16711680,rosybrown:12357519,royalblue:4286945,saddlebrown:9127187,salmon:16416882,sandybrown:16032864,seagreen:3050327,seashell:16774638,sienna:10506797,silver:12632256,skyblue:8900331,slateblue:6970061,slategray:7372944,slategrey:7372944,snow:16775930,springgreen:65407,steelblue:4620980,tan:13808780,teal:32896,thistle:14204888,tomato:16737095,turquoise:4251856,violet:15631086,wheat:16113331,white:16777215,whitesmoke:16119285,yellow:16776960,yellowgreen:10145074};Nt(St,Et,{displayable:function(){return this.rgb().displayable()},toString:function(){return this.rgb()+""}}),Nt(Rt,Pt,kt(St,{brighter:function(t){return t=null==t?1/.7:Math.pow(1/.7,t),new Rt(this.r*t,this.g*t,this.b*t,this.opacity)},darker:function(t){return t=null==t?.7:Math.pow(.7,t),new Rt(this.r*t,this.g*t,this.b*t,this.opacity)},rgb:function(){return this},displayable:function(){return 0<=this.r&&this.r<=255&&0<=this.g&&this.g<=255&&0<=this.b&&this.b<=255&&0<=this.opacity&&this.opacity<=1},toString:function(){var t=this.opacity;return(1===(t=isNaN(t)?1:Math.max(0,Math.min(1,t)))?"rgb(":"rgba(")+Math.max(0,Math.min(255,Math.round(this.r)||0))+", "+Math.max(0,Math.min(255,Math.round(this.g)||0))+", "+Math.max(0,Math.min(255,Math.round(this.b)||0))+(1===t?")":", "+t+")")}})),Nt(Dt,qt,kt(St,{brighter:function(t){return t=null==t?1/.7:Math.pow(1/.7,t),new Dt(this.h,this.s,this.l*t,this.opacity)},darker:function(t){return t=null==t?.7:Math.pow(.7,t),new Dt(this.h,this.s,this.l*t,this.opacity)},rgb:function(){var t=this.h%360+360*(this.h<0),n=isNaN(t)||isNaN(this.s)?0:this.s,e=this.l,r=e+(e<.5?e:1-e)*n,i=2*e-r;return new Rt(Ut(t>=240?t-240:t+120,i,r),Ut(t,i,r),Ut(t<120?t+240:t-120,i,r),this.opacity)},displayable:function(){return(0<=this.s&&this.s<=1||isNaN(this.s))&&0<=this.l&&this.l<=1&&0<=this.opacity&&this.opacity<=1}}));var wf=Math.PI/180,Mf=180/Math.PI,Tf=.95047,Nf=1,kf=1.08883,Sf=4/29,Ef=6/29,Af=3*Ef*Ef,Cf=Ef*Ef*Ef;Nt(It,Ft,kt(St,{brighter:function(t){return new It(this.l+18*(null==t?1:t),this.a,this.b,this.opacity)},darker:function(t){return new It(this.l-18*(null==t?1:t),this.a,this.b,this.opacity)},rgb:function(){var t=(this.l+16)/116,n=isNaN(this.a)?t:t+this.a/500,e=isNaN(this.b)?t:t-this.b/200;return t=Nf*Bt(t),n=Tf*Bt(n),e=kf*Bt(e),new Rt(Ht(3.2404542*n-1.5371385*t-.4985314*e),Ht(-.969266*n+1.8760108*t+.041556*e),Ht(.0556434*n-.2040259*t+1.0572252*e),this.opacity)}})),Nt(Vt,Xt,kt(St,{brighter:function(t){return new Vt(this.h,this.c,this.l+18*(null==t?1:t),this.opacity)},darker:function(t){return new Vt(this.h,this.c,this.l-18*(null==t?1:t),this.opacity)},rgb:function(){return Ot(this).rgb()}}));var zf=-.29227,Pf=-.90649,Rf=1.97294,Lf=Rf*Pf,qf=1.78277*Rf,Df=1.78277*zf- -.14861*Pf;Nt(Wt,$t,kt(St,{brighter:function(t){return t=null==t?1/.7:Math.pow(1/.7,t),new Wt(this.h,this.s,this.l*t,this.opacity)},darker:function(t){return t=null==t?.7:Math.pow(.7,t),new Wt(this.h,this.s,this.l*t,this.opacity)},rgb:function(){var t=isNaN(this.h)?0:(this.h+120)*wf,n=+this.l,e=isNaN(this.s)?0:this.s*n*(1-n),r=Math.cos(t),i=Math.sin(t);return new Rt(255*(n+e*(-.14861*r+1.78277*i)),255*(n+e*(zf*r+Pf*i)),255*(n+e*(Rf*r)),this.opacity)}}));var Uf,Of,Ff,If,Yf,Bf,Hf=function t(n){function e(t,n){var e=r((t=Pt(t)).r,(n=Pt(n)).r),i=r(t.g,n.g),o=r(t.b,n.b),u=en(t.opacity,n.opacity);return function(n){return t.r=e(n),t.g=i(n),t.b=o(n),t.opacity=u(n),t+""}}var r=nn(n);return e.gamma=t,e}(1),jf=rn(Gt),Xf=rn(Qt),Vf=/[-+]?(?:\d+\.?\d*|\.?\d+)(?:[eE][-+]?\d+)?/g,$f=new RegExp(Vf.source,"g"),Wf=180/Math.PI,Zf={translateX:0,translateY:0,rotate:0,skewX:0,scaleX:1,scaleY:1},Gf=pn(function(t){return"none"===t?Zf:(Uf||(Uf=document.createElement("DIV"),Of=document.documentElement,Ff=document.defaultView),Uf.style.transform=t,t=Ff.getComputedStyle(Of.appendChild(Uf),null).getPropertyValue("transform"),Of.removeChild(Uf),t=t.slice(7,-1).split(","),hn(+t[0],+t[1],+t[2],+t[3],+t[4],+t[5]))},"px, ","px)","deg)"),Qf=pn(function(t){return null==t?Zf:(If||(If=document.createElementNS("http://www.w3.org/2000/svg","g")),If.setAttribute("transform",t),(t=If.transform.baseVal.consolidate())?(t=t.matrix,hn(t.a,t.b,t.c,t.d,t.e,t.f)):Zf)},", ",")",")"),Jf=Math.SQRT2,Kf=2,tl=4,nl=1e-12,el=gn(tn),rl=gn(en),il=_n(tn),ol=_n(en),ul=yn(tn),al=yn(en),cl=0,sl=0,fl=0,ll=1e3,hl=0,pl=0,dl=0,vl="object"==typeof performance&&performance.now?performance:Date,gl="object"==typeof window&&window.requestAnimationFrame?window.requestAnimationFrame.bind(window):function(t){setTimeout(t,17)};bn.prototype=wn.prototype={constructor:bn,restart:function(t,n,e){if("function"!=typeof t)throw new TypeError("callback is not a function");e=(null==e?mn():+e)+(null==n?0:+n),this._next||Bf===this||(Bf?Bf._next=this:Yf=this,Bf=this),this._call=t,this._time=e,kn()},stop:function(){this._call&&(this._call=null,this._time=1/0,kn())}};var _l=N("start","end","interrupt"),yl=[],ml=0,xl=1,bl=2,wl=3,Ml=4,Tl=5,Nl=6,kl=at.prototype.constructor,Sl=0,El=at.prototype;qn.prototype=Dn.prototype={constructor:qn,select:function(t){var n=this._name,e=this._id;"function"!=typeof t&&(t=z(t));for(var r=this._groups,i=r.length,o=new Array(i),u=0;u<i;++u)for(var a,c,s=r[u],f=s.length,l=o[u]=new Array(f),h=0;h<f;++h)(a=s[h])&&(c=t.call(a,a.__data__,h,s))&&("__data__"in a&&(c.__data__=a.__data__),l[h]=c,En(l[h],n,e,h,l,zn(a,e)));return new qn(o,this._parents,n,e)},selectAll:function(t){var n=this._name,e=this._id;"function"!=typeof t&&(t=R(t));for(var r=this._groups,i=r.length,o=[],u=[],a=0;a<i;++a)for(var c,s=r[a],f=s.length,l=0;l<f;++l)if(c=s[l]){for(var h,p=t.call(c,c.__data__,l,s),d=zn(c,e),v=0,g=p.length;v<g;++v)(h=p[v])&&En(h,n,e,v,p,d);o.push(p),u.push(c)}return new qn(o,u,n,e)},filter:function(t){"function"!=typeof t&&(t=of(t));for(var n=this._groups,e=n.length,r=new Array(e),i=0;i<e;++i)for(var o,u=n[i],a=u.length,c=r[i]=[],s=0;s<a;++s)(o=u[s])&&t.call(o,o.__data__,s,u)&&c.push(o);return new qn(r,this._parents,this._name,this._id)},merge:function(t){if(t._id!==this._id)throw new Error;for(var n=this._groups,e=t._groups,r=n.length,i=e.length,o=Math.min(r,i),u=new Array(r),a=0;a<o;++a)for(var c,s=n[a],f=e[a],l=s.length,h=u[a]=new Array(l),p=0;p<l;++p)(c=s[p]||f[p])&&(h[p]=c);for(;a<r;++a)u[a]=n[a];return new qn(u,this._parents,this._name,this._id)},selection:function(){return new kl(this._groups,this._parents)},transition:function(){for(var t=this._name,n=this._id,e=Un(),r=this._groups,i=r.length,o=0;o<i;++o)for(var u,a=r[o],c=a.length,s=0;s<c;++s)if(u=a[s]){var f=zn(u,n);En(u,t,e,s,a,{time:f.time+f.delay+f.duration,delay:0,duration:f.duration,ease:f.ease})}return new qn(r,this._parents,t,e)},call:El.call,nodes:El.nodes,node:El.node,size:El.size,empty:El.empty,each:El.each,on:function(t,n){var e=this._id;return arguments.length<2?zn(this.node(),e).on.on(t):this.each(function(t,n,e){var r,i,o=function(t){return(t+"").trim().split(/^|\s+/).every(function(t){var n=t.indexOf(".");return n>=0&&(t=t.slice(0,n)),!t||"start"===t})}(n)?An:Cn;return function(){var u=o(this,t),a=u.on;a!==r&&(i=(r=a).copy()).on(n,e),u.on=i}}(e,t,n))},attr:function(t,n){var e=E(t),r="transform"===e?Qf:Ln;return this.attrTween(t,"function"==typeof n?(e.local?function(t,n,e){var r,i,o;return function(){var u,a=e(this);if(null!=a)return(u=this.getAttributeNS(t.space,t.local))===a?null:u===r&&a===i?o:o=n(r=u,i=a);this.removeAttributeNS(t.space,t.local)}}:function(t,n,e){var r,i,o;return function(){var u,a=e(this);if(null!=a)return(u=this.getAttribute(t))===a?null:u===r&&a===i?o:o=n(r=u,i=a);this.removeAttribute(t)}})(e,r,Rn(this,"attr."+t,n)):null==n?(e.local?function(t){return function(){this.removeAttributeNS(t.space,t.local)}}:function(t){return function(){this.removeAttribute(t)}})(e):(e.local?function(t,n,e){var r,i;return function(){var o=this.getAttributeNS(t.space,t.local);return o===e?null:o===r?i:i=n(r=o,e)}}:function(t,n,e){var r,i;return function(){var o=this.getAttribute(t);return o===e?null:o===r?i:i=n(r=o,e)}})(e,r,n+""))},attrTween:function(t,n){var e="attr."+t;if(arguments.length<2)return(e=this.tween(e))&&e._value;if(null==n)return this.tween(e,null);if("function"!=typeof n)throw new Error;var r=E(t);return this.tween(e,(r.local?function(t,n){function e(){var e=this,r=n.apply(e,arguments);return r&&function(n){e.setAttributeNS(t.space,t.local,r(n))}}return e._value=n,e}:function(t,n){function e(){var e=this,r=n.apply(e,arguments);return r&&function(n){e.setAttribute(t,r(n))}}return e._value=n,e})(r,n))},style:function(t,n,e){var r="transform"==(t+="")?Gf:Ln;return null==n?this.styleTween(t,function(t,n){var e,r,i;return function(){var o=I(this,t),u=(this.style.removeProperty(t),I(this,t));return o===u?null:o===e&&u===r?i:i=n(e=o,r=u)}}(t,r)).on("end.style."+t,function(t){return function(){this.style.removeProperty(t)}}(t)):this.styleTween(t,"function"==typeof n?function(t,n,e){var r,i,o;return function(){var u=I(this,t),a=e(this);return null==a&&(this.style.removeProperty(t),a=I(this,t)),u===a?null:u===r&&a===i?o:o=n(r=u,i=a)}}(t,r,Rn(this,"style."+t,n)):function(t,n,e){var r,i;return function(){var o=I(this,t);return o===e?null:o===r?i:i=n(r=o,e)}}(t,r,n+""),e)},styleTween:function(t,n,e){var r="style."+(t+="");if(arguments.length<2)return(r=this.tween(r))&&r._value;if(null==n)return this.tween(r,null);if("function"!=typeof n)throw new Error;return this.tween(r,function(t,n,e){function r(){var r=this,i=n.apply(r,arguments);return i&&function(n){r.style.setProperty(t,i(n),e)}}return r._value=n,r}(t,n,null==e?"":e))},text:function(t){return this.tween("text","function"==typeof t?function(t){return function(){var n=t(this);this.textContent=null==n?"":n}}(Rn(this,"text",t)):function(t){return function(){this.textContent=t}}(null==t?"":t+""))},remove:function(){return this.on("end.remove",function(t){return function(){var n=this.parentNode;for(var e in this.__transition)if(+e!==t)return;n&&n.removeChild(this)}}(this._id))},tween:function(t,n){var e=this._id;if(t+="",arguments.length<2){for(var r,i=zn(this.node(),e).tween,o=0,u=i.length;o<u;++o)if((r=i[o]).name===t)return r.value;return null}return this.each((null==n?function(t,n){var e,r;return function(){var i=Cn(this,t),o=i.tween;if(o!==e)for(var u=0,a=(r=e=o).length;u<a;++u)if(r[u].name===n){(r=r.slice()).splice(u,1);break}i.tween=r}}:function(t,n,e){var r,i;if("function"!=typeof e)throw new Error;return function(){var o=Cn(this,t),u=o.tween;if(u!==r){i=(r=u).slice();for(var a={name:n,value:e},c=0,s=i.length;c<s;++c)if(i[c].name===n){i[c]=a;break}c===s&&i.push(a)}o.tween=i}})(e,t,n))},delay:function(t){var n=this._id;return arguments.length?this.each(("function"==typeof t?function(t,n){return function(){An(this,t).delay=+n.apply(this,arguments)}}:function(t,n){return n=+n,function(){An(this,t).delay=n}})(n,t)):zn(this.node(),n).delay},duration:function(t){var n=this._id;return arguments.length?this.each(("function"==typeof t?function(t,n){return function(){Cn(this,t).duration=+n.apply(this,arguments)}}:function(t,n){return n=+n,function(){Cn(this,t).duration=n}})(n,t)):zn(this.node(),n).duration},ease:function(t){var n=this._id;return arguments.length?this.each(function(t,n){if("function"!=typeof n)throw new Error;return function(){Cn(this,t).ease=n}}(n,t)):zn(this.node(),n).ease}};var Al=function t(n){function e(t){return Math.pow(t,n)}return n=+n,e.exponent=t,e}(3),Cl=function t(n){function e(t){return 1-Math.pow(1-t,n)}return n=+n,e.exponent=t,e}(3),zl=function t(n){function e(t){return((t*=2)<=1?Math.pow(t,n):2-Math.pow(2-t,n))/2}return n=+n,e.exponent=t,e}(3),Pl=Math.PI,Rl=Pl/2,Ll=4/11,ql=6/11,Dl=8/11,Ul=.75,Ol=9/11,Fl=10/11,Il=.9375,Yl=21/22,Bl=63/64,Hl=1/Ll/Ll,jl=function t(n){function e(t){return t*t*((n+1)*t-n)}return n=+n,e.overshoot=t,e}(1.70158),Xl=function t(n){function e(t){return--t*t*((n+1)*t+n)+1}return n=+n,e.overshoot=t,e}(1.70158),Vl=function t(n){function e(t){return((t*=2)<1?t*t*((n+1)*t-n):(t-=2)*t*((n+1)*t+n)+2)/2}return n=+n,e.overshoot=t,e}(1.70158),$l=2*Math.PI,Wl=function t(n,e){function r(t){return n*Math.pow(2,10*--t)*Math.sin((i-t)/e)}var i=Math.asin(1/(n=Math.max(1,n)))*(e/=$l);return r.amplitude=function(n){return t(n,e*$l)},r.period=function(e){return t(n,e)},r}(1,.3),Zl=function t(n,e){function r(t){return 1-n*Math.pow(2,-10*(t=+t))*Math.sin((t+i)/e)}var i=Math.asin(1/(n=Math.max(1,n)))*(e/=$l);return r.amplitude=function(n){return t(n,e*$l)},r.period=function(e){return t(n,e)},r}(1,.3),Gl=function t(n,e){function r(t){return((t=2*t-1)<0?n*Math.pow(2,10*t)*Math.sin((i-t)/e):2-n*Math.pow(2,-10*t)*Math.sin((i+t)/e))/2}var i=Math.asin(1/(n=Math.max(1,n)))*(e/=$l);return r.amplitude=function(n){return t(n,e*$l)},r.period=function(e){return t(n,e)},r}(1,.3),Ql={time:null,delay:0,duration:250,ease:Fn};at.prototype.interrupt=function(t){return this.each(function(){Pn(this,t)})},at.prototype.transition=function(t){var n,e;t instanceof qn?(n=t._id,t=t._name):(n=Un(),(e=Ql).time=mn(),t=null==t?null:t+"");for(var r=this._groups,i=r.length,o=0;o<i;++o)for(var u,a=r[o],c=a.length,s=0;s<c;++s)(u=a[s])&&En(u,t,n,s,a,e||jn(u,n));return new qn(r,this._parents,t,n)};var Jl=[null],Kl={name:"drag"},th={name:"space"},nh={name:"handle"},eh={name:"center"},rh={name:"x",handles:["e","w"].map(Wn),input:function(t,n){return t&&[[t[0],n[0][1]],[t[1],n[1][1]]]},output:function(t){return t&&[t[0][0],t[1][0]]}},ih={name:"y",handles:["n","s"].map(Wn),input:function(t,n){return t&&[[n[0][0],t[0]],[n[1][0],t[1]]]},output:function(t){return t&&[t[0][1],t[1][1]]}},oh={name:"xy",handles:["n","e","s","w","nw","ne","se","sw"].map(Wn),input:function(t){return t},output:function(t){return t}},uh={overlay:"crosshair",selection:"move",n:"ns-resize",e:"ew-resize",s:"ns-resize",w:"ew-resize",nw:"nwse-resize",ne:"nesw-resize",se:"nwse-resize",sw:"nesw-resize"},ah={e:"w",w:"e",nw:"ne",ne:"nw",se:"sw",sw:"se"},ch={n:"s",s:"n",nw:"sw",ne:"se",se:"ne",sw:"nw"},sh={overlay:1,selection:1,n:null,e:1,s:null,w:-1,nw:-1,ne:1,se:1,sw:-1},fh={overlay:1,selection:1,n:-1,e:null,s:1,w:null,nw:-1,ne:-1,se:1,sw:1},lh=Math.cos,hh=Math.sin,ph=Math.PI,dh=ph/2,vh=2*ph,gh=Math.max,_h=Array.prototype.slice,yh=Math.PI,mh=2*yh,xh=mh-1e-6;ne.prototype=ee.prototype={constructor:ne,moveTo:function(t,n){this._+="M"+(this._x0=this._x1=+t)+","+(this._y0=this._y1=+n)},closePath:function(){null!==this._x1&&(this._x1=this._x0,this._y1=this._y0,this._+="Z")},lineTo:function(t,n){this._+="L"+(this._x1=+t)+","+(this._y1=+n)},quadraticCurveTo:function(t,n,e,r){this._+="Q"+ +t+","+ +n+","+(this._x1=+e)+","+(this._y1=+r)},bezierCurveTo:function(t,n,e,r,i,o){this._+="C"+ +t+","+ +n+","+ +e+","+ +r+","+(this._x1=+i)+","+(this._y1=+o)},arcTo:function(t,n,e,r,i){t=+t,n=+n,e=+e,r=+r,i=+i;var o=this._x1,u=this._y1,a=e-t,c=r-n,s=o-t,f=u-n,l=s*s+f*f;if(i<0)throw new Error("negative radius: "+i);if(null===this._x1)this._+="M"+(this._x1=t)+","+(this._y1=n);else if(l>1e-6)if(Math.abs(f*a-c*s)>1e-6&&i){var h=e-o,p=r-u,d=a*a+c*c,v=h*h+p*p,g=Math.sqrt(d),_=Math.sqrt(l),y=i*Math.tan((yh-Math.acos((d+l-v)/(2*g*_)))/2),m=y/_,x=y/g;Math.abs(m-1)>1e-6&&(this._+="L"+(t+m*s)+","+(n+m*f)),this._+="A"+i+","+i+",0,0,"+ +(f*h>s*p)+","+(this._x1=t+x*a)+","+(this._y1=n+x*c)}else this._+="L"+(this._x1=t)+","+(this._y1=n);else;},arc:function(t,n,e,r,i,o){t=+t,n=+n;var u=(e=+e)*Math.cos(r),a=e*Math.sin(r),c=t+u,s=n+a,f=1^o,l=o?r-i:i-r;if(e<0)throw new Error("negative radius: "+e);null===this._x1?this._+="M"+c+","+s:(Math.abs(this._x1-c)>1e-6||Math.abs(this._y1-s)>1e-6)&&(this._+="L"+c+","+s),e&&(l<0&&(l=l%mh+mh),l>xh?this._+="A"+e+","+e+",0,1,"+f+","+(t-u)+","+(n-a)+"A"+e+","+e+",0,1,"+f+","+(this._x1=c)+","+(this._y1=s):l>1e-6&&(this._+="A"+e+","+e+",0,"+ +(l>=yh)+","+f+","+(this._x1=t+e*Math.cos(i))+","+(this._y1=n+e*Math.sin(i))))},rect:function(t,n,e,r){this._+="M"+(this._x0=this._x1=+t)+","+(this._y0=this._y1=+n)+"h"+ +e+"v"+ +r+"h"+-e+"Z"},toString:function(){return this._}};ce.prototype=se.prototype={constructor:ce,has:function(t){return"$"+t in this},get:function(t){return this["$"+t]},set:function(t,n){return this["$"+t]=n,this},remove:function(t){var n="$"+t;return n in this&&delete this[n]},clear:function(){for(var t in this)"$"===t[0]&&delete this[t]},keys:function(){var t=[];for(var n in this)"$"===n[0]&&t.push(n.slice(1));return t},values:function(){var t=[];for(var n in this)"$"===n[0]&&t.push(this[n]);return t},entries:function(){var t=[];for(var n in this)"$"===n[0]&&t.push({key:n.slice(1),value:this[n]});return t},size:function(){var t=0;for(var n in this)"$"===n[0]&&++t;return t},empty:function(){for(var t in this)if("$"===t[0])return!1;return!0},each:function(t){for(var n in this)"$"===n[0]&&t(this[n],n.slice(1),this)}};var bh=se.prototype;de.prototype=ve.prototype={constructor:de,has:bh.has,add:function(t){return t+="",this["$"+t]=t,this},remove:bh.remove,clear:bh.clear,values:bh.keys,size:bh.size,empty:bh.empty,each:bh.each};var wh={},Mh={},Th=34,Nh=10,kh=13,Sh=_e(","),Eh=Sh.parse,Ah=Sh.parseRows,Ch=Sh.format,zh=Sh.formatRows,Ph=_e("\t"),Rh=Ph.parse,Lh=Ph.parseRows,qh=Ph.format,Dh=Ph.formatRows,Uh=Te.prototype=Ne.prototype;Uh.copy=function(){var t,n,e=new Ne(this._x,this._y,this._x0,this._y0,this._x1,this._y1),r=this._root;if(!r)return e;if(!r.length)return e._root=ke(r),e;for(t=[{source:r,target:e._root=new Array(4)}];r=t.pop();)for(var i=0;i<4;++i)(n=r.source[i])&&(n.length?t.push({source:n,target:r.target[i]=new Array(4)}):r.target[i]=ke(n));return e},Uh.add=function(t){var n=+this._x.call(null,t),e=+this._y.call(null,t);return xe(this.cover(n,e),n,e,t)},Uh.addAll=function(t){var n,e,r,i,o=t.length,u=new Array(o),a=new Array(o),c=1/0,s=1/0,f=-1/0,l=-1/0;for(e=0;e<o;++e)isNaN(r=+this._x.call(null,n=t[e]))||isNaN(i=+this._y.call(null,n))||(u[e]=r,a[e]=i,r<c&&(c=r),r>f&&(f=r),i<s&&(s=i),i>l&&(l=i));for(f<c&&(c=this._x0,f=this._x1),l<s&&(s=this._y0,l=this._y1),this.cover(c,s).cover(f,l),e=0;e<o;++e)xe(this,u[e],a[e],t[e]);return this},Uh.cover=function(t,n){if(isNaN(t=+t)||isNaN(n=+n))return this;var e=this._x0,r=this._y0,i=this._x1,o=this._y1;if(isNaN(e))i=(e=Math.floor(t))+1,o=(r=Math.floor(n))+1;else{if(!(e>t||t>i||r>n||n>o))return this;var u,a,c=i-e,s=this._root;switch(a=(n<(r+o)/2)<<1|t<(e+i)/2){case 0:do{u=new Array(4),u[a]=s,s=u}while(c*=2,i=e+c,o=r+c,t>i||n>o);break;case 1:do{u=new Array(4),u[a]=s,s=u}while(c*=2,e=i-c,o=r+c,e>t||n>o);break;case 2:do{u=new Array(4),u[a]=s,s=u}while(c*=2,i=e+c,r=o-c,t>i||r>n);break;case 3:do{u=new Array(4),u[a]=s,s=u}while(c*=2,e=i-c,r=o-c,e>t||r>n)}this._root&&this._root.length&&(this._root=s)}return this._x0=e,this._y0=r,this._x1=i,this._y1=o,this},Uh.data=function(){var t=[];return this.visit(function(n){if(!n.length)do{t.push(n.data)}while(n=n.next)}),t},Uh.extent=function(t){return arguments.length?this.cover(+t[0][0],+t[0][1]).cover(+t[1][0],+t[1][1]):isNaN(this._x0)?void 0:[[this._x0,this._y0],[this._x1,this._y1]]},Uh.find=function(t,n,e){var r,i,o,u,a,c,s,f=this._x0,l=this._y0,h=this._x1,p=this._y1,d=[],v=this._root;for(v&&d.push(new be(v,f,l,h,p)),null==e?e=1/0:(f=t-e,l=n-e,h=t+e,p=n+e,e*=e);c=d.pop();)if(!(!(v=c.node)||(i=c.x0)>h||(o=c.y0)>p||(u=c.x1)<f||(a=c.y1)<l))if(v.length){var g=(i+u)/2,_=(o+a)/2;d.push(new be(v[3],g,_,u,a),new be(v[2],i,_,g,a),new be(v[1],g,o,u,_),new be(v[0],i,o,g,_)),(s=(n>=_)<<1|t>=g)&&(c=d[d.length-1],d[d.length-1]=d[d.length-1-s],d[d.length-1-s]=c)}else{var y=t-+this._x.call(null,v.data),m=n-+this._y.call(null,v.data),x=y*y+m*m;if(x<e){var b=Math.sqrt(e=x);f=t-b,l=n-b,h=t+b,p=n+b,r=v.data}}return r},Uh.remove=function(t){if(isNaN(o=+this._x.call(null,t))||isNaN(u=+this._y.call(null,t)))return this;var n,e,r,i,o,u,a,c,s,f,l,h,p=this._root,d=this._x0,v=this._y0,g=this._x1,_=this._y1;if(!p)return this;if(p.length)for(;;){if((s=o>=(a=(d+g)/2))?d=a:g=a,(f=u>=(c=(v+_)/2))?v=c:_=c,n=p,!(p=p[l=f<<1|s]))return this;if(!p.length)break;(n[l+1&3]||n[l+2&3]||n[l+3&3])&&(e=n,h=l)}for(;p.data!==t;)if(r=p,!(p=p.next))return this;return(i=p.next)&&delete p.next,r?(i?r.next=i:delete r.next,this):n?(i?n[l]=i:delete n[l],(p=n[0]||n[1]||n[2]||n[3])&&p===(n[3]||n[2]||n[1]||n[0])&&!p.length&&(e?e[h]=p:this._root=p),this):(this._root=i,this)},Uh.removeAll=function(t){for(var n=0,e=t.length;n<e;++n)this.remove(t[n]);return this},Uh.root=function(){return this._root},Uh.size=function(){var t=0;return this.visit(function(n){if(!n.length)do{++t}while(n=n.next)}),t},Uh.visit=function(t){var n,e,r,i,o,u,a=[],c=this._root;for(c&&a.push(new be(c,this._x0,this._y0,this._x1,this._y1));n=a.pop();)if(!t(c=n.node,r=n.x0,i=n.y0,o=n.x1,u=n.y1)&&c.length){var s=(r+o)/2,f=(i+u)/2;(e=c[3])&&a.push(new be(e,s,f,o,u)),(e=c[2])&&a.push(new be(e,r,f,s,u)),(e=c[1])&&a.push(new be(e,s,i,o,f)),(e=c[0])&&a.push(new be(e,r,i,s,f))}return this},Uh.visitAfter=function(t){var n,e=[],r=[];for(this._root&&e.push(new be(this._root,this._x0,this._y0,this._x1,this._y1));n=e.pop();){var i=n.node;if(i.length){var o,u=n.x0,a=n.y0,c=n.x1,s=n.y1,f=(u+c)/2,l=(a+s)/2;(o=i[0])&&e.push(new be(o,u,a,f,l)),(o=i[1])&&e.push(new be(o,f,a,c,l)),(o=i[2])&&e.push(new be(o,u,l,f,s)),(o=i[3])&&e.push(new be(o,f,l,c,s))}r.push(n)}for(;n=r.pop();)t(n.node,n.x0,n.y0,n.x1,n.y1);return this},Uh.x=function(t){return arguments.length?(this._x=t,this):this._x},Uh.y=function(t){return arguments.length?(this._y=t,this):this._y};var Oh,Fh=10,Ih=Math.PI*(3-Math.sqrt(5)),Yh={"":function(t,n){t:for(var e,r=(t=t.toPrecision(n)).length,i=1,o=-1;i<r;++i)switch(t[i]){case".":o=e=i;break;case"0":0===o&&(o=i),e=i;break;case"e":break t;default:o>0&&(o=0)}return o>0?t.slice(0,o)+t.slice(e+1):t},"%":function(t,n){return(100*t).toFixed(n)},b:function(t){return Math.round(t).toString(2)},c:function(t){return t+""},d:function(t){return Math.round(t).toString(10)},e:function(t,n){return t.toExponential(n)},f:function(t,n){return t.toFixed(n)},g:function(t,n){return t.toPrecision(n)},o:function(t){return Math.round(t).toString(8)},p:function(t,n){return qe(100*t,n)},r:qe,s:function(t,n){var e=Re(t,n);if(!e)return t+"";var r=e[0],i=e[1],o=i-(Oh=3*Math.max(-8,Math.min(8,Math.floor(i/3))))+1,u=r.length;return o===u?r:o>u?r+new Array(o-u+1).join("0"):o>0?r.slice(0,o)+"."+r.slice(o):"0."+new Array(1-o).join("0")+Re(t,Math.max(0,n+o-1))[0]},X:function(t){return Math.round(t).toString(16).toUpperCase()},x:function(t){return Math.round(t).toString(16)}},Bh=/^(?:(.)?([<>=^]))?([+\-\( ])?([$#])?(0)?(\d+)?(,)?(\.\d+)?([a-z%])?$/i;De.prototype=Ue.prototype,Ue.prototype.toString=function(){return this.fill+this.align+this.sign+this.symbol+(this.zero?"0":"")+(null==this.width?"":Math.max(1,0|this.width))+(this.comma?",":"")+(null==this.precision?"":"."+Math.max(0,0|this.precision))+this.type};var Hh,jh=["y","z","a","f","p","n","µ","m","","k","M","G","T","P","E","Z","Y"];Ie({decimal:".",thousands:",",grouping:[3],currency:["$",""]}),Xe.prototype={constructor:Xe,reset:function(){this.s=this.t=0},add:function(t){Ve(wp,t,this.t),Ve(this,wp.s,this.s),this.s?this.t+=wp.t:this.s=wp.t},valueOf:function(){return this.s}};var Xh,Vh,$h,Wh,Zh,Gh,Qh,Jh,Kh,tp,np,ep,rp,ip,op,up,ap,cp,sp,fp,lp,hp,pp,dp,vp,gp,_p,yp,mp,xp,bp,wp=new Xe,Mp=1e-6,Tp=1e-12,Np=Math.PI,kp=Np/2,Sp=Np/4,Ep=2*Np,Ap=180/Np,Cp=Np/180,zp=Math.abs,Pp=Math.atan,Rp=Math.atan2,Lp=Math.cos,qp=Math.ceil,Dp=Math.exp,Up=Math.log,Op=Math.pow,Fp=Math.sin,Ip=Math.sign||function(t){return t>0?1:t<0?-1:0},Yp=Math.sqrt,Bp=Math.tan,Hp={Feature:function(t,n){Qe(t.geometry,n)},FeatureCollection:function(t,n){for(var e=t.features,r=-1,i=e.length;++r<i;)Qe(e[r].geometry,n)}},jp={Sphere:function(t,n){n.sphere()},Point:function(t,n){t=t.coordinates,n.point(t[0],t[1],t[2])},MultiPoint:function(t,n){for(var e=t.coordinates,r=-1,i=e.length;++r<i;)t=e[r],n.point(t[0],t[1],t[2])},LineString:function(t,n){Je(t.coordinates,n,0)},MultiLineString:function(t,n){for(var e=t.coordinates,r=-1,i=e.length;++r<i;)Je(e[r],n,0)},Polygon:function(t,n){Ke(t.coordinates,n)},MultiPolygon:function(t,n){for(var e=t.coordinates,r=-1,i=e.length;++r<i;)Ke(e[r],n)},GeometryCollection:function(t,n){for(var e=t.geometries,r=-1,i=e.length;++r<i;)Qe(e[r],n)}},Xp=je(),Vp=je(),$p={point:Ge,lineStart:Ge,lineEnd:Ge,polygonStart:function(){Xp.reset(),$p.lineStart=nr,$p.lineEnd=er},polygonEnd:function(){var t=+Xp;Vp.add(t<0?Ep+t:t),this.lineStart=this.lineEnd=this.point=Ge},sphere:function(){Vp.add(Ep)}},Wp=je(),Zp={point:hr,lineStart:dr,lineEnd:vr,polygonStart:function(){Zp.point=gr,Zp.lineStart=_r,Zp.lineEnd=yr,Wp.reset(),$p.polygonStart()},polygonEnd:function(){$p.polygonEnd(),Zp.point=hr,Zp.lineStart=dr,Zp.lineEnd=vr,Xp<0?(Gh=-(Jh=180),Qh=-(Kh=90)):Wp>Mp?Kh=90:Wp<-Mp&&(Qh=-90),op[0]=Gh,op[1]=Jh}},Gp={sphere:Ge,point:wr,lineStart:Tr,lineEnd:Sr,polygonStart:function(){Gp.lineStart=Er,Gp.lineEnd=Ar},polygonEnd:function(){Gp.lineStart=Tr,Gp.lineEnd=Sr}};Lr.invert=Lr;var Qp,Jp,Kp,td,nd,ed,rd,id,od,ud,ad,cd=je(),sd=Wr(function(){return!0},function(t){var n,e=NaN,r=NaN,i=NaN;return{lineStart:function(){t.lineStart(),n=1},point:function(o,u){var a=o>0?Np:-Np,c=zp(o-e);zp(c-Np)<Mp?(t.point(e,r=(r+u)/2>0?kp:-kp),t.point(i,r),t.lineEnd(),t.lineStart(),t.point(a,r),t.point(o,r),n=0):i!==a&&c>=Np&&(zp(e-i)<Mp&&(e-=i*Mp),zp(o-a)<Mp&&(o-=a*Mp),r=function(t,n,e,r){var i,o,u=Fp(t-e);return zp(u)>Mp?Pp((Fp(n)*(o=Lp(r))*Fp(e)-Fp(r)*(i=Lp(n))*Fp(t))/(i*o*u)):(n+r)/2}(e,r,o,u),t.point(i,r),t.lineEnd(),t.lineStart(),t.point(a,r),n=0),t.point(e=o,r=u),i=a},lineEnd:function(){t.lineEnd(),e=r=NaN},clean:function(){return 2-n}}},function(t,n,e,r){var i;if(null==t)i=e*kp,r.point(-Np,i),r.point(0,i),r.point(Np,i),r.point(Np,0),r.point(Np,-i),r.point(0,-i),r.point(-Np,-i),r.point(-Np,0),r.point(-Np,i);else if(zp(t[0]-n[0])>Mp){var o=t[0]<n[0]?Np:-Np;i=e*o/2,r.point(-o,i),r.point(0,i),r.point(o,i)}else r.point(n[0],n[1])},[-Np,-kp]),fd=1e9,ld=-fd,hd=je(),pd={sphere:Ge,point:Ge,lineStart:function(){pd.point=ti,pd.lineEnd=Kr},lineEnd:Ge,polygonStart:Ge,polygonEnd:Ge},dd=[null,null],vd={type:"LineString",coordinates:dd},gd={Feature:function(t,n){return ii(t.geometry,n)},FeatureCollection:function(t,n){for(var e=t.features,r=-1,i=e.length;++r<i;)if(ii(e[r].geometry,n))return!0;return!1}},_d={Sphere:function(){return!0},Point:function(t,n){return oi(t.coordinates,n)},MultiPoint:function(t,n){for(var e=t.coordinates,r=-1,i=e.length;++r<i;)if(oi(e[r],n))return!0;return!1},LineString:function(t,n){return ui(t.coordinates,n)},MultiLineString:function(t,n){for(var e=t.coordinates,r=-1,i=e.length;++r<i;)if(ui(e[r],n))return!0;return!1},Polygon:function(t,n){return ai(t.coordinates,n)},MultiPolygon:function(t,n){for(var e=t.coordinates,r=-1,i=e.length;++r<i;)if(ai(e[r],n))return!0;return!1},GeometryCollection:function(t,n){for(var e=t.geometries,r=-1,i=e.length;++r<i;)if(ii(e[r],n))return!0;return!1}},yd=je(),md=je(),xd={point:Ge,lineStart:Ge,lineEnd:Ge,polygonStart:function(){xd.lineStart=di,xd.lineEnd=_i},polygonEnd:function(){xd.lineStart=xd.lineEnd=xd.point=Ge,yd.add(zp(md)),md.reset()},result:function(){var t=yd/2;return yd.reset(),t}},bd=1/0,wd=bd,Md=-bd,Td=Md,Nd={point:function(t,n){t<bd&&(bd=t),t>Md&&(Md=t),n<wd&&(wd=n),n>Td&&(Td=n)},lineStart:Ge,lineEnd:Ge,polygonStart:Ge,polygonEnd:Ge,result:function(){var t=[[bd,wd],[Md,Td]];return Md=Td=-(wd=bd=1/0),t}},kd=0,Sd=0,Ed=0,Ad=0,Cd=0,zd=0,Pd=0,Rd=0,Ld=0,qd={point:yi,lineStart:mi,lineEnd:wi,polygonStart:function(){qd.lineStart=Mi,qd.lineEnd=Ti},polygonEnd:function(){qd.point=yi,qd.lineStart=mi,qd.lineEnd=wi},result:function(){var t=Ld?[Pd/Ld,Rd/Ld]:zd?[Ad/zd,Cd/zd]:Ed?[kd/Ed,Sd/Ed]:[NaN,NaN];return kd=Sd=Ed=Ad=Cd=zd=Pd=Rd=Ld=0,t}};Si.prototype={_radius:4.5,pointRadius:function(t){return this._radius=t,this},polygonStart:function(){this._line=0},polygonEnd:function(){this._line=NaN},lineStart:function(){this._point=0},lineEnd:function(){0===this._line&&this._context.closePath(),this._point=NaN},point:function(t,n){switch(this._point){case 0:this._context.moveTo(t,n),this._point=1;break;case 1:this._context.lineTo(t,n);break;default:this._context.moveTo(t+this._radius,n),this._context.arc(t,n,this._radius,0,Ep)}},result:Ge};var Dd,Ud,Od,Fd,Id,Yd=je(),Bd={point:Ge,lineStart:function(){Bd.point=Ei},lineEnd:function(){Dd&&Ai(Ud,Od),Bd.point=Ge},polygonStart:function(){Dd=!0},polygonEnd:function(){Dd=null},result:function(){var t=+Yd;return Yd.reset(),t}};Ci.prototype={_radius:4.5,_circle:zi(4.5),pointRadius:function(t){return(t=+t)!==this._radius&&(this._radius=t,this._circle=null),this},polygonStart:function(){this._line=0},polygonEnd:function(){this._line=NaN},lineStart:function(){this._point=0},lineEnd:function(){0===this._line&&this._string.push("Z"),this._point=NaN},point:function(t,n){switch(this._point){case 0:this._string.push("M",t,",",n),this._point=1;break;case 1:this._string.push("L",t,",",n);break;default:null==this._circle&&(this._circle=zi(this._radius)),this._string.push("M",t,",",n,this._circle)}},result:function(){if(this._string.length){var t=this._string.join("");return this._string=[],t}return null}},Ri.prototype={constructor:Ri,point:function(t,n){this.stream.point(t,n)},sphere:function(){this.stream.sphere()},lineStart:function(){this.stream.lineStart()},lineEnd:function(){this.stream.lineEnd()},polygonStart:function(){this.stream.polygonStart()},polygonEnd:function(){this.stream.polygonEnd()}};var Hd=16,jd=Lp(30*Cp),Xd=Pi({point:function(t,n){this.stream.point(t*Cp,n*Cp)}}),Vd=Vi(function(t){return Yp(2/(1+t))});Vd.invert=$i(function(t){return 2*We(t/2)});var $d=Vi(function(t){return(t=$e(t))&&t/Fp(t)});$d.invert=$i(function(t){return t}),Wi.invert=function(t,n){return[t,2*Pp(Dp(n))-kp]},Ji.invert=Ji,to.invert=$i(Pp),eo.invert=function(t,n){var e,r=n,i=25;do{var o=r*r,u=o*o;r-=e=(r*(1.007226+o*(.015085+u*(.028874*o-.044475-.005916*u)))-n)/(1.007226+o*(.045255+u*(.259866*o-.311325-.005916*11*u)))}while(zp(e)>Mp&&--i>0);return[t/(.8707+(o=r*r)*(o*(o*o*o*(.003971-.001529*o)-.013791)-.131979)),r]},ro.invert=$i(We),io.invert=$i(function(t){return 2*Pp(t)}),oo.invert=function(t,n){return[-n,2*Pp(Dp(t))-kp]},vo.prototype=fo.prototype={constructor:vo,count:function(){return this.eachAfter(so)},each:function(t){var n,e,r,i,o=this,u=[o];do{for(n=u.reverse(),u=[];o=n.pop();)if(t(o),e=o.children)for(r=0,i=e.length;r<i;++r)u.push(e[r])}while(u.length);return this},eachAfter:function(t){for(var n,e,r,i=this,o=[i],u=[];i=o.pop();)if(u.push(i),n=i.children)for(e=0,r=n.length;e<r;++e)o.push(n[e]);for(;i=u.pop();)t(i);return this},eachBefore:function(t){for(var n,e,r=this,i=[r];r=i.pop();)if(t(r),n=r.children)for(e=n.length-1;e>=0;--e)i.push(n[e]);return this},sum:function(t){return this.eachAfter(function(n){for(var e=+t(n.data)||0,r=n.children,i=r&&r.length;--i>=0;)e+=r[i].value;n.value=e})},sort:function(t){return this.eachBefore(function(n){n.children&&n.children.sort(t)})},path:function(t){for(var n=this,e=function(t,n){if(t===n)return t;var e=t.ancestors(),r=n.ancestors(),i=null;for(t=e.pop(),n=r.pop();t===n;)i=t,t=e.pop(),n=r.pop();return i}(n,t),r=[n];n!==e;)n=n.parent,r.push(n);for(var i=r.length;t!==e;)r.splice(i,0,t),t=t.parent;return r},ancestors:function(){for(var t=this,n=[t];t=t.parent;)n.push(t);return n},descendants:function(){var t=[];return this.each(function(n){t.push(n)}),t},leaves:function(){var t=[];return this.eachBefore(function(n){n.children||t.push(n)}),t},links:function(){var t=this,n=[];return t.each(function(e){e!==t&&n.push({source:e.parent,target:e})}),n},copy:function(){return fo(this).eachBefore(ho)}};var Wd=Array.prototype.slice,Zd="$",Gd={depth:-1},Qd={};Ho.prototype=Object.create(vo.prototype);var Jd=(1+Math.sqrt(5))/2,Kd=function t(n){function e(t,e,r,i,o){Xo(n,t,e,r,i,o)}return e.ratio=function(n){return t((n=+n)>1?n:1)},e}(Jd),tv=function t(n){function e(t,e,r,i,o){if((u=t._squarify)&&u.ratio===n)for(var u,a,c,s,f,l=-1,h=u.length,p=t.value;++l<h;){for(c=(a=u[l]).children,s=a.value=0,f=c.length;s<f;++s)a.value+=c[s].value;a.dice?qo(a,e,r,i,r+=(o-r)*a.value/p):jo(a,e,r,e+=(i-e)*a.value/p,o),p-=a.value}else t._squarify=u=Xo(n,t,e,r,i,o),u.ratio=n}return e.ratio=function(n){return t((n=+n)>1?n:1)},e}(Jd),nv=[].slice,ev={};Zo.prototype=Ko.prototype={constructor:Zo,defer:function(t){if("function"!=typeof t)throw new Error("invalid callback");if(this._call)throw new Error("defer after await");if(null!=this._error)return this;var n=nv.call(arguments,1);return n.push(t),++this._waiting,this._tasks.push(n),Go(this),this},abort:function(){return null==this._error&&Qo(this,new Error("abort")),this},await:function(t){if("function"!=typeof t)throw new Error("invalid callback");if(this._call)throw new Error("multiple await");return this._call=function(n,e){t.apply(null,[n].concat(e))},Jo(this),this},awaitAll:function(t){if("function"!=typeof t)throw new Error("invalid callback");if(this._call)throw new Error("multiple await");return this._call=t,Jo(this),this}};var rv=function t(n){function e(t,e){return t=null==t?0:+t,e=null==e?1:+e,1===arguments.length?(e=t,t=0):e-=t,function(){return n()*e+t}}return e.source=t,e}(tu),iv=function t(n){function e(t,e){var r,i;return t=null==t?0:+t,e=null==e?1:+e,function(){var o;if(null!=r)o=r,r=null;else do{r=2*n()-1,o=2*n()-1,i=r*r+o*o}while(!i||i>1);return t+e*o*Math.sqrt(-2*Math.log(i)/i)}}return e.source=t,e}(tu),ov=function t(n){function e(){var t=iv.source(n).apply(this,arguments);return function(){return Math.exp(t())}}return e.source=t,e}(tu),uv=function t(n){function e(t){return function(){for(var e=0,r=0;r<t;++r)e+=n();return e}}return e.source=t,e}(tu),av=function t(n){function e(t){var e=uv.source(n)(t);return function(){return e()/t}}return e.source=t,e}(tu),cv=function t(n){function e(t){return function(){return-Math.log(1-n())/t}}return e.source=t,e}(tu),sv=eu("text/html",function(t){return document.createRange().createContextualFragment(t.responseText)}),fv=eu("application/json",function(t){return JSON.parse(t.responseText)}),lv=eu("text/plain",function(t){return t.responseText}),hv=eu("application/xml",function(t){var n=t.responseXML;if(!n)throw new Error("parse error");return n}),pv=ru("text/csv",Eh),dv=ru("text/tab-separated-values",Rh),vv=Array.prototype,gv=vv.map,_v=vv.slice,yv={name:"implicit"},mv=[0,1],xv=new Date,bv=new Date,wv=Cu(function(){},function(t,n){t.setTime(+t+n)},function(t,n){return n-t});wv.every=function(t){return t=Math.floor(t),isFinite(t)&&t>0?t>1?Cu(function(n){n.setTime(Math.floor(n/t)*t)},function(n,e){n.setTime(+n+e*t)},function(n,e){return(e-n)/t}):wv:null};var Mv=wv.range,Tv=6e4,Nv=6048e5,kv=Cu(function(t){t.setTime(1e3*Math.floor(t/1e3))},function(t,n){t.setTime(+t+1e3*n)},function(t,n){return(n-t)/1e3},function(t){return t.getUTCSeconds()}),Sv=kv.range,Ev=Cu(function(t){t.setTime(Math.floor(t/Tv)*Tv)},function(t,n){t.setTime(+t+n*Tv)},function(t,n){return(n-t)/Tv},function(t){return t.getMinutes()}),Av=Ev.range,Cv=Cu(function(t){var n=t.getTimezoneOffset()*Tv%36e5;n<0&&(n+=36e5),t.setTime(36e5*Math.floor((+t-n)/36e5)+n)},function(t,n){t.setTime(+t+36e5*n)},function(t,n){return(n-t)/36e5},function(t){return t.getHours()}),zv=Cv.range,Pv=Cu(function(t){t.setHours(0,0,0,0)},function(t,n){t.setDate(t.getDate()+n)},function(t,n){return(n-t-(n.getTimezoneOffset()-t.getTimezoneOffset())*Tv)/864e5},function(t){return t.getDate()-1}),Rv=Pv.range,Lv=zu(0),qv=zu(1),Dv=zu(2),Uv=zu(3),Ov=zu(4),Fv=zu(5),Iv=zu(6),Yv=Lv.range,Bv=qv.range,Hv=Dv.range,jv=Uv.range,Xv=Ov.range,Vv=Fv.range,$v=Iv.range,Wv=Cu(function(t){t.setDate(1),t.setHours(0,0,0,0)},function(t,n){t.setMonth(t.getMonth()+n)},function(t,n){return n.getMonth()-t.getMonth()+12*(n.getFullYear()-t.getFullYear())},function(t){return t.getMonth()}),Zv=Wv.range,Gv=Cu(function(t){t.setMonth(0,1),t.setHours(0,0,0,0)},function(t,n){t.setFullYear(t.getFullYear()+n)},function(t,n){return n.getFullYear()-t.getFullYear()},function(t){return t.getFullYear()});Gv.every=function(t){return isFinite(t=Math.floor(t))&&t>0?Cu(function(n){n.setFullYear(Math.floor(n.getFullYear()/t)*t),n.setMonth(0,1),n.setHours(0,0,0,0)},function(n,e){n.setFullYear(n.getFullYear()+e*t)}):null};var Qv=Gv.range,Jv=Cu(function(t){t.setUTCSeconds(0,0)},function(t,n){t.setTime(+t+n*Tv)},function(t,n){return(n-t)/Tv},function(t){return t.getUTCMinutes()}),Kv=Jv.range,tg=Cu(function(t){t.setUTCMinutes(0,0,0)},function(t,n){t.setTime(+t+36e5*n)},function(t,n){return(n-t)/36e5},function(t){return t.getUTCHours()}),ng=tg.range,eg=Cu(function(t){t.setUTCHours(0,0,0,0)},function(t,n){t.setUTCDate(t.getUTCDate()+n)},function(t,n){return(n-t)/864e5},function(t){return t.getUTCDate()-1}),rg=eg.range,ig=Pu(0),og=Pu(1),ug=Pu(2),ag=Pu(3),cg=Pu(4),sg=Pu(5),fg=Pu(6),lg=ig.range,hg=og.range,pg=ug.range,dg=ag.range,vg=cg.range,gg=sg.range,_g=fg.range,yg=Cu(function(t){t.setUTCDate(1),t.setUTCHours(0,0,0,0)},function(t,n){t.setUTCMonth(t.getUTCMonth()+n)},function(t,n){return n.getUTCMonth()-t.getUTCMonth()+12*(n.getUTCFullYear()-t.getUTCFullYear())},function(t){return t.getUTCMonth()}),mg=yg.range,xg=Cu(function(t){t.setUTCMonth(0,1),t.setUTCHours(0,0,0,0)},function(t,n){t.setUTCFullYear(t.getUTCFullYear()+n)},function(t,n){return n.getUTCFullYear()-t.getUTCFullYear()},function(t){return t.getUTCFullYear()});xg.every=function(t){return isFinite(t=Math.floor(t))&&t>0?Cu(function(n){n.setUTCFullYear(Math.floor(n.getUTCFullYear()/t)*t),n.setUTCMonth(0,1),n.setUTCHours(0,0,0,0)},function(n,e){n.setUTCFullYear(n.getUTCFullYear()+e*t)}):null};var bg,wg=xg.range,Mg={"-":"",_:" ",0:"0"},Tg=/^\s*\d+/,Ng=/^%/,kg=/[\\^$*+?|[\]().{}]/g;Ha({dateTime:"%x, %X",date:"%-m/%-d/%Y",time:"%-I:%M:%S %p",periods:["AM","PM"],days:["Sunday","Monday","Tuesday","Wednesday","Thursday","Friday","Saturday"],shortDays:["Sun","Mon","Tue","Wed","Thu","Fri","Sat"],months:["January","February","March","April","May","June","July","August","September","October","November","December"],shortMonths:["Jan","Feb","Mar","Apr","May","Jun","Jul","Aug","Sep","Oct","Nov","Dec"]});var Sg="%Y-%m-%dT%H:%M:%S.%LZ",Eg=Date.prototype.toISOString?function(t){return t.toISOString()}:t.utcFormat(Sg),Ag=+new Date("2000-01-01T00:00:00.000Z")?function(t){var n=new Date(t);return isNaN(n)?null:n}:t.utcParse(Sg),Cg=1e3,zg=60*Cg,Pg=60*zg,Rg=24*Pg,Lg=7*Rg,qg=30*Rg,Dg=365*Rg,Ug=$a("1f77b4ff7f0e2ca02cd627289467bd8c564be377c27f7f7fbcbd2217becf"),Og=$a("393b795254a36b6ecf9c9ede6379398ca252b5cf6bcedb9c8c6d31bd9e39e7ba52e7cb94843c39ad494ad6616be7969c7b4173a55194ce6dbdde9ed6"),Fg=$a("3182bd6baed69ecae1c6dbefe6550dfd8d3cfdae6bfdd0a231a35474c476a1d99bc7e9c0756bb19e9ac8bcbddcdadaeb636363969696bdbdbdd9d9d9"),Ig=$a("1f77b4aec7e8ff7f0effbb782ca02c98df8ad62728ff98969467bdc5b0d58c564bc49c94e377c2f7b6d27f7f7fc7c7c7bcbd22dbdb8d17becf9edae5"),Yg=al($t(300,.5,0),$t(-240,.5,1)),Bg=al($t(-100,.75,.35),$t(80,1.5,.8)),Hg=al($t(260,.75,.35),$t(80,1.5,.8)),jg=$t(),Xg=Wa($a("44015444025645045745055946075a46085c460a5d460b5e470d60470e6147106347116447136548146748166848176948186a481a6c481b6d481c6e481d6f481f70482071482173482374482475482576482677482878482979472a7a472c7a472d7b472e7c472f7d46307e46327e46337f463480453581453781453882443983443a83443b84433d84433e85423f854240864241864142874144874045884046883f47883f48893e49893e4a893e4c8a3d4d8a3d4e8a3c4f8a3c508b3b518b3b528b3a538b3a548c39558c39568c38588c38598c375a8c375b8d365c8d365d8d355e8d355f8d34608d34618d33628d33638d32648e32658e31668e31678e31688e30698e306a8e2f6b8e2f6c8e2e6d8e2e6e8e2e6f8e2d708e2d718e2c718e2c728e2c738e2b748e2b758e2a768e2a778e2a788e29798e297a8e297b8e287c8e287d8e277e8e277f8e27808e26818e26828e26828e25838e25848e25858e24868e24878e23888e23898e238a8d228b8d228c8d228d8d218e8d218f8d21908d21918c20928c20928c20938c1f948c1f958b1f968b1f978b1f988b1f998a1f9a8a1e9b8a1e9c891e9d891f9e891f9f881fa0881fa1881fa1871fa28720a38620a48621a58521a68522a78522a88423a98324aa8325ab8225ac8226ad8127ad8128ae8029af7f2ab07f2cb17e2db27d2eb37c2fb47c31b57b32b67a34b67935b77937b87838b9773aba763bbb753dbc743fbc7340bd7242be7144bf7046c06f48c16e4ac16d4cc26c4ec36b50c46a52c56954c56856c66758c7655ac8645cc8635ec96260ca6063cb5f65cb5e67cc5c69cd5b6ccd5a6ece5870cf5773d05675d05477d1537ad1517cd2507fd34e81d34d84d44b86d54989d5488bd6468ed64590d74393d74195d84098d83e9bd93c9dd93ba0da39a2da37a5db36a8db34aadc32addc30b0dd2fb2dd2db5de2bb8de29bade28bddf26c0df25c2df23c5e021c8e020cae11fcde11dd0e11cd2e21bd5e21ad8e219dae319dde318dfe318e2e418e5e419e7e419eae51aece51befe51cf1e51df4e61ef6e620f8e621fbe723fde725")),Vg=Wa($a("00000401000501010601010802010902020b02020d03030f03031204041405041606051806051a07061c08071e0907200a08220b09240c09260d0a290e0b2b100b2d110c2f120d31130d34140e36150e38160f3b180f3d19103f1a10421c10441d11471e114920114b21114e22115024125325125527125829115a2a115c2c115f2d11612f116331116533106734106936106b38106c390f6e3b0f703d0f713f0f72400f74420f75440f764510774710784910784a10794c117a4e117b4f127b51127c52137c54137d56147d57157e59157e5a167e5c167f5d177f5f187f601880621980641a80651a80671b80681c816a1c816b1d816d1d816e1e81701f81721f817320817521817621817822817922827b23827c23827e24828025828125818326818426818627818827818928818b29818c29818e2a81902a81912b81932b80942c80962c80982d80992d809b2e7f9c2e7f9e2f7fa02f7fa1307ea3307ea5317ea6317da8327daa337dab337cad347cae347bb0357bb2357bb3367ab5367ab73779b83779ba3878bc3978bd3977bf3a77c03a76c23b75c43c75c53c74c73d73c83e73ca3e72cc3f71cd4071cf4070d0416fd2426fd3436ed5446dd6456cd8456cd9466bdb476adc4869de4968df4a68e04c67e24d66e34e65e44f64e55064e75263e85362e95462ea5661eb5760ec5860ed5a5fee5b5eef5d5ef05f5ef1605df2625df2645cf3655cf4675cf4695cf56b5cf66c5cf66e5cf7705cf7725cf8745cf8765cf9785df9795df97b5dfa7d5efa7f5efa815ffb835ffb8560fb8761fc8961fc8a62fc8c63fc8e64fc9065fd9266fd9467fd9668fd9869fd9a6afd9b6bfe9d6cfe9f6dfea16efea36ffea571fea772fea973feaa74feac76feae77feb078feb27afeb47bfeb67cfeb77efeb97ffebb81febd82febf84fec185fec287fec488fec68afec88cfeca8dfecc8ffecd90fecf92fed194fed395fed597fed799fed89afdda9cfddc9efddea0fde0a1fde2a3fde3a5fde5a7fde7a9fde9aafdebacfcecaefceeb0fcf0b2fcf2b4fcf4b6fcf6b8fcf7b9fcf9bbfcfbbdfcfdbf")),$g=Wa($a("00000401000501010601010802010a02020c02020e03021004031204031405041706041907051b08051d09061f0a07220b07240c08260d08290e092b10092d110a30120a32140b34150b37160b39180c3c190c3e1b0c411c0c431e0c451f0c48210c4a230c4c240c4f260c51280b53290b552b0b572d0b592f0a5b310a5c320a5e340a5f3609613809623909633b09643d09653e0966400a67420a68440a68450a69470b6a490b6a4a0c6b4c0c6b4d0d6c4f0d6c510e6c520e6d540f6d550f6d57106e59106e5a116e5c126e5d126e5f136e61136e62146e64156e65156e67166e69166e6a176e6c186e6d186e6f196e71196e721a6e741a6e751b6e771c6d781c6d7a1d6d7c1d6d7d1e6d7f1e6c801f6c82206c84206b85216b87216b88226a8a226a8c23698d23698f24699025689225689326679526679727669827669a28659b29649d29649f2a63a02a63a22b62a32c61a52c60a62d60a82e5fa92e5eab2f5ead305dae305cb0315bb1325ab3325ab43359b63458b73557b93556ba3655bc3754bd3853bf3952c03a51c13a50c33b4fc43c4ec63d4dc73e4cc83f4bca404acb4149cc4248ce4347cf4446d04545d24644d34743d44842d54a41d74b3fd84c3ed94d3dda4e3cdb503bdd513ade5238df5337e05536e15635e25734e35933e45a31e55c30e65d2fe75e2ee8602de9612bea632aeb6429eb6628ec6726ed6925ee6a24ef6c23ef6e21f06f20f1711ff1731df2741cf3761bf37819f47918f57b17f57d15f67e14f68013f78212f78410f8850ff8870ef8890cf98b0bf98c0af98e09fa9008fa9207fa9407fb9606fb9706fb9906fb9b06fb9d07fc9f07fca108fca309fca50afca60cfca80dfcaa0ffcac11fcae12fcb014fcb216fcb418fbb61afbb81dfbba1ffbbc21fbbe23fac026fac228fac42afac62df9c72ff9c932f9cb35f8cd37f8cf3af7d13df7d340f6d543f6d746f5d949f5db4cf4dd4ff4df53f4e156f3e35af3e55df2e661f2e865f2ea69f1ec6df1ed71f1ef75f1f179f2f27df2f482f3f586f3f68af4f88ef5f992f6fa96f8fb9af9fc9dfafda1fcffa4")),Wg=Wa($a("0d088710078813078916078a19068c1b068d1d068e20068f2206902406912605912805922a05932c05942e05952f059631059733059735049837049938049a3a049a3c049b3e049c3f049c41049d43039e44039e46039f48039f4903a04b03a14c02a14e02a25002a25102a35302a35502a45601a45801a45901a55b01a55c01a65e01a66001a66100a76300a76400a76600a76700a86900a86a00a86c00a86e00a86f00a87100a87201a87401a87501a87701a87801a87a02a87b02a87d03a87e03a88004a88104a78305a78405a78606a68707a68808a68a09a58b0aa58d0ba58e0ca48f0da4910ea3920fa39410a29511a19613a19814a099159f9a169f9c179e9d189d9e199da01a9ca11b9ba21d9aa31e9aa51f99a62098a72197a82296aa2395ab2494ac2694ad2793ae2892b02991b12a90b22b8fb32c8eb42e8db52f8cb6308bb7318ab83289ba3388bb3488bc3587bd3786be3885bf3984c03a83c13b82c23c81c33d80c43e7fc5407ec6417dc7427cc8437bc9447aca457acb4679cc4778cc4977cd4a76ce4b75cf4c74d04d73d14e72d24f71d35171d45270d5536fd5546ed6556dd7566cd8576bd9586ada5a6ada5b69db5c68dc5d67dd5e66de5f65de6164df6263e06363e16462e26561e26660e3685fe4695ee56a5de56b5de66c5ce76e5be76f5ae87059e97158e97257ea7457eb7556eb7655ec7754ed7953ed7a52ee7b51ef7c51ef7e50f07f4ff0804ef1814df1834cf2844bf3854bf3874af48849f48948f58b47f58c46f68d45f68f44f79044f79143f79342f89441f89540f9973ff9983ef99a3efa9b3dfa9c3cfa9e3bfb9f3afba139fba238fca338fca537fca636fca835fca934fdab33fdac33fdae32fdaf31fdb130fdb22ffdb42ffdb52efeb72dfeb82cfeba2cfebb2bfebd2afebe2afec029fdc229fdc328fdc527fdc627fdc827fdca26fdcb26fccd25fcce25fcd025fcd225fbd324fbd524fbd724fad824fada24f9dc24f9dd25f8df25f8e125f7e225f7e425f6e626f6e826f5e926f5eb27f4ed27f3ee27f3f027f2f227f1f426f1f525f0f724f0f921")),Zg=Math.abs,Gg=Math.atan2,Qg=Math.cos,Jg=Math.max,Kg=Math.min,t_=Math.sin,n_=Math.sqrt,e_=1e-12,r_=Math.PI,i_=r_/2,o_=2*r_;ic.prototype={areaStart:function(){this._line=0},areaEnd:function(){this._line=NaN},lineStart:function(){this._point=0},lineEnd:function(){(this._line||0!==this._line&&1===this._point)&&this._context.closePath(),this._line=1-this._line},point:function(t,n){switch(t=+t,n=+n,this._point){case 0:this._point=1,this._line?this._context.lineTo(t,n):this._context.moveTo(t,n);break;case 1:this._point=2;default:this._context.lineTo(t,n)}}};var u_=pc(oc);hc.prototype={areaStart:function(){this._curve.areaStart()},areaEnd:function(){this._curve.areaEnd()},lineStart:function(){this._curve.lineStart()},lineEnd:function(){this._curve.lineEnd()},point:function(t,n){this._curve.point(n*Math.sin(t),n*-Math.cos(t))}};var a_=Array.prototype.slice,c_={draw:function(t,n){var e=Math.sqrt(n/r_);t.moveTo(e,0),t.arc(0,0,e,0,o_)}},s_={draw:function(t,n){var e=Math.sqrt(n/5)/2;t.moveTo(-3*e,-e),t.lineTo(-e,-e),t.lineTo(-e,-3*e),t.lineTo(e,-3*e),t.lineTo(e,-e),t.lineTo(3*e,-e),t.lineTo(3*e,e),t.lineTo(e,e),t.lineTo(e,3*e),t.lineTo(-e,3*e),t.lineTo(-e,e),t.lineTo(-3*e,e),t.closePath()}},f_=Math.sqrt(1/3),l_=2*f_,h_={draw:function(t,n){var e=Math.sqrt(n/l_),r=e*f_;t.moveTo(0,-e),t.lineTo(r,0),t.lineTo(0,e),t.lineTo(-r,0),t.closePath()}},p_=Math.sin(r_/10)/Math.sin(7*r_/10),d_=Math.sin(o_/10)*p_,v_=-Math.cos(o_/10)*p_,g_={draw:function(t,n){var e=Math.sqrt(.8908130915292852*n),r=d_*e,i=v_*e;t.moveTo(0,-e),t.lineTo(r,i);for(var o=1;o<5;++o){var u=o_*o/5,a=Math.cos(u),c=Math.sin(u);t.lineTo(c*e,-a*e),t.lineTo(a*r-c*i,c*r+a*i)}t.closePath()}},__={draw:function(t,n){var e=Math.sqrt(n),r=-e/2;t.rect(r,r,e,e)}},y_=Math.sqrt(3),m_={draw:function(t,n){var e=-Math.sqrt(n/(3*y_));t.moveTo(0,2*e),t.lineTo(-y_*e,-e),t.lineTo(y_*e,-e),t.closePath()}},x_=Math.sqrt(3)/2,b_=1/Math.sqrt(12),w_=3*(b_/2+1),M_={draw:function(t,n){var e=Math.sqrt(n/w_),r=e/2,i=e*b_,o=r,u=e*b_+e,a=-o,c=u;t.moveTo(r,i),t.lineTo(o,u),t.lineTo(a,c),t.lineTo(-.5*r-x_*i,x_*r+-.5*i),t.lineTo(-.5*o-x_*u,x_*o+-.5*u),t.lineTo(-.5*a-x_*c,x_*a+-.5*c),t.lineTo(-.5*r+x_*i,-.5*i-x_*r),t.lineTo(-.5*o+x_*u,-.5*u-x_*o),t.lineTo(-.5*a+x_*c,-.5*c-x_*a),t.closePath()}},T_=[c_,s_,h_,__,g_,m_,M_];kc.prototype={areaStart:function(){this._line=0},areaEnd:function(){this._line=NaN},lineStart:function(){this._x0=this._x1=this._y0=this._y1=NaN,this._point=0},lineEnd:function(){switch(this._point){case 3:Nc(this,this._x1,this._y1);case 2:this._context.lineTo(this._x1,this._y1)}(this._line||0!==this._line&&1===this._point)&&this._context.closePath(),this._line=1-this._line},point:function(t,n){switch(t=+t,n=+n,this._point){case 0:this._point=1,this._line?this._context.lineTo(t,n):this._context.moveTo(t,n);break;case 1:this._point=2;break;case 2:this._point=3,this._context.lineTo((5*this._x0+this._x1)/6,(5*this._y0+this._y1)/6);default:Nc(this,t,n)}this._x0=this._x1,this._x1=t,this._y0=this._y1,this._y1=n}},Sc.prototype={areaStart:Tc,areaEnd:Tc,lineStart:function(){this._x0=this._x1=this._x2=this._x3=this._x4=this._y0=this._y1=this._y2=this._y3=this._y4=NaN,this._point=0},lineEnd:function(){switch(this._point){case 1:this._context.moveTo(this._x2,this._y2),this._context.closePath();break;case 2:this._context.moveTo((this._x2+2*this._x3)/3,(this._y2+2*this._y3)/3),this._context.lineTo((this._x3+2*this._x2)/3,(this._y3+2*this._y2)/3),this._context.closePath();break;case 3:this.point(this._x2,this._y2),this.point(this._x3,this._y3),this.point(this._x4,this._y4)}},point:function(t,n){switch(t=+t,n=+n,this._point){case 0:this._point=1,this._x2=t,this._y2=n;break;case 1:this._point=2,this._x3=t,this._y3=n;break;case 2:this._point=3,this._x4=t,this._y4=n,this._context.moveTo((this._x0+4*this._x1+t)/6,(this._y0+4*this._y1+n)/6);break;default:Nc(this,t,n)}this._x0=this._x1,this._x1=t,this._y0=this._y1,this._y1=n}},Ec.prototype={areaStart:function(){this._line=0},areaEnd:function(){this._line=NaN},lineStart:function(){this._x0=this._x1=this._y0=this._y1=NaN,this._point=0},lineEnd:function(){(this._line||0!==this._line&&3===this._point)&&this._context.closePath(),this._line=1-this._line},point:function(t,n){switch(t=+t,n=+n,this._point){case 0:this._point=1;break;case 1:this._point=2;break;case 2:this._point=3;var e=(this._x0+4*this._x1+t)/6,r=(this._y0+4*this._y1+n)/6;this._line?this._context.lineTo(e,r):this._context.moveTo(e,r);break;case 3:this._point=4;default:Nc(this,t,n)}this._x0=this._x1,this._x1=t,this._y0=this._y1,this._y1=n}},Ac.prototype={lineStart:function(){this._x=[],this._y=[],this._basis.lineStart()},lineEnd:function(){var t=this._x,n=this._y,e=t.length-1;if(e>0)for(var r,i=t[0],o=n[0],u=t[e]-i,a=n[e]-o,c=-1;++c<=e;)r=c/e,this._basis.point(this._beta*t[c]+(1-this._beta)*(i+r*u),this._beta*n[c]+(1-this._beta)*(o+r*a));this._x=this._y=null,this._basis.lineEnd()},point:function(t,n){this._x.push(+t),this._y.push(+n)}};var N_=function t(n){function e(t){return 1===n?new kc(t):new Ac(t,n)}return e.beta=function(n){return t(+n)},e}(.85);zc.prototype={areaStart:function(){this._line=0},areaEnd:function(){this._line=NaN},lineStart:function(){this._x0=this._x1=this._x2=this._y0=this._y1=this._y2=NaN,this._point=0},lineEnd:function(){switch(this._point){case 2:this._context.lineTo(this._x2,this._y2);break;case 3:Cc(this,this._x1,this._y1)}(this._line||0!==this._line&&1===this._point)&&this._context.closePath(),this._line=1-this._line},point:function(t,n){switch(t=+t,n=+n,this._point){case 0:this._point=1,this._line?this._context.lineTo(t,n):this._context.moveTo(t,n);break;case 1:this._point=2,this._x1=t,this._y1=n;break;case 2:this._point=3;default:Cc(this,t,n)}this._x0=this._x1,this._x1=this._x2,this._x2=t,this._y0=this._y1,this._y1=this._y2,this._y2=n}};var k_=function t(n){function e(t){return new zc(t,n)}return e.tension=function(n){return t(+n)},e}(0);Pc.prototype={areaStart:Tc,areaEnd:Tc,lineStart:function(){this._x0=this._x1=this._x2=this._x3=this._x4=this._x5=this._y0=this._y1=this._y2=this._y3=this._y4=this._y5=NaN,this._point=0},lineEnd:function(){switch(this._point){case 1:this._context.moveTo(this._x3,this._y3),this._context.closePath();break;case 2:this._context.lineTo(this._x3,this._y3),this._context.closePath();break;case 3:this.point(this._x3,this._y3),this.point(this._x4,this._y4),this.point(this._x5,this._y5)}},point:function(t,n){switch(t=+t,n=+n,this._point){case 0:this._point=1,this._x3=t,this._y3=n;break;case 1:this._point=2,this._context.moveTo(this._x4=t,this._y4=n);break;case 2:this._point=3,this._x5=t,this._y5=n;break;default:Cc(this,t,n)}this._x0=this._x1,this._x1=this._x2,this._x2=t,this._y0=this._y1,this._y1=this._y2,this._y2=n}};var S_=function t(n){function e(t){return new Pc(t,n)}return e.tension=function(n){return t(+n)},e}(0);Rc.prototype={areaStart:function(){this._line=0},areaEnd:function(){this._line=NaN},lineStart:function(){this._x0=this._x1=this._x2=this._y0=this._y1=this._y2=NaN,this._point=0},lineEnd:function(){(this._line||0!==this._line&&3===this._point)&&this._context.closePath(),this._line=1-this._line},point:function(t,n){switch(t=+t,n=+n,this._point){case 0:this._point=1;break;case 1:this._point=2;break;case 2:this._point=3,this._line?this._context.lineTo(this._x2,this._y2):this._context.moveTo(this._x2,this._y2);break;case 3:this._point=4;default:Cc(this,t,n)}this._x0=this._x1,this._x1=this._x2,this._x2=t,this._y0=this._y1,this._y1=this._y2,this._y2=n}};var E_=function t(n){function e(t){return new Rc(t,n)}return e.tension=function(n){return t(+n)},e}(0);qc.prototype={areaStart:function(){this._line=0},areaEnd:function(){this._line=NaN},lineStart:function(){this._x0=this._x1=this._x2=this._y0=this._y1=this._y2=NaN,this._l01_a=this._l12_a=this._l23_a=this._l01_2a=this._l12_2a=this._l23_2a=this._point=0},lineEnd:function(){switch(this._point){case 2:this._context.lineTo(this._x2,this._y2);break;case 3:this.point(this._x2,this._y2)}(this._line||0!==this._line&&1===this._point)&&this._context.closePath(),this._line=1-this._line},point:function(t,n){if(t=+t,n=+n,this._point){var e=this._x2-t,r=this._y2-n;this._l23_a=Math.sqrt(this._l23_2a=Math.pow(e*e+r*r,this._alpha))}switch(this._point){case 0:this._point=1,this._line?this._context.lineTo(t,n):this._context.moveTo(t,n);break;case 1:this._point=2;break;case 2:this._point=3;default:Lc(this,t,n)}this._l01_a=this._l12_a,this._l12_a=this._l23_a,this._l01_2a=this._l12_2a,this._l12_2a=this._l23_2a,this._x0=this._x1,this._x1=this._x2,this._x2=t,this._y0=this._y1,this._y1=this._y2,this._y2=n}};var A_=function t(n){function e(t){return n?new qc(t,n):new zc(t,0)}return e.alpha=function(n){return t(+n)},e}(.5);Dc.prototype={areaStart:Tc,areaEnd:Tc,lineStart:function(){this._x0=this._x1=this._x2=this._x3=this._x4=this._x5=this._y0=this._y1=this._y2=this._y3=this._y4=this._y5=NaN,this._l01_a=this._l12_a=this._l23_a=this._l01_2a=this._l12_2a=this._l23_2a=this._point=0},lineEnd:function(){switch(this._point){case 1:this._context.moveTo(this._x3,this._y3),this._context.closePath();break;case 2:this._context.lineTo(this._x3,this._y3),this._context.closePath();break;case 3:this.point(this._x3,this._y3),this.point(this._x4,this._y4),this.point(this._x5,this._y5)}},point:function(t,n){if(t=+t,n=+n,this._point){var e=this._x2-t,r=this._y2-n;this._l23_a=Math.sqrt(this._l23_2a=Math.pow(e*e+r*r,this._alpha))}switch(this._point){case 0:this._point=1,this._x3=t,this._y3=n;break;case 1:this._point=2,this._context.moveTo(this._x4=t,this._y4=n);break;case 2:this._point=3,this._x5=t,this._y5=n;break;default:Lc(this,t,n)}this._l01_a=this._l12_a,this._l12_a=this._l23_a,this._l01_2a=this._l12_2a,this._l12_2a=this._l23_2a,this._x0=this._x1,this._x1=this._x2,this._x2=t,this._y0=this._y1,this._y1=this._y2,this._y2=n}};var C_=function t(n){function e(t){return n?new Dc(t,n):new Pc(t,0)}return e.alpha=function(n){return t(+n)},e}(.5);Uc.prototype={areaStart:function(){this._line=0},areaEnd:function(){this._line=NaN},lineStart:function(){this._x0=this._x1=this._x2=this._y0=this._y1=this._y2=NaN,this._l01_a=this._l12_a=this._l23_a=this._l01_2a=this._l12_2a=this._l23_2a=this._point=0},lineEnd:function(){(this._line||0!==this._line&&3===this._point)&&this._context.closePath(),this._line=1-this._line},point:function(t,n){if(t=+t,n=+n,this._point){var e=this._x2-t,r=this._y2-n;this._l23_a=Math.sqrt(this._l23_2a=Math.pow(e*e+r*r,this._alpha))}switch(this._point){case 0:this._point=1;break;case 1:this._point=2;break;case 2:this._point=3,this._line?this._context.lineTo(this._x2,this._y2):this._context.moveTo(this._x2,this._y2);break;case 3:this._point=4;default:Lc(this,t,n)}this._l01_a=this._l12_a,this._l12_a=this._l23_a,this._l01_2a=this._l12_2a,this._l12_2a=this._l23_2a,this._x0=this._x1,this._x1=this._x2,this._x2=t,this._y0=this._y1,this._y1=this._y2,this._y2=n}};var z_=function t(n){function e(t){return n?new Uc(t,n):new Rc(t,0)}return e.alpha=function(n){return t(+n)},e}(.5);Oc.prototype={areaStart:Tc,areaEnd:Tc,lineStart:function(){this._point=0},lineEnd:function(){this._point&&this._context.closePath()},point:function(t,n){t=+t,n=+n,this._point?this._context.lineTo(t,n):(this._point=1,this._context.moveTo(t,n))}},Hc.prototype={areaStart:function(){this._line=0},areaEnd:function(){this._line=NaN},lineStart:function(){this._x0=this._x1=this._y0=this._y1=this._t0=NaN,this._point=0},lineEnd:function(){switch(this._point){case 2:this._context.lineTo(this._x1,this._y1);break;case 3:Bc(this,this._t0,Yc(this,this._t0))}(this._line||0!==this._line&&1===this._point)&&this._context.closePath(),this._line=1-this._line},point:function(t,n){var e=NaN;if(t=+t,n=+n,t!==this._x1||n!==this._y1){switch(this._point){case 0:this._point=1,this._line?this._context.lineTo(t,n):this._context.moveTo(t,n);break;case 1:this._point=2;break;case 2:this._point=3,Bc(this,Yc(this,e=Ic(this,t,n)),e);break;default:Bc(this,this._t0,e=Ic(this,t,n))}this._x0=this._x1,this._x1=t,this._y0=this._y1,this._y1=n,this._t0=e}}},(jc.prototype=Object.create(Hc.prototype)).point=function(t,n){Hc.prototype.point.call(this,n,t)},Xc.prototype={moveTo:function(t,n){this._context.moveTo(n,t)},closePath:function(){this._context.closePath()},lineTo:function(t,n){this._context.lineTo(n,t)},bezierCurveTo:function(t,n,e,r,i,o){this._context.bezierCurveTo(n,t,r,e,o,i)}},Vc.prototype={areaStart:function(){this._line=0},areaEnd:function(){this._line=NaN},lineStart:function(){this._x=[],this._y=[]},lineEnd:function(){var t=this._x,n=this._y,e=t.length;if(e)if(this._line?this._context.lineTo(t[0],n[0]):this._context.moveTo(t[0],n[0]),2===e)this._context.lineTo(t[1],n[1]);else for(var r=$c(t),i=$c(n),o=0,u=1;u<e;++o,++u)this._context.bezierCurveTo(r[0][o],i[0][o],r[1][o],i[1][o],t[u],n[u]);(this._line||0!==this._line&&1===e)&&this._context.closePath(),this._line=1-this._line,this._x=this._y=null},point:function(t,n){this._x.push(+t),this._y.push(+n)}},Wc.prototype={areaStart:function(){this._line=0},areaEnd:function(){this._line=NaN},lineStart:function(){this._x=this._y=NaN,this._point=0},lineEnd:function(){0<this._t&&this._t<1&&2===this._point&&this._context.lineTo(this._x,this._y),(this._line||0!==this._line&&1===this._point)&&this._context.closePath(),this._line>=0&&(this._t=1-this._t,this._line=1-this._line)},point:function(t,n){switch(t=+t,n=+n,this._point){case 0:this._point=1,this._line?this._context.lineTo(t,n):this._context.moveTo(t,n);break;case 1:this._point=2;default:if(this._t<=0)this._context.lineTo(this._x,n),this._context.lineTo(t,n);else{var e=this._x*(1-this._t)+t*this._t;this._context.lineTo(e,this._y),this._context.lineTo(e,n)}}this._x=t,this._y=n}},rs.prototype={constructor:rs,insert:function(t,n){var e,r,i;if(t){if(n.P=t,n.N=t.N,t.N&&(t.N.P=n),t.N=n,t.R){for(t=t.R;t.L;)t=t.L;t.L=n}else t.R=n;e=t}else this._?(t=as(this._),n.P=null,n.N=t,t.P=t.L=n,e=t):(n.P=n.N=null,this._=n,e=null);for(n.L=n.R=null,n.U=e,n.C=!0,t=n;e&&e.C;)e===(r=e.U).L?(i=r.R)&&i.C?(e.C=i.C=!1,r.C=!0,t=r):(t===e.R&&(os(this,e),e=(t=e).U),e.C=!1,r.C=!0,us(this,r)):(i=r.L)&&i.C?(e.C=i.C=!1,r.C=!0,t=r):(t===e.L&&(us(this,e),e=(t=e).U),e.C=!1,r.C=!0,os(this,r)),e=t.U;this._.C=!1},remove:function(t){t.N&&(t.N.P=t.P),t.P&&(t.P.N=t.N),t.N=t.P=null;var n,e,r,i=t.U,o=t.L,u=t.R;if(e=o?u?as(u):o:u,i?i.L===t?i.L=e:i.R=e:this._=e,o&&u?(r=e.C,e.C=t.C,e.L=o,o.U=e,e!==u?(i=e.U,e.U=t.U,t=e.R,i.L=t,e.R=u,u.U=e):(e.U=i,i=e,t=e.R)):(r=t.C,t=e),t&&(t.U=i),!r)if(t&&t.C)t.C=!1;else{do{if(t===this._)break;if(t===i.L){if((n=i.R).C&&(n.C=!1,i.C=!0,os(this,i),n=i.R),n.L&&n.L.C||n.R&&n.R.C){n.R&&n.R.C||(n.L.C=!1,n.C=!0,us(this,n),n=i.R),n.C=i.C,i.C=n.R.C=!1,os(this,i),t=this._;break}}else if((n=i.L).C&&(n.C=!1,i.C=!0,us(this,i),n=i.L),n.L&&n.L.C||n.R&&n.R.C){n.L&&n.L.C||(n.R.C=!1,n.C=!0,os(this,n),n=i.L),n.C=i.C,i.C=n.L.C=!1,us(this,i),t=this._;break}n.C=!0,t=i,i=i.U}while(!t.C);t&&(t.C=!1)}}};var P_,R_,L_,q_,D_,U_=[],O_=[],F_=1e-6,I_=1e-12;Ns.prototype={constructor:Ns,polygons:function(){var t=this.edges;return this.cells.map(function(n){var e=n.halfedges.map(function(e){return ds(n,t[e])});return e.data=n.site.data,e})},triangles:function(){var t=[],n=this.edges;return this.cells.forEach(function(e,r){if(o=(i=e.halfedges).length)for(var i,o,u,a=e.site,c=-1,s=n[i[o-1]],f=s.left===a?s.right:s.left;++c<o;)u=f,f=(s=n[i[c]]).left===a?s.right:s.left,u&&f&&r<u.index&&r<f.index&&Ms(a,u,f)<0&&t.push([a.data,u.data,f.data])}),t},links:function(){return this.edges.filter(function(t){return t.right}).map(function(t){return{source:t.left.data,target:t.right.data}})},find:function(t,n,e){for(var r,i,o=this,u=o._found||0,a=o.cells.length;!(i=o.cells[u]);)if(++u>=a)return null;var c=t-i.site[0],s=n-i.site[1],f=c*c+s*s;do{i=o.cells[r=u],u=null,i.halfedges.forEach(function(e){var r=o.edges[e],a=r.left;if(a!==i.site&&a||(a=r.right)){var c=t-a[0],s=n-a[1],l=c*c+s*s;l<f&&(f=l,u=a.index)}})}while(null!==u);return o._found=r,null==e||f<=e*e?i.site:null}},Ss.prototype={constructor:Ss,scale:function(t){return 1===t?this:new Ss(this.k*t,this.x,this.y)},translate:function(t,n){return 0===t&0===n?this:new Ss(this.k,this.x+this.k*t,this.y+this.k*n)},apply:function(t){return[t[0]*this.k+this.x,t[1]*this.k+this.y]},applyX:function(t){return t*this.k+this.x},applyY:function(t){return t*this.k+this.y},invert:function(t){return[(t[0]-this.x)/this.k,(t[1]-this.y)/this.k]},invertX:function(t){return(t-this.x)/this.k},invertY:function(t){return(t-this.y)/this.k},rescaleX:function(t){return t.copy().domain(t.range().map(this.invertX,this).map(t.invert,t))},rescaleY:function(t){return t.copy().domain(t.range().map(this.invertY,this).map(t.invert,t))},toString:function(){return"translate("+this.x+","+this.y+") scale("+this.k+")"}};var Y_=new Ss(1,0,0);Es.prototype=Ss.prototype,t.version="4.13.0",t.bisect=Os,t.bisectRight=Os,t.bisectLeft=Fs,t.ascending=n,t.bisector=e,t.cross=function(t,n,e){var i,o,u,a,c=t.length,s=n.length,f=new Array(c*s);for(null==e&&(e=r),i=u=0;i<c;++i)for(a=t[i],o=0;o<s;++o,++u)f[u]=e(a,n[o]);return f},t.descending=function(t,n){return n<t?-1:n>t?1:n>=t?0:NaN},t.deviation=u,t.extent=a,t.histogram=function(){function t(t){var i,o,u=t.length,a=new Array(u);for(i=0;i<u;++i)a[i]=n(t[i],i,t);var c=e(a),s=c[0],l=c[1],h=r(a,s,l);Array.isArray(h)||(h=p(s,l,h),h=f(Math.ceil(s/h)*h,Math.floor(l/h)*h,h));for(var d=h.length;h[0]<=s;)h.shift(),--d;for(;h[d-1]>l;)h.pop(),--d;var v,g=new Array(d+1);for(i=0;i<=d;++i)(v=g[i]=[]).x0=i>0?h[i-1]:s,v.x1=i<d?h[i]:l;for(i=0;i<u;++i)s<=(o=a[i])&&o<=l&&g[Os(h,o,0,d)].push(t[i]);return g}var n=s,e=a,r=d;return t.value=function(e){return arguments.length?(n="function"==typeof e?e:c(e),t):n},t.domain=function(n){return arguments.length?(e="function"==typeof n?n:c([n[0],n[1]]),t):e},t.thresholds=function(n){return arguments.length?(r="function"==typeof n?n:Array.isArray(n)?c(Ys.call(n)):c(n),t):r},t},t.thresholdFreedmanDiaconis=function(t,e,r){return t=Bs.call(t,i).sort(n),Math.ceil((r-e)/(2*(v(t,.75)-v(t,.25))*Math.pow(t.length,-1/3)))},t.thresholdScott=function(t,n,e){return Math.ceil((e-n)/(3.5*u(t)*Math.pow(t.length,-1/3)))},t.thresholdSturges=d,t.max=function(t,n){var e,r,i=t.length,o=-1;if(null==n){for(;++o<i;)if(null!=(e=t[o])&&e>=e)for(r=e;++o<i;)null!=(e=t[o])&&e>r&&(r=e)}else for(;++o<i;)if(null!=(e=n(t[o],o,t))&&e>=e)for(r=e;++o<i;)null!=(e=n(t[o],o,t))&&e>r&&(r=e);return r},t.mean=function(t,n){var e,r=t.length,o=r,u=-1,a=0;if(null==n)for(;++u<r;)isNaN(e=i(t[u]))?--o:a+=e;else for(;++u<r;)isNaN(e=i(n(t[u],u,t)))?--o:a+=e;if(o)return a/o},t.median=function(t,e){var r,o=t.length,u=-1,a=[];if(null==e)for(;++u<o;)isNaN(r=i(t[u]))||a.push(r);else for(;++u<o;)isNaN(r=i(e(t[u],u,t)))||a.push(r);return v(a.sort(n),.5)},t.merge=g,t.min=_,t.pairs=function(t,n){null==n&&(n=r);for(var e=0,i=t.length-1,o=t[0],u=new Array(i<0?0:i);e<i;)u[e]=n(o,o=t[++e]);return u},t.permute=function(t,n){for(var e=n.length,r=new Array(e);e--;)r[e]=t[n[e]];return r},t.quantile=v,t.range=f,t.scan=function(t,e){if(r=t.length){var r,i,o=0,u=0,a=t[u];for(null==e&&(e=n);++o<r;)(e(i=t[o],a)<0||0!==e(a,a))&&(a=i,u=o);return 0===e(a,a)?u:void 0}},t.shuffle=function(t,n,e){for(var r,i,o=(null==e?t.length:e)-(n=null==n?0:+n);o;)i=Math.random()*o--|0,r=t[o+n],t[o+n]=t[i+n],t[i+n]=r;return t},t.sum=function(t,n){var e,r=t.length,i=-1,o=0;if(null==n)for(;++i<r;)(e=+t[i])&&(o+=e);else for(;++i<r;)(e=+n(t[i],i,t))&&(o+=e);return o},t.ticks=l,t.tickIncrement=h,t.tickStep=p,t.transpose=y,t.variance=o,t.zip=function(){return y(arguments)},t.axisTop=function(t){return T($s,t)},t.axisRight=function(t){return T(Ws,t)},t.axisBottom=function(t){return T(Zs,t)},t.axisLeft=function(t){return T(Gs,t)},t.brush=function(){return Kn(oh)},t.brushX=function(){return Kn(rh)},t.brushY=function(){return Kn(ih)},t.brushSelection=function(t){var n=t.__brush;return n?n.dim.output(n.selection):null},t.chord=function(){function t(t){var o,u,a,c,s,l,h=t.length,p=[],d=f(h),v=[],g=[],_=g.groups=new Array(h),y=new Array(h*h);for(o=0,s=-1;++s<h;){for(u=0,l=-1;++l<h;)u+=t[s][l];p.push(u),v.push(f(h)),o+=u}for(e&&d.sort(function(t,n){return e(p[t],p[n])}),r&&v.forEach(function(n,e){n.sort(function(n,i){return r(t[e][n],t[e][i])})}),c=(o=gh(0,vh-n*h)/o)?n:vh/h,u=0,s=-1;++s<h;){for(a=u,l=-1;++l<h;){var m=d[s],x=v[m][l],b=t[m][x],w=u,M=u+=b*o;y[x*h+m]={index:m,subindex:x,startAngle:w,endAngle:M,value:b}}_[m]={index:m,startAngle:a,endAngle:u,value:p[m]},u+=c}for(s=-1;++s<h;)for(l=s-1;++l<h;){var T=y[l*h+s],N=y[s*h+l];(T.value||N.value)&&g.push(T.value<N.value?{source:N,target:T}:{source:T,target:N})}return i?g.sort(i):g}var n=0,e=null,r=null,i=null;return t.padAngle=function(e){return arguments.length?(n=gh(0,e),t):n},t.sortGroups=function(n){return arguments.length?(e=n,t):e},t.sortSubgroups=function(n){return arguments.length?(r=n,t):r},t.sortChords=function(n){return arguments.length?(null==n?i=null:(i=function(t){return function(n,e){return t(n.source.value+n.target.value,e.source.value+e.target.value)}}(n))._=n,t):i&&i._},t},t.ribbon=function(){function t(){var t,a=_h.call(arguments),c=n.apply(this,a),s=e.apply(this,a),f=+r.apply(this,(a[0]=c,a)),l=i.apply(this,a)-dh,h=o.apply(this,a)-dh,p=f*lh(l),d=f*hh(l),v=+r.apply(this,(a[0]=s,a)),g=i.apply(this,a)-dh,_=o.apply(this,a)-dh;if(u||(u=t=ee()),u.moveTo(p,d),u.arc(0,0,f,l,h),l===g&&h===_||(u.quadraticCurveTo(0,0,v*lh(g),v*hh(g)),u.arc(0,0,v,g,_)),u.quadraticCurveTo(0,0,p,d),u.closePath(),t)return u=null,t+""||null}var n=re,e=ie,r=oe,i=ue,o=ae,u=null;return t.radius=function(n){return arguments.length?(r="function"==typeof n?n:te(+n),t):r},t.startAngle=function(n){return arguments.length?(i="function"==typeof n?n:te(+n),t):i},t.endAngle=function(n){return arguments.length?(o="function"==typeof n?n:te(+n),t):o},t.source=function(e){return arguments.length?(n=e,t):n},t.target=function(n){return arguments.length?(e=n,t):e},t.context=function(n){return arguments.length?(u=null==n?null:n,t):u},t},t.nest=function(){function t(n,i,u,a){if(i>=o.length)return null!=e&&n.sort(e),null!=r?r(n):n;for(var c,s,f,l=-1,h=n.length,p=o[i++],d=se(),v=u();++l<h;)(f=d.get(c=p(s=n[l])+""))?f.push(s):d.set(c,[s]);return d.each(function(n,e){a(v,e,t(n,i,u,a))}),v}function n(t,e){if(++e>o.length)return t;var i,a=u[e-1];return null!=r&&e>=o.length?i=t.entries():(i=[],t.each(function(t,r){i.push({key:r,values:n(t,e)})})),null!=a?i.sort(function(t,n){return a(t.key,n.key)}):i}var e,r,i,o=[],u=[];return i={object:function(n){return t(n,0,fe,le)},map:function(n){return t(n,0,he,pe)},entries:function(e){return n(t(e,0,he,pe),0)},key:function(t){return o.push(t),i},sortKeys:function(t){return u[o.length-1]=t,i},sortValues:function(t){return e=t,i},rollup:function(t){return r=t,i}}},t.set=ve,t.map=se,t.keys=function(t){var n=[];for(var e in t)n.push(e);return n},t.values=function(t){var n=[];for(var e in t)n.push(t[e]);return n},t.entries=function(t){var n=[];for(var e in t)n.push({key:e,value:t[e]});return n},t.color=Et,t.rgb=Pt,t.hsl=qt,t.lab=Ft,t.hcl=Xt,t.cubehelix=$t,t.dispatch=N,t.drag=function(){function n(t){t.on("mousedown.drag",e).filter(g).on("touchstart.drag",o).on("touchmove.drag",u).on("touchend.drag touchcancel.drag",a).style("touch-action","none").style("-webkit-tap-highlight-color","rgba(0,0,0,0)")}function e(){if(!h&&p.apply(this,arguments)){var n=c("mouse",d.apply(this,arguments),pt,this,arguments);n&&(ct(t.event.view).on("mousemove.drag",r,!0).on("mouseup.drag",i,!0),_t(t.event.view),vt(),l=!1,s=t.event.clientX,f=t.event.clientY,n("start"))}}function r(){if(gt(),!l){var n=t.event.clientX-s,e=t.event.clientY-f;l=n*n+e*e>x}_.mouse("drag")}function i(){ct(t.event.view).on("mousemove.drag mouseup.drag",null),yt(t.event.view,l),gt(),_.mouse("end")}function o(){if(p.apply(this,arguments)){var n,e,r=t.event.changedTouches,i=d.apply(this,arguments),o=r.length;for(n=0;n<o;++n)(e=c(r[n].identifier,i,dt,this,arguments))&&(vt(),e("start"))}}function u(){var n,e,r=t.event.changedTouches,i=r.length;for(n=0;n<i;++n)(e=_[r[n].identifier])&&(gt(),e("drag"))}function a(){var n,e,r=t.event.changedTouches,i=r.length;for(h&&clearTimeout(h),h=setTimeout(function(){h=null},500),n=0;n<i;++n)(e=_[r[n].identifier])&&(vt(),e("end"))}function c(e,r,i,o,u){var a,c,s,f=i(r,e),l=y.copy();if(it(new xt(n,"beforestart",a,e,m,f[0],f[1],0,0,l),function(){return null!=(t.event.subject=a=v.apply(o,u))&&(c=a.x-f[0]||0,s=a.y-f[1]||0,!0)}))return function t(h){var p,d=f;switch(h){case"start":_[e]=t,p=m++;break;case"end":delete _[e],--m;case"drag":f=i(r,e),p=m}it(new xt(n,h,a,e,p,f[0]+c,f[1]+s,f[0]-d[0],f[1]-d[1],l),l.apply,l,[h,o,u])}}var s,f,l,h,p=bt,d=wt,v=Mt,g=Tt,_={},y=N("start","drag","end"),m=0,x=0;return n.filter=function(t){return arguments.length?(p="function"==typeof t?t:mt(!!t),n):p},n.container=function(t){return arguments.length?(d="function"==typeof t?t:mt(t),n):d},n.subject=function(t){return arguments.length?(v="function"==typeof t?t:mt(t),n):v},n.touchable=function(t){return arguments.length?(g="function"==typeof t?t:mt(!!t),n):g},n.on=function(){var t=y.on.apply(y,arguments);return t===y?n:t},n.clickDistance=function(t){return arguments.length?(x=(t=+t)*t,n):Math.sqrt(x)},n},t.dragDisable=_t,t.dragEnable=yt,t.dsvFormat=_e,t.csvParse=Eh,t.csvParseRows=Ah,t.csvFormat=Ch,t.csvFormatRows=zh,t.tsvParse=Rh,t.tsvParseRows=Lh,t.tsvFormat=qh,t.tsvFormatRows=Dh,t.easeLinear=function(t){return+t},t.easeQuad=On,t.easeQuadIn=function(t){return t*t},t.easeQuadOut=function(t){return t*(2-t)},t.easeQuadInOut=On,t.easeCubic=Fn,t.easeCubicIn=function(t){return t*t*t},t.easeCubicOut=function(t){return--t*t*t+1},t.easeCubicInOut=Fn,t.easePoly=zl,t.easePolyIn=Al,t.easePolyOut=Cl,t.easePolyInOut=zl,t.easeSin=In,t.easeSinIn=function(t){return 1-Math.cos(t*Rl)},t.easeSinOut=function(t){return Math.sin(t*Rl)},t.easeSinInOut=In,t.easeExp=Yn,t.easeExpIn=function(t){return Math.pow(2,10*t-10)},t.easeExpOut=function(t){return 1-Math.pow(2,-10*t)},t.easeExpInOut=Yn,t.easeCircle=Bn,t.easeCircleIn=function(t){return 1-Math.sqrt(1-t*t)},t.easeCircleOut=function(t){return Math.sqrt(1- --t*t)},t.easeCircleInOut=Bn,t.easeBounce=Hn,t.easeBounceIn=function(t){return 1-Hn(1-t)},t.easeBounceOut=Hn,t.easeBounceInOut=function(t){return((t*=2)<=1?1-Hn(1-t):Hn(t-1)+1)/2},t.easeBack=Vl,t.easeBackIn=jl,t.easeBackOut=Xl,t.easeBackInOut=Vl,t.easeElastic=Zl,t.easeElasticIn=Wl,t.easeElasticOut=Zl,t.easeElasticInOut=Gl,t.forceCenter=function(t,n){function e(){var e,i,o=r.length,u=0,a=0;for(e=0;e<o;++e)u+=(i=r[e]).x,a+=i.y;for(u=u/o-t,a=a/o-n,e=0;e<o;++e)(i=r[e]).x-=u,i.y-=a}var r;return null==t&&(t=0),null==n&&(n=0),e.initialize=function(t){r=t},e.x=function(n){return arguments.length?(t=+n,e):t},e.y=function(t){return arguments.length?(n=+t,e):n},e},t.forceCollide=function(t){function n(){for(var t,n,r,c,s,f,l,h=i.length,p=0;p<a;++p)for(n=Te(i,Se,Ee).visitAfter(e),t=0;t<h;++t)r=i[t],f=o[r.index],l=f*f,c=r.x+r.vx,s=r.y+r.vy,n.visit(function(t,n,e,i,o){var a=t.data,h=t.r,p=f+h;if(!a)return n>c+p||i<c-p||e>s+p||o<s-p;if(a.index>r.index){var d=c-a.x-a.vx,v=s-a.y-a.vy,g=d*d+v*v;g<p*p&&(0===d&&(d=me(),g+=d*d),0===v&&(v=me(),g+=v*v),g=(p-(g=Math.sqrt(g)))/g*u,r.vx+=(d*=g)*(p=(h*=h)/(l+h)),r.vy+=(v*=g)*p,a.vx-=d*(p=1-p),a.vy-=v*p)}})}function e(t){if(t.data)return t.r=o[t.data.index];for(var n=t.r=0;n<4;++n)t[n]&&t[n].r>t.r&&(t.r=t[n].r)}function r(){if(i){var n,e,r=i.length;for(o=new Array(r),n=0;n<r;++n)e=i[n],o[e.index]=+t(e,n,i)}}var i,o,u=1,a=1;return"function"!=typeof t&&(t=ye(null==t?1:+t)),n.initialize=function(t){i=t,r()},n.iterations=function(t){return arguments.length?(a=+t,n):a},n.strength=function(t){return arguments.length?(u=+t,n):u},n.radius=function(e){return arguments.length?(t="function"==typeof e?e:ye(+e),r(),n):t},n},t.forceLink=function(t){function n(n){for(var e=0,r=t.length;e<p;++e)for(var i,a,c,f,l,h,d,v=0;v<r;++v)a=(i=t[v]).source,f=(c=i.target).x+c.vx-a.x-a.vx||me(),l=c.y+c.vy-a.y-a.vy||me(),f*=h=((h=Math.sqrt(f*f+l*l))-u[v])/h*n*o[v],l*=h,c.vx-=f*(d=s[v]),c.vy-=l*d,a.vx+=f*(d=1-d),a.vy+=l*d}function e(){if(a){var n,e,l=a.length,h=t.length,p=se(a,f);for(n=0,c=new Array(l);n<h;++n)(e=t[n]).index=n,"object"!=typeof e.source&&(e.source=Ce(p,e.source)),"object"!=typeof e.target&&(e.target=Ce(p,e.target)),c[e.source.index]=(c[e.source.index]||0)+1,c[e.target.index]=(c[e.target.index]||0)+1;for(n=0,s=new Array(h);n<h;++n)e=t[n],s[n]=c[e.source.index]/(c[e.source.index]+c[e.target.index]);o=new Array(h),r(),u=new Array(h),i()}}function r(){if(a)for(var n=0,e=t.length;n<e;++n)o[n]=+l(t[n],n,t)}function i(){if(a)for(var n=0,e=t.length;n<e;++n)u[n]=+h(t[n],n,t)}var o,u,a,c,s,f=Ae,l=function(t){return 1/Math.min(c[t.source.index],c[t.target.index])},h=ye(30),p=1;return null==t&&(t=[]),n.initialize=function(t){a=t,e()},n.links=function(r){return arguments.length?(t=r,e(),n):t},n.id=function(t){return arguments.length?(f=t,n):f},n.iterations=function(t){return arguments.length?(p=+t,n):p},n.strength=function(t){return arguments.length?(l="function"==typeof t?t:ye(+t),r(),n):l},n.distance=function(t){return arguments.length?(h="function"==typeof t?t:ye(+t),i(),n):h},n},t.forceManyBody=function(){function t(t){var n,a=i.length,c=Te(i,ze,Pe).visitAfter(e);for(u=t,n=0;n<a;++n)o=i[n],c.visit(r)}function n(){if(i){var t,n,e=i.length;for(a=new Array(e),t=0;t<e;++t)n=i[t],a[n.index]=+c(n,t,i)}}function e(t){var n,e,r,i,o,u=0,c=0;if(t.length){for(r=i=o=0;o<4;++o)(n=t[o])&&(e=Math.abs(n.value))&&(u+=n.value,c+=e,r+=e*n.x,i+=e*n.y);t.x=r/c,t.y=i/c}else{(n=t).x=n.data.x,n.y=n.data.y;do{u+=a[n.data.index]}while(n=n.next)}t.value=u}function r(t,n,e,r){if(!t.value)return!0;var i=t.x-o.x,c=t.y-o.y,h=r-n,p=i*i+c*c;if(h*h/l<p)return p<f&&(0===i&&(i=me(),p+=i*i),0===c&&(c=me(),p+=c*c),p<s&&(p=Math.sqrt(s*p)),o.vx+=i*t.value*u/p,o.vy+=c*t.value*u/p),!0;if(!(t.length||p>=f)){(t.data!==o||t.next)&&(0===i&&(i=me(),p+=i*i),0===c&&(c=me(),p+=c*c),p<s&&(p=Math.sqrt(s*p)));do{t.data!==o&&(h=a[t.data.index]*u/p,o.vx+=i*h,o.vy+=c*h)}while(t=t.next)}}var i,o,u,a,c=ye(-30),s=1,f=1/0,l=.81;return t.initialize=function(t){i=t,n()},t.strength=function(e){return arguments.length?(c="function"==typeof e?e:ye(+e),n(),t):c},t.distanceMin=function(n){return arguments.length?(s=n*n,t):Math.sqrt(s)},t.distanceMax=function(n){return arguments.length?(f=n*n,t):Math.sqrt(f)},t.theta=function(n){return arguments.length?(l=n*n,t):Math.sqrt(l)},t},t.forceRadial=function(t,n,e){function r(t){for(var r=0,i=o.length;r<i;++r){var c=o[r],s=c.x-n||1e-6,f=c.y-e||1e-6,l=Math.sqrt(s*s+f*f),h=(a[r]-l)*u[r]*t/l;c.vx+=s*h,c.vy+=f*h}}function i(){if(o){var n,e=o.length;for(u=new Array(e),a=new Array(e),n=0;n<e;++n)a[n]=+t(o[n],n,o),u[n]=isNaN(a[n])?0:+c(o[n],n,o)}}var o,u,a,c=ye(.1);return"function"!=typeof t&&(t=ye(+t)),null==n&&(n=0),null==e&&(e=0),r.initialize=function(t){o=t,i()},r.strength=function(t){return arguments.length?(c="function"==typeof t?t:ye(+t),i(),r):c},r.radius=function(n){return arguments.length?(t="function"==typeof n?n:ye(+n),i(),r):t},r.x=function(t){return arguments.length?(n=+t,r):n},r.y=function(t){return arguments.length?(e=+t,r):e},r},t.forceSimulation=function(t){function n(){e(),p.call("tick",o),u<a&&(h.stop(),p.call("end",o))}function e(){var n,e,r=t.length;for(u+=(s-u)*c,l.each(function(t){t(u)}),n=0;n<r;++n)null==(e=t[n]).fx?e.x+=e.vx*=f:(e.x=e.fx,e.vx=0),null==e.fy?e.y+=e.vy*=f:(e.y=e.fy,e.vy=0)}function r(){for(var n,e=0,r=t.length;e<r;++e){if(n=t[e],n.index=e,isNaN(n.x)||isNaN(n.y)){var i=Fh*Math.sqrt(e),o=e*Ih;n.x=i*Math.cos(o),n.y=i*Math.sin(o)}(isNaN(n.vx)||isNaN(n.vy))&&(n.vx=n.vy=0)}}function i(n){return n.initialize&&n.initialize(t),n}var o,u=1,a=.001,c=1-Math.pow(a,1/300),s=0,f=.6,l=se(),h=wn(n),p=N("tick","end");return null==t&&(t=[]),r(),o={tick:e,restart:function(){return h.restart(n),o},stop:function(){return h.stop(),o},nodes:function(n){return arguments.length?(t=n,r(),l.each(i),o):t},alpha:function(t){return arguments.length?(u=+t,o):u},alphaMin:function(t){return arguments.length?(a=+t,o):a},alphaDecay:function(t){return arguments.length?(c=+t,o):+c},alphaTarget:function(t){return arguments.length?(s=+t,o):s},velocityDecay:function(t){return arguments.length?(f=1-t,o):1-f},force:function(t,n){return arguments.length>1?(null==n?l.remove(t):l.set(t,i(n)),o):l.get(t)},find:function(n,e,r){var i,o,u,a,c,s=0,f=t.length;for(null==r?r=1/0:r*=r,s=0;s<f;++s)(u=(i=n-(a=t[s]).x)*i+(o=e-a.y)*o)<r&&(c=a,r=u);return c},on:function(t,n){return arguments.length>1?(p.on(t,n),o):p.on(t)}}},t.forceX=function(t){function n(t){for(var n,e=0,u=r.length;e<u;++e)(n=r[e]).vx+=(o[e]-n.x)*i[e]*t}function e(){if(r){var n,e=r.length;for(i=new Array(e),o=new Array(e),n=0;n<e;++n)i[n]=isNaN(o[n]=+t(r[n],n,r))?0:+u(r[n],n,r)}}var r,i,o,u=ye(.1);return"function"!=typeof t&&(t=ye(null==t?0:+t)),n.initialize=function(t){r=t,e()},n.strength=function(t){return arguments.length?(u="function"==typeof t?t:ye(+t),e(),n):u},n.x=function(r){return arguments.length?(t="function"==typeof r?r:ye(+r),e(),n):t},n},t.forceY=function(t){function n(t){for(var n,e=0,u=r.length;e<u;++e)(n=r[e]).vy+=(o[e]-n.y)*i[e]*t}function e(){if(r){var n,e=r.length;for(i=new Array(e),o=new Array(e),n=0;n<e;++n)i[n]=isNaN(o[n]=+t(r[n],n,r))?0:+u(r[n],n,r)}}var r,i,o,u=ye(.1);return"function"!=typeof t&&(t=ye(null==t?0:+t)),n.initialize=function(t){r=t,e()},n.strength=function(t){return arguments.length?(u="function"==typeof t?t:ye(+t),e(),n):u},n.y=function(r){return arguments.length?(t="function"==typeof r?r:ye(+r),e(),n):t},n},t.formatDefaultLocale=Ie,t.formatLocale=Fe,t.formatSpecifier=De,t.precisionFixed=Ye,t.precisionPrefix=Be,t.precisionRound=He,t.geoArea=function(t){return Vp.reset(),tr(t,$p),2*Vp},t.geoBounds=function(t){var n,e,r,i,o,u,a;if(Kh=Jh=-(Gh=Qh=1/0),ip=[],tr(t,Zp),e=ip.length){for(ip.sort(xr),n=1,o=[r=ip[0]];n<e;++n)br(r,(i=ip[n])[0])||br(r,i[1])?(mr(r[0],i[1])>mr(r[0],r[1])&&(r[1]=i[1]),mr(i[0],r[1])>mr(r[0],r[1])&&(r[0]=i[0])):o.push(r=i);for(u=-1/0,n=0,r=o[e=o.length-1];n<=e;r=i,++n)i=o[n],(a=mr(r[1],i[0]))>u&&(u=a,Gh=i[0],Jh=r[1])}return ip=op=null,Gh===1/0||Qh===1/0?[[NaN,NaN],[NaN,NaN]]:[[Gh,Qh],[Jh,Kh]]},t.geoCentroid=function(t){up=ap=cp=sp=fp=lp=hp=pp=dp=vp=gp=0,tr(t,Gp);var n=dp,e=vp,r=gp,i=n*n+e*e+r*r;return i<Tp&&(n=lp,e=hp,r=pp,ap<Mp&&(n=cp,e=sp,r=fp),(i=n*n+e*e+r*r)<Tp)?[NaN,NaN]:[Rp(e,n)*Ap,We(r/Yp(i))*Ap]},t.geoCircle=function(){function t(){var t=r.apply(this,arguments),a=i.apply(this,arguments)*Cp,c=o.apply(this,arguments)*Cp;return n=[],e=qr(-t[0]*Cp,-t[1]*Cp,0).invert,Ir(u,a,c,1),t={type:"Polygon",coordinates:[n]},n=e=null,t}var n,e,r=Pr([0,0]),i=Pr(90),o=Pr(6),u={point:function(t,r){n.push(t=e(t,r)),t[0]*=Ap,t[1]*=Ap}};return t.center=function(n){return arguments.length?(r="function"==typeof n?n:Pr([+n[0],+n[1]]),t):r},t.radius=function(n){return arguments.length?(i="function"==typeof n?n:Pr(+n),t):i},t.precision=function(n){return arguments.length?(o="function"==typeof n?n:Pr(+n),t):o},t},t.geoClipAntimeridian=sd,t.geoClipCircle=Qr,t.geoClipExtent=function(){var t,n,e,r=0,i=0,o=960,u=500;return e={stream:function(e){return t&&n===e?t:t=Jr(r,i,o,u)(n=e)},extent:function(a){return arguments.length?(r=+a[0][0],i=+a[0][1],o=+a[1][0],u=+a[1][1],t=n=null,e):[[r,i],[o,u]]}}},t.geoClipRectangle=Jr,t.geoContains=function(t,n){return(t&&gd.hasOwnProperty(t.type)?gd[t.type]:ii)(t,n)},t.geoDistance=ri,t.geoGraticule=hi,t.geoGraticule10=function(){return hi()()},t.geoInterpolate=function(t,n){var e=t[0]*Cp,r=t[1]*Cp,i=n[0]*Cp,o=n[1]*Cp,u=Lp(r),a=Fp(r),c=Lp(o),s=Fp(o),f=u*Lp(e),l=u*Fp(e),h=c*Lp(i),p=c*Fp(i),d=2*We(Yp(Ze(o-r)+u*c*Ze(i-e))),v=Fp(d),g=d?function(t){var n=Fp(t*=d)/v,e=Fp(d-t)/v,r=e*f+n*h,i=e*l+n*p,o=e*a+n*s;return[Rp(i,r)*Ap,Rp(o,Yp(r*r+i*i))*Ap]}:function(){return[e*Ap,r*Ap]};return g.distance=d,g},t.geoLength=ei,t.geoPath=function(t,n){function e(t){return t&&("function"==typeof o&&i.pointRadius(+o.apply(this,arguments)),tr(t,r(i))),i.result()}var r,i,o=4.5;return e.area=function(t){return tr(t,r(xd)),xd.result()},e.measure=function(t){return tr(t,r(Bd)),Bd.result()},e.bounds=function(t){return tr(t,r(Nd)),Nd.result()},e.centroid=function(t){return tr(t,r(qd)),qd.result()},e.projection=function(n){return arguments.length?(r=null==n?(t=null,pi):(t=n).stream,e):t},e.context=function(t){return arguments.length?(i=null==t?(n=null,new Ci):new Si(n=t),"function"!=typeof o&&i.pointRadius(o),e):n},e.pointRadius=function(t){return arguments.length?(o="function"==typeof t?t:(i.pointRadius(+t),+t),e):o},e.projection(t).context(n)},t.geoAlbers=Xi,t.geoAlbersUsa=function(){function t(t){var n=t[0],e=t[1];return a=null,i.point(n,e),a||(o.point(n,e),a)||(u.point(n,e),a)}function n(){return e=r=null,t}var e,r,i,o,u,a,c=Xi(),s=ji().rotate([154,0]).center([-2,58.5]).parallels([55,65]),f=ji().rotate([157,0]).center([-3,19.9]).parallels([8,18]),l={point:function(t,n){a=[t,n]}};return t.invert=function(t){var n=c.scale(),e=c.translate(),r=(t[0]-e[0])/n,i=(t[1]-e[1])/n;return(i>=.12&&i<.234&&r>=-.425&&r<-.214?s:i>=.166&&i<.234&&r>=-.214&&r<-.115?f:c).invert(t)},t.stream=function(t){return e&&r===t?e:e=function(t){var n=t.length;return{point:function(e,r){for(var i=-1;++i<n;)t[i].point(e,r)},sphere:function(){for(var e=-1;++e<n;)t[e].sphere()},lineStart:function(){for(var e=-1;++e<n;)t[e].lineStart()},lineEnd:function(){for(var e=-1;++e<n;)t[e].lineEnd()},polygonStart:function(){for(var e=-1;++e<n;)t[e].polygonStart()},polygonEnd:function(){for(var e=-1;++e<n;)t[e].polygonEnd()}}}([c.stream(r=t),s.stream(t),f.stream(t)])},t.precision=function(t){return arguments.length?(c.precision(t),s.precision(t),f.precision(t),n()):c.precision()},t.scale=function(n){return arguments.length?(c.scale(n),s.scale(.35*n),f.scale(n),t.translate(c.translate())):c.scale()},t.translate=function(t){if(!arguments.length)return c.translate();var e=c.scale(),r=+t[0],a=+t[1];return i=c.translate(t).clipExtent([[r-.455*e,a-.238*e],[r+.455*e,a+.238*e]]).stream(l),o=s.translate([r-.307*e,a+.201*e]).clipExtent([[r-.425*e+Mp,a+.12*e+Mp],[r-.214*e-Mp,a+.234*e-Mp]]).stream(l),u=f.translate([r-.205*e,a+.212*e]).clipExtent([[r-.214*e+Mp,a+.166*e+Mp],[r-.115*e-Mp,a+.234*e-Mp]]).stream(l),n()},t.fitExtent=function(n,e){return qi(t,n,e)},t.fitSize=function(n,e){return Di(t,n,e)},t.fitWidth=function(n,e){return Ui(t,n,e)},t.fitHeight=function(n,e){return Oi(t,n,e)},t.scale(1070)},t.geoAzimuthalEqualArea=function(){return Ii(Vd).scale(124.75).clipAngle(179.999)},t.geoAzimuthalEqualAreaRaw=Vd,t.geoAzimuthalEquidistant=function(){return Ii($d).scale(79.4188).clipAngle(179.999)},t.geoAzimuthalEquidistantRaw=$d,t.geoConicConformal=function(){return Bi(Qi).scale(109.5).parallels([30,30])},t.geoConicConformalRaw=Qi,t.geoConicEqualArea=ji,t.geoConicEqualAreaRaw=Hi,t.geoConicEquidistant=function(){return Bi(Ki).scale(131.154).center([0,13.9389])},t.geoConicEquidistantRaw=Ki,t.geoEquirectangular=function(){return Ii(Ji).scale(152.63)},t.geoEquirectangularRaw=Ji,t.geoGnomonic=function(){return Ii(to).scale(144.049).clipAngle(60)},t.geoGnomonicRaw=to,t.geoIdentity=function(){function t(){return i=o=null,u}var n,e,r,i,o,u,a=1,c=0,s=0,f=1,l=1,h=pi,p=null,d=pi;return u={stream:function(t){return i&&o===t?i:i=h(d(o=t))},postclip:function(i){return arguments.length?(d=i,p=n=e=r=null,t()):d},clipExtent:function(i){return arguments.length?(d=null==i?(p=n=e=r=null,pi):Jr(p=+i[0][0],n=+i[0][1],e=+i[1][0],r=+i[1][1]),t()):null==p?null:[[p,n],[e,r]]},scale:function(n){return arguments.length?(h=no((a=+n)*f,a*l,c,s),t()):a},translate:function(n){return arguments.length?(h=no(a*f,a*l,c=+n[0],s=+n[1]),t()):[c,s]},reflectX:function(n){return arguments.length?(h=no(a*(f=n?-1:1),a*l,c,s),t()):f<0},reflectY:function(n){return arguments.length?(h=no(a*f,a*(l=n?-1:1),c,s),t()):l<0},fitExtent:function(t,n){return qi(u,t,n)},fitSize:function(t,n){return Di(u,t,n)},fitWidth:function(t,n){return Ui(u,t,n)},fitHeight:function(t,n){return Oi(u,t,n)}}},t.geoProjection=Ii,t.geoProjectionMutator=Yi,t.geoMercator=function(){return Zi(Wi).scale(961/Ep)},t.geoMercatorRaw=Wi,t.geoNaturalEarth1=function(){return Ii(eo).scale(175.295)},t.geoNaturalEarth1Raw=eo,t.geoOrthographic=function(){return Ii(ro).scale(249.5).clipAngle(90+Mp)},t.geoOrthographicRaw=ro,t.geoStereographic=function(){return Ii(io).scale(250).clipAngle(142)},t.geoStereographicRaw=io,t.geoTransverseMercator=function(){var t=Zi(oo),n=t.center,e=t.rotate;return t.center=function(t){return arguments.length?n([-t[1],t[0]]):(t=n(),[t[1],-t[0]])},t.rotate=function(t){return arguments.length?e([t[0],t[1],t.length>2?t[2]+90:90]):(t=e(),[t[0],t[1],t[2]-90])},e([0,0,90]).scale(159.155)},t.geoTransverseMercatorRaw=oo,t.geoRotation=Fr,t.geoStream=tr,t.geoTransform=function(t){return{stream:Pi(t)}},t.cluster=function(){function t(t){var o,u=0;t.eachAfter(function(t){var e=t.children;e?(t.x=function(t){return t.reduce(ao,0)/t.length}(e),t.y=function(t){return 1+t.reduce(co,0)}(e)):(t.x=o?u+=n(t,o):0,t.y=0,o=t)});var a=function(t){for(var n;n=t.children;)t=n[0];return t}(t),c=function(t){for(var n;n=t.children;)t=n[n.length-1];return t}(t),s=a.x-n(a,c)/2,f=c.x+n(c,a)/2;return t.eachAfter(i?function(n){n.x=(n.x-t.x)*e,n.y=(t.y-n.y)*r}:function(n){n.x=(n.x-s)/(f-s)*e,n.y=(1-(t.y?n.y/t.y:1))*r})}var n=uo,e=1,r=1,i=!1;return t.separation=function(e){return arguments.length?(n=e,t):n},t.size=function(n){return arguments.length?(i=!1,e=+n[0],r=+n[1],t):i?null:[e,r]},t.nodeSize=function(n){return arguments.length?(i=!0,e=+n[0],r=+n[1],t):i?[e,r]:null},t},t.hierarchy=fo,t.pack=function(){function t(t){return t.x=e/2,t.y=r/2,n?t.eachBefore(zo(n)).eachAfter(Po(i,.5)).eachBefore(Ro(1)):t.eachBefore(zo(Co)).eachAfter(Po(Eo,1)).eachAfter(Po(i,t.r/Math.min(e,r))).eachBefore(Ro(Math.min(e,r)/(2*t.r))),t}var n=null,e=1,r=1,i=Eo;return t.radius=function(e){return arguments.length?(n=function(t){return null==t?null:So(t)}(e),t):n},t.size=function(n){return arguments.length?(e=+n[0],r=+n[1],t):[e,r]},t.padding=function(n){return arguments.length?(i="function"==typeof n?n:Ao(+n),t):i},t},t.packSiblings=function(t){return ko(t),t},t.packEnclose=go,t.partition=function(){function t(t){var o=t.height+1;return t.x0=t.y0=r,t.x1=n,t.y1=e/o,t.eachBefore(function(t,n){return function(e){e.children&&qo(e,e.x0,t*(e.depth+1)/n,e.x1,t*(e.depth+2)/n);var i=e.x0,o=e.y0,u=e.x1-r,a=e.y1-r;u<i&&(i=u=(i+u)/2),a<o&&(o=a=(o+a)/2),e.x0=i,e.y0=o,e.x1=u,e.y1=a}}(e,o)),i&&t.eachBefore(Lo),t}var n=1,e=1,r=0,i=!1;return t.round=function(n){return arguments.length?(i=!!n,t):i},t.size=function(r){return arguments.length?(n=+r[0],e=+r[1],t):[n,e]},t.padding=function(n){return arguments.length?(r=+n,t):r},t},t.stratify=function(){function t(t){var r,i,o,u,a,c,s,f=t.length,l=new Array(f),h={};for(i=0;i<f;++i)r=t[i],a=l[i]=new vo(r),null!=(c=n(r,i,t))&&(c+="")&&(h[s=Zd+(a.id=c)]=s in h?Qd:a);for(i=0;i<f;++i)if(a=l[i],null!=(c=e(t[i],i,t))&&(c+="")){if(!(u=h[Zd+c]))throw new Error("missing: "+c);if(u===Qd)throw new Error("ambiguous: "+c);u.children?u.children.push(a):u.children=[a],a.parent=u}else{if(o)throw new Error("multiple roots");o=a}if(!o)throw new Error("no root");if(o.parent=Gd,o.eachBefore(function(t){t.depth=t.parent.depth+1,--f}).eachBefore(po),o.parent=null,f>0)throw new Error("cycle");return o}var n=Do,e=Uo;return t.id=function(e){return arguments.length?(n=So(e),t):n},t.parentId=function(n){return arguments.length?(e=So(n),t):e},t},t.tree=function(){function t(t){var c=function(t){for(var n,e,r,i,o,u=new Ho(t,0),a=[u];n=a.pop();)if(r=n._.children)for(n.children=new Array(o=r.length),i=o-1;i>=0;--i)a.push(e=n.children[i]=new Ho(r[i],i)),e.parent=n;return(u.parent=new Ho(null,0)).children=[u],u}(t);if(c.eachAfter(n),c.parent.m=-c.z,c.eachBefore(e),a)t.eachBefore(r);else{var s=t,f=t,l=t;t.eachBefore(function(t){t.x<s.x&&(s=t),t.x>f.x&&(f=t),t.depth>l.depth&&(l=t)});var h=s===f?1:i(s,f)/2,p=h-s.x,d=o/(f.x+h+p),v=u/(l.depth||1);t.eachBefore(function(t){t.x=(t.x+p)*d,t.y=t.depth*v})}return t}function n(t){var n=t.children,e=t.parent.children,r=t.i?e[t.i-1]:null;if(n){(function(t){for(var n,e=0,r=0,i=t.children,o=i.length;--o>=0;)(n=i[o]).z+=e,n.m+=e,e+=n.s+(r+=n.c)})(t);var o=(n[0].z+n[n.length-1].z)/2;r?(t.z=r.z+i(t._,r._),t.m=t.z-o):t.z=o}else r&&(t.z=r.z+i(t._,r._));t.parent.A=function(t,n,e){if(n){for(var r,o=t,u=t,a=n,c=o.parent.children[0],s=o.m,f=u.m,l=a.m,h=c.m;a=Io(a),o=Fo(o),a&&o;)c=Fo(c),(u=Io(u)).a=t,(r=a.z+l-o.z-s+i(a._,o._))>0&&(Yo(Bo(a,t,e),t,r),s+=r,f+=r),l+=a.m,s+=o.m,h+=c.m,f+=u.m;a&&!Io(u)&&(u.t=a,u.m+=l-f),o&&!Fo(c)&&(c.t=o,c.m+=s-h,e=t)}return e}(t,r,t.parent.A||e[0])}function e(t){t._.x=t.z+t.parent.m,t.m+=t.parent.m}function r(t){t.x*=o,t.y=t.depth*u}var i=Oo,o=1,u=1,a=null;return t.separation=function(n){return arguments.length?(i=n,t):i},t.size=function(n){return arguments.length?(a=!1,o=+n[0],u=+n[1],t):a?null:[o,u]},t.nodeSize=function(n){return arguments.length?(a=!0,o=+n[0],u=+n[1],t):a?[o,u]:null},t},t.treemap=function(){function t(t){return t.x0=t.y0=0,t.x1=i,t.y1=o,t.eachBefore(n),u=[0],r&&t.eachBefore(Lo),t}function n(t){var n=u[t.depth],r=t.x0+n,i=t.y0+n,o=t.x1-n,h=t.y1-n;o<r&&(r=o=(r+o)/2),h<i&&(i=h=(i+h)/2),t.x0=r,t.y0=i,t.x1=o,t.y1=h,t.children&&(n=u[t.depth+1]=a(t)/2,r+=l(t)-n,i+=c(t)-n,o-=s(t)-n,h-=f(t)-n,o<r&&(r=o=(r+o)/2),h<i&&(i=h=(i+h)/2),e(t,r,i,o,h))}var e=Kd,r=!1,i=1,o=1,u=[0],a=Eo,c=Eo,s=Eo,f=Eo,l=Eo;return t.round=function(n){return arguments.length?(r=!!n,t):r},t.size=function(n){return arguments.length?(i=+n[0],o=+n[1],t):[i,o]},t.tile=function(n){return arguments.length?(e=So(n),t):e},t.padding=function(n){return arguments.length?t.paddingInner(n).paddingOuter(n):t.paddingInner()},t.paddingInner=function(n){return arguments.length?(a="function"==typeof n?n:Ao(+n),t):a},t.paddingOuter=function(n){return arguments.length?t.paddingTop(n).paddingRight(n).paddingBottom(n).paddingLeft(n):t.paddingTop()},t.paddingTop=function(n){return arguments.length?(c="function"==typeof n?n:Ao(+n),t):c},t.paddingRight=function(n){return arguments.length?(s="function"==typeof n?n:Ao(+n),t):s},t.paddingBottom=function(n){return arguments.length?(f="function"==typeof n?n:Ao(+n),t):f},t.paddingLeft=function(n){return arguments.length?(l="function"==typeof n?n:Ao(+n),t):l},t},t.treemapBinary=function(t,n,e,r,i){function o(t,n,e,r,i,u,a){if(t>=n-1){var s=c[t];return s.x0=r,s.y0=i,s.x1=u,void(s.y1=a)}for(var l=f[t],h=e/2+l,p=t+1,d=n-1;p<d;){var v=p+d>>>1;f[v]<h?p=v+1:d=v}h-f[p-1]<f[p]-h&&t+1<p&&--p;var g=f[p]-l,_=e-g;if(u-r>a-i){var y=(r*_+u*g)/e;o(t,p,g,r,i,y,a),o(p,n,_,y,i,u,a)}else{var m=(i*_+a*g)/e;o(t,p,g,r,i,u,m),o(p,n,_,r,m,u,a)}}var u,a,c=t.children,s=c.length,f=new Array(s+1);for(f[0]=a=u=0;u<s;++u)f[u+1]=a+=c[u].value;o(0,s,t.value,n,e,r,i)},t.treemapDice=qo,t.treemapSlice=jo,t.treemapSliceDice=function(t,n,e,r,i){(1&t.depth?jo:qo)(t,n,e,r,i)},t.treemapSquarify=Kd,t.treemapResquarify=tv,t.interpolate=fn,t.interpolateArray=on,t.interpolateBasis=Gt,t.interpolateBasisClosed=Qt,t.interpolateDate=un,t.interpolateNumber=an,t.interpolateObject=cn,t.interpolateRound=ln,t.interpolateString=sn,t.interpolateTransformCss=Gf,t.interpolateTransformSvg=Qf,t.interpolateZoom=vn,t.interpolateRgb=Hf,t.interpolateRgbBasis=jf,t.interpolateRgbBasisClosed=Xf,t.interpolateHsl=el,t.interpolateHslLong=rl,t.interpolateLab=function(t,n){var e=en((t=Ft(t)).l,(n=Ft(n)).l),r=en(t.a,n.a),i=en(t.b,n.b),o=en(t.opacity,n.opacity);return function(n){return t.l=e(n),t.a=r(n),t.b=i(n),t.opacity=o(n),t+""}},t.interpolateHcl=il,t.interpolateHclLong=ol,t.interpolateCubehelix=ul,t.interpolateCubehelixLong=al,t.quantize=function(t,n){for(var e=new Array(n),r=0;r<n;++r)e[r]=t(r/(n-1));return e},t.path=ee,t.polygonArea=function(t){for(var n,e=-1,r=t.length,i=t[r-1],o=0;++e<r;)n=i,i=t[e],o+=n[1]*i[0]-n[0]*i[1];return o/2},t.polygonCentroid=function(t){for(var n,e,r=-1,i=t.length,o=0,u=0,a=t[i-1],c=0;++r<i;)n=a,a=t[r],c+=e=n[0]*a[1]-a[0]*n[1],o+=(n[0]+a[0])*e,u+=(n[1]+a[1])*e;return c*=3,[o/c,u/c]},t.polygonHull=function(t){if((e=t.length)<3)return null;var n,e,r=new Array(e),i=new Array(e);for(n=0;n<e;++n)r[n]=[+t[n][0],+t[n][1],n];for(r.sort($o),n=0;n<e;++n)i[n]=[r[n][0],-r[n][1]];var o=Wo(r),u=Wo(i),a=u[0]===o[0],c=u[u.length-1]===o[o.length-1],s=[];for(n=o.length-1;n>=0;--n)s.push(t[r[o[n]][2]]);for(n=+a;n<u.length-c;++n)s.push(t[r[u[n]][2]]);return s},t.polygonContains=function(t,n){for(var e,r,i=t.length,o=t[i-1],u=n[0],a=n[1],c=o[0],s=o[1],f=!1,l=0;l<i;++l)e=(o=t[l])[0],(r=o[1])>a!=s>a&&u<(c-e)*(a-r)/(s-r)+e&&(f=!f),c=e,s=r;return f},t.polygonLength=function(t){for(var n,e,r=-1,i=t.length,o=t[i-1],u=o[0],a=o[1],c=0;++r<i;)n=u,e=a,n-=u=(o=t[r])[0],e-=a=o[1],c+=Math.sqrt(n*n+e*e);return c},t.quadtree=Te,t.queue=Ko,t.randomUniform=rv,t.randomNormal=iv,t.randomLogNormal=ov,t.randomBates=av,t.randomIrwinHall=uv,t.randomExponential=cv,t.request=nu,t.html=sv,t.json=fv,t.text=lv,t.xml=hv,t.csv=pv,t.tsv=dv,t.scaleBand=ou,t.scalePoint=function(){return uu(ou().paddingInner(1))},t.scaleIdentity=gu,t.scaleLinear=vu,t.scaleLog=Tu,t.scaleOrdinal=iu,t.scaleImplicit=yv,t.scalePow=ku,t.scaleSqrt=function(){return ku().exponent(.5)},t.scaleQuantile=Su,t.scaleQuantize=Eu,t.scaleThreshold=Au,t.scaleTime=function(){return Va(Gv,Wv,Lv,Pv,Cv,Ev,kv,wv,t.timeFormat).domain([new Date(2e3,0,1),new Date(2e3,0,2)])},t.scaleUtc=function(){return Va(xg,yg,ig,eg,tg,Jv,kv,wv,t.utcFormat).domain([Date.UTC(2e3,0,1),Date.UTC(2e3,0,2)])},t.schemeCategory10=Ug,t.schemeCategory20b=Og,t.schemeCategory20c=Fg,t.schemeCategory20=Ig,t.interpolateCubehelixDefault=Yg,t.interpolateRainbow=function(t){(t<0||t>1)&&(t-=Math.floor(t));var n=Math.abs(t-.5);return jg.h=360*t-100,jg.s=1.5-1.5*n,jg.l=.8-.9*n,jg+""},t.interpolateWarm=Bg,t.interpolateCool=Hg,t.interpolateViridis=Xg,t.interpolateMagma=Vg,t.interpolateInferno=$g,t.interpolatePlasma=Wg,t.scaleSequential=Za,t.create=function(t){return ct(A(t).call(document.documentElement))},t.creator=A,t.local=st,t.matcher=of,t.mouse=pt,t.namespace=E,t.namespaces=tf,t.clientPoint=ht,t.select=ct,t.selectAll=function(t){return"string"==typeof t?new ut([document.querySelectorAll(t)],[document.documentElement]):new ut([null==t?[]:t],cf)},t.selection=at,t.selector=z,t.selectorAll=R,t.style=I,t.touch=dt,t.touches=function(t,n){null==n&&(n=lt().touches);for(var e=0,r=n?n.length:0,i=new Array(r);e<r;++e)i[e]=ht(t,n[e]);return i},t.window=F,t.customEvent=it,t.arc=function(){function t(){var t,s,f=+n.apply(this,arguments),l=+e.apply(this,arguments),h=o.apply(this,arguments)-i_,p=u.apply(this,arguments)-i_,d=Zg(p-h),v=p>h;if(c||(c=t=ee()),l<f&&(s=l,l=f,f=s),l>e_)if(d>o_-e_)c.moveTo(l*Qg(h),l*t_(h)),c.arc(0,0,l,h,p,!v),f>e_&&(c.moveTo(f*Qg(p),f*t_(p)),c.arc(0,0,f,p,h,v));else{var g,_,y=h,m=p,x=h,b=p,w=d,M=d,T=a.apply(this,arguments)/2,N=T>e_&&(i?+i.apply(this,arguments):n_(f*f+l*l)),k=Kg(Zg(l-f)/2,+r.apply(this,arguments)),S=k,E=k;if(N>e_){var A=Qa(N/f*t_(T)),C=Qa(N/l*t_(T));(w-=2*A)>e_?(A*=v?1:-1,x+=A,b-=A):(w=0,x=b=(h+p)/2),(M-=2*C)>e_?(C*=v?1:-1,y+=C,m-=C):(M=0,y=m=(h+p)/2)}var z=l*Qg(y),P=l*t_(y),R=f*Qg(b),L=f*t_(b);if(k>e_){var q=l*Qg(m),D=l*t_(m),U=f*Qg(x),O=f*t_(x);if(d<r_){var F=w>e_?function(t,n,e,r,i,o,u,a){var c=e-t,s=r-n,f=u-i,l=a-o,h=(f*(n-o)-l*(t-i))/(l*c-f*s);return[t+h*c,n+h*s]}(z,P,U,O,q,D,R,L):[R,L],I=z-F[0],Y=P-F[1],B=q-F[0],H=D-F[1],j=1/t_(function(t){return t>1?0:t<-1?r_:Math.acos(t)}((I*B+Y*H)/(n_(I*I+Y*Y)*n_(B*B+H*H)))/2),X=n_(F[0]*F[0]+F[1]*F[1]);S=Kg(k,(f-X)/(j-1)),E=Kg(k,(l-X)/(j+1))}}M>e_?E>e_?(g=rc(U,O,z,P,l,E,v),_=rc(q,D,R,L,l,E,v),c.moveTo(g.cx+g.x01,g.cy+g.y01),E<k?c.arc(g.cx,g.cy,E,Gg(g.y01,g.x01),Gg(_.y01,_.x01),!v):(c.arc(g.cx,g.cy,E,Gg(g.y01,g.x01),Gg(g.y11,g.x11),!v),c.arc(0,0,l,Gg(g.cy+g.y11,g.cx+g.x11),Gg(_.cy+_.y11,_.cx+_.x11),!v),c.arc(_.cx,_.cy,E,Gg(_.y11,_.x11),Gg(_.y01,_.x01),!v))):(c.moveTo(z,P),c.arc(0,0,l,y,m,!v)):c.moveTo(z,P),f>e_&&w>e_?S>e_?(g=rc(R,L,q,D,f,-S,v),_=rc(z,P,U,O,f,-S,v),c.lineTo(g.cx+g.x01,g.cy+g.y01),S<k?c.arc(g.cx,g.cy,S,Gg(g.y01,g.x01),Gg(_.y01,_.x01),!v):(c.arc(g.cx,g.cy,S,Gg(g.y01,g.x01),Gg(g.y11,g.x11),!v),c.arc(0,0,f,Gg(g.cy+g.y11,g.cx+g.x11),Gg(_.cy+_.y11,_.cx+_.x11),v),c.arc(_.cx,_.cy,S,Gg(_.y11,_.x11),Gg(_.y01,_.x01),!v))):c.arc(0,0,f,b,x,v):c.lineTo(R,L)}else c.moveTo(0,0);if(c.closePath(),t)return c=null,t+""||null}var n=Ja,e=Ka,r=Ga(0),i=null,o=tc,u=nc,a=ec,c=null;return t.centroid=function(){var t=(+n.apply(this,arguments)+ +e.apply(this,arguments))/2,r=(+o.apply(this,arguments)+ +u.apply(this,arguments))/2-r_/2;return[Qg(r)*t,t_(r)*t]},t.innerRadius=function(e){return arguments.length?(n="function"==typeof e?e:Ga(+e),t):n},t.outerRadius=function(n){return arguments.length?(e="function"==typeof n?n:Ga(+n),t):e},t.cornerRadius=function(n){return arguments.length?(r="function"==typeof n?n:Ga(+n),t):r},t.padRadius=function(n){return arguments.length?(i=null==n?null:"function"==typeof n?n:Ga(+n),t):i},t.startAngle=function(n){return arguments.length?(o="function"==typeof n?n:Ga(+n),t):o},t.endAngle=function(n){return arguments.length?(u="function"==typeof n?n:Ga(+n),t):u},t.padAngle=function(n){return arguments.length?(a="function"==typeof n?n:Ga(+n),t):a},t.context=function(n){return arguments.length?(c=null==n?null:n,t):c},t},t.area=sc,t.line=cc,t.pie=function(){function t(t){var a,c,s,f,l,h=t.length,p=0,d=new Array(h),v=new Array(h),g=+i.apply(this,arguments),_=Math.min(o_,Math.max(-o_,o.apply(this,arguments)-g)),y=Math.min(Math.abs(_)/h,u.apply(this,arguments)),m=y*(_<0?-1:1);for(a=0;a<h;++a)(l=v[d[a]=a]=+n(t[a],a,t))>0&&(p+=l);for(null!=e?d.sort(function(t,n){return e(v[t],v[n])}):null!=r&&d.sort(function(n,e){return r(t[n],t[e])}),a=0,s=p?(_-h*m)/p:0;a<h;++a,g=f)c=d[a],f=g+((l=v[c])>0?l*s:0)+m,v[c]={data:t[c],index:a,value:l,startAngle:g,endAngle:f,padAngle:y};return v}var n=lc,e=fc,r=null,i=Ga(0),o=Ga(o_),u=Ga(0);return t.value=function(e){return arguments.length?(n="function"==typeof e?e:Ga(+e),t):n},t.sortValues=function(n){return arguments.length?(e=n,r=null,t):e},t.sort=function(n){return arguments.length?(r=n,e=null,t):r},t.startAngle=function(n){return arguments.length?(i="function"==typeof n?n:Ga(+n),t):i},t.endAngle=function(n){return arguments.length?(o="function"==typeof n?n:Ga(+n),t):o},t.padAngle=function(n){return arguments.length?(u="function"==typeof n?n:Ga(+n),t):u},t},t.areaRadial=gc,t.radialArea=gc,t.lineRadial=vc,t.radialLine=vc,t.pointRadial=_c,t.linkHorizontal=function(){return xc(bc)},t.linkVertical=function(){return xc(wc)},t.linkRadial=function(){var t=xc(Mc);return t.angle=t.x,delete t.x,t.radius=t.y,delete t.y,t},t.symbol=function(){function t(){var t;if(r||(r=t=ee()),n.apply(this,arguments).draw(r,+e.apply(this,arguments)),t)return r=null,t+""||null}var n=Ga(c_),e=Ga(64),r=null;return t.type=function(e){return arguments.length?(n="function"==typeof e?e:Ga(e),t):n},t.size=function(n){return arguments.length?(e="function"==typeof n?n:Ga(+n),t):e},t.context=function(n){return arguments.length?(r=null==n?null:n,t):r},t},t.symbols=T_,t.symbolCircle=c_,t.symbolCross=s_,t.symbolDiamond=h_,t.symbolSquare=__,t.symbolStar=g_,t.symbolTriangle=m_,t.symbolWye=M_,t.curveBasisClosed=function(t){return new Sc(t)},t.curveBasisOpen=function(t){return new Ec(t)},t.curveBasis=function(t){return new kc(t)},t.curveBundle=N_,t.curveCardinalClosed=S_,t.curveCardinalOpen=E_,t.curveCardinal=k_,t.curveCatmullRomClosed=C_,t.curveCatmullRomOpen=z_,t.curveCatmullRom=A_,t.curveLinearClosed=function(t){return new Oc(t)},t.curveLinear=oc,t.curveMonotoneX=function(t){return new Hc(t)},t.curveMonotoneY=function(t){return new jc(t)},t.curveNatural=function(t){return new Vc(t)},t.curveStep=function(t){return new Wc(t,.5)},t.curveStepAfter=function(t){return new Wc(t,1)},t.curveStepBefore=function(t){return new Wc(t,0)},t.stack=function(){function t(t){var o,u,a=n.apply(this,arguments),c=t.length,s=a.length,f=new Array(s);for(o=0;o<s;++o){for(var l,h=a[o],p=f[o]=new Array(c),d=0;d<c;++d)p[d]=l=[0,+i(t[d],h,d,t)],l.data=t[d];p.key=h}for(o=0,u=e(f);o<s;++o)f[u[o]].index=o;return r(f,u),f}var n=Ga([]),e=Gc,r=Zc,i=Qc;return t.keys=function(e){return arguments.length?(n="function"==typeof e?e:Ga(a_.call(e)),t):n},t.value=function(n){return arguments.length?(i="function"==typeof n?n:Ga(+n),t):i},t.order=function(n){return arguments.length?(e=null==n?Gc:"function"==typeof n?n:Ga(a_.call(n)),t):e},t.offset=function(n){return arguments.length?(r=null==n?Zc:n,t):r},t},t.stackOffsetExpand=function(t,n){if((r=t.length)>0){for(var e,r,i,o=0,u=t[0].length;o<u;++o){for(i=e=0;e<r;++e)i+=t[e][o][1]||0;if(i)for(e=0;e<r;++e)t[e][o][1]/=i}Zc(t,n)}},t.stackOffsetDiverging=function(t,n){if((a=t.length)>1)for(var e,r,i,o,u,a,c=0,s=t[n[0]].length;c<s;++c)for(o=u=0,e=0;e<a;++e)(i=(r=t[n[e]][c])[1]-r[0])>=0?(r[0]=o,r[1]=o+=i):i<0?(r[1]=u,r[0]=u+=i):r[0]=o},t.stackOffsetNone=Zc,t.stackOffsetSilhouette=function(t,n){if((e=t.length)>0){for(var e,r=0,i=t[n[0]],o=i.length;r<o;++r){for(var u=0,a=0;u<e;++u)a+=t[u][r][1]||0;i[r][1]+=i[r][0]=-a/2}Zc(t,n)}},t.stackOffsetWiggle=function(t,n){if((i=t.length)>0&&(r=(e=t[n[0]]).length)>0){for(var e,r,i,o=0,u=1;u<r;++u){for(var a=0,c=0,s=0;a<i;++a){for(var f=t[n[a]],l=f[u][1]||0,h=(l-(f[u-1][1]||0))/2,p=0;p<a;++p){var d=t[n[p]];h+=(d[u][1]||0)-(d[u-1][1]||0)}c+=l,s+=h*l}e[u-1][1]+=e[u-1][0]=o,c&&(o-=s/c)}e[u-1][1]+=e[u-1][0]=o,Zc(t,n)}},t.stackOrderAscending=Jc,t.stackOrderDescending=function(t){return Jc(t).reverse()},t.stackOrderInsideOut=function(t){var n,e,r=t.length,i=t.map(Kc),o=Gc(t).sort(function(t,n){return i[n]-i[t]}),u=0,a=0,c=[],s=[];for(n=0;n<r;++n)e=o[n],u<a?(u+=i[e],c.push(e)):(a+=i[e],s.push(e));return s.reverse().concat(c)},t.stackOrderNone=Gc,t.stackOrderReverse=function(t){return Gc(t).reverse()},t.timeInterval=Cu,t.timeMillisecond=wv,t.timeMilliseconds=Mv,t.utcMillisecond=wv,t.utcMilliseconds=Mv,t.timeSecond=kv,t.timeSeconds=Sv,t.utcSecond=kv,t.utcSeconds=Sv,t.timeMinute=Ev,t.timeMinutes=Av,t.timeHour=Cv,t.timeHours=zv,t.timeDay=Pv,t.timeDays=Rv,t.timeWeek=Lv,t.timeWeeks=Yv,t.timeSunday=Lv,t.timeSundays=Yv,t.timeMonday=qv,t.timeMondays=Bv,t.timeTuesday=Dv,t.timeTuesdays=Hv,t.timeWednesday=Uv,t.timeWednesdays=jv,t.timeThursday=Ov,t.timeThursdays=Xv,t.timeFriday=Fv,t.timeFridays=Vv,t.timeSaturday=Iv,t.timeSaturdays=$v,t.timeMonth=Wv,t.timeMonths=Zv,t.timeYear=Gv,t.timeYears=Qv,t.utcMinute=Jv,t.utcMinutes=Kv,t.utcHour=tg,t.utcHours=ng,t.utcDay=eg,t.utcDays=rg,t.utcWeek=ig,t.utcWeeks=lg,t.utcSunday=ig,t.utcSundays=lg,t.utcMonday=og,t.utcMondays=hg,t.utcTuesday=ug,t.utcTuesdays=pg,t.utcWednesday=ag,t.utcWednesdays=dg,t.utcThursday=cg,t.utcThursdays=vg,t.utcFriday=sg,t.utcFridays=gg,t.utcSaturday=fg,t.utcSaturdays=_g,t.utcMonth=yg,t.utcMonths=mg,t.utcYear=xg,t.utcYears=wg,t.timeFormatDefaultLocale=Ha,t.timeFormatLocale=Du,t.isoFormat=Eg,t.isoParse=Ag,t.now=mn,t.timer=wn,t.timerFlush=Mn,t.timeout=Sn,t.interval=function(t,n,e){var r=new bn,i=n;return null==n?(r.restart(t,n,e),r):(n=+n,e=null==e?mn():+e,r.restart(function o(u){u+=i,r.restart(o,i+=n,e),t(u)},n,e),r)},t.transition=Dn,t.active=function(t,n){var e,r,i=t.__transition;if(i){n=null==n?null:n+"";for(r in i)if((e=i[r]).state>xl&&e.name===n)return new qn([[t]],Jl,n,+r)}return null},t.interrupt=Pn,t.voronoi=function(){function t(t){return new Ns(t.map(function(r,i){var o=[Math.round(n(r,i,t)/F_)*F_,Math.round(e(r,i,t)/F_)*F_];return o.index=i,o.data=r,o}),r)}var n=ns,e=es,r=null;return t.polygons=function(n){return t(n).polygons()},t.links=function(n){return t(n).links()},t.triangles=function(n){return t(n).triangles()},t.x=function(e){return arguments.length?(n="function"==typeof e?e:ts(+e),t):n},t.y=function(n){return arguments.length?(e="function"==typeof n?n:ts(+n),t):e},t.extent=function(n){return arguments.length?(r=null==n?null:[[+n[0][0],+n[0][1]],[+n[1][0],+n[1][1]]],t):r&&[[r[0][0],r[0][1]],[r[1][0],r[1][1]]]},t.size=function(n){return arguments.length?(r=null==n?null:[[0,0],[+n[0],+n[1]]],t):r&&[r[1][0]-r[0][0],r[1][1]-r[0][1]]},t},t.zoom=function(){function n(t){t.property("__zoom",Rs).on("wheel.zoom",c).on("mousedown.zoom",s).on("dblclick.zoom",f).filter(x).on("touchstart.zoom",l).on("touchmove.zoom",h).on("touchend.zoom touchcancel.zoom",p).style("touch-action","none").style("-webkit-tap-highlight-color","rgba(0,0,0,0)")}function e(t,n){return(n=Math.max(b[0],Math.min(b[1],n)))===t.k?t:new Ss(n,t.x,t.y)}function r(t,n,e){var r=n[0]-e[0]*t.k,i=n[1]-e[1]*t.k;return r===t.x&&i===t.y?t:new Ss(t.k,r,i)}function i(t){return[(+t[0][0]+ +t[1][0])/2,(+t[0][1]+ +t[1][1])/2]}function o(t,n,e){t.on("start.zoom",function(){u(this,arguments).start()}).on("interrupt.zoom end.zoom",function(){u(this,arguments).end()}).tween("zoom",function(){var t=arguments,r=u(this,t),o=_.apply(this,t),a=e||i(o),c=Math.max(o[1][0]-o[0][0],o[1][1]-o[0][1]),s=this.__zoom,f="function"==typeof n?n.apply(this,t):n,l=T(s.invert(a).concat(c/s.k),f.invert(a).concat(c/f.k));return function(t){if(1===t)t=f;else{var n=l(t),e=c/n[2];t=new Ss(e,a[0]-n[0]*e,a[1]-n[1]*e)}r.zoom(null,t)}})}function u(t,n){for(var e,r=0,i=k.length;r<i;++r)if((e=k[r]).that===t)return e;return new a(t,n)}function a(t,n){this.that=t,this.args=n,this.index=-1,this.active=0,this.extent=_.apply(t,n)}function c(){if(g.apply(this,arguments)){var t=u(this,arguments),n=this.__zoom,i=Math.max(b[0],Math.min(b[1],n.k*Math.pow(2,m.apply(this,arguments)))),o=pt(this);if(t.wheel)t.mouse[0][0]===o[0]&&t.mouse[0][1]===o[1]||(t.mouse[1]=n.invert(t.mouse[0]=o)),clearTimeout(t.wheel);else{if(n.k===i)return;t.mouse=[o,n.invert(o)],Pn(this),t.start()}Cs(),t.wheel=setTimeout(function(){t.wheel=null,t.end()},A),t.zoom("mouse",y(r(e(n,i),t.mouse[0],t.mouse[1]),t.extent,w))}}function s(){if(!v&&g.apply(this,arguments)){var n=u(this,arguments),e=ct(t.event.view).on("mousemove.zoom",function(){if(Cs(),!n.moved){var e=t.event.clientX-o,i=t.event.clientY-a;n.moved=e*e+i*i>C}n.zoom("mouse",y(r(n.that.__zoom,n.mouse[0]=pt(n.that),n.mouse[1]),n.extent,w))},!0).on("mouseup.zoom",function(){e.on("mousemove.zoom mouseup.zoom",null),yt(t.event.view,n.moved),Cs(),n.end()},!0),i=pt(this),o=t.event.clientX,a=t.event.clientY;_t(t.event.view),As(),n.mouse=[i,this.__zoom.invert(i)],Pn(this),n.start()}}function f(){if(g.apply(this,arguments)){var i=this.__zoom,u=pt(this),a=i.invert(u),c=i.k*(t.event.shiftKey?.5:2),s=y(r(e(i,c),u,a),_.apply(this,arguments),w);Cs(),M>0?ct(this).transition().duration(M).call(o,s,u):ct(this).call(n.transform,s)}}function l(){if(g.apply(this,arguments)){var n,e,r,i,o=u(this,arguments),a=t.event.changedTouches,c=a.length;for(As(),e=0;e<c;++e)i=[i=dt(this,a,(r=a[e]).identifier),this.__zoom.invert(i),r.identifier],o.touch0?o.touch1||(o.touch1=i):(o.touch0=i,n=!0);if(d&&(d=clearTimeout(d),!o.touch1))return o.end(),void((i=ct(this).on("dblclick.zoom"))&&i.apply(this,arguments));n&&(d=setTimeout(function(){d=null},E),Pn(this),o.start())}}function h(){var n,i,o,a,c=u(this,arguments),s=t.event.changedTouches,f=s.length;for(Cs(),d&&(d=clearTimeout(d)),n=0;n<f;++n)o=dt(this,s,(i=s[n]).identifier),c.touch0&&c.touch0[2]===i.identifier?c.touch0[0]=o:c.touch1&&c.touch1[2]===i.identifier&&(c.touch1[0]=o);if(i=c.that.__zoom,c.touch1){var l=c.touch0[0],h=c.touch0[1],p=c.touch1[0],v=c.touch1[1],g=(g=p[0]-l[0])*g+(g=p[1]-l[1])*g,_=(_=v[0]-h[0])*_+(_=v[1]-h[1])*_;i=e(i,Math.sqrt(g/_)),o=[(l[0]+p[0])/2,(l[1]+p[1])/2],a=[(h[0]+v[0])/2,(h[1]+v[1])/2]}else{if(!c.touch0)return;o=c.touch0[0],a=c.touch0[1]}c.zoom("touch",y(r(i,o,a),c.extent,w))}function p(){var n,e,r=u(this,arguments),i=t.event.changedTouches,o=i.length;for(As(),v&&clearTimeout(v),v=setTimeout(function(){v=null},E),n=0;n<o;++n)e=i[n],r.touch0&&r.touch0[2]===e.identifier?delete r.touch0:r.touch1&&r.touch1[2]===e.identifier&&delete r.touch1;r.touch1&&!r.touch0&&(r.touch0=r.touch1,delete r.touch1),r.touch0?r.touch0[1]=this.__zoom.invert(r.touch0[0]):r.end()}var d,v,g=zs,_=Ps,y=Ds,m=Ls,x=qs,b=[0,1/0],w=[[-1/0,-1/0],[1/0,1/0]],M=250,T=vn,k=[],S=N("start","zoom","end"),E=500,A=150,C=0;return n.transform=function(t,n){var e=t.selection?t.selection():t;e.property("__zoom",Rs),t!==e?o(t,n):e.interrupt().each(function(){u(this,arguments).start().zoom(null,"function"==typeof n?n.apply(this,arguments):n).end()})},n.scaleBy=function(t,e){n.scaleTo(t,function(){return this.__zoom.k*("function"==typeof e?e.apply(this,arguments):e)})},n.scaleTo=function(t,o){n.transform(t,function(){var t=_.apply(this,arguments),n=this.__zoom,u=i(t),a=n.invert(u),c="function"==typeof o?o.apply(this,arguments):o;return y(r(e(n,c),u,a),t,w)})},n.translateBy=function(t,e,r){n.transform(t,function(){return y(this.__zoom.translate("function"==typeof e?e.apply(this,arguments):e,"function"==typeof r?r.apply(this,arguments):r),_.apply(this,arguments),w)})},n.translateTo=function(t,e,r){n.transform(t,function(){var t=_.apply(this,arguments),n=this.__zoom,o=i(t);return y(Y_.translate(o[0],o[1]).scale(n.k).translate("function"==typeof e?-e.apply(this,arguments):-e,"function"==typeof r?-r.apply(this,arguments):-r),t,w)})},a.prototype={start:function(){return 1==++this.active&&(this.index=k.push(this)-1,this.emit("start")),this},zoom:function(t,n){return this.mouse&&"mouse"!==t&&(this.mouse[1]=n.invert(this.mouse[0])),this.touch0&&"touch"!==t&&(this.touch0[1]=n.invert(this.touch0[0])),this.touch1&&"touch"!==t&&(this.touch1[1]=n.invert(this.touch1[0])),this.that.__zoom=n,this.emit("zoom"),this},end:function(){return 0==--this.active&&(k.splice(this.index,1),this.index=-1,this.emit("end")),this},emit:function(t){it(new function(t,n,e){this.target=t,this.type=n,this.transform=e}(n,t,this.that.__zoom),S.apply,S,[t,this.that,this.args])}},n.wheelDelta=function(t){return arguments.length?(m="function"==typeof t?t:ks(+t),n):m},n.filter=function(t){return arguments.length?(g="function"==typeof t?t:ks(!!t),n):g},n.touchable=function(t){return arguments.length?(x="function"==typeof t?t:ks(!!t),n):x},n.extent=function(t){return arguments.length?(_="function"==typeof t?t:ks([[+t[0][0],+t[0][1]],[+t[1][0],+t[1][1]]]),n):_},n.scaleExtent=function(t){return arguments.length?(b[0]=+t[0],b[1]=+t[1],n):[b[0],b[1]]},n.translateExtent=function(t){return arguments.length?(w[0][0]=+t[0][0],w[1][0]=+t[1][0],w[0][1]=+t[0][1],w[1][1]=+t[1][1],n):[[w[0][0],w[0][1]],[w[1][0],w[1][1]]]},n.constrain=function(t){return arguments.length?(y=t,n):y},n.duration=function(t){return arguments.length?(M=+t,n):M},n.interpolate=function(t){return arguments.length?(T=t,n):T},n.on=function(){var t=S.on.apply(S,arguments);return t===S?n:t},n.clickDistance=function(t){return arguments.length?(C=(t=+t)*t,n):Math.sqrt(C)},n},t.zoomTransform=Es,t.zoomIdentity=Y_,Object.defineProperty(t,"__esModule",{value:!0})});
```
