## Supplementary material for "ClusTRace, a bioinformatic pipeline for analyzing clusters in virus phylogenies": SFile 1: g3-lollipop.js

```
(function (global, factory) {
	typeof exports === 'object' && typeof module !== 'undefined' ? factory(exports, require('d3')) :
	typeof define === 'function' && define.amd ? define(['exports', 'd3'], factory) :
	(factory((global.g3 = global.g3 || {}),global.d3));
}(this, (function (exports,d3) { 'use strict';

d3 = d3 && d3.hasOwnProperty('default') ? d3['default'] : d3;

function getTextWidth(text, font) {
    var canvas = document.getElementById("canvas") || document.createElement("canvas");
    var context = canvas.getContext("2d");
    context.font = font;
    return context.measureText(text).width;
}

function getUniqueID(left, right) {
    left = left || 1e5;
    right = right || 1e6 - 1;
    return Math.floor(Math.random() * (right - left) + left);
}

var palettes = {
    bottlerocket1: ["#A42820", "#5F5647", "#9B110E", "#3F5151", "#4E2A1E",
        "#550307", "#0C1707"
    ],
    bottlerocket2: ["#FAD510", "#CB2314", "#273046", "#354823", "#1E1E1E"],
    rushmore1: ["#E1BD6D", "#F2300F", "#35274A", "#EABE94", "#0B775E"],
    royal1: ["#899DA4", "#C93312", "#FAEFD1", "#DC863B"],
    royal2: ["#9A8822", "#F5CDB4", "#F8AFA8", "#FDDDA0", "#74A089"],
    zissou1: ["#3B9AB2", "#78B7C5", "#EBCC2A", "#E1AF00", "#F21A00"],
    darjeeling1: ["#FF0000", "#00A08A", "#F2AD00", "#F98400", "#5BBCD6"],
    darjeeling2: ["#ECCBAE", "#046C9A", "#D69C4E", "#ABDDDE", "#000000"],
    chevalier1: ["#446455", "#FDD262", "#D3DDDC", "#C7B19C"],
    fantasticfox1: ["#DD8D29", "#E2D200", "#46ACC8", "#E58601", "#B40F20"],
    moonrise1: ["#F3DF6C", "#CEAB07", "#D5D5D3", "#24281A"],
    moonrise2: ["#798E87", "#C27D38", "#CCC591", "#29211F"],
    moonrise3: ["#85D4E3", "#F4B5BD", "#9C964A", "#CDC08C", "#FAD77B"],
    cavalcanti1: ["#D8B70A", "#02401B", "#A2A475", "#81A88D", "#972D15"],
    grandbudapest1: ["#F1BB7B", "#FD6467", "#5B1A18", "#D67236"],
    grandbudapest2: ["#E6A0C4", "#C6CDF7", "#D8A499", "#7294D4"],
    google16: ["#3366cc", "#dc3912", "#ff9900", "#109618", "#990099",
        "#0099c6", "#dd4477", "#66aa00", "#b82e2e", "#316395",
        "#994499", "#22aa99", "#aaaa11", "#6633cc", "#e67300",
        "#8b0707", "#651067", "#329262", "#5574a6", "#3b3eac"
    ],
    google5: ["#008744", "#0057e7", "#d62d20", "#ffa700", "#ffffff"],
    material1: ["#263238", "#212121", "#3e2723", "#dd2c00", "#ff6d00",
        "#ffab00", "#ffd600", "#aeea00", "#64dd17", "#00c853",
        "#00bfa5", "#00b8d4", "#0091ea", "#2962ff", "#304ffe",
        "#6200ea", "#aa00ff", "#c51162", "#d50000"
    ],
    pie1: ["#a6cee3", "#1f78b4", "#b2df8a", "#33a02c", "#fb9a99"],
    pie2: ["#e41a1c", "#377eb8", "#4daf4a", "#984ea3", "#ff7f00"],
    pie3: ["#495769", "#A0C2BB", "#F4A775", "#F4C667", "#F37361"],
    pie4: ["#FA7921", "#E55934", "#9BC53D", "#FDE74C", "#5BC0EB"],
    pie5: ["#5DA5DA", "#4D4D4D", "#60BD68", "#B2912F", "#B276B2",
        "#F15854", "#FAA43A"
    ],
    pie6: ["#537ea2", "#42a593", "#9f1a1a", "#7c5f95", "#61a070"],
    pie7: ["#bddff9", "#1e72bf", "#ead1ab", "#a2dbc5", "#c7ae7d"],
    breakfast: ["#b6411a", "#eec3d8", "#4182dd", "#ecf0c8", "#2d6328"],
    set1: ["#e41a1c", "#377eb8", "#4daf4a", "#984ea3", "#ff7f00",
        "#ffff33", "#a65628", "#f781bf", "#999999"
    ],
    set2: ["#66c2a5", "#fc8d62", "#8da0cb", "#e78ac3", "#a6d854",
        "#ffd92f", "#e5c494", "#b3b3b3"
    ],
    set3: ["#8dd3c7", "#ffffb3", "#bebada", "#fb8072", "#80b1d3",
        "#fdb462", "#b3de69", "#fccde5", "#d9d9d9", "#bc80bd",
        "#ccebc5", "#ffed6f"
    ],
    category10: ["#1f77b4", "#2ca02c", "#d62728", "#ff7f0e", "#9467bd",
        "#8c564b", "#e377c2", "#7f7f7f", "#bcbd22", "#17becf"
    ],
    pastel1: ["#fbb4ae", "#b3cde3", "#ccebc5", "#decbe4", "#fed9a6",
        "#ffffcc", "#e5d8bd", "#fddaec", "#f2f2f2"
    ],
    pastel2: ["#b3e2cd", "#fdcdac", "#cbd5e8", "#f4cae4", "#e6f5c9",
        "#fff2ae", "#f1e2cc", "#cccccc"
    ],
    accent: ["#7fc97f", "#beaed4", "#fdc086", "#ffff99", "#bf5b17",
        "#386cb0", "#f0027f", "#666666"
    ],
    dark2: ["#1b9e77", "#d95f02", "#7570b3", "#e7298a", "#66a61e",
        "#e6ab02", "#a6761d", "#666666"
    ],
    rainbow: ["#e6261f", "#eb7532", "#f7d038", "#a3e048", "#49da9a", "#34bbe6", "#4355db", "#d23be7"],
    chineseWaterColor: [
        "#832f0e", "#0c0a08", "#594a40", "#8e7967", "#e3c2a0", "#deaa6e", "#81947a"
    ]
};

function defaultPalette() {
    return palettes["google16"];
}

function getPalette(paletteName) {
    return (paletteName in palettes) ? palettes[paletteName] : defaultPalette;
}

function listPalettes() {
    return Object.keys(palettes);
}

function scaleOrdinal(paletteName) {
    return d3.scaleOrdinal(getPalette(paletteName));
}

// ==========================================
// g3.utils.output module 
// ==========================================
//import { saveAs } from 'file-saver';

function output () {
    var defaultFilename = "output";
    
    var output = function(){};
    /*
    function output(nodeId) {
        svgNode = nodeId || svgNode;
    }
    */

    // helper functions
    function _getSVGByID(nodeId) {
        if (nodeId == "svg") {
            return document.querySelector('svg');
        } else {
            return document.getElementsByTagName('svg')[nodeId];
        }
    }

    function _parseFilename(filename, fileExt) {
        filename = filename || defaultFilename;
        return filename + "." + fileExt;
    }

    function _getSVGString(svgNode) {
        svgNode.setAttribute("xmlns", "http://www.w3.org/2000/svg");
        svgNode.setAttribute("xmlns:xlink", "http://www.w3.org/1999/xlink");
        var cssStyvarext = _getCSSStyles(svgNode);

        _appendCSS(cssStyvarext, svgNode);

        var serializer = new XMLSerializer(),
            svgString = serializer.serializeToString(svgNode);
        svgString = svgString.replace(/(\w+)?:?xlink=/g, "xmlns:xlink="); // Fix root xlink without namespace
        svgString = svgString.replace(/NS\d+:href/g, "xlink:href"); // Safari NS namespace fix

        return svgString;

        function _getCSSStyles(parentElement) {
            var selectorTextArr = [];

            // Add Parent element Id and Classes to the list
            selectorTextArr.push("#" + parentElement.id);
            for (let c = 0; c < parentElement.classList.length; c++)
                if (!contains("." + parentElement.classList[c], selectorTextArr))
                    selectorTextArr.push("." + parentElement.classList[c]);

            // Add Children element Ids and Classes to the list
            var nodes = parentElement.getElementsByTagName("*");
            for (let i = 0; i < nodes.length; i++) {
                var id = nodes[i].id;
                if (!contains("#" + id, selectorTextArr))
                    selectorTextArr.push("#" + id);

                var classes = nodes[i].classList;
                for (let c = 0; c < classes.length; c++)
                    if (!contains("." + classes[c], selectorTextArr))
                        selectorTextArr.push("." + classes[c]);
            }

            // Extract CSS Rules
            var extractedCSSText = "";
            for (let i = 0; i < document.styleSheets.length; i++) {
                var s = document.styleSheets[i];

                try {
                    if (!s.cssRules)
                        continue;
                } catch (e) {
                    if (e.name !== "SecurityError")
                        throw e; // for Firefox
                    continue;
                }

                var cssRules = s.cssRules;
                for (var r = 0; r < cssRules.length; r++) {
                    if (contains(cssRules[r].selectorText, selectorTextArr))
                        extractedCSSText += cssRules[r].cssText;
                }
            }

            return extractedCSSText;

            function contains(str, arr) {
                return arr.indexOf(str) === -1 ? false : true;
            }
        }

        function _appendCSS(cssText, element) {
            var styleElement = document.createElement("style");
            styleElement.setAttribute("type", "text/css");
            styleElement.innerHTML = cssText;
            var refNode = element.hasChildNodes() ? element.children[0] : null;
            element.insertBefore(styleElement, refNode);
        }
    }

    output.toSVG = function (filename, nodeId) {
        nodeId = nodeId || "svg";
        var svg = _getSVGByID(nodeId);

        filename = _parseFilename(filename, "svg");

        var svg_string = _getSVGString(svg);
        var blod = new Blob([svg_string], {
            type: "image/svg+xml"
        });

        saveAs(blod, filename);
    };

    output.toPNG = function (filename, nodeId) {
        nodeId = nodeId || "svg";
        var svg = _getSVGByID(nodeId);

        filename = _parseFilename(filename, "png");

        var width = +svg.getAttribute("width"),
            height = +svg.getAttribute("height"),
            svg_string = _getSVGString(svg);

        // Convert SVG string to data URL
        var imgsrc = "data:image/svg+xml;base64," +
            btoa(unescape(encodeURIComponent(svg_string)));
        var canvas = document.createElement("canvas");
        canvas.width = width;
        canvas.height = height;

        var canvas_content = canvas.getContext("2d");
        var image = new Image();
        image.src = imgsrc;
        image.onload = function () {
            canvas_content.clearRect(0, 0, width, height);
            canvas_content.drawImage(image, 0, 0, width, height);

            canvas.toBlob(function (blob) {
                saveAs(blob, filename);
            });
        };
    };

    /* FileSaver.js
     * A saveAs() FileSaver implementation.
     * 1.3.8
     * 2018-03-22 14:03:47
     *
     * By Eli Grey, https://eligrey.com
     * License: MIT
     *   See https://github.com/eligrey/FileSaver.js/blob/master/LICENSE.md
     */

    /*global self */
    /*jslint bitwise: true, indent: 4, laxbreak: true, laxcomma: true, smarttabs: true, plusplus: true */

    /*! @source http://purl.eligrey.com/github/FileSaver.js/blob/master/src/FileSaver.js */

    var saveAs = saveAs || (function (view) {
        if (typeof view === "undefined" || typeof navigator !== "undefined" && /MSIE [1-9]\./.test(navigator.userAgent)) {
            return;
        }
        var
            doc = view.document
            // only get URL when necessary in case Blob.js hasn't overridden it yet
            ,
            get_URL = function () {
                return view.URL || view.webkitURL || view;
            },
            save_link = doc.createElementNS("http://www.w3.org/1999/xhtml", "a"),
            can_use_save_link = "download" in save_link,
            click = function (node) {
                var event = new MouseEvent("click");
                node.dispatchEvent(event);
            },
            is_safari = /constructor/i.test(view.HTMLElement) || view.safari,
            is_chrome_ios = /CriOS\/[\d]+/.test(navigator.userAgent),
            setImmediate = view.setImmediate || view.setTimeout,
            throw_outside = function (ex) {
                setImmediate(function () {
                    throw ex;
                }, 0);
            },
            force_saveable_type = "application/octet-stream"
            // the Blob API is fundamentally broken as there is no "downloadfinished" event to subscribe to
            ,
            arbitrary_revoke_timeout = 1000 * 40 // in ms
            ,
            revoke = function (file) {
                var revoker = function () {
                    if (typeof file === "string") { // file is an object URL
                        get_URL().revokeObjectURL(file);
                    } else { // file is a File
                        file.remove();
                    }
                };
                setTimeout(revoker, arbitrary_revoke_timeout);
            },
            dispatch = function (filesaver, event_types, event) {
                event_types = [].concat(event_types);
                var i = event_types.length;
                while (i--) {
                    var listener = filesaver["on" + event_types[i]];
                    if (typeof listener === "function") {
                        try {
                            listener.call(filesaver, event || filesaver);
                        } catch (ex) {
                            throw_outside(ex);
                        }
                    }
                }
            },
            auto_bom = function (blob) {
                // prepend BOM for UTF-8 XML and text/* types (including HTML)
                // note: your browser will automatically convert UTF-16 U+FEFF to EF BB BF
                if (/^\s*(?:text\/\S*|application\/xml|\S*\/\S*\+xml)\s*;.*charset\s*=\s*utf-8/i.test(blob.type)) {
                    return new Blob([String.fromCharCode(0xFEFF), blob], {
                        type: blob.type
                    });
                }
                return blob;
            },
            FileSaver = function (blob, name, no_auto_bom) {
                if (!no_auto_bom) {
                    blob = auto_bom(blob);
                }
                // First try a.download, then web filesystem, then object URLs
                var
                    filesaver = this,
                    type = blob.type,
                    force = type === force_saveable_type,
                    object_url, dispatch_all = function () {
                        dispatch(filesaver, "writestart progress write writeend".split(" "));
                    }
                    // on any filesys errors revert to saving with object URLs
                    ,
                    fs_error = function () {
                        if ((is_chrome_ios || (force && is_safari)) && view.FileReader) {
                            // Safari doesn't allow downloading of blob urls
                            var reader = new FileReader();
                            reader.onloadend = function () {
                                var url = is_chrome_ios ? reader.result : reader.result.replace(/^data:[^;]*;/, 'data:attachment/file;');
                                var popup = view.open(url, '_blank');
                                if (!popup) view.location.href = url;
                                url = undefined; // release reference before dispatching
                                filesaver.readyState = filesaver.DONE;
                                dispatch_all();
                            };
                            reader.readAsDataURL(blob);
                            filesaver.readyState = filesaver.INIT;
                            return;
                        }
                        // don't create more object URLs than needed
                        if (!object_url) {
                            object_url = get_URL().createObjectURL(blob);
                        }
                        if (force) {
                            view.location.href = object_url;
                        } else {
                            var opened = view.open(object_url, "_blank");
                            if (!opened) {
                                // Apple does not allow window.open, see https://developer.apple.com/library/safari/documentation/Tools/Conceptual/SafariExtensionGuide/WorkingwithWindowsandTabs/WorkingwithWindowsandTabs.html
                                view.location.href = object_url;
                            }
                        }
                        filesaver.readyState = filesaver.DONE;
                        dispatch_all();
                        revoke(object_url);
                    };
                filesaver.readyState = filesaver.INIT;

                if (can_use_save_link) {
                    object_url = get_URL().createObjectURL(blob);
                    setImmediate(function () {
                        save_link.href = object_url;
                        save_link.download = name;
                        click(save_link);
                        dispatch_all();
                        revoke(object_url);
                        filesaver.readyState = filesaver.DONE;
                    }, 0);
                    return;
                }

                fs_error();
            },
            FS_proto = FileSaver.prototype,
            saveAs = function (blob, name, no_auto_bom) {
                return new FileSaver(blob, name || blob.name || "download", no_auto_bom);
            };

        // IE 10+ (native saveAs)
        if (typeof navigator !== "undefined" && navigator.msSaveOrOpenBlob) {
            return function (blob, name, no_auto_bom) {
                name = name || blob.name || "download";

                if (!no_auto_bom) {
                    blob = auto_bom(blob);
                }
                return navigator.msSaveOrOpenBlob(blob, name);
            };
        }

        // todo: detect chrome extensions & packaged apps
        //save_link.target = "_blank";

        FS_proto.abort = function () {};
        FS_proto.readyState = FS_proto.INIT = 0;
        FS_proto.WRITING = 1;
        FS_proto.DONE = 2;

        FS_proto.error =
            FS_proto.onwritestart =
            FS_proto.onprogress =
            FS_proto.onwrite =
            FS_proto.onabort =
            FS_proto.onerror =
            FS_proto.onwriteend =
            null;

        return saveAs;
    }(
        typeof self !== "undefined" && self ||
        typeof window !== "undefined" && window ||
        undefined
    ));

    return output
}

function legend (target, title, series) {
    const Enabled = "enabled", Disabled = "disabled";

    // use setter and getter to control these variables
    // so that "this" keyword can be ignored in legend object
    var interactive = true,
        margin = { left: 10, right: 0, top: 5, bottom: 5, },
        dispatch = d3.dispatch(
            'legendClick',
            'legendDblclick',
            'legendMouseover',
            'legendMouseout');

    title = title || false;
    series = series || [];

    var svg, height = 0;

    var labelFont = "normal 12px sans-serif", labelColor = "black",
        titleFont = "bold 12px sans-serif", titleColor = "black";

    var legend = {
        set target(_) { target = _; }, get target() { return target; },
        set title(_) { title = _; }, get title() { return title; },
        set series(_) { series = _; }, get series() { return series; },
        set margin(_) { margin = _; }, get margin() { return margin; },
        set dispatch(_) { dispatch = _; }, get dispatch() { return dispatch; },
        set interactive(_) { interactive = _; }, get interactive() { return interactive; },
        set labelFont(_) { labelFont = _; }, get labelFont() { return labelFont; },
        set labelColor(_) { labelColor = _; }, get labelColor() { return labelColor; },
        set titleFont(_) { titleFont = _; }, get titleFont() { return titleFont; },
        set titleColor(_) { titleColor = _; }, get titleColor() { return titleColor; },
        get height() { return height; },
        addSeries: function (datum) {
            // key : legend name
            // value: legend related information
            // (text: label, fill: color, class: class_name controlled)
            if (!(datum.key in Object.keys(series))) {
                series.push(datum);
            }
        },
        getLegendStatus: function () {
            return series.map(function (d) {
                return {
                    key: d.key,
                    status: d._status || true
                };
            });
        },
        destroy: function () {
            svg.select(".g3-legend").selectAll("*").remove();
        },
        draw: function () {
            series.forEach(d => d._status = true);
            //let _counter = series.length;

            svg = d3.select("#"+target);
            let _width = +svg.attr("width"),
                _height = +svg.attr("height"),
                _totalW = _width - (margin.left || 0) - (margin.right || 0),
                _wrap = svg.append("g").attr("class", "g3-legend")
                    .attr("transform", "translate(" + margin.left + "," + (_height + margin.top) + ")"),
                _lineHeight = 16;

            let _titleLen = (title) ? getTextWidth(title, titleFont) : 0;

            let _radioLeft = 12, _radioRight = 5, _radioRadius = 3,
                _radioStrokeWidth = 1, _radioStroke = "grey";

            let _titleInterval = 6,
                _titleWidth = Math.min(_titleLen + _titleInterval, 1 / 3 * _totalW),
                _availableW = _totalW - _titleWidth,
                _radioW = _radioLeft + _radioRight + 2 * _radioRadius;

            let _curPos = { x: 0, y: _lineHeight / 2, },
                _nextPos = { x: 0, y: _lineHeight / 2, };

            // add title
            if (title) {
                _wrap.append("g").attr("class", "g3-legend-title")
                    .append("text")
                    .attr("x", _titleWidth - _titleInterval).attr("y", _lineHeight / 2)
                    .attr("text-anchor", "end")
                    .attr("fill", titleColor)
                    .style("font", titleFont)
                    .attr("dy", "0.35em")
                    .text(title);
            }

            var _updatePosition = function (width) {
                // if longer than availabelW
                width += _radioW;
                if (width > _availableW - _nextPos.x) {
                    // new line
                    _curPos.x = 0, _curPos.y = _nextPos.y + _lineHeight;
                    _nextPos.x = width, _nextPos.y = _curPos.y;
                } else {
                    // same line
                    _curPos.x = _nextPos.x;
                    _nextPos.x += width;
                }
            };

            var _toggleState = function (g) {
                var _color = g.attr("defaultColor");
                var _c = g.select("circle"), _t = g.select("text");

                if (g.attr("state") == Enabled) {
                    g.attr("state", Disabled);

                    _c.attr("fill", "transparent")
                        .attr("stroke", _color);
                    _t.style("opacity", 0.5);
                } else {
                    g.attr("state", Enabled);

                    _c.attr("fill", _color)
                        .attr("stroke", _radioStroke);
                    _t.style("opacity", 1);
                }
            };

            var _legends = _wrap.append("g")
                .attr("class", "g3-legend-item-container")
                .attr("transform", "translate(" + _titleWidth + ", 0)");

            var _addOneLegend = function (d) {
                let _g = d3.select(this);
                let _text = d.value.label || d.key;
                _updatePosition(getTextWidth(_text, labelFont));

                let _x = _curPos.x + _radioLeft + _radioRadius, _y = _curPos.y;

                // add legend circle
                _g.attr("state", Enabled)
                    .attr("defaultColor", d.value.fill);

                _g.append("circle")
                    .attr("cx", _x).attr("cy", _y)
                    .attr("r", _radioRadius)
                    .attr("fill", d.value.fill || "transparent")
                    .attr("stroke-width", _radioStrokeWidth)
                    .attr("stroke", _radioStroke);

                _x += (_radioRadius + _radioRight);

                // add text
                _g.append("text")
                    .attr("x", _x).attr("y", _y)
                    .attr("text-anchor", "start")
                    .attr("fill", labelColor)
                    .style("font", labelFont)
                    .attr("dy", "0.35em")
                    .text(_text);

                _g.on("click", function () {
                    if (interactive) {
                        _toggleState(_g);
                        dispatch.call("legendClick", this, d.key, _g.attr("state") == Enabled);
                        d._status = (_g.attr("state") == Enabled);
                    }
                });
            };

            _legends.selectAll(".g3-legend-item")
                .data(series).enter()
                .append("g").attr("class", "g3 g3-legend-item")
                .style("cursor", interactive ? "pointer" : "default")
                .each(_addOneLegend);

            height = margin.top + _curPos.y + _lineHeight / 2 + margin.bottom;
            +svg.attr("height", height + _height);
        },
    };

    return legend;
}

/**
 * d3.tip
 * Copyright (c) 2013-2017 Justin Palmer
 *
 * Tooltips for d3.js SVG visualizations
 * modified
 */
// Public - constructs a new tooltip
//
// Returns a tip
function tooltip () {
  var direction = d3TipDirection,
    offset = d3TipOffset,
    html = d3TipHTML,
    rootElement = document.body,
    node = initNode(),
    svg = null,
    point = null,
    target = null;

  function tip(vis) {
    svg = selectSVGNode(vis);
    if (!svg) return;
    point = svg.createSVGPoint();
    rootElement.appendChild(node);
  }

  // Public - show the tooltip on the screen
  //
  // Returns a tip
  tip.show = function () {
    var args = Array.prototype.slice.call(arguments);
    if (args[args.length - 1] instanceof SVGElement) target = args.pop();

    var content = html.apply(this, args),
      poffset = offset.apply(this, args),
      dir = direction.apply(this, args),
      nodel = getNodeEl(),
      i = directions.length,
      coords,
      scrollTop = document.documentElement.scrollTop ||
      rootElement.scrollTop,
      scrollLeft = document.documentElement.scrollLeft ||
      rootElement.scrollLeft;

    nodel.html(content)
      .style('opacity', 1).style('pointer-events', 'all');

    while (i--) nodel.classed(directions[i], false);
    coords = directionCallbacks.get(dir).apply(this);
    nodel.classed(dir, true)
      .style('top', (coords.top + poffset[0]) + scrollTop + 'px')
      .style('left', (coords.left + poffset[1]) + scrollLeft + 'px');

    return tip
  };

  // Public - hide the tooltip
  //
  // Returns a tip
  tip.hide = function () {
    var nodel = getNodeEl();
    nodel.style('opacity', 0).style('pointer-events', 'none');
    return tip
  };

  // Public: Proxy attr calls to the d3 tip container.
  // Sets or gets attribute value.
  //
  // n - name of the attribute
  // v - value of the attribute
  //
  // Returns tip or attribute value
  // eslint-disable-next-line no-unused-vars
  tip.attr = function (n, v) {
    if (arguments.length < 2 && typeof n === 'string') {
      return getNodeEl().attr(n)
    }

    var args = Array.prototype.slice.call(arguments);
    d3.selection.prototype.attr.apply(getNodeEl(), args);
    return tip
  };

  // Public: Proxy style calls to the d3 tip container.
  // Sets or gets a style value.
  //
  // n - name of the property
  // v - value of the property
  //
  // Returns tip or style property value
  // eslint-disable-next-line no-unused-vars
  tip.style = function (n, v) {
    if (arguments.length < 2 && typeof n === 'string') {
      return getNodeEl().style(n)
    }

    var args = Array.prototype.slice.call(arguments);
    d3.selection.prototype.style.apply(getNodeEl(), args);
    return tip
  };

  // Public: Set or get the direction of the tooltip
  //
  // v - One of n(north), s(south), e(east), or w(west), nw(northwest),
  //     sw(southwest), ne(northeast) or se(southeast)
  //
  // Returns tip or direction
  tip.direction = function (v) {
    if (!arguments.length) return direction
    direction = v == null ? v : functor(v);

    return tip
  };

  // Public: Sets or gets the offset of the tip
  //
  // v - Array of [x, y] offset
  //
  // Returns offset or
  tip.offset = function (v) {
    if (!arguments.length) return offset
    offset = v == null ? v : functor(v);

    return tip
  };

  // Public: sets or gets the html value of the tooltip
  //
  // v - String value of the tip
  //
  // Returns html value or tip
  tip.html = function (v) {
    if (!arguments.length) return html
    html = v == null ? v : functor(v);

    return tip
  };

  // Public: sets or gets the root element anchor of the tooltip
  //
  // v - root element of the tooltip
  //
  // Returns root node of tip
  tip.rootElement = function (v) {
    if (!arguments.length) return rootElement
    rootElement = v == null ? v : functor(v);

    return tip
  };

  // Public: destroys the tooltip and removes it from the DOM
  //
  // Returns a tip
  tip.destroy = function () {
    if (node) {
      getNodeEl().remove();
      node = null;
    }
    return tip
  };

  function d3TipDirection() {
    return 'n'
  }

  function d3TipOffset() {
    return [0, 0]
  }

  function d3TipHTML() {
    return ' '
  }

  var directionCallbacks = d3.map({
      n: directionNorth,
      s: directionSouth,
      e: directionEast,
      w: directionWest,
      nw: directionNorthWest,
      ne: directionNorthEast,
      sw: directionSouthWest,
      se: directionSouthEast
    }),
    directions = directionCallbacks.keys();

  function directionNorth() {
    var bbox = getScreenBBox(this);
    return {
      top: bbox.n.y - node.offsetHeight,
      left: bbox.n.x - node.offsetWidth / 2
    }
  }

  function directionSouth() {
    var bbox = getScreenBBox(this);
    return {
      top: bbox.s.y,
      left: bbox.s.x - node.offsetWidth / 2
    }
  }

  function directionEast() {
    var bbox = getScreenBBox(this);
    return {
      top: bbox.e.y - node.offsetHeight / 2,
      left: bbox.e.x
    }
  }

  function directionWest() {
    var bbox = getScreenBBox(this);
    return {
      top: bbox.w.y - node.offsetHeight / 2,
      left: bbox.w.x - node.offsetWidth
    }
  }

  function directionNorthWest() {
    var bbox = getScreenBBox(this);
    return {
      top: bbox.nw.y - node.offsetHeight,
      left: bbox.nw.x - node.offsetWidth
    }
  }

  function directionNorthEast() {
    var bbox = getScreenBBox(this);
    return {
      top: bbox.ne.y - node.offsetHeight,
      left: bbox.ne.x
    }
  }

  function directionSouthWest() {
    var bbox = getScreenBBox(this);
    return {
      top: bbox.sw.y,
      left: bbox.sw.x - node.offsetWidth
    }
  }

  function directionSouthEast() {
    var bbox = getScreenBBox(this);
    return {
      top: bbox.se.y,
      left: bbox.se.x
    }
  }

  function initNode() {
    var div = d3.select(document.createElement('div'));
    div
      .style('position', 'absolute')
      .style('top', 0)
      .style('opacity', 0)
      .style('pointer-events', 'none')
      .style('box-sizing', 'border-box');

    return div.node()
  }

  function getNodeEl() {
    if (node == null) {
      node = initNode();
      // re-add node to DOM
      rootElement.appendChild(node);
    }
    return d3.select(node)
  }

  // Private - gets the screen coordinates of a shape
  //
  // Given a shape on the screen, will return an SVGPoint for the directions
  // n(north), s(south), e(east), w(west), ne(northeast), se(southeast),
  // nw(northwest), sw(southwest).
  //
  //    +-+-+
  //    |   |
  //    +   +
  //    |   |
  //    +-+-+
  //
  // Returns an Object {n, s, e, w, nw, sw, ne, se}
  function getScreenBBox(targetShape) {
    var targetel = target || d3.event.target; //targetShape

    while (targetel.getScreenCTM == null && targetel.parentNode != null) {
      targetel = targetel.parentNode;
    }

    var bbox = {},
      matrix = targetel.getScreenCTM(),
      tbbox = targetel.getBBox(),
      width = tbbox.width,
      height = tbbox.height,
      x = tbbox.x,
      y = tbbox.y;

    point.x = x;
    point.y = y;
    bbox.nw = point.matrixTransform(matrix);
    point.x += width;
    bbox.ne = point.matrixTransform(matrix);
    point.y += height;
    bbox.se = point.matrixTransform(matrix);
    point.x -= width;
    bbox.sw = point.matrixTransform(matrix);
    point.y -= height / 2;
    bbox.w = point.matrixTransform(matrix);
    point.x += width;
    bbox.e = point.matrixTransform(matrix);
    point.x -= width / 2;
    point.y -= height / 2;
    bbox.n = point.matrixTransform(matrix);
    point.y += height;
    bbox.s = point.matrixTransform(matrix);

    return bbox
  }

  function selectSVGNode(element) {
    var svgNode = element.node();
    if (!svgNode) {
      return null;
    }
    if (svgNode.tagName.toLowerCase() === 'svg') {
      return svgNode;
    }
    return svgNode.ownerSVGElement;
  }

  // Private - replace D3JS 3.X d3.functor() function
  function functor(v) {
    return typeof v === 'function' ? v : function () {
      return v
    }
  }

  return tip
}

function Lollipop(target, chartType, width) {
    const Add = 1, Remove = -1,
        Prefix = "g3_lollipop",
        Undefined = "__undefined__";

    const ChartTypes = { "circle": 0, "pie": 1 };
    const ChartTypeDefault = "circle";
    const WidthDefault = 800;
    const LollipopTrackHeightDefault = 420, DomainTractHeightDefault = 30;

    const SnvDataDefaultFormat = {
        x: "AA_Position",
        y: "Protein_Change",
        factor: "Mutation_Class",
    };

    const DomainDataDefaultFormat = {
        symbol: "hgnc_symbol",
        name: "protein_name",
        length: "length",
        domainType: "pfam",
        details: {
            start: "pfam_start",
            end: "pfam_end",
            ac: "pfam_ac",
            name: "pfam_id",
        },
    };

    // public parameters
    width = width || WidthDefault;
    chartType = chartType || ChartTypeDefault;

    if (!(chartType in ChartTypes)) {
        chartType = ChartTypeDefault;
    }

    var uniqueID = getUniqueID();
    var options = {
        chartID: Prefix + "_chart_" + uniqueID,
        className: "g3-chart",
        tooltip: true,
        margin: { left: 40, right: 20, top: 15, bottom: 25 },
        background: "transparent",
        transitionTime: 600,
        highlightTextAngle: 90,
        legend: true,
        legendOpt: {
            margin: {},
            interactive: true,
            title: Undefined,
        },
    };

    // lollipop settings
    var lollipopOpt = {
        id: Prefix + "-main-" + uniqueID,
        defsId: Prefix + "-main-defs-" + uniqueID,
        xAxisDefsId: Prefix + "-xAxis-defs-" + uniqueID,
        height: LollipopTrackHeightDefault,
        background: "rgb(233,233,233)",
        lollipopClassName: {
            group: "lollipop",
            line: "lollipopLine",
            pop: "pop",
        },
        lollipopLine: {
            "stroke": "rgb(42,42,42)",
            "stroke-width": 0.5,
        },
        popClassName: {
            pie: "popPie",
            slice: "pieSlice",
            text: "popText",
            circle: "popCircle",
            label: "popLabel",
        },
        // pop circle
        popCircle: {
            "stroke": "wheat",
            "stroke-width": 0.5,
        },
        // text in middle of pops
        popText: {
            fill: "#EEE",
            "font-family": "Sans",
            "font-weight": "normal",
            "text-anchor": "middle",
            dy: "0.35em",
        },
        popLabel: {
            "font-family": "Arial",
            "font-weight": "normal",
            minFontSize: 10,
            fontsizeToRadius: 1.4,
            padding: 5,
        },
        popColorSchemeName: "accent",
        popColorScheme: scaleOrdinal("accent"),
        yPaddingRatio: 1.1,
        popParams: {
            rMin: 2,
            rMax: 12,
            addNumRCutoff: 8,
            yLowerQuantile: .25,
            yUpperQuantile: .99,
        },
        title: {
            id: "g3-chart-title",
            text: "",
            font: "normal 16px Arial",
            color: "#424242",
            alignment: "middle",
            dy: "0.35em",
        },
        axisLabel: {
            font: "normal 12px Arial",
            fill: "#4f4f4f",
            "text-anchor": "middle",
            dy: "-2em",
            alignment: "middle",
        },
        ylab: {
            text: "# of mutations",
            lineColor: "#c4c8ca",
            lineWidth: 1,
            lineStyle: "dash",
        },
    };

    // domain information settings
    var domainOpt = {
        id: Prefix + "-domain-" + uniqueID,
        defsId: Prefix + "-annotation-defs-" + uniqueID,
        height: DomainTractHeightDefault,
        margin: { top: 4, bottom: 0 },
        background: "transparent",
        className: {
            track: "domain_track",
            bar: "domain_bar",
            domain: Prefix + "_domain",
            brush: "domain-x-brush",
            zoom: "main-viz-zoom",
        },
        domain: {
            colorSchemeName: "category10",
            colorScheme: scaleOrdinal("category10"),
            margin: { top: 0, bottom: 0 },
            label: {
                font: "normal 11px Arial",
                color: "#f2f2f2",
            },
        },
        bar: {
            background: "#e5e3e1",
            margin: { top: 2, bottom: 2 }
        },
        brush: {
            enabled: true,
            fill: "#666",
            opacity: 0.2,
            stroke: "#969696",
            strokeWidth: 1,
            handler: "#666"
        },
        zoom: true,
    };

    // data and settings
    var snvData = [], domainData = {};
    var snvDataFormat, domainDataFormat;

    /* private variables */
    // chart settings
    var svg;
    var _viz, _mainViz, _domainViz,
        _lollipops, _popLines, _pops,
        _popYUpper, _popYLower, // lollipop viz components
        _domainRect, _domainText,
        _snvInit = false, // data initiallization
        _yUpperValueArray = [], _yValueMax,
        _xRange, _xScale, _xAxis, _xTicks, _domXAxis,
        _yRange, _yScale, _yAxis, _yValues, _domYAxis,
        _popTooltip = null,
        _chartInit = false,
        _byFactor = false,
        _initStates = {}, _currentStates = {},
        _lollipopLegend;

    var _domainBrush, _xScaleOrig, _domainZoom, _legendHeight;
    var _domainH, _domainW, _mainH, _mainW, _svgH, _svgW;

    var _appendValueToDomain = function (tickValues, value) {
        if (value === undefined || tickValues.length <= 1 || tickValues[tickValues.length - 1] >= value)
            return tickValues;

        var interval = Math.abs(tickValues[1] - tickValues[0]);

        if (Math.abs(value - tickValues[tickValues.length - 1]) > interval / 2) {
            tickValues.push(value);
        } else {
            tickValues[tickValues.length - 1] = value;
        }

        return tickValues;
    };

    // event handlers
    var _getDomainRectWidth = function (d) {
        return _getDomainEnd(d) - _getDomainStart(d);
    };

    var _getDomainStart = function (d) {
        return _xScale(d[domainDataFormat.details.start]);
    };

    var _getDomainEnd = function (d) {
        return _xScale(d[domainDataFormat.details.end]);
    };

    var _updateX = function () {
        _xScale.domain(_xRange);

        if (!domainOpt.brush.enabled){
            _domXAxis.call(_xReAxis);

            // update domains
            _domainRect
                .attr("x", function (d) { return _getDomainStart(d); })
                .attr("width", function (d) { return _getDomainRectWidth(d); });

            _domainText
                .attr("x", function (d) { return (_getDomainStart(d) + _getDomainEnd(d)) / 2; });
        }

        // update lines of lollipops
        _popLines//.transition().duration(_options.transitionTime)
            .attr("x1", function (d) { return _getPopX(d); })
            .attr("x2", function (d) { return _getPopX(d); });

        // update pop arcs
        _pops//.transition().duration(_options.transitionTime)
            .attr("transform", function (d) {
                return "translate(" + _getPopX(d) + "," + _getPopY(d) + ")";
            });
    };

    var _domainBrushMove = function () {
        if (d3.event.sourceEvent && d3.event.sourceEvent.type === "zoom") return;

        let _selection = d3.event.selection || _xScaleOrig.range();
        _xRange = _selection.map(_xScaleOrig.invert);

        if (domainOpt.zoom) {
            _mainViz.select(".main-viz-zoom").call(_domainZoom.transform,
                d3.zoomIdentity
                    .scale(_mainW / (_selection[1] - _selection[0]))
                    .translate(-_selection[0], 0));
        }

        _updateX();
    };

    var _drawDomain = function () {
        // domain protein bar
        let _barH = _domainH - domainOpt.bar.margin.top - domainOpt.bar.margin.bottom;

        _domainViz.append("rect").classed(domainOpt.className.bar, true)
            .attr("x", 0).attr("y", domainOpt.bar.margin.top)
            .attr("width", _domainW).attr("height", _barH)
            .attr("fill", domainOpt.bar.background);

        // draw protein domains
        let _dH = _domainH - domainOpt.domain.margin.top - domainOpt.domain.margin.bottom;

        // draw domains
        let _domainG = _domainViz.append("g").attr("class", domainOpt.className.track)
            .selectAll(domainOpt.className.domain)
            .data(domainData[domainDataFormat.domainType]).enter().append("g")
            .attr("clip-path", "url(#" + domainOpt.defsId + ")")
            .attr("class", domainOpt.className.domain);

        _domainRect = _domainG.append("rect")
            .attr("x", function (d) { return _getDomainStart(d); })
            .attr("y", domainOpt.domain.margin.top)
            .attr("height", _dH)
            .attr("width", function (d) { return _getDomainRectWidth(d); })
            .attr("fill", function (d) { return domainOpt.domain.colorScheme(d[domainDataFormat.details.name]); });

        _domainText = _domainG.append("text")
            .attr("x", function (d) { return (_getDomainStart(d) + _getDomainEnd(d)) / 2; })
            .attr("y", domainOpt.domain.margin.top + _dH / 2)
            .attr("dy", "0.35em")
            .attr("text-anchor", "middle")
            .attr("fill", domainOpt.domain.label.color || "white")
            .style("font", domainOpt.domain.label.font)
            .text(function (d) { return d[domainDataFormat.details.name]; });

        if (domainOpt.brush.enabled) {
            _domainBrush = d3.brushX()
                .extent([[0, 0], [_domainW, _domainH]])
                .on("brush end", _domainBrushMove);
        }

        if (domainOpt.zoom) {
            _domainZoom = d3.zoom()
                .scaleExtent([1, Infinity])
                .translateExtent([[0, 0], [_mainW, _mainH]])
                .extent([[0, 0], [_mainW, _mainH]])
                .on("zoom", _mainVizZoomed);
        }

        if (domainOpt.brush.enabled) {
            var _brush = _domainViz.append("g")
                .attr("class", domainOpt.className.brush)
                /*
                .attr("fill", domainOpt.brush.fill)
                .attr("fill-opacity", domainOpt.brush.opacity)
                .attr("stroke", domainOpt.brush.stroke)
                .attr("stroke-width", domainOpt.brush.strokeWidth)
                */
                .call(_domainBrush);

            _brush.selectAll("rect.selection")
                .attr("fill", domainOpt.brush.fill)
                .attr("fill-opacity", domainOpt.brush.opacity)
                .attr("stroke", domainOpt.brush.stroke)
                .attr("stroke-width", domainOpt.brush.strokeWidth);

            _brush.selectAll("rect.handle")
                .attr("fill", domainOpt.brush.handler);
        }
        if (domainOpt.zoom) {
            _mainViz.append("rect")
                .attr("class", domainOpt.className.zoom)
                .attr("width", _mainW)
                .attr("height", _mainH)
                .attr("fill", "none")
                .attr("cursor", "move")
                .attr("pointer-events", "all")
                .call(_domainZoom);
        }
    };

    var _getPopX = function (_datum) {
        return _xScale(_datum.position);
    };

    var _getPopY = function (_datum) {
        return _yScale(_datum._currentState.count);
    };

    var _getPopR = function (_datum) {
        let count = _datum._currentState.count;

        if (count == 0)
            return 0;

        if (count >= _popYUpper)
            return lollipopOpt.popParams.rMax;
        else if (count <= _popYLower)
            return lollipopOpt.popParams.rMin;
        else
            return ((count - _popYLower) / (_popYUpper - _popYLower) *
                (lollipopOpt.popParams.rMax - lollipopOpt.popParams.rMin) +
                lollipopOpt.popParams.rMin);
    };

    var _popDetailHtml = function (d) {
        let _html = "";

        if (_byFactor) {
            _html += `<div class="note">
            <table class="pure-table pure-table-bordered">
                <caption> Position ${d.position} </caption>
                <thead>
                    <tr>
                        <th scope="col">Class</th>
                        <th scope="col">#</th>
                        <th scope="col">Change</th>
                        <th scope="col">&#37;</th>
                    </tr>
                </thead>
                <tbody>`;

            d._currentState.summary.forEach(function (ent) {
                if (ent.value.count > 0) {
                    _html += `
                    <tr>
                        <th scope="row"> ${ent.key} </th>
                        <td> ${ent.value.count} </td>
                        <td>` + _formatProteinChange(ent.value.byY) + `</td>
                        <td>` + d3.format(".1f")(ent.value.count / d._currentState.count * 100) + `&#37;</td>
                    </tr>`;
                }
            });

            _html += `
                </tbody>
            </table></div>`;
        } else {
            _html += `
            <table class="pure-table pure-table-bordered">
                <caption> Position ${d.position} </caption>
                <thead>
                    <tr>
                        <th scope="col">Change</th>
                        <th scope="col">&#37;</th>
                    </tr>
                </thead>
                <tbody>`;

            d._currentState.summary[0].value.byY.forEach(function (ent) {
                _html += `
                    <tr>
                        <th> ${ent.key} </th>
                        <td>` + d3.format(".1f")(ent.value / d._currentState.count * 100) + `&#37;</td>
                    </tr>`;
            });
            _html += `
                </tbody>
            </table>`;
        }

        return _html;
    };

    var _formatProteinChange = function (arr) {
        var info = "";
        arr.forEach(function (e) {
            info += `${e.key} (n=${e.value})<br>`;
        });
        return info;
    };

    var _popMouseOver = function (d, target) {
        if (options.tooltip) {
            _popTooltip.show(d, target);
        }
    };

    var _popMouseOut = function (d) {
        if (options.tooltip) {
            _popTooltip.hide(d);
        }
    };

    var _getDataDominant = function (d) {
        let _snv, _color, _cur_max = -1, _class;
        d._currentState.summary.forEach(function (_c) {
            if (_c.value.count > 0 && _c.value.count > _cur_max) {
                _snv = _c.value.byY[0].key;
                _class = _c;
                _color = lollipopOpt.popColorScheme(_c.key);
                _cur_max = _c.value.count;
            }
        });

        return { "class": _class, "entry": _snv, "count": _cur_max, "color": _color };
    };

    var _popClick = function (d, parentG) {
        // show the most representative node
        if (d._currentState.showLabel) {
            // hide label
            d._currentState.showLabel = false;
            let _label = parentG.select("." + lollipopOpt.popClassName.label);

            _updateYBahavior(_label.attr("yMaxValue"), Remove);

            // text rotation
            _label.transition().duration(options.transitionTime).attr("transform", "rotate(0)").remove();
        } else {
            d._currentState.showLabel = true;

            // (1) pick the lastest mutation class
            // (2) pick the the most representative snvs in this class
            let _dominant = _getDataDominant(d);

            // get text length
            let _fontSize = Math.max(d._currentState.radius * lollipopOpt.popLabel.fontsizeToRadius, lollipopOpt.popLabel.minFontSize) + "px";
            let _font = lollipopOpt.popLabel["font-weight"] + " " + _fontSize + " " + lollipopOpt.popLabel["font-family"];

            // text length in dimention
            let _totalTxtLen = getTextWidth(_dominant.entry, _font) + lollipopOpt.popLabel.padding + d._currentState.radius;

            let _rotatedTxtLen = _totalTxtLen * Math.sin(options.highlightTextAngle / 180 * Math.PI) + 
                                 d._currentState.radius * Math.cos(options.highlightTextAngle / 180 * Math.PI);

            // note: y axis 0 is at top-left cornor
            let _txtYMax = _mainH * d._currentState.count / (_mainH - _rotatedTxtLen);

            _updateYBahavior(_txtYMax, Add);

            var txtHolder = parentG
                .append("text")
                .classed(lollipopOpt.popClassName.label, true)
                .attr("x", - d._currentState.radius - lollipopOpt.popLabel.padding)
                .attr("y", 0)
                .attr("yMaxValue", _txtYMax)
                .text(_dominant.entry)
                .style("fill", _dominant.color)
                .style("font", _font)
                .attr("dy", ".35em")
                .attr("text-anchor", "end");

            // text rotation
            txtHolder.transition().duration(options.transitionTime).attr("transform", "rotate("+options.highlightTextAngle+")");
        }
    };

    var _addLollipopLegend = function () {
        // register legend
        if (!options.legend) return;

        _lollipopLegend = new legend(options.chartID);

        if ((Object.keys(options.legendOpt.margin)).length == 0) {
            options.legendOpt.margin = {
                left: options.margin.left,
                right: options.margin.right,
                top: 2,
                bottom: 2,
            };
        }

        _lollipopLegend.margin = options.legendOpt.margin;
        _lollipopLegend.interactive = options.legendOpt.interactive;
        _lollipopLegend.title = (options.legendOpt.title === Undefined) ? 
            snvDataFormat.factor : options.legendOpt.title;

        for (let _d in _currentStates) {
            _lollipopLegend.addSeries({
                "key": _d,
                "value": {
                    fill: lollipopOpt.popColorScheme(_d),
                    label: _d + " (" + _currentStates[_d] + ")",
                }
            });
        }

        _lollipopLegend.dispatch.on("legendClick", function (key, selected) {
            !selected ? delete _currentStates[key] : _currentStates[key] = selected;

            _prepareData();
            // calculate y range based on the current yValues
            _calcYRange();

            // update (1) y axis (2) lollipop lines (3) pop group
            _updateY();

            _pops.each(_updatePop);
        });

        _lollipopLegend.draw();
        _legendHeight = _lollipopLegend.height;
    };

    var _pieFun = function () {
        return d3.pie().sort(null);
    };

    var _pieTransition = function (d, popGroup, popR) {
        let _arc = d3.arc().innerRadius(0).outerRadius(popR);
        let _pie = _pieFun().value(function (_d) { return _d.value.count; });

        let _pieSlice = popGroup.select("." + lollipopOpt.popClassName.pie)
            .selectAll("path." + lollipopOpt.popClassName.slice)
            .data(_pie(d._currentState.summary));

        _pieSlice.enter().insert("path");

        _pieSlice.transition().duration(options.transitionTime)
            .attrTween("d", function (_d) {
                this._current = this._current || _d;
                let _interpolate = d3.interpolate(this._current, _d);
                this._current = _interpolate(0);

                return function (t) {
                    return _arc(_interpolate(t));
                }
            });

        _pieSlice.exit().remove();
    };

    var _circleTransition = function (d, popGroup, popR) {
        let _circle = popGroup.select("." + lollipopOpt.popClassName.circle);
        let _dominant = _getDataDominant(d);
        let _fill = _dominant.color || "transparent";

        _circle.transition().duration(options.transitionTime)
            .attr("r", popR).attr("fill", _fill);
    };

    var _updatePop = function (d) {
        let _popGroup = d3.select(this);

        // remove label
        if (d._currentState.showLabel) {
            _popGroup.select("." + lollipopOpt.popClassName.label).remove();
            d._currentState.showLabel = false;
        }

        // update pop
        let _popR = d._currentState.radius = _getPopR(d);

        if(chartType == "pie"){
            _pieTransition(d, _popGroup, _popR);
        }
        
        _circleTransition(d, _popGroup, _popR);

        let _popText = _popGroup.select("." + lollipopOpt.popClassName.text);
        // add text if large enough
        if (_popR >= lollipopOpt.popParams.addNumRCutoff) {
            let _popTextFontSize = _calcPopTextFontSize(_popR, d._currentState.count, lollipopOpt.popText["font-weight"],
                lollipopOpt.popText["font-family"]);
            let _popTextFont = lollipopOpt.popText["font-weight"] + " " + _popTextFontSize + " " + lollipopOpt.popText["font-family"];

            _popText.transition().duration(options.transitionTime)
                .text(d._currentState.count)
                .style("font", _popTextFont);
        } else {
            _popText.text("");
        }
    };

    var _calcPopTextFontSize = function (r, number, fontWeight, fontFamily) {
        let _w = r * 1.75;
        for (let _f = Math.ceil(_w); _f >= 0; _f--) {
            let _textwidth = getTextWidth(number, fontWeight + " " + _f + "px " + fontFamily);
            if (_textwidth <= _w) {
                return _f + "px";
            }
        }
        return 0;
    };

    var _getPopGroupId = function (idx) {
        return Prefix + "-" + uniqueID + "-lollipop-group-" + idx;
    };

    var _drawPie = function (d) {
        let _chartG = d3.select(this);

        let _popR = d._currentState.radius = _getPopR(d);

        // add popCircle
        _chartG.append("circle")
            .attr("class", lollipopOpt.popClassName.circle)
            .attr("cx", 0).attr("cy", 0)
            .attr("r", _popR)
            .attr("stroke", lollipopOpt.popCircle.stroke)
            .attr("stroke-width", lollipopOpt.popCircle["stroke-width"])
            .attr("fill", "none");

        // add pie (no change on axis change)
        let _arc = d3.arc().innerRadius(0).outerRadius(_popR);
        let _pie = _pieFun().value(function (_d) { return _d.value.count; });

        // pie arc
        let _pieSlice = _chartG.append("g").attr("class", lollipopOpt.popClassName.pie);

        // add pie arcs
        _pieSlice.selectAll(lollipopOpt.popClassName.slice)
            .data(_pie(d._currentState.summary))
            .enter().append("path")
            .attr("d", _arc)
            .attr("class", lollipopOpt.popClassName.slice)
            .attr("fill", function (d) {
                return lollipopOpt.popColorScheme(d.data.key);
            });

        _pieSlice.exit().remove();

        // add text if large enough
        let _popText = _chartG.append("text")
            .attr("class", lollipopOpt.popClassName.text)
            .attr("fill", lollipopOpt.popText.fill)
            .style("font-family", lollipopOpt.popText["font-family"])
            .style("font-weight", lollipopOpt.popText["font-weight"])
            .attr("text-anchor", lollipopOpt.popText["text-anchor"])
            .attr("dy", lollipopOpt.popText.dy);

        if (_popR >= lollipopOpt.popParams.addNumRCutoff) {
            let _popTextFontSize = _calcPopTextFontSize(_popR,
                d._currentState.count, lollipopOpt.popText["font-weight"],
                lollipopOpt.popText["font-family"]);
            _popText.transition().duration(options.transitionTime)
                .text(d._currentState.count)
                .style("font-size", _popTextFontSize);
        } else {
            _popText.text("");
        }

        _chartG.on("mouseover", function (d) { _popMouseOver(d); })
            .on("mouseout", function (d) { _popMouseOut(d); })
            .on("click", function (d) { _popClick(d, _chartG); });
    };

    var _drawCircle = function (d) {
        let _chartG = d3.select(this);

        let _popR = d._currentState.radius = _getPopR(d);

        // circle
        let _dominant = _getDataDominant(d);
        // circle
        _chartG.append("circle")
            .attr("class", lollipopOpt.popClassName.circle)
            .attr("cx", 0).attr("cy", 0)
            .attr("r", _popR)
            .attr("stroke", lollipopOpt.popCircle.stroke)
            .attr("stroke-width", lollipopOpt.popCircle["stroke-width"])
            .attr("fill", _dominant.color);

        // add text if large enough
        let _popText = _chartG.append("text")
            .attr("class", lollipopOpt.popClassName.text)
            .style("font-family", lollipopOpt.popText["font-family"])
            .style("font-weight", lollipopOpt.popText["font-weight"])
            .attr("text-anchor", lollipopOpt.popText["text-anchor"])
            .attr("dy", lollipopOpt.popText.dy);

        if (_popR >= lollipopOpt.popParams.addNumRCutoff) {
            let _popTextFontSize = _calcPopTextFontSize(_popR,
                d._currentState.count, lollipopOpt.popText["font-weight"],
                lollipopOpt.popText["font-family"]);
            _popText.transition().duration(options.transitionTime)
                .text(d._currentState.count)
                .attr("font-size", _popTextFontSize);
        } else {
            _popText.text("");
        }

        _chartG.on("mouseover", function (d) { _popMouseOver(d); })
            .on("mouseout", function (d) { _popMouseOut(d); })
            .on("click", function (d) { _popClick(d, _chartG); });
    };

    var _drawLollipops = function () {
        // lollipop groups
        _lollipops = _mainViz.append("g")
            .attr("class", lollipopOpt.lollipopClassName.group);

        // draw popline first
        _popLines = _lollipops.append("g")
            .attr("clip-path", "url(#" + lollipopOpt.defsId + ")")
            .selectAll(lollipopOpt.lollipopClassName.line)
            .data(snvData).enter()
            .append("line")
            .attr("x1", function (d) { return _getPopX(d); })
            .attr("x2", function (d) { return _getPopX(d); })
            .attr("y1", _yScale(0) + domainOpt.margin.top)
            .attr("y2", function (d) { return _getPopY(d); })
            .attr("class", lollipopOpt.lollipopClassName.line)
            .attr("stroke", lollipopOpt.lollipopLine.stroke)
            .attr("stroke-width", lollipopOpt.lollipopLine["stroke-width"]);

        // then draw pops
        _pops = _lollipops.append("g")
            .attr("clip-path", "url(#" + lollipopOpt.defsId + ")")
            .selectAll(lollipopOpt.lollipopClassName.pop)
            .data(snvData).enter()
            .append("g")
            .attr("transform", function (d) { return "translate(" + _getPopX(d) + "," + _getPopY(d) + ")"; })
            .classed(lollipopOpt.lollipopClassName.pop, true)
            .attr("id", function (d, i) { return _getPopGroupId(i); });

        switch (chartType) {
            case "pie":
                _pops.each(_drawPie);
                break;
            default:
                _pops.each(_drawCircle);
        }
        // add title
        let _anchor, _xPos, _yPos;
        if (lollipopOpt.title.text.length > 0) {
            if (lollipopOpt.title.alignment == "start") {
                _anchor = "start"; _xPos = 0;
            } else if (lollipopOpt.title.alignment == "middle") {
                _anchor = "middle"; _xPos = _mainW / 2;
            } else {
                _anchor = "end"; _xPos = _mainW;
            }

            _yPos = -options.margin.top / 2;

            _mainViz.append("text")
                .attr("id", lollipopOpt.title.id)
                .style("font", lollipopOpt.title.font)
                .attr("fill", lollipopOpt.title.color)
                .attr("text-anchor", _anchor)
                .attr("alignment-base", "alphabetic")
                .attr("transform", "translate(" + _xPos + "," + _yPos + ")")
                .attr("dy", lollipopOpt.title.dy)
                .text(lollipopOpt.title.text);
        }
    };

    var _mainVizZoomed = function () {
        if (d3.event.sourceEvent && d3.event.sourceEvent.type === "brush") return;

        let _t = d3.event.transform;
        _xRange = _t.rescaleX(_xScaleOrig).domain();

        if (domainOpt.brush.enabled) {
            _domainViz.select("." + domainOpt.className.brush)
                .call(_domainBrush.move, _xScaleOrig.range().map(_t.invertX, _t));
        }
        _updateX();
    };

    var _calcOptions = function () {
        // viz height
        _svgH = lollipopOpt.height + domainOpt.height;
        _svgW = width;
        _mainH = lollipopOpt.height - options.margin.top - options.margin.bottom;
        _mainW = _svgW - options.margin.left - options.margin.right;
        _domainH = domainOpt.height - domainOpt.margin.top - domainOpt.margin.bottom;
        _domainW = _mainW;
    };

    var _prepareData = function () {
        if (!snvDataFormat.hasOwnProperty('x') || !snvDataFormat.hasOwnProperty('y')) {
            throw "No X or Y columns specified in data set";
        }
        // for the first time, summarize mutation factors and counts
        if (!_snvInit) {
            // => factor (or undefined): count
            _initStates = d3.nest()
                .key(d => d[snvDataFormat.factor])
                .rollup(function (d) { return +d.length; })
                .object(snvData);
            
            _currentStates = JSON.parse(JSON.stringify(_initStates));

            // group by postion, sort
            snvData = d3.nest()
                .key(d => +d[snvDataFormat.x])
                .entries(snvData)
                .sort((a, b) => a.key - b.key);

            // position and total
            snvData.forEach(function (d) {
                d["position"] = +d["key"]; delete d["key"];
                d["total"] = +d.values.length;
            });
        }

        // parse each position
        // use _currentState to record the current information for current
        snvData.forEach(function (d) {
            // group by factor, group by y, sort
            //let _d = d.values.filter(function (d) { return d[snvOpt.factor] in _currentStates; });
            let _factorSummary = d3.nest()
                .key(function (_) { return _[snvDataFormat.factor]; })
                .rollup(function (_) {
                    let _d = _.filter(function (_) { return _[snvDataFormat.factor] in _currentStates; });
                    return {
                        count: +_d.length,
                        byY: d3.nest().key(function (_) { return _[snvDataFormat.y]; })
                            .rollup(function (_) { return +_.length; })
                            .entries(_d).sort(function (a, b) { return b.value - a.value; }),
                    };
                }).entries(d.values)
                .sort(function (a, b) { return a.key > b.key; });
            //.sort(function (a, b) { return b.value.count - a.value.count; });

            if (!_snvInit) {
                d._currentState = {};
                d._currentState.showLabel = false;
            }
            d._currentState.count = d3.sum(_factorSummary.map(function (_) { return _.value.count; }));
            d._currentState.summary = _factorSummary;
        });

        _snvInit = true;

        // calculate counts
        _yValues = snvData
            .map(function (_) { return +_._currentState.count; })
            .filter(function (_) { return _ > 0; })
            .sort(function (a, b) { return a - b; });

        // calculate pop size boundary
        _popYLower = Math.ceil(d3.quantile(_yValues, lollipopOpt.popParams.yLowerQuantile));
        _popYUpper = Math.floor(d3.quantile(_yValues, lollipopOpt.popParams.yUpperQuantile));

        _yUpperValueArray = [];
    };

    var _getYMaxAfterNice = function (yMax) {
//        return d3.scaleLinear().domain([0, yMax]).range([_mainH, 0]).nice().domain()[1];
        return d3.scaleLinear().domain([0, yMax]).nice().domain()[1];
    };

    var _calcYRange = function () {
        if (_yValues.length == 0) {
            _yRange = [0, 1];
        } else {
            _yRange = d3.extent(_yValues);
        }
        // get yRange
        _yRange[0] = Math.min(0, _yRange[0]);
        // add lollipopOpt.popParams.rMax

//        lollipopOpt.popParams.rMax / _mainH
//        _yValueMax = _yRange[1] = _getYMaxAfterNice(_yRange[1] * lollipopOpt.yPaddingRatio);
        _yValueMax = _yRange[1] = _getYMaxAfterNice(
            _yRange[1] * (lollipopOpt.yPaddingRatio + lollipopOpt.popParams.rMax / _mainH)
        );
    };

    var _calcAxis = function () {
        // y
        _calcYRange();
        _yScale = d3.scaleLinear().domain(_yRange).range([_mainH, 0]);
        _yAxis = d3.axisLeft(_yScale).tickSize(-_mainW).ticks(6).tickFormat(d3.format("d"));

        // x
        _xRange = [0, domainData[domainDataFormat.length]];
        _xScale = d3.scaleLinear().domain(_xRange).range([0, _domainW]);
        _xScaleOrig = d3.scaleLinear().domain(_xRange).range([0, _domainW]);
        _xTicks = _appendValueToDomain(_xScale.ticks(), domainData[domainDataFormat.length], "aa");
        _xAxis = d3.axisBottom(_xScale).tickValues(_xTicks)
            .tickFormat(function (_) { return this.parentNode.nextSibling ? _ : _ + " aa"; });
    };

    var _updateYBahavior = function (yValue, behavior) {
        // check behavior
        behavior = behavior || Add; // default is to add
        if (behavior != Add && behavior != Remove) behavior = Add;

        // calculate new yMax
        yValue = _getYMaxAfterNice(yValue * lollipopOpt.yPaddingRatio);
        yValue = Math.max(_yValueMax, yValue);

        // if yValue > yValueMax
        if (behavior === Add) {
            _yUpperValueArray.push(yValue);
            _yUpperValueArray = _yUpperValueArray.sort(function(a, b){return a - b});
        } else {
            let _idx = _yUpperValueArray.indexOf(yValue);
            if (_idx != -1) {
                _yUpperValueArray = _yUpperValueArray.slice(0, _idx).concat(_yUpperValueArray.slice(_idx + 1));
            }
        }

        // if need to
        let _yMax = (_yUpperValueArray.length == 0) ? _yValueMax : _yUpperValueArray[_yUpperValueArray.length - 1];

        if (_yMax != _yRange[1]) {
            _yRange[1] = _yMax;
            _updateY();
        }
    };

    var _updateY = function () {
        _yScale.domain(_yRange);
        _domYAxis.transition().duration(options.transitionTime).call(_yReAxis);

        // update lines of lollipops
        _popLines.transition().duration(options.transitionTime).attr("y2", function (d) { return _getPopY(d); });

        // update pop group
        _pops.transition().duration(options.transitionTime).attr("transform", function (d) {
            return "translate(" + _getPopX(d) + "," + _getPopY(d) + ")";
        });
    };

    var _xReAxis = function (g) {
        var s = g.selection ? g.selection() : g;
        g.call(_xAxis);

        s.select(".domain").remove();
        s.selectAll(".tick line")
            .attr("stroke", "#c4c8ca")
            .attr("stroke-width", 1);
    };

    var _yReAxis = function (g) {
        var s = g.selection ? g.selection() : g;
        g.call(_yAxis);

        s.select(".domain").remove();
        let __tickLine = s.selectAll(".tick line")//.filter(Number)
            .attr("stroke", lollipopOpt.ylab.lineColor)
            .attr("stroke-width", lollipopOpt.ylab.lineWidth);
        if(lollipopOpt.ylab.lineStyle == "dash"){
            __tickLine.attr("stroke-dasharray", "3,3");
        }

        s.selectAll(".tick text").attr("x", -2).attr("dy", 2);
        if (s !== g) g.selectAll(".tick text").attrTween("x", null).attrTween("dy", null);
    };

    var _drawAxis = function () {
        //yRange[1] *= _lollipopOpt.yRatio;
        _calcAxis();

        _domYAxis = _mainViz.append("g").attr("class", "axis axis--y").call(_yReAxis);

        // y label text
        let _anchor, _yPos;
        if (lollipopOpt.axisLabel.alignment == "start") {
            _anchor = "end"; _yPos = 0;
        } else if (lollipopOpt.axisLabel.alignment == "middle") {
            _anchor = "middle"; _yPos = _mainH / 2;
        } else {
            _anchor = "start"; _yPos = _mainH;
        }

        _mainViz.append("text")
            .attr("class", "yaxis axis-label")
            .attr("transform", "translate(0," + _yPos + ")rotate(-90)")
            .style("font", lollipopOpt.axisLabel.font)
            .attr("fill", lollipopOpt.axisLabel.fill)
            .attr("text-anchor", _anchor)
            .attr("alignment-base", "middle")
            .attr("dy", lollipopOpt.axisLabel.dy)
            .text(lollipopOpt.ylab.text);

        // add x axis
        _domXAxis = _mainViz.append("g")
            .attr("clip-path", "url(#" + lollipopOpt.xAxisDefsId + ")")
            .attr("class", "axis axis--x")
            .attr("transform", "translate(0, " + (_mainH + domainOpt.height) + ")")
            .call(_xReAxis);
    };

    var _addBackground = function (g, bgColor, height, width) {
        var bgLayer = g.append("g").attr("id", "background-layer");
        bgLayer.append("rect")
            .attr("x", 0).attr("y", 0)
            .attr("width", width).attr("height", height)
            .attr("fill", bgColor);
    };

    var _initMainViz = function () {
        _addBackground(_mainViz, lollipopOpt.background, _mainH, _mainW);
    };

    var _initDomainViz = function () {
        _addBackground(_domainViz, domainOpt.background, _domainH, _domainW);
    };

    var _initTooltip = function () {
        if (options.tooltip) {
            _popTooltip = tooltip().attr("class", "d3-tip").offset([8, 0])
                .direction(
                    function (d) {
                        if (d.count > _yRange[1] / 2) return "s";
                        else return "n";
                    })
                .offset(function (d) {
                    if (d.count > _yRange[1] / 2) return [16, 0];
                    else return [-12, 0];
                })
                .html(_popDetailHtml);

            _mainViz.call(_popTooltip);
        }
    };

    var _initViz = function () {
        if (!target) {
            target = "g3_" + uniqueID;
            d3.select("body").append("div").attr("id", target);
        }

        svg = d3.select("#" + target).append("svg");
        svg.attr("width", _svgW).attr("height", _svgH)
            .attr("xmlns", "http://www.w3.org/2000/svg")
            .attr("xmlns:xlink", "http://www.w3.org/1999/xlink");

        svg.classed(options.className, true)
            .attr("id", options.chartID)
            .style("background-color", options.background || "transparent");

        // viz region
        _viz = svg.append("g")
            .attr("transform", "translate(" + options.margin.left + "," + options.margin.top + ")");

        // lollipop viz
        _mainViz = _viz.append("g").attr("id", lollipopOpt.id);
        _initMainViz();

        // annotation viz
        _domainViz = _viz.append("g").attr("id", domainOpt.id)
            .attr("transform", "translate(0, " + (_mainH + domainOpt.margin.top) + ")");
        _initDomainViz();

        // tooltips and menu
        _initTooltip();
    };

    var _initBrush = function () {
        _domainViz.select(".domain-x-brush").call(_domainBrush.move, [0, _domainW]);
    };

    var _addDefs = function () {
        // defs
        _mainViz.append("clipPath")
            .attr("id", lollipopOpt.defsId)
            .append("rect")
            .attr("width", _mainW)
            .attr("height", _mainH + domainOpt.margin.top);

        _mainViz.append("clipPath")
            .attr("id", lollipopOpt.xAxisDefsId)
            .append("rect")
            .attr("x", -5)
            .attr("width", _mainW + 20) // TODO
            .attr("height", 30);

        // defs
        _domainViz.append("clipPath")
            .attr("id", domainOpt.defsId)
            .append("rect")
            .attr("width", _domainW)
            .attr("height", _domainH);
    };

    // =====================
    // public functions
    // =====================
    var lollipop = {
    };

    // getter and setter, e.g., lollipop.options.width; lollipop.options.snvData = [];
    lollipop.options = {
        // set target svg node
        set chartTarget(_) { target = _; }, get chartTarget() { return target; },
        // set chart width
        set chartWidth(_) { width = _; }, get chartWidth() { return width; },
        // set chart type (pie or circle)
        set chartType(_) { if (_ && _ in ChartTypes) chartType = _; }, get chartType() { return chartType; },
        // get chart ID
        get chartID() { return options.chartID },
        get chartHeight() { return +svg.attr("height") },

        // chart margin (top / bottom / left / right)
        set chartMargin(_) { options.margin = _; }, get chartMargin() { return options.margin; },
        // chart background
        set chartBackground(_) { options.background = _; }, get chartBackground() { return options.background; },

        // chart animation transition time (ms)
        set transitionTime(_) { options.transitionTime = _; }, get transitionTime() { return options.transitionTime; },

        // highlight text angle
        set highlightTextAngle(_) { options.highlightTextAngle = _; }, get highlightTextAngle() { return options.highlightTextAngle; },

        // if enable tooltip
        set tooltip(_) { options.tooltip = _; }, get tooltip() { return options.tooltip; },

        // ylabel text
        set yAxisLabel(_) { lollipopOpt.ylab.text = _; }, get yAxisLabel() { return lollipopOpt.ylab.text; },
        // y-axis line color
        set yAxisLineColor(_) { lollipopOpt.ylab.lineColor = _; }, get yAxisLineColor() { return lollipopOpt.ylab.lineColor; },
        set yAxisLineWidth(_) { lollipopOpt.ylab.lineWidth = _; }, get yAxisLineWidth() { return lollipopOpt.ylab.lineWidth; },
        set yAxisLineStyle(_) { lollipopOpt.ylab.lineStyle = _; }, get yAxisLineStyle() { return lollipopOpt.ylab.lineStyle; },
        set yMaxRangeRatio(_) { lollipopOpt.yPaddingRatio = _; }, get yMaxRangeRatio() { return lollipopOpt.yPaddingRatio; },

        // axis settings (label font / color / alignment / y-adjustment)
        set axisLabelFont(_) { lollipopOpt.axisLabel.font = _; }, get axisLabelFont() { return lollipopOpt.axisLabel.font; },
        set axisLabelColor(_) { lollipopOpt.axisLabel.fill = _; }, get axisLabelColor() { return lollipopOpt.axisLabel.fill; },
        set axisLabelAlignment(_) { lollipopOpt.axisLabel.alignment = _; }, get axisLabelAlignment() { return lollipopOpt.axisLabel.alignment; },
        set axisLabelDy(_) { lollipopOpt.axisLabel.dy = _; }, get axisLabelDy() { return lollipopOpt.axisLabel.dy; },

        // legend settings (show legend or not / legend margin / interactive legend or not)
        set legend(_) { options.legend = _; }, get legend() { return options.legend; },
        set legendMargin(_) { options.legendOpt.margin = _; }, get legendMargin() { return options.legendOpt.margin; },
        set legendInteractive(_) { options.legendOpt.interactive = _; }, get legendInteractive() { return options.legendOpt.interactive; },
        set legendTitle(_) { options.legendOpt.title = _; }, get legendTitle() { return options.legendOpt.title; },
        get legendHeight() { return _legendHeight; },

        // get lollipopTrack ID
        get lollipopTrackID() { return lollipopOpt.id; },

        // lollipop track height and background
        set lollipopTrackHeight(_) { lollipopOpt.height = _; }, 
        get lollipopTrackHeight() { return lollipopOpt.height; },

        set lollipopTrackBackground(_) { lollipopOpt.background = _; }, 
        get lollipopTrackBackground() { return lollipopOpt.background; },

        // pop circle size (min / max)
        set lollipopPopMinSize(_) { lollipopOpt.popParams.rMin = _; }, 
        get lollipopPopMinSize() { return lollipopOpt.popParams.rMin; },

        set lollipopPopMaxSize(_) { lollipopOpt.popParams.rMax = _; }, 
        get lollipopPopMaxSize() { return lollipopOpt.popParams.rMax; },

        // pop cicle text (radius cutoff to show info, info text color)
        set lollipopPopInfoLimit(_) { lollipopOpt.popParams.addNumRCutoff = _; }, 
        get lollipopPopInfoLimit() { return lollipopOpt.popParams.addNumRCutoff; },

        set lollipopPopInfoColor(_) { lollipopOpt.popText.fill = _; }, 
        get lollipopPopInfoColor() { return lollipopOpt.popText.fill; },

        set lollipopPopInfoDy(_) { lollipopOpt.popText.dy = _; }, 
        get lollipopPopInfoDy() { return lollipopOpt.popText.dy; },

        // lollipop line (color / width)
        set lollipopLineColor(_) { lollipopOpt.lollipopLine.stroke = _; }, 
        get lollipopLineColor() { return lollipopOpt.lollipopLine.stroke; },

        set lollipopLineWidth(_) { lollipopOpt.lollipopLine["stroke-width"] = _; }, 
        get lollipopLineWidth() { return lollipopOpt.lollipopLine["stroke-width"]; },

        // lollipop circle (color / width)
        set lollipopCircleColor(_) { lollipopOpt.popCircle.stroke = _; }, 
        get lollipopCircleColor() { return lollipopOpt.popCircle.stroke; },

        set lollipopCircleWidth(_) { lollipopOpt.popCircle["stroke-width"] = _; }, 
        get lollipopCircleWidth() { return lollipopOpt.popCircle["stroke-width"]; },

        // pop click label (font size ratio to pop size / minimal font size)
        set lollipopLabelRatio(_) { lollipopOpt.popLabel.fontsizeToRadius = _; }, 
        get lollipopLabelRatio() { return lollipopOpt.popLabel.fontsizeToRadius; },
        
        set lollipopLabelMinFontSize(_) { lollipopOpt.popLabel.minFontSize = _; }, 
        get lollipopLabelMinFontSize() { return lollipopOpt.popLabel.minFontSize; },

        // pop color scheme
        set lollipopColorScheme(_) { lollipopOpt.popColorSchemeName = _; lollipopOpt.popColorScheme = scaleOrdinal(_); },
        get lollipopColorScheme() { return lollipopOpt.popColorSchemeName; },

        // title related settings (text / font / color / alignment / y-adjustment)
        // note : font (font-style font-variant font-weight font-size/line-height font-family)
        // i.e., italic small-caps normal 13px sans-serif
        set titleText(_) { lollipopOpt.title.text = _; }, 
        get titleText() { return lollipopOpt.title.text; },

        set titleFont(_) { lollipopOpt.title.font = _; }, 
        get titleFont() { return lollipopOpt.title.font; },

        set titleColor(_) { lollipopOpt.title.color = _; }, 
        get titleColor() { return lollipopOpt.title.color; },

        set titleAlignment(_) { lollipopOpt.title.alignment = _; }, 
        get titleAlignment() { return lollipopOpt.title.alignment; },

        set titleDy(_) { lollipopOpt.title.dy = _; }, 
        get titleDy() { return lollipopOpt.title.dy; },

        get annoID() { return domainOpt.id },

        set annoHeight(_) { domainOpt.height = _; }, 
        get annoHeight() { return domainOpt.height; },

        set annoMargin(_) { domainOpt.margin = _; }, 
        get annoMargin() { return domainOpt.margin; },

        set annoBackground(_) { domainOpt.background = _; }, 
        get annoBackground() { return domainOpt.background; },

        set annoBarFill(_) { domainOpt.bar.background = _; }, 
        get annoBarFill() { return domainOpt.bar.background; },

        set annoBarMargin(_) { domainOpt.bar.margin = _; }, 
        get annoBarMargin() { return domainOpt.bar.margin; },

        set domainColorScheme(_) { domainOpt.domain.colorSchemeName = _; domainOpt.domain.colorScheme = scaleOrdinal(_); }, 
        get domainColorScheme() { return domainOpt.domain.colorSchemeName; },

        set domainMargin(_) { domainOpt.domain.margin = _; }, 
        get domainMargin() { return domainOpt.domain.margin; },

        set domainTextFont(_) { domainOpt.domain.label.font = _; }, 
        get domainTextFont() { return domainOpt.domain.label.font; },

        set domainTextColor(_) { domainOpt.domain.label.color = _; }, 
        get domainTextColor() { return domainOpt.domain.label.color; },

        set brush(_) { domainOpt.brush.enabled = _; }, 
        get brush() { return domainOpt.brush.enabled; },

        set brushBackground(_) { domainOpt.brush.fill = _; }, 
        get brushBackground() { return domainOpt.brush.fill; },

        set brushOpacity(_) { domainOpt.brush.opacity = _; }, 
        get brushOpacity() { return domainOpt.brush.opacity; },

        set brushBorderColor(_) { domainOpt.brush.stroke = _; }, 
        get brushBorderColor() { return domainOpt.brush.stroke; },

        set brushBorderWidth(_) { domainOpt.brush.strokeWdith = _; }, 
        get brushBorderWidth() { return domainOpt.brush.strokeWidth; },

        set brushHandler(_) { domainOpt.brush.handler = _; }, 
        get brushHandler() { return domainOpt.brush.handler; }, 

        set zoom(_) { domainOpt.zoom = _; }, 
        get zoom() { return domainOpt.zoom; },
    };

    lollipop.setOptions = function (options) {
        for (let _key in options) {
            this.options[_key] = options[_key];
        }
    };

    lollipop.destroy = function () {
        svg.selectAll("*").remove();
        svg.remove();

        if (options.tooltip && _chartInit) {
            _popTooltip.destroy();
        }
    };

    lollipop.refresh = function () {
        this.destroy();

        // reset states
        _currentStates = JSON.parse(JSON.stringify(_initStates));
        this.draw();
    };

    lollipop.data = {
        set snvData(_) { snvData = _; }, get snvData() { return snvData; },
        set domainData(_) { domainData = _; }, get domainData() { return domainData; },
    };

    lollipop.format = {
        set snvData(_) { snvDataFormat = _; }, get snvData() { return snvDataFormat; },
        set domainData(_) { domainDataFormat = _; }, get domainData() { return domainDataFormat; },
    };

    lollipop.draw = function() {
        // check data format
        snvDataFormat = snvDataFormat || SnvDataDefaultFormat;
        domainDataFormat = domainDataFormat || DomainDataDefaultFormat;

        if (snvDataFormat.hasOwnProperty('factor')) {
            _byFactor = true;
        } else {
            _byFactor = false;
            lollipop.format.snvData.factor = snvDataFormat.factor = Undefined;
            snvData.forEach(function (d) { d[snvDataFormat.factor] = Undefined; });
        }

        _calcOptions();

        // prepare data
        _prepareData();

        // initalize viz
        _initViz();

        // calculate range, Axis
        _drawAxis();

        // add defs
        _addDefs();

        // add domain panel
        _drawDomain();

        // add lollipops
        _drawLollipops();
        if (options.legend) {
            _addLollipopLegend();
        }

        if (domainOpt.brush.enabled) {
            _initBrush();
        }

        _chartInit = true;
    };

    return lollipop;
}

exports.getTextWidth = getTextWidth;
exports.getUniqueID = getUniqueID;
exports.palettes = palettes;
exports.defaultPalette = defaultPalette;
exports.getPalette = getPalette;
exports.listPalettes = listPalettes;
exports.scaleOrdinal = scaleOrdinal;
exports.output = output;
exports.legend = legend;
exports.tooltip = tooltip;
exports.Lollipop = Lollipop;

Object.defineProperty(exports, '__esModule', { value: true });

})));
```
