## Supplementary material for "ClusTRace, a bioinformatic pipeline for analyzing clusters in virus phylogenies": SFile 1: g3-lollipop.min.js

```
!function(t,e){"object"==typeof exports&&"undefined"!=typeof module?e(exports,require("d3")):"function"==typeof define&&define.amd?define(["exports","d3"],e):e(t.g3=t.g3||{},t.d3)}(this,function(t,e){"use strict";function n(t,e){var n=(document.getElementById("canvas")||document.createElement("canvas")).getContext("2d");return n.font=e,n.measureText(t).width}function r(t,e){return t=t||1e5,e=e||1e6-1,Math.floor(Math.random()*(e-t)+t)}e=e&&e.hasOwnProperty("default")?e.default:e;var a={bottlerocket1:["#A42820","#5F5647","#9B110E","#3F5151","#4E2A1E","#550307","#0C1707"],bottlerocket2:["#FAD510","#CB2314","#273046","#354823","#1E1E1E"],rushmore1:["#E1BD6D","#F2300F","#35274A","#EABE94","#0B775E"],royal1:["#899DA4","#C93312","#FAEFD1","#DC863B"],royal2:["#9A8822","#F5CDB4","#F8AFA8","#FDDDA0","#74A089"],zissou1:["#3B9AB2","#78B7C5","#EBCC2A","#E1AF00","#F21A00"],darjeeling1:["#FF0000","#00A08A","#F2AD00","#F98400","#5BBCD6"],darjeeling2:["#ECCBAE","#046C9A","#D69C4E","#ABDDDE","#000000"],chevalier1:["#446455","#FDD262","#D3DDDC","#C7B19C"],fantasticfox1:["#DD8D29","#E2D200","#46ACC8","#E58601","#B40F20"],moonrise1:["#F3DF6C","#CEAB07","#D5D5D3","#24281A"],moonrise2:["#798E87","#C27D38","#CCC591","#29211F"],moonrise3:["#85D4E3","#F4B5BD","#9C964A","#CDC08C","#FAD77B"],cavalcanti1:["#D8B70A","#02401B","#A2A475","#81A88D","#972D15"],grandbudapest1:["#F1BB7B","#FD6467","#5B1A18","#D67236"],grandbudapest2:["#E6A0C4","#C6CDF7","#D8A499","#7294D4"],google16:["#3366cc","#dc3912","#ff9900","#109618","#990099","#0099c6","#dd4477","#66aa00","#b82e2e","#316395","#994499","#22aa99","#aaaa11","#6633cc","#e67300","#8b0707","#651067","#329262","#5574a6","#3b3eac"],google5:["#008744","#0057e7","#d62d20","#ffa700","#ffffff"],material1:["#263238","#212121","#3e2723","#dd2c00","#ff6d00","#ffab00","#ffd600","#aeea00","#64dd17","#00c853","#00bfa5","#00b8d4","#0091ea","#2962ff","#304ffe","#6200ea","#aa00ff","#c51162","#d50000"],pie1:["#a6cee3","#1f78b4","#b2df8a","#33a02c","#fb9a99"],pie2:["#e41a1c","#377eb8","#4daf4a","#984ea3","#ff7f00"],pie3:["#495769","#A0C2BB","#F4A775","#F4C667","#F37361"],pie4:["#FA7921","#E55934","#9BC53D","#FDE74C","#5BC0EB"],pie5:["#5DA5DA","#4D4D4D","#60BD68","#B2912F","#B276B2","#F15854","#FAA43A"],pie6:["#537ea2","#42a593","#9f1a1a","#7c5f95","#61a070"],pie7:["#bddff9","#1e72bf","#ead1ab","#a2dbc5","#c7ae7d"],breakfast:["#b6411a","#eec3d8","#4182dd","#ecf0c8","#2d6328"],set1:["#e41a1c","#377eb8","#4daf4a","#984ea3","#ff7f00","#ffff33","#a65628","#f781bf","#999999"],set2:["#66c2a5","#fc8d62","#8da0cb","#e78ac3","#a6d854","#ffd92f","#e5c494","#b3b3b3"],set3:["#8dd3c7","#ffffb3","#bebada","#fb8072","#80b1d3","#fdb462","#b3de69","#fccde5","#d9d9d9","#bc80bd","#ccebc5","#ffed6f"],category10:["#1f77b4","#2ca02c","#d62728","#ff7f0e","#9467bd","#8c564b","#e377c2","#7f7f7f","#bcbd22","#17becf"],pastel1:["#fbb4ae","#b3cde3","#ccebc5","#decbe4","#fed9a6","#ffffcc","#e5d8bd","#fddaec","#f2f2f2"],pastel2:["#b3e2cd","#fdcdac","#cbd5e8","#f4cae4","#e6f5c9","#fff2ae","#f1e2cc","#cccccc"],accent:["#7fc97f","#beaed4","#fdc086","#ffff99","#bf5b17","#386cb0","#f0027f","#666666"],dark2:["#1b9e77","#d95f02","#7570b3","#e7298a","#66a61e","#e6ab02","#a6761d","#666666"],rainbow:["#e6261f","#eb7532","#f7d038","#a3e048","#49da9a","#34bbe6","#4355db","#d23be7"],chineseWaterColor:["#832f0e","#0c0a08","#594a40","#8e7967","#e3c2a0","#deaa6e","#81947a"]};function o(){return a.google16}function i(t){return t in a?a[t]:o}function l(t){return e.scaleOrdinal(i(t))}function s(t,r,a){var o=!0,i={left:10,right:0,top:5,bottom:5},l=e.dispatch("legendClick","legendDblclick","legendMouseover","legendMouseout");r=r||!1,a=a||[];var s,c=0,u="normal 12px sans-serif",p="black",d="bold 12px sans-serif",f="black";return{set target(e){t=e},get target(){return t},set title(t){r=t},get title(){return r},set series(t){a=t},get series(){return a},set margin(t){i=t},get margin(){return i},set dispatch(t){l=t},get dispatch(){return l},set interactive(t){o=t},get interactive(){return o},set labelFont(t){u=t},get labelFont(){return u},set labelColor(t){p=t},get labelColor(){return p},set titleFont(t){d=t},get titleFont(){return d},set titleColor(t){f=t},get titleColor(){return f},get height(){return c},addSeries:function(t){t.key in Object.keys(a)||a.push(t)},getLegendStatus:function(){return a.map(function(t){return{key:t.key,status:t._status||!0}})},destroy:function(){s.select(".g3-legend").selectAll("*").remove()},draw:function(){a.forEach(t=>t._status=!0);let g=+(s=e.select("#"+t)).attr("width"),h=+s.attr("height"),m=g-(i.left||0)-(i.right||0),b=s.append("g").attr("class","g3-legend").attr("transform","translate("+i.left+","+(h+i.top)+")"),y=r?n(r,d):0,x=Math.min(y+6,1/3*m),v=m-x,k={x:0,y:8},C={x:0,y:8};r&&b.append("g").attr("class","g3-legend-title").append("text").attr("x",x-6).attr("y",8).attr("text-anchor","end").attr("fill",f).style("font",d).attr("dy","0.35em").text(r);b.append("g").attr("class","g3-legend-item-container").attr("transform","translate("+x+", 0)").selectAll(".g3-legend-item").data(a).enter().append("g").attr("class","g3 g3-legend-item").style("cursor",o?"pointer":"default").each(function(t){let r=e.select(this),a=t.value.label||t.key;var i;i=n(a,u),(i+=23)>v-C.x?(k.x=0,k.y=C.y+16,C.x=i,C.y=k.y):(k.x=C.x,C.x+=i);let s=k.x+12+3,c=k.y;r.attr("state","enabled").attr("defaultColor",t.value.fill),r.append("circle").attr("cx",s).attr("cy",c).attr("r",3).attr("fill",t.value.fill||"transparent").attr("stroke-width",1).attr("stroke","grey"),s+=8,r.append("text").attr("x",s).attr("y",c).attr("text-anchor","start").attr("fill",p).style("font",u).attr("dy","0.35em").text(a),r.on("click",function(){var e,n,a,i;o&&(n=(e=r).attr("defaultColor"),a=e.select("circle"),i=e.select("text"),"enabled"==e.attr("state")?(e.attr("state","disabled"),a.attr("fill","transparent").attr("stroke",n),i.style("opacity",.5)):(e.attr("state","enabled"),a.attr("fill",n).attr("stroke","grey"),i.style("opacity",1)),l.call("legendClick",this,t.key,"enabled"==r.attr("state")),t._status="enabled"==r.attr("state"))})}),c=i.top+k.y+8+i.bottom,s.attr("height",c+h)}}}function c(){var t=function(){return"n"},n=function(){return[0,0]},r=function(){return" "},a=document.body,o=d(),i=null,l=null,s=null;function c(t){(i=function(t){var e=t.node();if(!e)return null;if("svg"===e.tagName.toLowerCase())return e;return e.ownerSVGElement}(t))&&(l=i.createSVGPoint(),a.appendChild(o))}c.show=function(){var e=Array.prototype.slice.call(arguments);e[e.length-1]instanceof SVGElement&&(s=e.pop());var o,i=r.apply(this,e),l=n.apply(this,e),d=t.apply(this,e),g=f(),h=p.length,m=document.documentElement.scrollTop||a.scrollTop,b=document.documentElement.scrollLeft||a.scrollLeft;for(g.html(i).style("opacity",1).style("pointer-events","all");h--;)g.classed(p[h],!1);return o=u.get(d).apply(this),g.classed(d,!0).style("top",o.top+l[0]+m+"px").style("left",o.left+l[1]+b+"px"),c},c.hide=function(){return f().style("opacity",0).style("pointer-events","none"),c},c.attr=function(t,n){if(arguments.length<2&&"string"==typeof t)return f().attr(t);var r=Array.prototype.slice.call(arguments);return e.selection.prototype.attr.apply(f(),r),c},c.style=function(t,n){if(arguments.length<2&&"string"==typeof t)return f().style(t);var r=Array.prototype.slice.call(arguments);return e.selection.prototype.style.apply(f(),r),c},c.direction=function(e){return arguments.length?(t=null==e?e:h(e),c):t},c.offset=function(t){return arguments.length?(n=null==t?t:h(t),c):n},c.html=function(t){return arguments.length?(r=null==t?t:h(t),c):r},c.rootElement=function(t){return arguments.length?(a=null==t?t:h(t),c):a},c.destroy=function(){return o&&(f().remove(),o=null),c};var u=e.map({n:function(){var t=g(this);return{top:t.n.y-o.offsetHeight,left:t.n.x-o.offsetWidth/2}},s:function(){var t=g(this);return{top:t.s.y,left:t.s.x-o.offsetWidth/2}},e:function(){var t=g(this);return{top:t.e.y-o.offsetHeight/2,left:t.e.x}},w:function(){var t=g(this);return{top:t.w.y-o.offsetHeight/2,left:t.w.x-o.offsetWidth}},nw:function(){var t=g(this);return{top:t.nw.y-o.offsetHeight,left:t.nw.x-o.offsetWidth}},ne:function(){var t=g(this);return{top:t.ne.y-o.offsetHeight,left:t.ne.x}},sw:function(){var t=g(this);return{top:t.sw.y,left:t.sw.x-o.offsetWidth}},se:function(){var t=g(this);return{top:t.se.y,left:t.se.x}}}),p=u.keys();function d(){var t=e.select(document.createElement("div"));return t.style("position","absolute").style("top",0).style("opacity",0).style("pointer-events","none").style("box-sizing","border-box"),t.node()}function f(){return null==o&&(o=d(),a.appendChild(o)),e.select(o)}function g(t){for(var n=s||e.event.target;null==n.getScreenCTM&&null!=n.parentNode;)n=n.parentNode;var r={},a=n.getScreenCTM(),o=n.getBBox(),i=o.width,c=o.height,u=o.x,p=o.y;return l.x=u,l.y=p,r.nw=l.matrixTransform(a),l.x+=i,r.ne=l.matrixTransform(a),l.y+=c,r.se=l.matrixTransform(a),l.x-=i,r.sw=l.matrixTransform(a),l.y-=c/2,r.w=l.matrixTransform(a),l.x+=i,r.e=l.matrixTransform(a),l.x-=i/2,l.y-=c/2,r.n=l.matrixTransform(a),l.y+=c,r.s=l.matrixTransform(a),r}function h(t){return"function"==typeof t?t:function(){return t}}return c}t.getTextWidth=n,t.getUniqueID=r,t.palettes=a,t.defaultPalette=o,t.getPalette=i,t.listPalettes=function(){return Object.keys(a)},t.scaleOrdinal=l,t.output=function(){var t="output",e=function(){};function n(t){return"svg"==t?document.querySelector("svg"):document.getElementsByTagName("svg")[t]}function r(e,n){return(e=e||t)+"."+n}function a(t){t.setAttribute("xmlns","http://www.w3.org/2000/svg"),t.setAttribute("xmlns:xlink","http://www.w3.org/1999/xlink"),function(t,e){var n=document.createElement("style");n.setAttribute("type","text/css"),n.innerHTML=t;var r=e.hasChildNodes()?e.children[0]:null;e.insertBefore(n,r)}(function(t){var e=[];e.push("#"+t.id);for(let n=0;n<t.classList.length;n++)c("."+t.classList[n],e)||e.push("."+t.classList[n]);var n=t.getElementsByTagName("*");for(let t=0;t<n.length;t++){var r=n[t].id;c("#"+r,e)||e.push("#"+r);var a=n[t].classList;for(let t=0;t<a.length;t++)c("."+a[t],e)||e.push("."+a[t])}var o="";for(let t=0;t<document.styleSheets.length;t++){var i=document.styleSheets[t];try{if(!i.cssRules)continue}catch(t){if("SecurityError"!==t.name)throw t;continue}for(var l=i.cssRules,s=0;s<l.length;s++)c(l[s].selectorText,e)&&(o+=l[s].cssText)}return o;function c(t,e){return-1!==e.indexOf(t)}}(t),t);var e=(new XMLSerializer).serializeToString(t);return e=(e=e.replace(/(\w+)?:?xlink=/g,"xmlns:xlink=")).replace(/NS\d+:href/g,"xlink:href")}e.toSVG=function(t,e){var i=n(e=e||"svg");t=r(t,"svg");var l=a(i),s=new Blob([l],{type:"image/svg+xml"});o(s,t)},e.toPNG=function(t,e){var i=n(e=e||"svg");t=r(t,"png");var l=+i.getAttribute("width"),s=+i.getAttribute("height"),c=a(i),u="data:image/svg+xml;base64,"+btoa(unescape(encodeURIComponent(c))),p=document.createElement("canvas");p.width=l,p.height=s;var d=p.getContext("2d"),f=new Image;f.src=u,f.onload=function(){d.clearRect(0,0,l,s),d.drawImage(f,0,0,l,s),p.toBlob(function(e){o(e,t)})}};var o=o||function(t){if(!(void 0===t||"undefined"!=typeof navigator&&/MSIE [1-9]\./.test(navigator.userAgent))){var e=t.document,n=function(){return t.URL||t.webkitURL||t},r=e.createElementNS("http://www.w3.org/1999/xhtml","a"),a="download"in r,o=/constructor/i.test(t.HTMLElement)||t.safari,i=/CriOS\/[\d]+/.test(navigator.userAgent),l=t.setImmediate||t.setTimeout,s=function(t){l(function(){throw t},0)},c=function(t){setTimeout(function(){"string"==typeof t?n().revokeObjectURL(t):t.remove()},4e4)},u=function(t){return/^\s*(?:text\/\S*|application\/xml|\S*\/\S*\+xml)\s*;.*charset\s*=\s*utf-8/i.test(t.type)?new Blob([String.fromCharCode(65279),t],{type:t.type}):t},p=function(e,p,d){d||(e=u(e));var f,g=this,h="application/octet-stream"===e.type,m=function(){!function(t,e,n){for(var r=(e=[].concat(e)).length;r--;){var a=t["on"+e[r]];if("function"==typeof a)try{a.call(t,n||t)}catch(t){s(t)}}}(g,"writestart progress write writeend".split(" "))};if(g.readyState=g.INIT,a)return f=n().createObjectURL(e),void l(function(){var t,e;r.href=f,r.download=p,t=r,e=new MouseEvent("click"),t.dispatchEvent(e),m(),c(f),g.readyState=g.DONE},0);!function(){if((i||h&&o)&&t.FileReader){var r=new FileReader;return r.onloadend=function(){var e=i?r.result:r.result.replace(/^data:[^;]*;/,"data:attachment/file;");t.open(e,"_blank")||(t.location.href=e),e=void 0,g.readyState=g.DONE,m()},r.readAsDataURL(e),void(g.readyState=g.INIT)}f||(f=n().createObjectURL(e)),h?t.location.href=f:t.open(f,"_blank")||(t.location.href=f),g.readyState=g.DONE,m(),c(f)}()},d=p.prototype;return"undefined"!=typeof navigator&&navigator.msSaveOrOpenBlob?function(t,e,n){return e=e||t.name||"download",n||(t=u(t)),navigator.msSaveOrOpenBlob(t,e)}:(d.abort=function(){},d.readyState=d.INIT=0,d.WRITING=1,d.DONE=2,d.error=d.onwritestart=d.onprogress=d.onwrite=d.onabort=d.onerror=d.onwriteend=null,function(t,e,n){return new p(t,e||t.name||"download",n)})}}("undefined"!=typeof self&&self||"undefined"!=typeof window&&window||void 0);return e},t.legend=s,t.tooltip=c,t.Lollipop=function(t,a,o){const i="g3_lollipop",u={circle:0,pie:1},p={x:"AA_Position",y:"Protein_Change",factor:"Mutation_Class"},d={symbol:"hgnc_symbol",name:"protein_name",length:"length",domainType:"pfam",details:{start:"pfam_start",end:"pfam_end",ac:"pfam_ac",name:"pfam_id"}};o=o||800,(a=a||"circle")in u||(a="circle");var f,g,h,m,b,y,x,v,k,C,w,S,T,A,L,D,_,N,E,B,M,F,P,O,I,z,R,W,H,j,U,V,$,G,q,X=r(),Y={chartID:i+"_chart_"+X,className:"g3-chart",tooltip:!0,margin:{left:40,right:20,top:15,bottom:25},background:"transparent",transitionTime:600,highlightTextAngle:90,legend:!0,legendOpt:{margin:{},interactive:!0,title:"__undefined__"}},J={id:i+"-main-"+X,defsId:i+"-main-defs-"+X,xAxisDefsId:i+"-xAxis-defs-"+X,height:420,background:"rgb(233,233,233)",lollipopClassName:{group:"lollipop",line:"lollipopLine",pop:"pop"},lollipopLine:{stroke:"rgb(42,42,42)","stroke-width":.5},popClassName:{pie:"popPie",slice:"pieSlice",text:"popText",circle:"popCircle",label:"popLabel"},popCircle:{stroke:"wheat","stroke-width":.5},popText:{fill:"#EEE","font-family":"Sans","font-weight":"normal","text-anchor":"middle",dy:"0.35em"},popLabel:{"font-family":"Arial","font-weight":"normal",minFontSize:10,fontsizeToRadius:1.4,padding:5},popColorSchemeName:"accent",popColorScheme:l("accent"),yPaddingRatio:1.1,popParams:{rMin:2,rMax:12,addNumRCutoff:8,yLowerQuantile:.25,yUpperQuantile:.99},title:{id:"g3-chart-title",text:"",font:"normal 16px Arial",color:"#424242",alignment:"middle",dy:"0.35em"},axisLabel:{font:"normal 12px Arial",fill:"#4f4f4f","text-anchor":"middle",dy:"-2em",alignment:"middle"},ylab:{text:"# of mutations",lineColor:"#c4c8ca",lineWidth:1,lineStyle:"dash"}},Q={id:i+"-domain-"+X,defsId:i+"-annotation-defs-"+X,height:30,margin:{top:4,bottom:0},background:"transparent",className:{track:"domain_track",bar:"domain_bar",domain:i+"_domain",brush:"domain-x-brush",zoom:"main-viz-zoom"},domain:{colorSchemeName:"category10",colorScheme:l("category10"),margin:{top:0,bottom:0},label:{font:"normal 11px Arial",color:"#f2f2f2"}},bar:{background:"#e5e3e1",margin:{top:2,bottom:2}},brush:{enabled:!0,fill:"#666",opacity:.2,stroke:"#969696",strokeWidth:1,handler:"#666"},zoom:!0},K=[],Z={},tt=!1,et=[],nt=null,rt=!1,at=!1,ot={},it={},lt=function(t){return ct(t)-st(t)},st=function(t){return D(t[g.details.start])},ct=function(t){return D(t[g.details.end])},ut=function(){D.domain(L),Q.brush.enabled||(E.call(Ft),S.attr("x",function(t){return st(t)}).attr("width",function(t){return lt(t)}),T.attr("x",function(t){return(st(t)+ct(t))/2})),v.attr("x1",function(t){return dt(t)}).attr("x2",function(t){return dt(t)}),k.attr("transform",function(t){return"translate("+dt(t)+","+ft(t)+")"})},pt=function(){if(e.event.sourceEvent&&"zoom"===e.event.sourceEvent.type)return;let t=e.event.selection||R.range();L=t.map(R.invert),Q.zoom&&b.select(".main-viz-zoom").call(W.transform,e.zoomIdentity.scale($/(t[1]-t[0])).translate(-t[0],0)),ut()},dt=function(t){return D(t.position)},ft=function(t){return M(t._currentState.count)},gt=function(t){let e=t._currentState.count;return 0==e?0:e>=C?J.popParams.rMax:e<=w?J.popParams.rMin:(e-w)/(C-w)*(J.popParams.rMax-J.popParams.rMin)+J.popParams.rMin},ht=function(t){let n="";return at?(n+=`<div class="note">\n            <table class="pure-table pure-table-bordered">\n                <caption> Position ${t.position} </caption>\n                <thead>\n                    <tr>\n                        <th scope="col">Class</th>\n                        <th scope="col">#</th>\n                        <th scope="col">Change</th>\n                        <th scope="col">&#37;</th>\n                    </tr>\n                </thead>\n                <tbody>`,t._currentState.summary.forEach(function(r){r.value.count>0&&(n+=`\n                    <tr>\n                        <th scope="row"> ${r.key} </th>\n                        <td> ${r.value.count} </td>\n                        <td>`+mt(r.value.byY)+"</td>\n                        <td>"+e.format(".1f")(r.value.count/t._currentState.count*100)+"&#37;</td>\n                    </tr>")}),n+="\n                </tbody>\n            </table></div>"):(n+=`\n            <table class="pure-table pure-table-bordered">\n                <caption> Position ${t.position} </caption>\n                <thead>\n                    <tr>\n                        <th scope="col">Change</th>\n                        <th scope="col">&#37;</th>\n                    </tr>\n                </thead>\n                <tbody>`,t._currentState.summary[0].value.byY.forEach(function(r){n+=`\n                    <tr>\n                        <th> ${r.key} </th>\n                        <td>`+e.format(".1f")(r.value/t._currentState.count*100)+"&#37;</td>\n                    </tr>"}),n+="\n                </tbody>\n            </table>"),n},mt=function(t){var e="";return t.forEach(function(t){e+=`${t.key} (n=${t.value})<br>`}),e},bt=function(t,e){Y.tooltip&&nt.show(t,e)},yt=function(t){Y.tooltip&&nt.hide(t)},xt=function(t){let e,n,r,a=-1;return t._currentState.summary.forEach(function(t){t.value.count>0&&t.value.count>a&&(e=t.value.byY[0].key,r=t,n=J.popColorScheme(t.key),a=t.value.count)}),{class:r,entry:e,count:a,color:n}},vt=function(t,e){if(t._currentState.showLabel){t._currentState.showLabel=!1;let n=e.select("."+J.popClassName.label);Bt(n.attr("yMaxValue"),-1),n.transition().duration(Y.transitionTime).attr("transform","rotate(0)").remove()}else{t._currentState.showLabel=!0;let r=xt(t),a=Math.max(t._currentState.radius*J.popLabel.fontsizeToRadius,J.popLabel.minFontSize)+"px",o=J.popLabel["font-weight"]+" "+a+" "+J.popLabel["font-family"],i=(n(r.entry,o)+J.popLabel.padding+t._currentState.radius)*Math.sin(Y.highlightTextAngle/180*Math.PI)+t._currentState.radius*Math.cos(Y.highlightTextAngle/180*Math.PI),l=V*t._currentState.count/(V-i);Bt(l,1),e.append("text").classed(J.popClassName.label,!0).attr("x",-t._currentState.radius-J.popLabel.padding).attr("y",0).attr("yMaxValue",l).text(r.entry).style("fill",r.color).style("font",o).attr("dy",".35em").attr("text-anchor","end").transition().duration(Y.transitionTime).attr("transform","rotate("+Y.highlightTextAngle+")")}},kt=function(){return e.pie().sort(null)},Ct=function(t){let n=e.select(this);t._currentState.showLabel&&(n.select("."+J.popClassName.label).remove(),t._currentState.showLabel=!1);let r=t._currentState.radius=gt(t);"pie"==a&&function(t,n,r){let a=e.arc().innerRadius(0).outerRadius(r),o=kt().value(function(t){return t.value.count}),i=n.select("."+J.popClassName.pie).selectAll("path."+J.popClassName.slice).data(o(t._currentState.summary));i.enter().insert("path"),i.transition().duration(Y.transitionTime).attrTween("d",function(t){this._current=this._current||t;let n=e.interpolate(this._current,t);return this._current=n(0),function(t){return a(n(t))}}),i.exit().remove()}(t,n,r),function(t,e,n){let r=e.select("."+J.popClassName.circle),a=xt(t).color||"transparent";r.transition().duration(Y.transitionTime).attr("r",n).attr("fill",a)}(t,n,r);let o=n.select("."+J.popClassName.text);if(r>=J.popParams.addNumRCutoff){let e=wt(r,t._currentState.count,J.popText["font-weight"],J.popText["font-family"]),n=J.popText["font-weight"]+" "+e+" "+J.popText["font-family"];o.transition().duration(Y.transitionTime).text(t._currentState.count).style("font",n)}else o.text("")},wt=function(t,e,r,a){let o=1.75*t;for(let t=Math.ceil(o);t>=0;t--)if(n(e,r+" "+t+"px "+a)<=o)return t+"px";return 0},St=function(t){let n=e.select(this),r=t._currentState.radius=gt(t);n.append("circle").attr("class",J.popClassName.circle).attr("cx",0).attr("cy",0).attr("r",r).attr("stroke",J.popCircle.stroke).attr("stroke-width",J.popCircle["stroke-width"]).attr("fill","none");let a=e.arc().innerRadius(0).outerRadius(r),o=kt().value(function(t){return t.value.count}),i=n.append("g").attr("class",J.popClassName.pie);i.selectAll(J.popClassName.slice).data(o(t._currentState.summary)).enter().append("path").attr("d",a).attr("class",J.popClassName.slice).attr("fill",function(t){return J.popColorScheme(t.data.key)}),i.exit().remove();let l=n.append("text").attr("class",J.popClassName.text).attr("fill",J.popText.fill).style("font-family",J.popText["font-family"]).style("font-weight",J.popText["font-weight"]).attr("text-anchor",J.popText["text-anchor"]).attr("dy",J.popText.dy);if(r>=J.popParams.addNumRCutoff){let e=wt(r,t._currentState.count,J.popText["font-weight"],J.popText["font-family"]);l.transition().duration(Y.transitionTime).text(t._currentState.count).style("font-size",e)}else l.text("");n.on("mouseover",function(t){bt(t)}).on("mouseout",function(t){yt(t)}).on("click",function(t){vt(t,n)})},Tt=function(t){let n=e.select(this),r=t._currentState.radius=gt(t),a=xt(t);n.append("circle").attr("class",J.popClassName.circle).attr("cx",0).attr("cy",0).attr("r",r).attr("stroke",J.popCircle.stroke).attr("stroke-width",J.popCircle["stroke-width"]).attr("fill",a.color);let o=n.append("text").attr("class",J.popClassName.text).style("font-family",J.popText["font-family"]).style("font-weight",J.popText["font-weight"]).attr("text-anchor",J.popText["text-anchor"]).attr("dy",J.popText.dy);if(r>=J.popParams.addNumRCutoff){let e=wt(r,t._currentState.count,J.popText["font-weight"],J.popText["font-family"]);o.transition().duration(Y.transitionTime).text(t._currentState.count).attr("font-size",e)}else o.text("");n.on("mouseover",function(t){bt(t)}).on("mouseout",function(t){yt(t)}).on("click",function(t){vt(t,n)})},At=function(){switch(x=b.append("g").attr("class",J.lollipopClassName.group),v=x.append("g").attr("clip-path","url(#"+J.defsId+")").selectAll(J.lollipopClassName.line).data(K).enter().append("line").attr("x1",function(t){return dt(t)}).attr("x2",function(t){return dt(t)}).attr("y1",M(0)+Q.margin.top).attr("y2",function(t){return ft(t)}).attr("class",J.lollipopClassName.line).attr("stroke",J.lollipopLine.stroke).attr("stroke-width",J.lollipopLine["stroke-width"]),k=x.append("g").attr("clip-path","url(#"+J.defsId+")").selectAll(J.lollipopClassName.pop).data(K).enter().append("g").attr("transform",function(t){return"translate("+dt(t)+","+ft(t)+")"}).classed(J.lollipopClassName.pop,!0).attr("id",function(t,e){return i+"-"+X+"-lollipop-group-"+e}),a){case"pie":k.each(St);break;default:k.each(Tt)}let t,e,n;J.title.text.length>0&&("start"==J.title.alignment?(t="start",e=0):"middle"==J.title.alignment?(t="middle",e=$/2):(t="end",e=$),n=-Y.margin.top/2,b.append("text").attr("id",J.title.id).style("font",J.title.font).attr("fill",J.title.color).attr("text-anchor",t).attr("alignment-base","alphabetic").attr("transform","translate("+e+","+n+")").attr("dy",J.title.dy).text(J.title.text))},Lt=function(){if(e.event.sourceEvent&&"brush"===e.event.sourceEvent.type)return;let t=e.event.transform;L=t.rescaleX(R).domain(),Q.brush.enabled&&y.select("."+Q.className.brush).call(z.move,R.range().map(t.invertX,t)),ut()},Dt=function(){if(!f.hasOwnProperty("x")||!f.hasOwnProperty("y"))throw"No X or Y columns specified in data set";tt||(ot=e.nest().key(t=>t[f.factor]).rollup(function(t){return+t.length}).object(K),it=JSON.parse(JSON.stringify(ot)),(K=e.nest().key(t=>+t[f.x]).entries(K).sort((t,e)=>t.key-e.key)).forEach(function(t){t.position=+t.key,delete t.key,t.total=+t.values.length})),K.forEach(function(t){let n=e.nest().key(function(t){return t[f.factor]}).rollup(function(t){let n=t.filter(function(t){return t[f.factor]in it});return{count:+n.length,byY:e.nest().key(function(t){return t[f.y]}).rollup(function(t){return+t.length}).entries(n).sort(function(t,e){return e.value-t.value})}}).entries(t.values).sort(function(t,e){return t.key>e.key});tt||(t._currentState={},t._currentState.showLabel=!1),t._currentState.count=e.sum(n.map(function(t){return t.value.count})),t._currentState.summary=n}),tt=!0,P=K.map(function(t){return+t._currentState.count}).filter(function(t){return t>0}).sort(function(t,e){return t-e}),w=Math.ceil(e.quantile(P,J.popParams.yLowerQuantile)),C=Math.floor(e.quantile(P,J.popParams.yUpperQuantile)),et=[]},_t=function(t){return e.scaleLinear().domain([0,t]).nice().domain()[1]},Nt=function(){(B=0==P.length?[0,1]:e.extent(P))[0]=Math.min(0,B[0]),A=B[1]=_t(B[1]*(J.yPaddingRatio+J.popParams.rMax/V))},Et=function(){Nt(),M=e.scaleLinear().domain(B).range([V,0]),F=e.axisLeft(M).tickSize(-$).ticks(6).tickFormat(e.format("d")),L=[0,Z[g.length]],D=e.scaleLinear().domain(L).range([0,U]),R=e.scaleLinear().domain(L).range([0,U]),N=function(t,e){if(void 0===e||t.length<=1||t[t.length-1]>=e)return t;var n=Math.abs(t[1]-t[0]);return Math.abs(e-t[t.length-1])>n/2?t.push(e):t[t.length-1]=e,t}(D.ticks(),Z[g.length]),_=e.axisBottom(D).tickValues(N).tickFormat(function(t){return this.parentNode.nextSibling?t:t+" aa"})},Bt=function(t,e){if(1!=(e=e||1)&&-1!=e&&(e=1),t=_t(t*J.yPaddingRatio),t=Math.max(A,t),1===e)et.push(t),et=et.sort(function(t,e){return t-e});else{let e=et.indexOf(t);-1!=e&&(et=et.slice(0,e).concat(et.slice(e+1)))}let n=0==et.length?A:et[et.length-1];n!=B[1]&&(B[1]=n,Mt())},Mt=function(){M.domain(B),O.transition().duration(Y.transitionTime).call(Pt),v.transition().duration(Y.transitionTime).attr("y2",function(t){return ft(t)}),k.transition().duration(Y.transitionTime).attr("transform",function(t){return"translate("+dt(t)+","+ft(t)+")"})},Ft=function(t){var e=t.selection?t.selection():t;t.call(_),e.select(".domain").remove(),e.selectAll(".tick line").attr("stroke","#c4c8ca").attr("stroke-width",1)},Pt=function(t){var e=t.selection?t.selection():t;t.call(F),e.select(".domain").remove();let n=e.selectAll(".tick line").attr("stroke",J.ylab.lineColor).attr("stroke-width",J.ylab.lineWidth);"dash"==J.ylab.lineStyle&&n.attr("stroke-dasharray","3,3"),e.selectAll(".tick text").attr("x",-2).attr("dy",2),e!==t&&t.selectAll(".tick text").attrTween("x",null).attrTween("dy",null)},Ot=function(t,e,n,r){t.append("g").attr("id","background-layer").append("rect").attr("x",0).attr("y",0).attr("width",r).attr("height",n).attr("fill",e)},It=function(){t||(t="g3_"+X,e.select("body").append("div").attr("id",t)),(h=e.select("#"+t).append("svg")).attr("width",q).attr("height",G).attr("xmlns","http://www.w3.org/2000/svg").attr("xmlns:xlink","http://www.w3.org/1999/xlink"),h.classed(Y.className,!0).attr("id",Y.chartID).style("background-color",Y.background||"transparent"),m=h.append("g").attr("transform","translate("+Y.margin.left+","+Y.margin.top+")"),b=m.append("g").attr("id",J.id),Ot(b,J.background,V,$),y=m.append("g").attr("id",Q.id).attr("transform","translate(0, "+(V+Q.margin.top)+")"),Ot(y,Q.background,j,U),Y.tooltip&&(nt=c().attr("class","d3-tip").offset([8,0]).direction(function(t){return t.count>B[1]/2?"s":"n"}).offset(function(t){return t.count>B[1]/2?[16,0]:[-12,0]}).html(ht),b.call(nt))},zt={};return zt.options={set chartTarget(e){t=e},get chartTarget(){return t},set chartWidth(t){o=t},get chartWidth(){return o},set chartType(t){t&&t in u&&(a=t)},get chartType(){return a},get chartID(){return Y.chartID},get chartHeight(){return+h.attr("height")},set chartMargin(t){Y.margin=t},get chartMargin(){return Y.margin},set chartBackground(t){Y.background=t},get chartBackground(){return Y.background},set transitionTime(t){Y.transitionTime=t},get transitionTime(){return Y.transitionTime},set highlightTextAngle(t){Y.highlightTextAngle=t},get highlightTextAngle(){return Y.highlightTextAngle},set tooltip(t){Y.tooltip=t},get tooltip(){return Y.tooltip},set yAxisLabel(t){J.ylab.text=t},get yAxisLabel(){return J.ylab.text},set yAxisLineColor(t){J.ylab.lineColor=t},get yAxisLineColor(){return J.ylab.lineColor},set yAxisLineWidth(t){J.ylab.lineWidth=t},get yAxisLineWidth(){return J.ylab.lineWidth},set yAxisLineStyle(t){J.ylab.lineStyle=t},get yAxisLineStyle(){return J.ylab.lineStyle},set yMaxRangeRatio(t){J.yPaddingRatio=t},get yMaxRangeRatio(){return J.yPaddingRatio},set axisLabelFont(t){J.axisLabel.font=t},get axisLabelFont(){return J.axisLabel.font},set axisLabelColor(t){J.axisLabel.fill=t},get axisLabelColor(){return J.axisLabel.fill},set axisLabelAlignment(t){J.axisLabel.alignment=t},get axisLabelAlignment(){return J.axisLabel.alignment},set axisLabelDy(t){J.axisLabel.dy=t},get axisLabelDy(){return J.axisLabel.dy},set legend(t){Y.legend=t},get legend(){return Y.legend},set legendMargin(t){Y.legendOpt.margin=t},get legendMargin(){return Y.legendOpt.margin},set legendInteractive(t){Y.legendOpt.interactive=t},get legendInteractive(){return Y.legendOpt.interactive},set legendTitle(t){Y.legendOpt.title=t},get legendTitle(){return Y.legendOpt.title},get legendHeight(){return H},get lollipopTrackID(){return J.id},set lollipopTrackHeight(t){J.height=t},get lollipopTrackHeight(){return J.height},set lollipopTrackBackground(t){J.background=t},get lollipopTrackBackground(){return J.background},set lollipopPopMinSize(t){J.popParams.rMin=t},get lollipopPopMinSize(){return J.popParams.rMin},set lollipopPopMaxSize(t){J.popParams.rMax=t},get lollipopPopMaxSize(){return J.popParams.rMax},set lollipopPopInfoLimit(t){J.popParams.addNumRCutoff=t},get lollipopPopInfoLimit(){return J.popParams.addNumRCutoff},set lollipopPopInfoColor(t){J.popText.fill=t},get lollipopPopInfoColor(){return J.popText.fill},set lollipopPopInfoDy(t){J.popText.dy=t},get lollipopPopInfoDy(){return J.popText.dy},set lollipopLineColor(t){J.lollipopLine.stroke=t},get lollipopLineColor(){return J.lollipopLine.stroke},set lollipopLineWidth(t){J.lollipopLine["stroke-width"]=t},get lollipopLineWidth(){return J.lollipopLine["stroke-width"]},set lollipopCircleColor(t){J.popCircle.stroke=t},get lollipopCircleColor(){return J.popCircle.stroke},set lollipopCircleWidth(t){J.popCircle["stroke-width"]=t},get lollipopCircleWidth(){return J.popCircle["stroke-width"]},set lollipopLabelRatio(t){J.popLabel.fontsizeToRadius=t},get lollipopLabelRatio(){return J.popLabel.fontsizeToRadius},set lollipopLabelMinFontSize(t){J.popLabel.minFontSize=t},get lollipopLabelMinFontSize(){return J.popLabel.minFontSize},set lollipopColorScheme(t){J.popColorSchemeName=t,J.popColorScheme=l(t)},get lollipopColorScheme(){return J.popColorSchemeName},set titleText(t){J.title.text=t},get titleText(){return J.title.text},set titleFont(t){J.title.font=t},get titleFont(){return J.title.font},set titleColor(t){J.title.color=t},get titleColor(){return J.title.color},set titleAlignment(t){J.title.alignment=t},get titleAlignment(){return J.title.alignment},set titleDy(t){J.title.dy=t},get titleDy(){return J.title.dy},get annoID(){return Q.id},set annoHeight(t){Q.height=t},get annoHeight(){return Q.height},set annoMargin(t){Q.margin=t},get annoMargin(){return Q.margin},set annoBackground(t){Q.background=t},get annoBackground(){return Q.background},set annoBarFill(t){Q.bar.background=t},get annoBarFill(){return Q.bar.background},set annoBarMargin(t){Q.bar.margin=t},get annoBarMargin(){return Q.bar.margin},set domainColorScheme(t){Q.domain.colorSchemeName=t,Q.domain.colorScheme=l(t)},get domainColorScheme(){return Q.domain.colorSchemeName},set domainMargin(t){Q.domain.margin=t},get domainMargin(){return Q.domain.margin},set domainTextFont(t){Q.domain.label.font=t},get domainTextFont(){return Q.domain.label.font},set domainTextColor(t){Q.domain.label.color=t},get domainTextColor(){return Q.domain.label.color},set brush(t){Q.brush.enabled=t},get brush(){return Q.brush.enabled},set brushBackground(t){Q.brush.fill=t},get brushBackground(){return Q.brush.fill},set brushOpacity(t){Q.brush.opacity=t},get brushOpacity(){return Q.brush.opacity},set brushBorderColor(t){Q.brush.stroke=t},get brushBorderColor(){return Q.brush.stroke},set brushBorderWidth(t){Q.brush.strokeWdith=t},get brushBorderWidth(){return Q.brush.strokeWidth},set brushHandler(t){Q.brush.handler=t},get brushHandler(){return Q.brush.handler},set zoom(t){Q.zoom=t},get zoom(){return Q.zoom}},zt.setOptions=function(t){for(let e in t)this.options[e]=t[e]},zt.destroy=function(){h.selectAll("*").remove(),h.remove(),Y.tooltip&&rt&&nt.destroy()},zt.refresh=function(){this.destroy(),it=JSON.parse(JSON.stringify(ot)),this.draw()},zt.data={set snvData(t){K=t},get snvData(){return K},set domainData(t){Z=t},get domainData(){return Z}},zt.format={set snvData(t){f=t},get snvData(){return f},set domainData(t){g=t},get domainData(){return g}},zt.draw=function(){g=g||d,(f=f||p).hasOwnProperty("factor")?at=!0:(at=!1,zt.format.snvData.factor=f.factor="__undefined__",K.forEach(function(t){t[f.factor]="__undefined__"})),G=J.height+Q.height,q=o,V=J.height-Y.margin.top-Y.margin.bottom,$=q-Y.margin.left-Y.margin.right,j=Q.height-Q.margin.top-Q.margin.bottom,U=$,Dt(),It(),function(){let t,e;Et(),O=b.append("g").attr("class","axis axis--y").call(Pt),"start"==J.axisLabel.alignment?(t="end",e=0):"middle"==J.axisLabel.alignment?(t="middle",e=V/2):(t="start",e=V),b.append("text").attr("class","yaxis axis-label").attr("transform","translate(0,"+e+")rotate(-90)").style("font",J.axisLabel.font).attr("fill",J.axisLabel.fill).attr("text-anchor",t).attr("alignment-base","middle").attr("dy",J.axisLabel.dy).text(J.ylab.text),E=b.append("g").attr("clip-path","url(#"+J.xAxisDefsId+")").attr("class","axis axis--x").attr("transform","translate(0, "+(V+Q.height)+")").call(Ft)}(),b.append("clipPath").attr("id",J.defsId).append("rect").attr("width",$).attr("height",V+Q.margin.top),b.append("clipPath").attr("id",J.xAxisDefsId).append("rect").attr("x",-5).attr("width",$+20).attr("height",30),y.append("clipPath").attr("id",Q.defsId).append("rect").attr("width",U).attr("height",j),function(){let t=j-Q.bar.margin.top-Q.bar.margin.bottom;y.append("rect").classed(Q.className.bar,!0).attr("x",0).attr("y",Q.bar.margin.top).attr("width",U).attr("height",t).attr("fill",Q.bar.background);let n=j-Q.domain.margin.top-Q.domain.margin.bottom,r=y.append("g").attr("class",Q.className.track).selectAll(Q.className.domain).data(Z[g.domainType]).enter().append("g").attr("clip-path","url(#"+Q.defsId+")").attr("class",Q.className.domain);if(S=r.append("rect").attr("x",function(t){return st(t)}).attr("y",Q.domain.margin.top).attr("height",n).attr("width",function(t){return lt(t)}).attr("fill",function(t){return Q.domain.colorScheme(t[g.details.name])}),T=r.append("text").attr("x",function(t){return(st(t)+ct(t))/2}).attr("y",Q.domain.margin.top+n/2).attr("dy","0.35em").attr("text-anchor","middle").attr("fill",Q.domain.label.color||"white").style("font",Q.domain.label.font).text(function(t){return t[g.details.name]}),Q.brush.enabled&&(z=e.brushX().extent([[0,0],[U,j]]).on("brush end",pt)),Q.zoom&&(W=e.zoom().scaleExtent([1,1/0]).translateExtent([[0,0],[$,V]]).extent([[0,0],[$,V]]).on("zoom",Lt)),Q.brush.enabled){var a=y.append("g").attr("class",Q.className.brush).call(z);a.selectAll("rect.selection").attr("fill",Q.brush.fill).attr("fill-opacity",Q.brush.opacity).attr("stroke",Q.brush.stroke).attr("stroke-width",Q.brush.strokeWidth),a.selectAll("rect.handle").attr("fill",Q.brush.handler)}Q.zoom&&b.append("rect").attr("class",Q.className.zoom).attr("width",$).attr("height",V).attr("fill","none").attr("cursor","move").attr("pointer-events","all").call(W)}(),At(),Y.legend&&function(){if(Y.legend){I=new s(Y.chartID),0==Object.keys(Y.legendOpt.margin).length&&(Y.legendOpt.margin={left:Y.margin.left,right:Y.margin.right,top:2,bottom:2}),I.margin=Y.legendOpt.margin,I.interactive=Y.legendOpt.interactive,I.title="__undefined__"===Y.legendOpt.title?f.factor:Y.legendOpt.title;for(let t in it)I.addSeries({key:t,value:{fill:J.popColorScheme(t),label:t+" ("+it[t]+")"}});I.dispatch.on("legendClick",function(t,e){e?it[t]=e:delete it[t],Dt(),Nt(),Mt(),k.each(Ct)}),I.draw(),H=I.height}}(),Q.brush.enabled&&y.select(".domain-x-brush").call(z.move,[0,U]),rt=!0},zt},Object.defineProperty(t,"__esModule",{value:!0})});
```
