## Supplementary material for "ClusTRace, a bioinformatic pipeline for analyzing clusters in virus phylogenies": SFile 1: g3Lollipop.js

```
HTMLWidgets.widget({
  name: 'g3Lollipop',
  type: 'output',
  factory: function (el) {
    return {
      renderValue: function (x) {
        //console.log(x);
        //console.log("root id = ", el.id);

        // root div
        var root_div = document.getElementById(el.id);

        // clear div (issue #2)
        while(root_div.firstChild){
          root_div.removeChild(root_div.firstChild);
        }

        // add lollipop contents
        var btnStyle = x.btnStyle;

        // if add button
        var btn_div, svg_btn, png_btn;

        if(btnStyle != null){
          btnStyle = btnStyle + "Btn";
        }

        if (x.pngButton || x.svgButton) {
          btn_div = document.createElement('div');
          btn_div.style.height = "40px";
          btn_div.style.width = "100%";
          //btn_div.className = "btn-group";

          if (x.svgButton) {
            svg_btn = document.createElement('BUTTON');
            svg_btn.innerHTML = 'save as SVG';
            svg_btn.id = 'save-as-svg';
            if(btnStyle != null){
              svg_btn.className = btnStyle;
            }

            btn_div.appendChild(svg_btn);
          }

          if (x.pngButton) {
            png_btn = document.createElement('BUTTON');
            png_btn.innerHTML = 'save as PNG';
            png_btn.id = 'save-as-png';
            if(btnStyle != null){
              png_btn.className = btnStyle;
            }

            btn_div.appendChild(png_btn);
          }

          root_div.appendChild(btn_div);
        }

        // lollipop div
        var main_div = document.createElement('div');
        var _id = g3.getUniqueID();
        main_div.id = "lollipop-container-" + _id;
        root_div.appendChild(main_div);

        var lollipop = g3.Lollipop(main_div.id);

        lollipop.data.snvData = x.snvData;
        lollipop.data.domainData = x.domainData;
        lollipop.format.snvData = x.snvDataFormat;
        lollipop.format.domainData = x.domainDataFormat;
        lollipop.setOptions(x.plotSettings);

        lollipop.draw();

        var lollipop_height = lollipop.options.chartHeight,
          lollipop_width = lollipop.options.chartWidth;

        //console.log("lollipop h = ", lollipop_height, "  w = ", lollipop_width);

        // main_div height and width
        main_div.style.height = lollipop_height + 'px';
        main_div.style.width = lollipop_width + 'px';

        // root div height and width
        var root_height = lollipop_height + 40;
        root_width = lollipop_width;
        root_div.style.height = root_height + 'px';
        root_div.style.width = root_width + 'px';

        // add button function
        var chartID = lollipop.options.chartID;
        //console.log("chart id = ", chartID);

        if (x.svgButton) {
          svg_btn.onclick = function (e) {
            g3.output().toSVG(x.outputFN, chartID);
          }
        }

        if (x.pngButton) {
          png_btn.onclick = function (e) {
            g3.output().toPNG(x.outputFN, chartID);
          }
        }

        el.chart = lollipop;
      },
      resize: function (width, height) {
        // TODO: code to re-render the widget with a new size
      },
    };
  }
});
```
